## Supplemental File 1 for "Lung lineage transcription factor NKX2-1 epigenetically resolves opposing cell fates in vivo": scATACandscRNAprocessing.R

```
##The following code was used for procssing scATAC-seq and scRNA-seq for figures from: Lung lineage transcription factor NKX2-1 epigenetically resolves opposing cell fates in vivo

###### This section is for Supplement Figure 1 #####
# This file is for processing 7wk old Adult lungs for scATAC-seq 
#preprocessing pipeline
#1.Sample preperation
#lungs were sorted for CD45+ cells, CD45- ICAM2+ cells, CD45- ICAM2- ECAD+ cells, and CD45- ICAM2- ECAD- cells
#these cells were then recombined in equal proportions prior to processing for scATAc-seq. 


#2.Sequencing information
#The sample was sequenced using Illumina hi-seq paired end 
#In the initial round of sequencing the sample had lower depth than desired so the sample was resequenced for further depth. 
#reads were then processed following cellranger-atac pipeline.


#3.Processing raw reads 
# the filtered_peak_bc_matrix.h5 with fragments.tsv.gz and fragments.tsv.gz. tbi was read into Signac version 0.1.6 and Seurat 3.1.4
# the data was then processed following https://satijalab.org/signac/articles/pbmc_vignette.html 
#cell types were then identified  

#4.Initial processing by Seurat and Signac for cell typing
library(Signac)
library(Seurat)
library(GenomeInfoDb)
library(EnsDb.Mmusculus.v79)
library(ggplot2)
library(dplyr)
library(harmony)
library(JASPAR2018)
library(TFBSTools)
library(BSgenome.Mmusculus.UCSC.mm10)
set.seed(1234)


counts <- Read10X_h5("Y:/scAtac-seq_7wks/filtered_peak_bc_matrix.h5")
metadata <- read.csv(file = "Y:/scAtac-seq_7wks/singlecell.csv", header= TRUE, row.names = 1)
fragment.path <- 'Y:/scAtac-seq_7wks/fragments.tsv.gz'

brain <- CreateSeuratObject(
  counts = counts,
  assay = 'peaks',
  project = 'ATAC',
  min.cells = 1,
  meta.data = metadata)


brain <- SetFragments(
  object = brain,
  file = fragment.path
)

###################################################
#QC and preprocessing
###################################################
brain <- NucleosomeSignal(object = brain)

brain$pct_reads_in_peaks <- brain$peak_region_fragments / brain$passed_filters * 100
brain$blacklist_ratio <- brain$blacklist_region_fragments / brain$peak_region_fragments

VlnPlot(object = brain, features = c('pct_reads_in_peaks', 'blacklist_ratio', 'nucleosome_signal', 'peak_region_fragments')
        ,  pt.size = 0.1, ncol= 4) + NoLegend()

brain$nucleosome_group <- ifelse(brain$nucleosome_signal > 10, 'NS > 10', 'NS < 10')
FragmentHistogram(object = brain, group.by = 'nucleosome_group', region = 'chr1-10000000')

gene.ranges <- genes(EnsDb.Mmusculus.v79)
gene.ranges <- gene.ranges[gene.ranges$gene_biotype == 'protein_coding', ]

tss.ranges <- GRanges(
  seqnames = seqnames(gene.ranges),
  ranges = IRanges(start = start(gene.ranges), width = 2),
  strand = strand(gene.ranges)
)

seqlevelsStyle(tss.ranges) <- 'UCSC'
tss.ranges <- keepStandardChromosomes(tss.ranges, pruning.mode = 'coarse')

brain <- TSSEnrichment(object = brain, tss.positions = tss.ranges[1:2000])

brain$high.tss <- ifelse(brain$TSS.enrichment > 2, 'High', 'Low')

TSSPlot(brain, group.by = 'high.tss') + ggtitle("Tss enrichment Score") + NoLegend()

#before subsetting check what kind of numbers you are getting 
brain
#An object of class Seurat 
#135926 features across 11225 samples within 1 assay 
#Active assay: peaks (135926 features)

brain <- subset(brain, subset = peak_region_fragments > 1000 & peak_region_fragments < 100000 & pct_reads_in_peaks > 25 & blacklist_ratio < 0.025 & nucleosome_signal < 10 & TSS.enrichment > 2)

brain  # with peak_region_framents >1000
#135926 features across 9331 samples within 1 assay 
#Active assay: peaks (135926 features)
#2 dimensional reductions calculated: lsi, umap

brain <- RunTFIDF(brain)

brain <- FindTopFeatures(brain, min.cutoff = 'q0')

brain <- RunSVD( object = brain, assay = 'peaks', reduction.key = 'LSI_', reduction.name = 'lsi')

#there should be a correlations between the first LSi compoenet and sequencing depth 
#the first Lsi component often captures sequencing depth (technical variation) rathat than biological variation,
#if this is the case it should be removed from the downstream analysis 
FeatureScatter(brain, 'LSI_1', 'nCount_peaks')
#the correlation for it to be taken out in the example on the mouse_brain_vignette suggests .93 is a correlation enough that it should be taken out
#this would be accomplished by using dimensions 2:30 instead of 1:30 
#here this is 0.09 which is not a correlation so this first LSi component should not be taken out 
brain <- RunUMAP(object = brain, reduction = 'lsi', dims = 1:30)

brain <- FindNeighbors( object = brain, reduction = 'lsi', dims = 1:30)

#here the seurat tutorial uses reolution 1.2 check with both
brain <- FindClusters(object = brain, algorithm = 3, resolution = 1.2, verbose = FALSE)

#difference between 1 and 1.2 was 23 clusters vs 24 clusters 
DimPlot(object = brain, label = T) +NoLegend()
DimPlot(object = brain, label = F) 

DimPlot(object = brain, label = TRUE) + NoLegend()
table(brain$seurat_clusters)
#0    1    2    3    4    5    6    7    8    9   10   11   12   13   14   15   16   17 
#1215  533  525  525  521  510  482  450  430  428  347  341  317  316  281  265  257  256 
#18   19   20   21   22   23   24   25   26 
#248  232  191  154  150  101   99   89   68

gene.coords <- genes(EnsDb.Mmusculus.v79, filter = ~ gene_biotype == "protein_coding")
seqlevelsStyle(gene.coords) <- 'UCSC'
genebody.coords <- keepStandardChromosomes(gene.coords, pruning.mode = 'coarse')
genebodyandpromoter.coords <- Extend(x = gene.coords, upstream = 2000, downstream = 0)

# build a gene by cell matrix; this is slow
gene.activities <- FeatureMatrix( fragments = fragment.path,
                                  features = genebodyandpromoter.coords,cells = colnames(brain), chunk = 10)

# convert rownames from chromsomal coordinates into gene names
gene.key <- genebodyandpromoter.coords$gene_name
names(gene.key) <- GRangesToString(grange = genebodyandpromoter.coords)
rownames(gene.activities) <- make.unique(gene.key[rownames(gene.activities)])
gene.activities <- gene.activities[rownames(gene.activities)!="",]

#Add the gene activity matrix to the Seurat object as a new assay, and normalize it
brain[['RNA']] <- CreateAssayObject(counts = gene.activities)

brain <- NormalizeData(  object = brain, assay = 'RNA',
                         normalization.method = 'LogNormalize', scale.factor = median(brain$nCount_RNA))

DefaultAssay(brain) <- 'RNA'

FeaturePlot(object = brain, features = c('Spock2','Lamp3','Scgb1a1', "Foxj1"),
            pt.size = 0.1, max.cutoff = 'q95', ncol = 3, cols = c('lightgrey', 'red'))

FeaturePlot( object = brain, features = c('Spock2','Lamp3','Trp63', "Shh"), 
             pt.size = 0.1, max.cutoff = 'q95', ncol = 3, cols = c('lightgrey', 'red'))

DotPlot(brain, features = c("Fgf18","Pdgfra","Pdgfrb","Notch3","Meox2","Acta2","Actc1","Msln",
                            "Rtkn2","Lamp3","Sox9","Sox2","Foxj1","Scgb1a1","Trp63","Ascl1","Nkx2-1",
                            "Plvap","Car4","Gja5","Prox1",
                            "Cd79a","Cd9","Cd3e","Ccl4","Ear1","S100a9","Cd300e","Mycl","Xcr1","Irak2","Cd83","Runx1",
                            "Tbx4","Epcam","Cdh5","Ptprc"), group.by = "seurat_clusters") + coord_flip()

brain@meta.data$cell_type <- brain@meta.data$seurat_clusters
brain@meta.data$cell_type <- plyr::mapvalues(x = brain@meta.data$cell_type,
                                             from = c(0,1,2,3,4,5,6,7,8,9,10,11,12,13,14,15,16,17,18,19,20,21,22,23,24,25,26 ),
                                             to = c("AT2","AT1","Plvap","AT2","Sox2","Mes1","Mes2","Mes3","AT1","Imm1",
                                                    "Mes4","AT2","Mes4","Mes1","Mes3", "Mesothelial",
                                                    "Sox2","Mes3","Imm2","Imm3","Mes3","Mes3","Imm4","Car4","Sox2",
                                                    "Plvap","Lymphatics"))
Idents(brain) <- "cell_type"
DimPlot(brain, label = T)
brain <- subset(brain, idents=c("AT1", "AT2", "Plvap", "Sox2", "Car4","Mes1", "Mes2", "Mes3","Mes4","Mesothelial", "Imm1", "Imm2","Imm4"))

#Sup. Fig 2a
#cell types UMAP
DimPlot(brain, label = F)

#Dotplot
levels(brain) <- c("AT1", "AT2", "Airway", "Plvap", "Car4", "Dendritic_B", "T_NK", "Macrophage_Monocyte_Neutrophils", "AVSM", "Wnt5a", "Interstitial", "Pericytes", "Mesothelial")
DotPlot(brain, features = c("Spock2","Lamp3","Sox2", 
                            "Plvap", "Car4",
                            "Cd79a", "Cd3e", "Ccl4", "Cd9", "Ear1", "Cd300e", "S100a9",
                            "Acta2","Actc1","Cdh4","Fgf18","Pdgfra","Pdgfrb", "Notch3", "Msln",
                            "Nkx2-1" ,"Cdh5","Ptprc","Tbx4")) + coord_flip()

#rename clusters into lineages based on Nkx2-1, Cdh5, Ptprc, and Tbx4 accessibility
brain@meta.data$lineages <- brain@meta.data$cell_type
brain@meta.data$lineages <- plyr::mapvalues(x = brain@meta.data$lineage,
                                            from = c("AT2", "AT1", "Plvap", "Sox2", "Mes1", "Mes2", "Mes3", "Imm1","Mes4", "Mesothelial","Imm2","Imm4","Car4"),
                                            to = c("Epi","Epi","Endo","Epi","Mes","Mes","Mes","Imm","Mes","Mes",
                                                   "Imm","Imm","Endo"))
#Lineages UMAP
Idents(brain) <- "lineages"
DimPlot(brain, label = F, group.by = "lineages", cols = c('salmon','violet', 'turquoise3','darkorchid1'))

#the celltypes associated barcodes were then pulled label by alphabet per cell type and used by Sinto Version 0.5.1 to split the post processed cellranger-atac possorted_bam.bam indexed by possorted_bam.bam.bai
#These files are alphabetically ordering barcodes, so splitting the each file in half can result in production of pseudobulk replicates. 
#AT1
cells.use <- WhichCells(brain, ident = 'AT1')
cells.barcodes <- as.data.frame(cells.use)
cells.barcodes$type <- "A"
cells.AT1 <- as.data.frame(cells.barcodes)
write.table(cells.AT1, file = "AT1_scATAC_barcodes_CT.bed", row.names = F, col.names = F, quote = F, sep = "\t" )


#AT2
cells.use <- WhichCells(brain, ident = 'AT2')
cells.barcodes2 <- as.data.frame(cells.use)
cells.barcodes2$type <- "B"
cells.AT2 <- as.data.frame(cells.barcodes2)
write.table(cells.AT2, file = "AT2_scATAC_barcodes_CT.bed", row.names = F, col.names = F, quote = F, sep = "\t" )


#Sox2
cells.use <- WhichCells(brain, ident = 'Sox2')
cells.barcodes3 <- as.data.frame(cells.use)
cells.barcodes3$type <- "C"
cells.Sox2 <- as.data.frame(cells.barcodes3)
write.table(cells.Sox2, file = "Sox2_scATAC_barcodes_CT.bed", row.names = F, col.names = F, quote = F, sep = "\t" )


#Epithelial
cells.use <- WhichCells(brain, ident = 'Epi')
cells.barcodes4 <- as.data.frame(cells.use)
cells.barcodes4$type <- "D"
cells.Epi <- as.data.frame(cells.barcodes4)
write.table(cells.Epi, file = "Epi_scATAC_barcodes_Lin.bed", row.names = F, col.names = F, quote = F, sep = "\t" )


#Endothelial
cells.use <- WhichCells(brain, ident = 'Endo')
cells.barcodes5 <- as.data.frame(cells.use)
cells.barcodes5$type <- "E"
cells.Endo <- as.data.frame(cells.barcodes5)
write.table(cells.Endo, file = "Endo_scATAC_barcodes_Lin.bed", row.names = F, col.names = F, quote = F, sep = "\t" )


#Immune
cells.use <- WhichCells(brain, ident = 'Imm')
cells.barcodes6 <- as.data.frame(cells.use)
cells.barcodes6$type <- "F"
cells.Imm <- as.data.frame(cells.barcodes6)
write.table(cells.Imm, file = "Imm_scATAC_barcodes_Lin.bed", row.names = F, col.names = F, quote = F, sep = "\t" )


#Mesenchymal
cells.use <- WhichCells(brain, ident = 'Mes')
cells.barcodes7 <- as.data.frame(cells.use)
cells.barcodes7$type <- "I"
cells.Mes <- as.data.frame(cells.barcodes7)
write.table(cells.Mes, file = "Mes_scATAC_barcodes_Lin.bed", row.names = F, col.names = F, quote = F, sep = "\t" )


#For cell types and lineages full barcodes sets and 2 halfed barcode sets were uploaded to our high performance computing cluster for sinto, filtering, and peakcalling, 
#Sinto commands: sinto filterbarcodes -b possorted_bam.bam -c x_scATAC_barcodes_y.bed


#5.Filtering for MACs2 peak calling 
#split bams were then moved to indepedent folders per cell type and relabeled as scATAC.bam
#each filed was then filtered following the steps below via samtools 1.10 and picard 2.9.0:
#samtools sort -o scATAC_s.bam scATAC.bam
#java -Xmx8g -jar /rsrch3/scratch/pul_med-rsch/drlittle/picard-2.18.27/picard.jar MarkDuplicates I=scATAC_s.bam ASSUME_SORTED=true METRICS_FILE=scATAC_s.markdup.metrics VALIDATION_STRINGENCY=LENIENT OUTPUT=scATAC_s.md.bam
#samtools index scATAC_s.md.bam
#samtools view -b -h -F 4 -F 1024 -F 2048 -q 30 scATAC_s.md.bam > scATAC_sf.md.bam

#6. MACs2 peak calling per cell type
#samples were then processed using MACS2 commands for broadpeak calling: macs2 callpeak -t sampleID -g mm -q 0.05 --nomodel --shift -100 --extsize 200 -n sampleID -B --broad -f BAM
#called peaks were then filetered by the mm10 blacklist using bedtools: bedtools subtract -a sampleID.bt.macs2_peaks.broadPeak -b mm10.blacklist.bed > sampleID_Blfilt.bt.mac2s_peaks.broadPeak
#To check frip bedtools was used: bedtools sort -i sampleID.bt.macs2_peaks.broadPeak | bedtools merge -i stdin | bedtools intersect -u -a scATAC_sf.md.bam -b stdin -ubam | samtools view -c
#Due to the low number of reads a -log10 Q cuttoff of 2 was used prior to differential accessibility analysis.

#Differential accessibility analysis was carried out as:

#1. pseudobulk Airway 1 and pseudobulk Airway 2 both with Airway sites vs pseudobulk AT1 1 and pseudobulk AT1 2 both with AT1 sites and pseudobulk AT2 1 and pseudobulk AT2 2 both with AT2 sites
#2. pseudobulk Epi 1 and pseudobulk Epi 2 both with Epi sites vs pseudobulk Endo 1 and pseudobulk Endo 2 both with Endo sites, pseudobulk Imm 1 and pseudobulk Imm 2 both with Imm sites, and pseudobulk Mes 1 and pseudobulk Mes 2 both with Mes sites


###### This section is for Figure 4 and Supplement Figure 4 and 5 #####
#Samples for E14.5, E16.5, E18.5, P4, P6, P7, P8, P10, P15, P20, 10wk, and 15wk are processed here

library(Seurat)
library(cowplot)
library(dplyr)
library(ggplot2)
library(sctransform)
library(monocle)
library(VGAM)
library(grid)
library(EnhancedVolcano)
library(stringr)
library(MAST)

#load the 10x dataset and initialize the Seurat object:
samples <- read.csv(file.path("/Users/drlittle/Desktop/Agg12controls_8-6-20/Agg12_8-6-20filtered_feature_bc_matrix/", "aggregation_csv.csv"), stringsAsFactors=F)
#Load the 10x dataset and initialize the Seurat object:
data <- Read10X("/Users/drlittle/Desktop/Agg12controls_8-6-20/Agg12_8-6-20filtered_feature_bc_matrix/mm10/")
cells <- new("seurat", raw.data=cells.data)

integrate <- CreateSeuratObject(data, min.cells = 3, min.features = 200) %>% PercentageFeatureSet(pattern = "^mt-", col.name = "percent.mt") %>% NormalizeData() %>% FindVariableFeatures() %>% ScaleData()
integrate <- subset(integrate, subset = nFeature_RNA > 200 & nFeature_RNA < 4000 & percent.mt < 10)
integrate <- ScaleData(integrate) %>% RunPCA(npcs = 30) %>% RunUMAP(dims=1:30) %>% FindNeighbors() %>% FindClusters()

#label each of the datasets
integrate$ID <- str_sub(rownames(integrate@meta.data),-2,-1)
integrate@meta.data$ID2 <- plyr::mapvalues(x= integrate@meta.data$ID,
                                           from=c("-1","-2", "-3", "-4", "-5", "-6", "-7", "-8","-9","10", "11", "12"),
                                           to= c("E145","E165", "E185", "P4", "P6", "P7", "P8","P10", "P15", "P20", "10wk", "15wk"))
Idents(object = integrate) <- "ID"
DimPlot(integrate, label = T)

#identify where epithelial cells types are
FeaturePlot(integrate, features = c("Sox9","Spock2","Lamp3","Foxj1", "Scgb1a1", "Ascl1", "Trp63"))
#identify lineage clusters and proliferating clusters
FeaturePlot(integrate, features = c("Cdh5","Cdh1","Mki67","Ptprc","Col3a1"))


#add metadata for clusters into lineages
integrate@meta.data$lineage <- integrate@meta.data$seurat_clusters
integrate@meta.data$lineage <- plyr::mapvalues(x= integrate@meta.data$lineage,
                                               from=c("0","1","2","3","4","5","6","7","8","9","10","11","12","13","14", "15","16","17","18","19","20","21","22","23","24","25","26","27","28","29","30","31","32","33","34","35","36","37","38","39","40","41","42"),
                                               to= c("Cdh1","Cdh5", "Col3a1", "Cdh5", "Cdh1", "Ptprc","Col3a1", "Cdh1", "Col3a1", "Ptprc", "Col3a1", "Ptprc", "Ptprc", "Cdh1", "Cdh5", "Cdh1", "Cdh1", "Ptprc", "Cdh5", "Col3a1","Col3a1", "Cdh1", "Cdh1", "Cdh5", "Cdh1", "Cdh1", 
                                                     "Ptprc", "Col3a1", "Ptprc", "Cdh1", "Col3a1","Col3a1", "Ptprc","Col3a1", "Ptprc", "Cdh1", "Cdh5", "Ptprc", "Cdh5", "Col3a1", "Col3a1", "Ptprc", "Ptprc"))
#confirm new labels
Idents(object = integrate) <- "lineage"

#save file
saveRDS(integrate, file = "/Users/drlittle/Desktop/Agg12_lineages.rds")
DimPlot(integrate, label = T, group.by = "lineage")
DimPlot(integrate, label = T, split.by = "ID")
#subset the epithelial cells 
integrate_epi <- subset(integrate, idents=c("Cdh1"))
#verify
DimPlot(integrate_epi, label = T)
DimPlot(integrate_epi, label = T, group.by = "ID")

#Recluster epithelial cells
integrate_epi <- FindVariableFeatures(integrate_epi)
integrate_epi <- ScaleData(integrate_epi) %>% RunPCA(npcs = 30) %>% RunUMAP(dims=1:30) %>% FindNeighbors() %>% FindClusters(resolution = 1)
Idents(object = integrate_epi) <- "seurat_clusters"
DimPlot(integrate_epi, label = T)

#recheck for doublets lineage genes that should not be present
FeaturePlot(integrate_epi, features = c("Cdh5","Cdh1","Mki67","Ptprc","Col3a1"))
#reidentify where epithelial cell types are
FeaturePlot(integrate_epi, features = c("Sox9","Spock2","Lamp3","Ascl1","Foxj1", "Scgb1a1", "Trp63", "Sox2"))


DimPlot(integrate_epi, label = T, group.by = "ID")

#save file
saveRDS(integrate_epi, file = "/Users/drlittle/Desktop/Agg12_Epi_1.rds")
integrate_epi <- readRDS( file = "/Users/drlittle/Desktop/Agg12_Epi_1.rds")
DimPlot(integrate_epi)
integrate_epi
#add metadata to rename clusters into cell types
integrate_epi@meta.data$cell_type <- integrate_epi@meta.data$seurat_clusters
integrate_epi@meta.data$cell_type <- plyr::mapvalues(x= integrate_epi@meta.data$cell_type,
                                                     from=c("0","1","2","3","4","5","6","7","8","9","10","11","12","13","14", "15","16","17","18", "19", "20", "21","22","23"),
                                                     to= c("AT1","AT2","AT2", "AT1", "AT2", "AT2", "AT2","Sox9", "AT2", "AT2", "AT1", "Sox9", "AT1", "Sox9", "Sox9", "Sox9", "AT1", "AT1", "Sox9", "Ciliated", "Prolif AT2","AT2", "Secretory", "Secretory"
                                                     ))

Idents(object = integrate_epi) <- "cell_type"
DimPlot(integrate_epi, label = T)

DimPlot(integrate_epi, label = F, group.by = "cell_type", cols = c('green4', 'darkmagenta', 'blue4', 'salmon', 'darkorchid1', 'goldenrod1'))

#subset to alveolar cells only
integrate_epi2 <- subset(integrate_epi, idents=c("AT1", "AT2", "Sox9", "Prolif AT2"))
#verify
DimPlot(integrate_epi2, label = T)
#recluster epithelial cells
integrate_epi2 <- ScaleData(integrate_epi2) %>% RunPCA(npcs = 30) %>% RunUMAP(dims=1:30) %>% FindNeighbors() %>% FindClusters(resolution = 1)
DimPlot(integrate_epi2, label = T)

#check where epithelial cell types are
FeaturePlot(integrate_epi2, features = c("Sox9","Spock2","Lamp3","Ascl1","Foxj1", "Scgb1a1", "Trp63", "Sox2"))

#renames clusters into cell_types
integrate_epi2@meta.data$cell_type <- integrate_epi2@meta.data$seurat_clusters
integrate_epi2@meta.data$cell_type <- plyr::mapvalues(x= integrate_epi2@meta.data$cell_type,
                                                      from=c("0","1","2","3","4","5","6","7","8","9","10","11","12","13","14", "15","16","17","18", "19", "20"),
                                                      to= c("AT2","AT1","AT2", "AT2", "AT2", "AT2", "AT2","Sox9", "AT2", "Sox9", "AT1", "AT1", "Sox2", "Sox2", "AT1", "AT1", "Prolif AT2", "Sox9", "Sox9", "AT1", "AT1"
                                                      ))
#verify
Idents(object = integrate_epi2) <- "cell_type"
DimPlot(integrate_epi2, label = T)  

#save file
saveRDS(integrate_epi2, file = "/Users/drlittle/Desktop/Agg12_Epi_2.rds")
integrate_epi2 <- readRDS(file = "/Users/drlittle/Desktop/Agg12_Epi_2.rds")

#subset to AT1, AT2, Sox9, and proliferating AT2
integrate_epi3 <- subset(integrate_epi2, idents=c("AT1", "AT2", "Sox9", "Prolif AT2"))
DimPlot(integrate_epi3, label = F)

#recluster AT1, AT2, Sox9, and proliferating AT2
integrate_epi3 <- ScaleData(integrate_epi3) %>% RunPCA(npcs = 30) %>% RunUMAP(dims=1:30) %>% FindNeighbors() %>% FindClusters(resolution = 1)
DimPlot(integrate_epi3 , label = T)
DimPlot(integrate_epi3 , label = T, group.by = "ID")

#check for doublets again with lineage expressed genes
FeaturePlot(integrate_epi3 , features = c("Cdh5","Cdh1","Mki67","Ptprc","Col3a1"))
#verify locations of epithelial cell types and absence of Sox2 cells
FeaturePlot(integrate_epi3 , features = c("Sox9","Spock2","Lamp3","Sox2"))

#rename clusters based for cell types
integrate_epi3@meta.data$cell_type <- integrate_epi3@meta.data$seurat_clusters
integrate_epi3@meta.data$cell_type <- plyr::mapvalues(x= integrate_epi3@meta.data$cell_type,
                                                      from=c("0","1","2","3","4","5","6","7","8","9","10","11","12","13","14", "15","16","17"),
                                                      to= c("AT2","AT2","AT2", "AT1", "AT1", "AT2","Sox9", "Sox9", "AT1", "AT2", "AT2", "AT1", "AT1","Prolif AT2", "Sox9", "Sox9", "AT1", "AT1"
                                                      ))
#verify
Idents(object = integrate_epi3) <- "cell_type"
DimPlot(integrate_epi3, label = T)

#Figure 4A
#Figure 4A UMAP
DimPlot(integrate_epi3, label = F, group.by = "cell_type", cols = c('darkmagenta', 'green4', 'blue4', 'darkorchid1'))

table(integrate_epi3$cell_type)
#  AT2        AT1       Sox9 Prolif AT2 
#10558       6786       3060        556 

#Figure 4A DotPlot
levels(integrate_epi3) <- c("AT1", "AT2", "Sox9" , "Prolif AT2")
DotPlot(integrate_epi3, features = c("Spock2", "Lamp3", "Sox9", "Mki67")) + coord_flip()

#save file
integrate_epi3 <- FindVariableFeatures(integrate_epi3)
saveRDS(integrate_epi3, file = "/Path/to/file/Agg12_Epi_3.rds")
integrate_epi3 <-readRDS(file = "/Path/to/file/Agg12_Epi_3.rds")

FeaturePlot(integrate_epi3, features = c("Spock2", "Lamp3", "Sox9", "Sox2", "Foxj1", "Scgb1a1"), cols = c('lightgrey', 'red'))

Idents(object = integrate_epi3) <- "cell_type"
levels(integrate_epi3) <- c("AT1", "AT2", "Sox9" , "Prolif AT2")
VlnPlot(integrate_epi3, features =c("Tead1"), cols= c('green4', 'darkmagenta', 'blue4', 'darkorchid1'))
VlnPlot(integrate_epi3, features =c("Tead4"), cols= c('green4', 'darkmagenta', 'blue4', 'darkorchid1'))
VlnPlot(integrate_epi3, features =c("Ctgf"), cols= c('green4', 'darkmagenta', 'blue4', 'darkorchid1'))
VlnPlot(integrate_epi3, features =c("Cyr61"), cols= c('green4', 'darkmagenta', 'blue4', 'darkorchid1'))
VlnPlot(integrate_epi3, features =c("Cebpa"), cols= c('green4', 'darkmagenta', 'blue4', 'darkorchid1'))
VlnPlot(integrate_epi3, features =c("Cebpd"), cols= c('green4', 'darkmagenta', 'blue4', 'darkorchid1'))


FeaturePlot(integrate_epi3, features = c("Tead1", "Tead2", "Tead3", 'Tead4'), cols = c('lightgrey', 'red'))
FeaturePlot(integrate_epi3, features = c("Ctgf", "Cyr61", "Cebpa", 'Cebpd'), cols = c('lightgrey', 'red'))


FeaturePlot(Epi, features = c("Cebpa","Cebpd","Elf5","Etv5","Tead1", "Tead2"))
FeaturePlot(Epi, features = c("Myrf","Batf","Ankrd1","Tead3","Tead4", "Sox9"))
FeaturePlot(Epi, features = c("Trp63","Ascl1","Robo2","Scgb3a2","Sox2", "Foxp3"))

#Plots for NKX2-1 binding changes over time from progenitors to mature AT1 and AT2 cells
FeaturePlot(integrate_epi3, features = c("Spock2", "Lamp3", "Sox9"),cols = c('lightgrey', 'red'))

FeaturePlot(integrate_epi3, features = c("Pdpn", "Hopx", "Sftpb"), cols = c('lightgrey', 'red'))


#subset to AT2, AT1 and prog #without prolifating AT2 cells 
integrate_epi4 <- subset(integrate_epi3, idents=c( "AT2", "Sox9", "AT1"))
#verify
DimPlot(integrate_epi4, label = T)

#save file
integate_epi4 <- FindVariableFeatures(integate_epi4)
saveRDS(integrate_epi4, file = "/Users/drlittle/Desktop/Agg12_Epi_4woprolif.rds")
integrate_epi4 <- readRDS( file = "/Users/drlittle/Desktop/Agg12_Epi_4woprolif.rds")


####add module score to seurat object for genes associated with increased NKX2-1 binding from progenitor to mature AT1 or AT2 cells and genes associated with retained NKX2-1 binding from progenitor to mature AT1 or AT2 cells
#As we can only add a score for genes deetected in scRNA-seq we want to export a list of all genes to cross refernce with ChIP-seq annotated genes
allgenesaccountedfor <- integrate_epi4@assays[["RNA"]]@counts@Dimnames[[1]]
write.csv(allgenesaccountedfor, file = "/Users/drlittle/Desktop/Agg12_allgenesdetected.csv")
genestocheck  <- as.data.frame(integrate_epi4@assays$RNA@var.features)
write.csv(genestocheck, file = "/Users/drlittle/Desktop/Agg12_variablegenes.csv")

DimPlot(integrate_epi4)

allgenesforanalysis <- c('Retnla',	'Lyz1',	'Saa3',	'Lcn2',	'Hbb-bs',	'Hba-a1',	'Dlk1',	'Scgb3a2',	'Hba-a2',	'Orm1',	'Hbb-bt',	'Ube2c',	'Lrg1',	'H19',	'Scgb1a1',	'Sprr1a',	'Cdkn1c',	'Cyp4b1',	'Cyp2b10',	'Orm2',	'Hopx',	'Chia1',	'Cym',	'Hspa1a',	'Ctla2a',	'Msln',	'Krtap17-1',	'Pf4',	'Cxcl1',	'Gm42418',	'Igfbp2',	'Cxcl5',	'Pclaf',	'Tm4sf1',	'Ly6i',	'Pon1',	'Meg3',	'Cks2',	'Mdk',	'Ppbp',	'Dmkn',	'Gm26870',	'Chil1',	'Areg',	'Clu',	'Atf3',	'Fabp5',	'H2afz',	'Tmem213',	'Hmgb2',	'8430408G22Rik',	'Tubb5',	'Cenpa',	'Cdc20',	'Mgp',	'S100a6',	'Igfbp6',	'Slc26a4',	'Ccnb1',	'Birc5',	'Lyz2',	'Slpi',	'Gp9',	'Hist1h2ap',	'Fgl1',	'Gadd45g',	'Hspa1b',	'Eln',	'Isg15',	'H2afx',	'Cks1b',	'Top2a',	'Cdk1',	'Apoe',	'Hp',	'Cenpf',	'Hspb1',	'Cd24a',	'F3',	'Cxcl2',	'Ass1',	'Krt19',	'Mt2',	'Ran',	'Ranbp1',	'Mif',	'Scd1',	'Rgcc',	'Ctgf',	'Clec1b',	'Gm12840',	'Lgals3',	'Cldn6',	'Rrm2',	'Nrn1',	'Pthlh',	'Pgam2',	'Hmgn2',	'Bc1',	'Pmp22',	'Stmn1',	'Npm1',	'Gpm6a',	'AY036118',	'Tnfrsf9',	'Myl7',	'Gm26825',	'Marcksl1',	'Ndnf',	'Nusap1',	'Pxdc1',	'Krtap28-13',	'Tppp3',	'Socs3',	'S100a14',	'Cldn10',	'Cyr61',	'C2cd4b',	'Gm20186',	'Ctsh',	'Cdca8',	'Fam13a',	'Ankrd1',	'Ptma',	'Trf',	'Ier3',	'Cbx3',	'Ifi27l2a',	'Cxcl15',	'Col18a1',	'Aqp1',	'Tuba1b',	'Apoc1',	'Phgr1',	'Jpt1',	'Hdc',	'Dut',	'Gkn2',	'Ager',	'Dkk1',	'Klf2',	'Bdnf',	'Tnni3',	'Sfn',	'Clic5',	'Cavin2',	'Pbk',	'Cav1',	'Ccl21a',	'Sftpa1',	'Cdca3',	'Phgdh',	'Mfap4',	'Plac1',	'Edn1',	'Il4i1',	'Rgs18',	'Hist1h2bc',	'Cebpd',	'Igfbp7',	'Mt1',	'Spink5',	'Tuba1a',	'Olfr1369-ps1',	'Prdx6',	'Cdk4',	'Cst8',	'Ccnb2',	'Abi3bp',	'Spc25',	'Hnrnpa1',	'Gkn3',	'Fos',	'Smc2',	'Junb',	'Mest',	'Mia',	'Cnn3',	'Cox6a2',	'Igfbp5',	'Tagln',	'Mirg',	'Col6a1',	'Gm47969',	'Gem',	'Spock2',	'Prmt1',	'Ccna2',	'Anp32b',	'Tyms',	'Fmod',	'Adamts1',	'Cyp2f2',	'Itih4',	'Cavin3',	'Krt7',	'Emp2',	'Cryab',	'Ung',	'Knstrn',	'Nupr1',	'Vim',	'Fbln5',	'Sox9',	'Akap5',	'Id1',	'Fosb',	'Cytl1',	'H2-Ab1',	'Ptgs2',	'Aqp5',	'Spc24',	'Alas2',	'Ly6e',	'Ccnd2',	'Bex1',	'Vegfa',	'Snca',	'Col6a2',	'Edn2',	'H1fx',	'Tubb2b',	'Sftpd',	'Il33',	'Cxcl10',	'Cd74',	'Trbc1',	'Mmrn1',	'Cxcl12',	'Hmgb3',	'Aurka',	'H2-Q1',	'Nme1',	'Sox11',	'Aurkb',	'Cbr2',	'Peg3',	'Rtkn2',	'Grb10',	'Gmnn',	'Pecam1',	'Rgs16',	'Sparc',	'Gm10260',	'Hmgb1',	'Cd14',	
                         'Sox2',	'Gm14964',	'Mki67',	'Gm26532',	'Col4a3',	'Plac8',	'Smoc2',	'Gapdh',	'Actb',	'Bex2',	'Cited2',	'Dusp1',	'Tk1',	'Hmgn1',	'Snrpd1',	'Mad2l1',	'Agl',	'Ifit3',	'Selenbp1',	'Tacstd2',	'Cdkn2b',	'B930036N10Rik',	'Hist1h1c',	'Egr1',	'Dnajb1',	'Plek',	'Olfr655',	'Set',	'Cxcl3',	'Ascl1',	'Serpine1',	'Thbs1',	'Foxp2',	'Prcd',	'Gp5',	'Pdpn',	'Gm1673',	'Pimreg',	'Tipin',	'S100g',	'Timp3',	'Nrgn',	'Gm33489',	'Lsm3',	'Cdca2',	'Cdkn1a',	'Anxa1',	'Nasp',	'Lig1',	'Fermt3',	'Steap4',	'Ifit1',	'Dctpp1',	'Hspe1',	'Col4a4',	'Rbp1',	'Col3a1',	'Jun',	'Hspd1',	'Tead2',	'Nbl1',	'Lsm2',	'Cenpe',	'Lmnb1',	'Gm26522',	'Ly6h',	'Plk1',	'Sec14l3',	'Gp1bb',	'Hmga2',	'Acta2',	'Krtap1-5',	'Tmem37',	'Tinagl1',	'Cd69',	'Krt8',	'Prc1',	'Igkc',	'Retnlg',	'Lockd',	'Ccdc34',	'Angptl8',	'Zfp36',	'Fkbp3',	'Mis18bp1',	'Ahnak',	'Aqp3',	'Ier2',	'Gins2',	'Tnnt2',	'Anxa3',	'6430710C18Rik',	'Fstl1',	'Ereg',	'Tpx2',	'Anxa5',	'Ahnak2',	'Napsa',	'Gata2',	'Cldn4',	'Xist',	'Id2',	'Gdpd2',	'Gm17669',	'Phox2a',	'Cd9',	'Psat1',	'Sftpb',	'Ifi47',	'Nap1l1',	'Fjx1',	'H2-Aa',	'Tcf21',	'H2afy2',	'Gpx2',	'Fxyd4',	'Inmt',	'Ybx1',	'Odc1',	'Gsn',	'Fbn2',	'Rapgef5',	'Nr4a1',	'Alox12',	'Serpinb9b',	'Ncl',	'Gng11',	'Iigp1',	'Ms4a4b',	'Dtl',	'Treml1',	'Lgals1',	'Hbegf',	'Nt5dc2',	'Gprc5a',	'Phlda1',	'H2-Eb1',	'Cdkn3',	'G0s2',	'Racgap1',	'Ptcra',	'Srgn',	'Gsta3',	'Sema3e',	'Hells',	'Atp6v0a4',	'Krt18',	'Psrc1',	'Cav2',	'Pcolce',	'Gp1ba',	'Krt83',	'Il23a',	'Trbc2',	'Pfn2',	'Slc6a4',	'Scnn1g',	'Gm15915',	'Selenoh',	'Ppia',	'Abhd2',	'Krt15',	'Kif23',	'Srm',	'Fbxo5',	'Uhrf1',	'Mthfd1',	'Il7r',	'Arg1',	'Dlk2',	'Slbp',	'Fbln1',	'Ptn',	'Tst',	'2210011C24Rik',	'Efcc1',	'Id3',	'Apex1',	'Gar1',	'Bin2',	'Cd177',	'H1f0',	'Eno1',	'Hmmr',	'Hoxd8',	'Col27a1',	'Nhp2',	'Hist3h2ba',	'Acot7',	'Hsp90aa1',	'Eef1g',	'Fscn1',	'Nop58',	'Rac2',	'Actc1',	'Plod2',	'Aldh1a1',	'Cfb',	'Myl4',	'Fgg',	'Acta1',	'Ifitm3',	'Nfkbia',	'AC140186.1',	'Ptpre',	'Ppa1',	'Tshb',	'Mid1ip1',	'Ccl20',	'Tcf19',	'Mfap2',	'Rad51',	'Flrt3',	'Selenom',	'3830403N18Rik',	'Frat2',	'Lyar',	'Wnt10b',	'Meis2',	'Rnd1',	'Sumo2',	'Smc4',	'Gnmt',	'Adm',	'Smpx',	'Glod5',	'Lyve1',	'Mcm6',	'Pdlim3',	'Ezh2',	'Gm32585',	'1700049E15Rik',	'Nnat',	'Srsf7',	'Il6',	'Lmo7',	'H2al1b',	'Tspan1',	'Cd247',
                         'Hmgcs2',	'Ccne1',	'Wfdc1',	'Btg2',	'Hck',	'Noa1',	'Srsf3',	'Ccne2',	'Timp2',	'Rpsa',	'Gsto1',	'Ldha',	'Clic3',	'Mcm5',	'Rpgrip1',	'Lypd2',	'B2m',	'Ptges3',	'Lin7a',	'Cenpm',	'Nmral1',	'Hsp90ab1',	'Pigr',	'Tspan8',	'Tspan3',	'Ube2s',	'Lama4',	'Nt5e',	'Gipc2',	'Klf4',	'Mok',	'Nppc',	'Dek',	'Igfbp4',	'Col9a3',	'Ccl11',	'Dpep1',	'Tnfrsf12a',	'Phyh',	'Mcm3',	'Cd28',	'Cdh13',	'Cdca7',	'Rpa2',	'B230312C02Rik',	'Kif22',	'Elovl1',	'Rps2',	'Rfc4',	'Ckap2l',	'Cldn5',	'Fam174b',	'Dnajc9',	'Tnfrsf23',	'Egr2',	'Itm2a',	'Hist1h4i',	'Tnni1',	'Snrpe',	'Psip1',	'Pkm',	'Nucks1',	'Cebpa',	'Trp63',	'Itgb6',	'Pmch',	'Soat2',	'Cd52',	'Lsm4',	'Tceanc',	'Dhfr',	'Ckb',	'Lims2',	'Bzw2',	'Zfp42',	'Cd300e',	'Eif5a',	'Cenpw',	'Serpina3g',	'Qpct',	'Syce2',	'Cavin4',	'Lhfp',	'Marcks',	'Uchl1',	'Fabp12',	'Stmn2',	'Asf1b',	'Dtymk',	'Tpm2',	'Lars2',	'Gja4',	'Timp4',	'Ehd2',	'2200002D01Rik',	'Hhip',	'Fgfbp1',	'Nudcd2',	'Elf5',	'Hnrnpab',	'Tacc3',	'Car3',	'Plin2',	'Rgs4',	'Cacybp',	'Pa2g4',	'Nudt21',	'Zeb2',	'Rfc5',	'Sema3a',	'Tpi1',	'Limch1',	'Pygl',	'Guca2a',	'Hc',	'Akr1c14',	'Kcne2',	'Cxcl13',	'Fst',	'Pxdn',	'Nkg7',	'Dpysl2',	'Cped1',	'Gm29480',	'Gstm5',	'Bambi',	'Cd93',	'Insig1',	'Rpl29',	'Prim1',	'Plscr2',	'Sox17',	'C1qbp',	'Fbl',	'Fam46a',	'Rasl11a',	'Lsm6',	'Scnn1b',	'Bub3',	'Mcm7',	'Cenpk',	'Scnn1a',	'Sln',	'Prdx4',	'Eif4a1',	'Lsm5',	'Npy',	'Csf3',	'Rcn3',	'Uchl3',	'Pmf1',	'Akap2',	'Mgarp',	'Vtcn1',	'Gm15987',	'Emp1',	'Myl9',	'Ly6c1',	'1500015O10Rik',	'Gclc',	'Mbd1',	'Ramp2',	'Orc6',	'Homer2',	'Pgc',	'Gadd45b',	'Piwil4',	'Icam1',	'Cthrc1',	'Fbp2',	'Reg3g',	'Rasd1',	'Pgam1',	'Gpihbp1',	'Cpne9',	'Gp6',	'Fbxo40',	'Hmgn3',	'Timm8a1',	'Parvb',	'C6',	'F2rl2',	'Npnt',	'Hat1',	'Aplnr',	'Pglyrp1',	'Myo1b',	'Rhof',	'Ccnd1',	'Kif20b',	'Agrn',	'Csrp2',	'Fen1',	'Esco2',	'Adrb2',	'Nebl',	'Ywhaq',	'Pim3',	'Klf6',	'Rasl11b',	'Snrpf',	'Tnfrsf19',	'Rpl14',	'Slc38a1',	'Car8',	'Klhl4',	'Cd163l1',	'Muc13',	'Cdsn',	'Pcdhb18',	'Ctsc',	'Fxyd6',	'Agpat4',	'Cct8',	'Ddc',	'Sgk1',	'Pitx2',	'Kif2c',	'Dcakd',	'Incenp',	'Pgk1',	'Jund',	'Lama3',	'Bcl2a1b',	'Rorb',	'Clspn',	'Alyref',	'Shcbp1',	'Basp1',	'Dnajc2',	'Tspan6',	'Gm17056',	'Nme4',	'Fosl1',	'Ssrp1',	'Rad51ap1',	'Cttnbp2',	'Cdca5',	'Snrpa1',	'Tubb2a',	'Zfp422',	'Dqx1',	'Wnt7b',	
                         'Kcnj13',	'2310007B03Rik',	'Galnt18',	'Batf',	'Fasn',	'Ddx39',	'Gm17224',	'Gm20652',	'Adh7',	'Gm19610',	'Nr5a2',	'8030474K03Rik',	'Pex5l',	'Trpc4',	'Fga',	'B230334C09Rik',	'Crybb1',	'Gm9946',	'Dscaml1',	'Chrnb4',	'Lrrc2',	'Slc4a1',	'Pcdhb8',	'Plb1',	'Ndufa4l2',	'Rgs2',	'Cxcl16',	'Serpinb6b',	'Cfi',	'Ppih',	'Mfge8',	'Pdlim2',	'Qk',	'Mmp23',	'Dapk2',	'Snrpd3',	'Id4',	'Serpine2',	'Iglc2',	'A330069E16Rik',	'Noxa1',	'Tspan2',	'Cenph',	'Bok',	'Ptx3',	'Rsad2',	'Slc34a2',	'Slfn9',	'Kpna2',	'Acoxl',	'Nek2',	'Chaf1b',	'Mis18a',	'Adam28',	'Col5a2',	'Cdc6',	'Fabp3',	'Ldoc1',	'Gpr84',	'Bora',	'Tmem243',	'Tpd52l1',	'Fam89a',	'Sgo1',	'Gstm1',	'Polr3k',	'Arg2',	'Mgst2',	'Ppef2',	'Fam213a',	'Msn',	'Ckap2',	'Col4a5',	'Aox3',	'Arc',	'G3bp1',	'Nefl',	'Tnf',	'Rbm3',	'Spon1',	'Fhl1',	'Nop56',	'Gm8797',	'Kif20a',	'Emg1',	'Tfdp1',	'Tmpo',	'Lox',	'Anp32e',	'Mex3a',	'Rgs1',	'Lbp',	'Gm32051',	'Cxcr6',	'Cbx1',	'Lyl1',	'Mxd3',	'Tgfb2',	'Impdh2',	'Pdgfa',	'Slc6a14',	'Serpinf1',	'Tmem140',	'Adamts18',	'Ifit1bl2',	'Nup37',	'Zfyve21',	'Cit',	'Cxcl17',	'Atad2',	'Supt16',	'Sap30',	'Ifitm1',	'Plk2',	'Wwtr1',	'Upk3bl',	'Nr4a2',	'Lsm7',	'Rpa3',	'Tnfrsf4',	'Kcnk2',	'Slco3a1',	'Usp1',	'Col1a2',	'Exosc8',	'Gucy1a1',	'Pebp1',	'Snrnp40',	'Ednrb',	'Procr',	'Adam33',	'Trat1',	'Mal',	'Ska1',	'Rhox5',	'Hes1',	'Ckap4',	'Emilin1',	'Sapcd2',	'Rpl4',	'Col9a1',	'Vnn1',	'Synpr',	'Erh',	'Serbp1',	'Npy2r',	'Ppil1',	'Hoxa5',	'Kcnj2',	'Abra',	'Cd83',	'Csf2',	'Cck',	'Hist1h1b',	'Ano1',	'Pcna',	'Alpk2',	'Slc22a19',	'Smpdl3a',	'Aldh1a7',	'Rps4x',	'Dlc1',	'Rbmxl1',	'Abca5',	'Postn',	'Stx11',	'Tmem54',	'Gpx3',	'Gipr',	'Lamc2',	'Tgm2',	'Ppp1r3b',	'Mpig6b',	'Fam136a',	'Reln',	'Pmvk',	'Cd302',	'Itga2b',	'Ckmt1',	'Gm11361',	'Wdr45',	'Bmp2',	'Nde1',	'Spata31d1a',	'Depdc1a',	'Skp2',	'Xlr4b',	'Prdx2',	'Ntm',	'Pde7a',	'Trim59',	'H2afv',	'Ctxn1',	'2410017I17Rik',	'Sfxn1',	'Cdh5',	'Hs2st1',	'Nppa',	'C1qa',	'Gm6213',	'Ppp1r16b',	'Pcdhb17',	'Rrm1',	'Pax6',	'Cst7',	'4930502E18Rik',	'Il13ra2',	'Gm26793',	'Slc17a8',	'Foxr1',	'Hoxb8',	'Sectm1a',	'Gm42477',	'Slc22a3',	'Cd226',	'Batf2',	'Gm26902',	'Agbl4',	'Defb1',	'AW551984',	'Gm35188',	'A730017C20Rik',	'Mcm2',	'Mthfd2',	'Cenps',	'Ddah1',	'Hoxb4',	'Chst12',	'Chst8',	'Npm3',	'Scgb1c1',	'Tubb6',	'Rplp0',
                         'Rps5',	'Phactr1',	'Ier5',	'Hnrnpc',	'Cep55',	'Cd36',	'Ppm1n',	'Eif4e',	'Nuf2',	'Col12a1',	'Srsf2',	'Gm609',	'Sphk1',	'Chaf1a',	'Rnf186',	'Dnph1',	'Crabp2',	'A2ml1',	'Wee1',	'Acer2',	'Ppp2r2b',	'Catsperz',	'Gm10457',	'1700061H18Rik',	'Lsm8',	'Ndst1',	'Slc39a2',	'Cttnbp2nl',	'Mycn',	'Fignl1',	'Cela1',	'Dctd',	'Rpl15',	'Il2rg',	'Hpf1',	'Sh2d1a',	'Hcar2',	'Oasl2',	'Ifi27',	'Sema3c',	'Nanos1',	'Sox21',	'Lypd1',	'Ppid',	'Slc24a5',	'Tspan15',	'Lamp3',	'Cavin1',	'Grem2',	'Rgs10',	'Anxa2',	'Mettl7a1',	'Hjurp',	'Mettl1',	'Trim16',	'Samhd1',	'B830012L14Rik',	'Sgo2a',	'Fam189a2',	'Kdr',	'Fdps',	'Ackr1',	'Gfi1b',	'4930441J16Rik',	'Gm43569',	'E330017L17Rik',	'Matn1',	'Gm43698',	'Gdf3',	'Gm45602',	'Gm15890',	'Kctd19',	'Pkd1l3',	'Ebf2',	'Nr2e3',	'Aqp9',	'Gm11195',	'Krt17',	'Nt5c1b',	'Tbx1',	'Scn7a',	'Eef1b2',	'Zdhhc3',	'Peg10',	'Jpt2',	'Asns',	'Npc2',	'Gm45805',	'Sh3d21',	'Dkc1',	'Popdc3',	'Pam',	'Wtip',	'Sox4',	'Hspg2',	'Auts2',	'Kif11',	'Bcl6b',	'Fgf20',	'Nedd4l',	'Mal2',	'Bspry',	'Mustn1',	'Gm19990',	'Nkain4',	'Ddit4',	'2610028H24Rik',	'Smarcc1',	'Amt',	'Arhgef26',	'Nono',	'Pla2g1b',	'Ces1d',	'Melk',	'Fam83d',	'Mfsd2a',	'Ndrg1',	'Psmd14',	'Thyn1',	'C3',	'Pdgfb',	'Csf1r',	'Lmo1',	'Mrpl42',	'Enah',	'Ndn',	'Rangrf',	'Tnfsf9',	'Trp53',	'2610318N02Rik',	'C330027C09Rik',	'Inpp4b',	'Bgn',	'Htra1',	'Txnip',	'Errfi1',	'Cdc26',	'Mcm4',	'Pla2g12a',	'Sdf2l1',	'Magoh',	'Rbbp7',	'Dag1',	'Egr4',	'Mirt1',	'Sema3b',	'Xlr3b',	'Snhg11',	'Aunip',	'Pbdc1',	'1500009L16Rik',	'Cdt1',	'Thbs3',	'Hhex',	'Rgs5',	'Abca1',	'Gm26802',	'Rpl13',	'Eif1ax',	'Plat',	'Snx5',	'Eef1e1',	'Cotl1',	'Mgll',	'Ddah2',	'Rps6',	'Rrp15',	'U2af1',	'Rfc3',	'Edn3',	'Acy3',	'Pcsk6',	'Tgm1',	'Myo1c',	'Slc39a4',	'Lpl',	'Gdf15',	'Egfl6',	'Cenpl',	'Tmsb10',	'Gramd2',	'Aldh3b1',	'Mad2l1bp',	'Ly6c2',	'Wfdc12',	'Soat1',	'Mycl',	'2810474O19Rik',	'Fam84a',	'Fibin',	'Polr2m',	'Calb2',	'Pitpnm2os2',	'Retn',	'Slc23a1',	'Pgrmc1',	'Ccnf',	'Nefm',	'Spink4',	'Gata1',	'Gm45542',	'43900',	'Ict1os',	'Gm29156',	'Aox4',	'Gm10489',	'Draxin',	'Svop',	'Cldnd2',	'Cngb1',	'Gm6980',	'Adarb2',	'Gm4814',	'Gm48610',	'Esr2',	'Krt5',	'1600014C23Rik',	'Ms4a7',	'Spry2',	'Myh7',	'Gtf2a2',	'Fxyd2',	'Lmcd1',	'Ubc',	'Rpf2',	'Cdca7l',	'Fmo1',	'Kcnq1ot1',	'AY512931',	'Ppp1r14b',	'AC191865.2',
                         'Cd3d',	'Rbp4',	'Wdr86',	'Hmgcs1',	'Map1b',	'Dscc1',	'Efna3',	'Sapcd1',	'Paqr7',	'Dcn',	'Bcas2',	'Mmp14',	'Fxyd3',	'F11r',	'Dbf4',	'Hist1h2ac',	'Rangap1',	'Tmod3',	'Fras1',	'Hif3a',	'Ankle1',	'Nrk',	'Arl4d',	'Rrs1',	'Slc39a1',	'Tmem108',	'Cdkn2c',	'Gm17182',	'Poc1a',	'Cdipt',	'Naa50',	'Col4a1',	'Tnip3',	'Polr2h',	'Gm10827',	'Snrpa',	'Gnb4',	'1700011H14Rik',	'Tcf3',	'Cckar',	'Egln1',	'Gjb2',	'B9d1',	'Epas1',	'Nup43',	'Epcam',	'Gas2l3',	'Tril',	'Fam46b',	'Gm3776',	'Gpc3',	'Nol4l',	'Crlf3',	'Ang',	'Glrx',	'Col2a1',	'Cdc25c',	'Flt1',	'Gm29065',	'Tgtp1',	'Mns1',	'Pard6b',	'Plagl1',	'Clps',	'Acot2',	'Rhob',	'Rab12',	'Pold2',	'Lrpap1',	'Dnah7a',	'Aspm',	'Stbd1',	'Cited1',	'Dcxr',	'Acp1',	'Dynll1',	'Ska2',	'Nqo1',	'Gcnt2',	'Ly6a',	'Rtp4',	'Rab25',	'Rps27a',	'Ssb',	'Hnrnpdl',	'Gm32200',	'Cox4i2',	'Tie1',	'Tuba1c',	'Ms4a6b',	'Bub1b',	'Ighm',	'Cftr',	'Cenpp',	'Fstl3',	'Cth',	'Cep78',	'Msx1',	'2410004I01Rik',	'Atp6v1g3',	'Knl1',	'Hist4h4',	'Pfn1',	'Mafb',	'Hpgd',	'Rnps1',	'Fzd2',	'Prmt8',	'Rbbp4',	'Rbp2',	'Anxa8',	'Paqr5',	'Ncapg2',	'Cd84',	'Sae1',	'Mmp11',	'Srsf1',	'Anapc15',	'Fam212a',	'Lrr1',	'Rfc1',	'Ppp1r3a',	'Cd200',	'Rab27b',	'Trip13',	'Dlgap5',	'P2rx7',	'Ltbp4',	'Ncapg',	'Osbpl6',	'S1pr3',	'Gm49077',	'1700097N02Rik',	'Nrm',	'Mbd3',	'Fam111a',	'Srxn1',	'Tcp1',	'Eif2s1',	'Mpped2',	'Lipa',	'Dbn1',	'Rnf26',	'Mrto4',	'Creb5',	'Cenpq',	'Abhd5',	'Ddr1',	'Dusp6',	'Cygb',	'Pdap1',	'Nipal1',	'Rab11fip1',	'Hnrnpa0',	'Lifr',	'Dio1',	'Asphd2',	'Gm44096',	'Gprc5d',	'1700011D18Rik',	'Gm20655',	'Gm12027',	'Gm30411',	'Twist1',	'Ptger3',	'Ermap',	'Gpr3',	'Ngfr',	'Apoh',	'1810049J17Rik',	'Lta',	'Fgf18',	'Lrp2',	'Suv39h2',	'Stip1',	'Celf4',	'Lamb1',	'Trim47',	'Txnl1',	'Nfkbiz',	'Espn',	'Cacna1s',	'C1qb',	'Dusp9',	'Snx29',	'Gstp3',	'Ccdc80',	'Hnrnpk',	'Ctdspl',	'Rcn1',	'Rfc2',	'9230117E06Rik',	'Tspan13',	'Ptges',	'Stab1',	'Ppp1r3c',	'St3gal6',	'Rasgrp2',	'Pif1',	'Nucb2',	'Rabgap1l',	'Serpinb9',	'Praf2',	'Lrrc26',	'Fn1',	'Pcbp4',	'Gpc6',	'Cdk6',	'Slamf6',	'Dnah12',	'Crygs',	'Plod1',	'Dck',	'Snu13',	'Cenpn',	'Gna14',	'Msmo1',	'Neil3',	'Mpp6',	'Cdo1',	'AW112010',	'Hoxb3os',	'Ly6k',	'Nfe2',	'Ndc80',	'Kif15',	'Rasgrf2',	'Ppp1r1b',	'Sumo3',	'Hmox1',	'Ptgds',	'E2f8',	'Hist2h4',	'Epgn',	'Lrrc31',	'Gm3550',	'Bpifb5',	'Psmc6',
                         'Car1',	'Rem2',	'Cited4',	'Capn2',	'Glul',	'B3gnt7',	'Rpl3',	'Hmgn5',	'E2f1',	'Parp1',	'H2-DMa',	'Ola1',	'Spag5',	'Thbd',	'Neat1',	'1200007C13Rik',	'Ube2e3',	'Rpl8',	'Ctnnbip1',	'Itgb3bp',	'Prkag3',	'Gemin6',	'Eva1a',	'Bcl11a',	'Cebpb',	'Cald1',	'44083',	'Orm3',	'Cyp2s1',	'Bhlhe40',	'Rusc2',	'Dnm3',	'2410018L13Rik',	'Wnt3a',	'Polr2d',	'B430010I23Rik',	'Aqp4',	'Tmod1',	'S100a3',	'Ptprn2',	'BC028528',	'Lrrc59',	'Brix1',	'Cfl1',	'Mindy2',	'Siva1',	'Cdkn1b',	'Mrpl11',	'Sorbs2',	'Cox6b2',	'H2-Q6',	'Cdk2ap1',	'Tomm40',	'Nsg1',	'Adh1',	'Gjc1',	'S100a8',	'Tubb4b',	'Fam198b',	'Eif2s3y',	'S100a9',	'Gm17171',	'Gm42743',	'Kbtbd12',	'Ly6f',	'Apol7a',	'Ano7',	'Gm13293',	'Dnaic1',	'Vwa3a',	'Fabp7',	'Gypa',	'4930533D04Rik',	'Plxdc1',	'Gm30893',	'1700054A03Rik',	'Cd3g',	'Tead1',	'Gnl3',	'Itga3',	'Klk13',	'Gif',	'Tpbg',	'Scrn1',	'1190002N15Rik',	'Flot1',	'Adam9',	'Tnfrsf21',	'Pdzrn3',	'6330403K07Rik',	'Fam92a',	'Krtap28-10',	'Apol11b',	'Enpp2',	'Dapl1',	'Samd9l',	'Mrpl18',	'Osgin1',	'Psma7',	'Acbd7',	'Mybl2',	'Isca1',	'Ska3',	'Syn3',	'Xlr3a',	'Cct3',	'Mme',	'Ube2t',	'Bmp4',	'Pla2g4c',	'Clns1a',	'Ppp1r15a',	'Eprn',	'Adamtsl4',	'Ecscr',	'Glrx3',	'She',	'Rgs17',	'Gcm1',	'Arxes2',	'Eddm3b',	'Thy1',	'Ifi209',	'Myh10',	'Wdr89',	'Dpt',	'Gm15241',	'Ifit3b',	'Wipf3',	'Plvap',	'Gimap6',	'Ivl',	'Bzw1',	'Scx',	'Fcer2a',	'Ly6d',	'4930523C07Rik',	'Six1',	'2310015K22Rik',	'Armcx2',	'Tex30',	'Icam2',	'Trp53bp2',	'Cdkl5',	'Mrc2',	'Gm26759',	'Idi1',	'Scara3',	'Psmc3ip',	'Cdca4',	'Gins1',	'Syp',	'Vpreb3',	'Gm47794',	'Ncapd2',	'Sycp1',	'Osr2',	'Uchl5',	'F2r',	'Cdh2',	'Serpinb1a',	'Hk2',	'Ugp2',	'Cdc45',	'Psmd7',	'Ccdc88b',	'Exo1',	'Plce1',	'Pdzd2',	'Gm9493',	'Cldn23',	'Traf4',	'Lpcat1',	'Lmo2',	'Zfp637',	'Myh14',	'Tmem119',	'Snx7',	'Ttk',	'4430402I18Rik',	'Hdac2',	'Ect2',	'Emcn',	'Ctsz',	'Mrc1',	'Lbhd2',	'St6galnac4',	'Cd164l2',	'Plxna1',	'Rassf9',	'Dram1',	'Etv4',	'Gsr',	'Topbp1',	'Gm29508',	'Aard',	'Wdr18',	'Kcnk3',	'Otud1',	'Pi4k2b',	'Ilf2',	'C1qc',	'Arhgef39',	'Rnaseh2b',	'Rhno1',	'2610524H06Rik',	'Krt23',	'Fblim1',	'Phlda3',	'Foxq1',	'Pkig',	'Usp18',	'Gm9856',	'Noxo1',	'Rps3',	'Tor3a',	'Dhrs7',	'Ddx39b',	'Gm32950',	'Ntng1',	'Grid2',	'Gm32913',	'Kcnj15',	'Brinp2',	'Itln1',	'Tgm5',	'Chgb',	'Gm5544',	'Gm43825',	'Epha5',	'Oas2',	'Gm43332',	'Cyp2e1',
                         'Tnni2',	'Rnase12',	'Casp12',	'Gm6657',	'Nrxn3',	'Mkx',	'Cd6',	'Gtpbp4',	'Rassf10',	'Col15a1',	'Slc12a3',	'Atf5',	'Tspan11',	'Nrp2',	'Lamb2',	'Ccdc58',	'H2-DMb1',	'Foxc1',	'P4ha1',	'Hirip3',	'BC018473',	'Fads3',	'Slc1a3',	'Gtf2e2',	'Vnn3',	'Slc44a2',	'Myc',	'Tgif1',	'6030408B16Rik',	'Gm13383',	'Ooep',	'Bcl2a1a',	'Ppp1r14a',	'Tpt1',	'Anln',	'Pim1',	'Nip7',	'Lrtm2',	'Sumo1',	'Ffar4',	'Wnt7a',	'Lama5',	'Asah1',	'Btg3',	'Myef2',	'Akap7',	'Cd248',	'Bub1',	'Gm16486',	'Prl',	'Mylk',	'Umps',	'Ildr2',	'Pno1',	'Rbm15',	'Mms22l',	'Zmat4',	'Ankrd37',	'Wdr76',	'Pole4',	'Dclk1',	'Sema6d',	'Mslnl',	'Fgf9',	'Akr1b8',	'Adora2b',	'Prss23',	'Igfbp3',	'Ptprn',	'Gm15965',	'Gm20512',	'Cdc7',	'Hoxb9',	'Bpifa1',	'Trdc',	'Ntn4',	'Nexmif',	'Morf4l2',	'Isyna1',	'Cpne4',	'Dusp14',	'Ruvbl1',	'Kntc1',	'Foxf2',	'Gm19276',	'Xlr4a',	'Perp',	'Pdcd4',	'Acsl4',	'Pnliprp1',	'Map7d1',	'Tmem100',	'Hoxb2',	'Prpf19',	'Cd59a',	'Nr4a3',	'Pdlim7',	'Pbld2',	'Mettl9',	'Plaur',	'Ruvbl2',	'Tmem163',	'Cpm',	'Sp5',	'Mrap2',	'Efcab1',	'Gm15635',	'Nolc1',	'Pabpc1',	'Gadd45a',	'Cldn18',	'Ncf1',	'Pon3',	'S100a4',	'Rps7',	'Lif',	'Psap',	'Ghrl',	'Arhgdib',	'Nbeal2',	'Rcc2',	'Smc3',	'Mcm10',	'BC023719',	'9130015G15Rik',	'Greb1',	'Snrnp25',	'Cbr3',	'Npw',	'Slc25a29',	'1810010H24Rik',	'Tuba8',	'Cbx5',	'Lcp1',	'2900060B14Rik',	'Ube2n',	'Hnrnpf',	'Gm9112',	'Aldh1a3',	'Gm26772',	'Gpc5',	'Psca',	'2810442N19Rik',	'Phactr3',	'Asb17',	'Gckr',	'Azgp1',	'Gm42788',	'Gm48053',	'Gm29773',	'Ccl12',	'Hoxb5os',	'Ly6g',	'Trim31',	'Kcng2',	'Crip2',	'Tnn',	'BC022687',	'Gmps',	'Phf5a',	'Igf1r',	'Ptgr1',	'Filip1',	'F5',	'Gm13010',	'Gm15704',	'Pgpep1l',	'Plk4',	'Actg1',	'Cd2',	'Atcayos',	'Pcsk9',	'Agr2',	'Nfil3',	'M1ap',	'Slc17a2',	'Tef',	'Larp7',	'Cd151',	'Gpm6b',	'Arhgap22',	'Pole3',	'Troap',	'Kras',	'Hpcal1',	'Sf3b4',	'Gm42047',	'Rnaseh2a',	'Arhgap11a',	'Nectin3',	'Fam216a',	'Gm3336',	'Prr5l',	'Samd4',	'Rhov',	'Higd1b',	'Maff',	'Rhobtb3',	'Tgoln1',	'AC160336.1',	'Pou2f2',	'A430073D23Rik',	'Elavl1',	'Sema5a',	'Rps11',	'Pfas',	'Fli1',	'Kctd12',	'Mdfi',	'Gjb6',	'Actg2',	'Dnmt1',	'Hist1h2ae',	'Synm',	'Gcat',	'Emp3',	'Tspan18',	'Hnrnpd',	'Kifc1',	'Wnk4',	'Itpripl1',	'Hspa9',	'Akr1c19',	'Sh3rf2',	'Samd5',	'Lpar1',	'Stk36',	'Gpam',	'Gabarapl1',	'AU021092',	'Map6',	'Slco4c1',
                         'Fzd1',	'Trmt6',	'Islr',	'Trim28',	'Zwint',	'Adssl1',	'Slc39a8',	'Ptprc',	'Bcar3',	'Gsta4',	'Sptssb',	'Prlr',	'A330048O09Rik',	'Slc7a11',	'Gm29292',	'Wdr77',	'Stx19',	'Peg12',	'Gm8186',	'Arhgef38',	'Tulp1',	'Thoc3',	'Dynlt1f',	'Snrpn',	'Asf1a',	'Utp14b',	'Rps6ka6',	'D030018L15Rik',	'Lat',	'Vegfc',	'Itgb3',	'Sf3b6',	'Scin',	'4631405K08Rik',	'Rcc1',	'Usp51',	'Capn6',	'Rpp30',	'Dusp2',	'Chek1',	'Tyrobp',	'Psmd6',	'Kctd14',	'Thrsp',	'Hist1h3d',	'Tcof1',	'Zfp703',	'Smim6',	'Slc43a3',	'Dubr',	'Serpinb6a',	'Fzr1',	'Kcnk1',	'Enpep',	'Nr1d1',	'Eno3',	'Tnfaip3',	'Lmna',	'Pcx',	'Vrk1',	'Ncaph',	'Gm42722',	'Egr3',	'Timeless',	'Gm4117',	'B3galt2',	'A930018P22Rik',	'Cela3b',	'Hspb7',	'4930594M22Rik',	'Casp4',	'Nek11',	'Alox8',	'Gm11615',	'Rgs7bp',	'Xlr4c',	'Gm12525',	'6030469F06Rik',	'Tagap',	'AC154200.1',	'Ipcef1',	'Prune2',	'Myl12a',	'Psma1',	'Aif1l',	'Ak1',	'Rusc1',	'Mcc',	'Pcdh10',	'Lsmem1',	'Pdgfc',	'Rnd3',	'Vbp1',	'Zfp428',	'Eps8l2',	'Wdr12',	'Sik1',	'Ppp4c',	'Reps2',	'Tspan5',	'Rpl10a',	'Ankrd22',	'Hlf',	'Krt34',	'Arfgef1',	'Slco5a1',	'Stau2',	'Pkhd1',	'Tmem131',	'Inpp4a',	'2010300C02Rik',	'Rpl31',	'Il1rl2',	'Slc39a10',	'1700019D03Rik',	'Stk17b',	'Spats2l',	'Acadl',	'Lancl1',	'Cps1',	'Ikzf2',	'Rpl37a',	'Tmbim1',	'Speg',	'Epha4',	'Fam124b',	'Dner',	'Cab39',	'Gpc1',	'Slc35f5',	'Gpr39',	'Nckap5',	'Fam72a',	'Slc26a9',	'Cntn2',	'Nfasc',	'Mdm4',	'Pik3c2b',	'Prelp',	'Arl8a',	'Fam129a',	'Edem3',	'Lamc1',	'Gm10031',	'Fam20b',	'Angptl1',	'Kifap3',	'Mpzl1',	'Gpa33',	'B4galt3',	'Fmn2',	'1700016C15Rik',	'Smyd3',	'Spata17',	'Kctd3',	'Hhat',	'A130010J15Rik',	'Camk1g',	'Cd34',	'Optn',	'Usp6nl',	'Dnajc1',	'Msrb2',	'Arhgap21',	'Zdhhc12',	'Hmcn2',	'Qrfp',	'Lmx1b',	'Mvb12b',	'Pbx3',	'Dab2ip',	'Strbp',	'Nek6',	'Arhgap15',	'Lypd6b',	'Rbm43',	'Tank',	'Fign',	'Stk39',	'Tlk1',	'Itgav',	'Tfpi',	'Ctnnd1',	'Ypel4',	'Ube2l6',	'Gm9821',	'Chst1',	'Prdm11',	'Ehf',	'Depdc7',	'Prrg4',	'Aven',	'Fsip1',	'Bmf',	'Ccdc32',	'Ino80',	'Ehd4',	'Eid1',	'Ap4e1',	'Mrps5',	'Gpcpd1',	'Plcb4',	'Foxa2',	'Pygb',	'Bcl2l1',	'Efcab8',	'Chmp4b',	'Bpi',	'Adig',	'Chd6',	'Ncoa3',	'Sulf2',	'Arfgef2',	'B4galt5',	'Tshz2',	'Pck1',	'Rab22a',	'Zfp831',	'Cables2',	'Clcn5',	'Gripap1',	'Maob',	'Slc9a7',	'44080',	'Xpnpep2',	'Gpc4',	'Hprt',	'Abcd1',	'Prkx',	'Arhgef9',	'Amer1',
                         'Pja1',	'Ogt',	'Gm379',	'Gm6377',	'Pof1b',	'Chm',	'Hnrnph2',	'Gprasp1',	'Bex4',	'Tceal8',	'Rnf128',	'Huwe1',	'Spin2c',	'Gemin8',	'Msl3',	'Mrps28',	'Tpd52',	'Zbtb10',	'Car13',	'Tnik',	'Mecom',	'Actl6a',	'Gm11549',	'Bbs12',	'Spry1',	'Pgrmc2',	'P2ry13',	'Mbnl1',	'Lekr1',	'Rapgef2',	'Gucy1b1',	'Rbm46',	'Dchs2',	'Fhdc1',	'Arfip1',	'Tigd4',	'Fam160a1',	'Etv3',	'Mrpl24',	'Rhbg',	'Atp8b2',	'Vps45',	'Mtmr11',	'Acp6',	'Reg4',	'Ptgfrn',	'Igsf3',	'Casq2',	'Syt6',	'Olfml3',	'Magi3',	'Wnt2b',	'Vav3',	'S1pr1',	'Slc44a3',	'Abca4',	'Gclm',	'1810037I17Rik',	'Camk2d',	'Rrh',	'Dkk2',	'Bdh2',	'Bank1',	'Mttp',	'4930579F01Rik',	'Bmpr1b',	'Pdlim5',	'Ssx2ip',	'Miga1',	'Zzz3',	'Wls',	'Fam110b',	'Asph',	'Tmem64',	'Gpr63',	'Bach2',	'Unc13b',	'Atp8b5',	'Glipr2',	'Gne',	'Fbxo10',	'Trmo',	'Nans',	'Tgfbr1',	'Tal2',	'D630039A03Rik',	'Susd1',	'Slc46a2',	'Slc31a2',	'Rgs3',	'Pappa',	'Tlr4',	'Lurap1l',	'Mpdz',	'Nfib',	'Dennd4c',	'Fggy',	'Ttc22',	'Lrrc42',	'Foxd2',	'Ptprf',	'Foxo6',	'Kcnq4',	'Ak2',	'C77080',	'Bsdc1',	'Epb41',	'Trim63',	'Srrm1',	'Rcan3',	'Ifnlr1',	'Rap1gap',	'Szrd1',	'Dnajc16',	'Dhrs3',	'Lzic',	'Car6',	'B230104I21Rik',	'Mmel1',	'Fam213b',	'Morn1',	'Cdk14',	'Dmtf1',	'Srpk2',	'Shh',	'Garem2',	'Tbc1d14',	'C1qtnf7',
                         'Cc2d2a',	'Qdpr',	'Slit2',	'Ppargc1a',	'Stim2',	'Pcdh7',	'Klf3',	'N4bp2',	'Rbm47',	'Slain2',	'Cwh43',	'Thegl',	'Parm1',	'Scarb2',	'Shroom3',	'Ccni',	'Gpat3',	'Arhgap24',	'Ptpn13',	'Tgfbr3',	'Fgfrl1',	'Crybb3',	'Ssh1',	'Nos1',	'Med13l',	'Atp2a2',	'Tmem120b',	'Tmem132d',	'Gtf2i',	'Sh2b2',	'Cux1',	'Myl10',	'Prkar1b',	'Mad1l1',	'Slc29a4',	'Tnrc18',	'Rbak',	'Lmtk2',	'Bhlha15',	'Alox5ap',	'Tex26',	'Fry',	'Pds5b',	'Stard13',	'Ppp1r9a',	'Asb4',	'Fam3c',	'Lmod2',	'Tspan33',	'Cep41',	'Mkln1',	'Lrguk',	'Bpgm',	'Agbl3',	'Dgki',	'Ephb6',	'Gsdme',	'Nfe2l3',	'Adcyap1r1',	'Thnsl2',	'Sema4f',	'Rtkn',	'Tet3',	'Ankrd53',	'Tgfa',	'Antxr1',	'Slc6a6',	'Prickle2',	'Kbtbd8',	'Mitf',	'Foxp1',	'Eif4e3',	'Shq1',	'Rad18',	'Setd5',	'Atp2b2',	'Vgll4',	'Cand2',	'Wnt5b',	'C1s2',	'Acrbp',	'Iffo1',	'Styk1',	'Bcl2l14',	'E330021D16Rik',	'Atf7ip',	'Pik3c2g',	'Plekha5',	'St8sia1',	'Sox5',	'Smco2',	'Klhl42',	'Bicd1',	'Zfp773',	'Dact3',	'Zfp60',	'Zfp568',	'Zfp536',	'Plekhf1',	'Med25',	'Irf3',	'Spty2d1',	'Dbx1',	'Mkrn3',	'Nr2f2',	'Gm10295',	'Akap13',	'Isg20',	'Wdr73',	'Mex3b',	'Fzd4',	'Ccdc90b',	'Nars2',	'Uvrag',	'Serpinh1',	'P4ha3',	'Arhgef17',	'Pde2a',	'Olfr571',	'Scube2',	'Plekha7',	'Rps15a',	'Dnah3',	'Ndufab1',	'Tbx6',	'Zfp629',	'Uros',	'Tcerg1l',	'Pwwp2b',	'Tspan32',	'Shank2',	'Iyd',	'Plekhg1',	'Ptprk',	'Wisp3',	'Traf3ip2',	'Foxo3',	'Afg1l',	'Rtn4ip1',	'Prdm1',	'Dcbld1',	'Pln',	'Unc5b',	'Pcbd1',	'Tacr2',	'Nrbf2',	'Arid5b',	'Ccdc6',	'Specc1l',	'Zfp280b',	'Dnmt3l',	'AU041133',	'Nfyb',	'Nuak1',	'Chpt1',	'Nr1h4',	'Nudt4',	'Atp2b1',	'Poc1b',	'Tph2',	'Zfc3h1',	'Frs2',	'Cpsf6',	'Rap1b',	'Grip1',	'Tbc1d30',	'Slc16a7',	'Shmt2',	'Baz2a',	'Rab20',	'Cdc16',	'Kbtbd11',	'Xkr5',	'Smim19',	'Eri1',	'Mtnr1a',	'Tlr3',	'Ccdc110',	'Irf2',	'Ing2',	'Cldn22',	'Cpe',	'Psd3',	'Pbx4',	'Mmaa',	'Adgrl1',	'G430095P16Rik',	'Dand5',	'Dnaja2',	'Ces1a',	'Ces1g',	'Ogfod1',	'Pllp',	'Ccdc102a',	'Cfap20',	'Cmtm4',	'Terf2',	'Tmem170',	'Sdr42e1',	'Rhou',	'Rab4a',	'Sipa1l2',	'Pcnx2',	'Pard3',	'Flnb',	'Kctd6',	'Dlg5',	'Il17rd',	'Hacl1',	'Gm8113',	'Prmt5',	'Zfp395',	'Trim35',	'Bnip3l',	'Dock5',	'Chmp7',	'Sorbs3',	'Lpar6',	'Siah3',	'Cog3',	'Serp2',	'Dnajc15',	'Pou4f1',	'Dct',	'Mbnl2',	'Farp1',	'Gm11168',	'Gm10718',	'St14',	'Ets1',	'Fez1',	'Scn3b',	'Sorl1',	'Mpzl3',	'Scn2b',	'Sik3',	'Cadm1',
                         'Zbtb16',	'Ppp2r1b',	'Colca2',	'Tmem266',	'Isl2',	'Peak1',	'1700017B05Rik',	'Stra6',	'Adpgk',	'Lrrc49',	'Tle3',	'Anp32a',	'Coro2b',	'Calml4',	'Pias1',	'Smad6',	'Plekho2',	'Tpm1',	'Adam10',	'Hmgcll1',	'Klhl31',	'Gsta2',	'Eef1a1',	'Irak1bp1',	'Ibtk',	'Spsb4',	'Dbr1',	'Slc35g2',	'Pccb',	'Cep63',	'Amotl2',	'Srprb',	'Atp2c1',	'Wdr82',	'Cmtm8',	'Stt3b',	'Tgfbr2',	'Scn10a',	'Lyzl4',	'Snrk',	'Zfp105',	'Limd1',	'Ccr5',	'Kremen1',	'Rtn4',	'Bod1',	'Sh3pxd2b',	'Ubtd2',	'Fbxw11',	'Ranbp17',	'Slit3',	'Rars',	'Wwc1',	'Mat2b',	'Trim7',	'Ccdc69',	'Galnt10',	'Larp1',	'Btnl10',	'Rai1',	'Zfp286',	'Hs3st3b1',	'Cox10',	'Shisa6',	'Dhrs7c',	'Scimp',	'Med31',	'Rap1gap2',	'Ccdc92b',	'Srr',	'Timm22',	'Rab34',	'Lyrm9',	'Rasl10b',	'Car4',	'Tbx2',	'Hsf5',	'Rnf43',	'Kat7',	'B4galnt2',	'Lasp1',	'Krt25',	'Krt36',	'Plekhh3',	'Ccr10',	'Meox1',	'Cd300lg',	'Mpp2',	'BC030867',	'Ubtf',	'Arhgap27',	'Mettl2',	'Tanc2',	'Cyb561',	'Ern1',	'Bptf',	'Pitpnc1',	'Wipi1',	'Nat9',	'Mgat5b',	'Slc16a3',	'Zfp750',	'Zmynd11',	'Arid4b',	'Inhba',	'Trim27',	'Gpx6',	'Hist1h2bl',	'Hist1h2an',	'Btn1a1',	'Hist1h2bh',	'Hist1h3e',	'Hist1h2bf',	'Hist1h4d',	'Hist1h3a',	'Gpld1',	'Agtr1a',	'Uqcrfs1',	'Gmds',	'Bphl',	'Rreb1',	'Sycp2l',	'Nedd9',	'Tbc1d7',	'Rnf182',	'Jarid2',	'Mylip',	'Atxn1',	'Nup153',	'Zfp169',	'Diras2',	'Syk',	'Cplx2',	'Ddx46',	'2010111I01Rik',	'Aaed1',	'Gm28041',	'Irx1',	'Vcan',	'Lhfpl2',	'Zbed3',	'Nsa2',	'Arhgef28',	'Utp15',	'Fcho2',	'Srek1',	'Snx18',	'Hspb3',	'Arl15',	'Parp8',	'Gm17509',	'Apob',	'Osr1',	'Rock2',	'Cys1',	'Tmem18',	'Alkal2',	'Immp2l',	'Lrrn3',	'Stxbp6',	'Mbip',	'Pole2',	'Klhdc1',	'Atp5s',	'Tmx1',	'Frmd6',	'Six4',	'Tmem30b',	'Prkch',	'Syne2',	'Max',	'Dcaf5',	'Map3k9',	'Sipa1l1',	'Rgs6',	'Fcf1',	'Ttll5',	'Ston2',	'Gpr68',	'Ccdc88c',	'Golga5',	'Ppp4r4',	'Dicer1',	'Bdkrb1',	'Dnah11',	'Ccdc152',	'Osmr',	'Nadk2',	'Rai14',	'Myo10',	'Ctnnd2',	'Nipal2',	'Zfp706',	'Azin1',	'Eif3h',	'Deptor',	'Mrpl13',	'Zhx2',	'Fer1l6',	'Nsmce2',	'Fam84b',	'Adgrb1',	'Eppk1',	'Hsf1',	'Rbfox2',	'Myh9',	'Sstr3',	'Cdc42ep1',	'Syngr1',	'Chadl',	'Atxn10',	'Gramd4',	'Ano6',	'Arid2',	'Nckap5l',	'Lima1',	'Dip2b',	'Sp1',	'Zfp385a',	'Crebbp',	'Ercc4',	'Fopnl',	'Map3k13',	'Igf2bp2',	'St6gal1',	'P3h2',	'Mb21d2',	'Dlg1',	'Stxbp5l',	'Gpr156',	'Gsk3b',	'Upk1b',	'Abhd10',	'Phldb2',	'Morc1',	'Nsun3',	'4930453N24Rik',
                         'Robo1',	'Robo2',	'Grik1',	'Mx1',	'Tmem181a',	'Dynlt1a',	'Rsph3b',	'Pacrg',	'Afdn',	'Zfp820',	'Zfp947',	'Zfp945',	'Traf7',	'Neurl1b',	'Rsph1',	'Cryaa',	'Brd2',	'C2',	'Ppp1r10',	'Adgrf5',	'Supt3',	'Unc5cl',	'Pp2d1',	'Ptprs',	'Efna5',	'Man2a1',	'Themis3',	'Epb41l3',	'Vit',	'Cdc42ep3',	'Pkdcc',	'Eml4',	'Haao',	'Lrpprc',	'Svil',	'Ccny',	'Dtna',	'Nrep',	'Reep2',	'Sil1',	'Psd2',	'Pcdh1',	'Nr3c1',	'Gm5689',	'Stk32a',	'Snx24',	'Zfp608',	'Tex43',	'Slc27a6',	'Sh3tc2',	'Wdr7',	'Zfp532',	'Oacyl',	'Grp',	'Lman1',	'Smad4',	'Mro',	'Gm9925',	'Myo5b',	'Acaa2',	'Zbtb7c',	'Nfatc1',	'Zfp516',	'Cndp1',	'Ppp6r3',	'Kdm2a',	'Mus81',	'Pcnx3',	'Pla2g16',	'Chrm1',	'Fads2',	'1110059E24Rik',	'Pum3',	'Slc16a12',	'Ccnj',	'Tm9sf3',	'Frat1',	'Ina',	'Sfr1',	'Afap1l2',	'Gfra1',	'Pnlip',	'Prdx3',	'Xkr4',	'Jph1',	'Fam178b',	'Gm3646',	'Pou3f3',	'Nck2',	'Rpe',	'Erbb4',	'Acsl3',	'Pid1',	'Glrp1',	'Cops8',	'Twist2',	'Serpinb8',	'Cxcr4',	'Lemd1',	'Lgr6',	'Cdc73',	'Hmcn1',	'Teddm2',	'Fam163a',	'Rxrg',	'1700056E22Rik',	'Lyplal1',	'Esrrg',	'Smyd2',	'Prox1',	'Plxna2',	'Camk1d',	'St8sia6',	'Hacd1',	'Notch1',	'Sardh',	'Rxra',	'Ppp1r26',	'Barhl1',	'Med27',	'Zbtb43',	'Crb2',	'Nr5a1',	'Fmnl2',	'Dpp4',	'Zfp385b',	'Prdx6b',	'Zfp804a',	'Slc1a2',	'Wt1',	'Them7',	'Arl14ep',	'Ryr3',	'Rasgrp1',	'Frmd5',	'Sord',	'Gatm',	'Sqor',	'Gm9913',	'Bcl2l11',	'Mrps26',	'Snap25',	'Ovol2',	'Syndig1',	'Gm14226',	'Src',	'Dhx35',	'Tox2',	'Kcng1',	'Sall4',	'Tfap2c',	'Bmp7',	'Ankrd60',	'Stx16',	'Ccdc160',	'Asb12',	'Snx12',	'Pak3',	'Zfp704',	'Pag1',	'Dnajc5b',	'Fndc3b',	'Tmem212',	'Egfem1',	'Slc7a14',	'Mab21l1',	'Gpr87',	'Plch1',	'Iqcj',	'Rxfp1',	'Sfrp2',	'Tmem131l',	'Kirrel',	'Fcgr1',	'Vangl1',	'A630076J17Rik',	'Col11a1',	'Cdc14a',	'Plppr5',	'Dpyd',	'Ptbp2',	'Ugt8a',	'Elovl6',	'Lef1',	'Adh6a',	'Unc5c',	'Pkn2',	'Col24a1',	'St6galnac5',	'Triqk',	'Fut9',	'Lingo2',	'Pax5',	'Igfbpl1',	'Xpa',	'Rad23b',	'Ambp',	'Astn2',	'Trim32',	'Brinp1',	'Gm11232',	'Slc24a2',	'Mtap',	'Hook1',	'E130114P18Rik',	'Lepr',	'Pde4b',	'Dmrta2',	'3100002H09Rik',	'Nkain1',	'Sdc3',	'Ptpru',	'Ephb2',	'Wnt4',	'Igsf21',	'Camta1',	'A430005L14Rik',	'Fam133b',	'Speer4c',	'Mnx1',	'Cad',	'Babam2',	'Cpz',	'Sorcs2',	'Zfp518b',	'Gm16223',	'Prom1',	'Smim20',	'9130230L23Rik',	'Lnx1',	'Adgrl3',	'Npffr2',	'Prkg2',	'Mapk10',	'Sez6l',	'Myo18b',	'Aym1',	'Gcn1l1',	
                         'Hspb8',	'Ksr2',	'2410131K14Rik',	'Rbm19',	'Rflna',	'Tmem132c',	'Fzd10',	'Galnt17',	'Gtf2ird1',	'Ephb4',	'Slc25a13',	'Glcci1',	'Aass',	'Plxna4',	'Chchd3',	'Hipk2',	'Tmem178b',	'Tpk1',	'Cntnap2',	'Gimap7',	'Igf2bp3',	'Snx10',	'Herc6',	'Atoh1',	'Gfpt1',	'Iqsec1',	'1700123L14Rik',	'Frmd4b',	'Rybp',	'Lrrn1',	'Hrh1',	'Ntf3',	'Etv6',	'Grin2b',	'Gucy2c',	'Ptpro',	'Lmo3',	'Pde3a',	'Etnk1',	'Usp29',	'Ccdc8',	'Tshz3',	'Kcnj11',	'Htatip2',	'Chrna7',	'St8sia2',	'Tmem135',	'Rab30',	'Emsy',	'Wnt11',	'Insc',	'Acsm1',	'Acsm4',	'Eef2k',	'Zkscan2',	'Wdr11',	'Nps',	'Mgmt',	'Kndc1',	'Ifitm10',	'Mrpl23',	'Cttn',	'Akap12',	'Myb',	'Echdc1',	'Amd2',	'Crybg1',	'Chst3',	'Ctnna3',	'Lrrtm3',	'Phyhipl',	'Pofut2',	'Chst11',	'Appl2',	'1700113H08Rik',	'Cep83',	'Btg1',	'Lum',	'Kitl',	'Otogl',	'Bbs10',	'Ptprr',	'Efnb2',	'Mcf2l',	'Csmd1',	'Unc5d',	'Cenpu',	'Enpp6',	'Tenm3',	'Galnt7',	'Cbr4',	'Cers1',	'Comp',	'Il15',	'Ndufb7',	'Sall1',	'Tox3',	'Rbl2',	'Irx5',	'Drc7',	'Setd6',	'Cdh8',	'Cdh11',	'Cdh3',	'Zfhx3',	'Fa2h',	'Maf',	'Tldc1',	'Foxl1',	'Map3k21',	'Tomm20',	'Nrp1',	'Top2b',	'Gng2',	'Slc35f4',	'Gata4',	'Gfra2',	'Htr2a',	'Olfm4',	'Pcdh20',	'Pcdh9',	'Klf12',	'Ipo5',	'Stk24',	'Nalcn',	'Fgf14',	'Sesn3',	'Jam3',	'Igsf9b',	'Opcml',	'Barx2',	'St3gal4',	'Pknox2',	'Tbcel',	'Grik4',	'Pou2f3',	'Nxpe4',	'Elmod1',	'Cspg4',	'Loxl1',	'Uaca',	'Igdcc3',	'Rora',	'Impg1',	'Sh3bgrl2',	'Slc9a9',	'Rasa2',	'Mrps22',	'Mrpl3',	'Arpp21',	'Trim71',	'Scn5a',	'Mobp',	'Pkd1l1',	'Hus1',	'Vwc2',	'Meis1',	'Ccdc85a',	'Sptbn1',	'Srebf1',	'Dnah9',	'Glp2r',	'Ntn1',	'Smtnl2',	'1700016K19Rik',	'Wsb1',	'Mrm1',	'Bcas3',	'Nog',	'Ankfn1',	'Pctp',	'Hils1',	'Tac4',	'Phb',	'Igf2bp1',	'Ikzf3',	'Krt20',	'Krt13',	'G6pc3',	'Wnt9b',	'1700012B07Rik',	'Map2k6',	'Kcnj16',	'Sdk2',	'Tnrc6c',	'Cyth1',	'Cbx4',	'Chrm3',	'Elmo1',	'Nrsn1',	'Cdkal1',	'Mylk4',	'Hivep1',	'Rnf144b',	'Sema4d',	'Ror2',	'Msx2',	'Pitx1',	'Spock1',	'Agtpbp1',	'Gas1',	'Dapk1',	'Ercc6l2',	'Hsd17b3',	'Irx2',	'Poc5',	'Cartpt',	'Naip6',	'Isl1',	'B430218F22Rik',	'Dnmt3a',	'Pomc',	'Dnajc27',	'Fam49a',	'Sntg2',	'Sostdc1',	'Dgkb',	'Npas3',	'Egln3',	'Dact1',	'Slc38a6',	'Actn1',	'Dpf3',	'Jdp2',	'Tgfb3',	'Esrrb',	'Sel1l',	'Akt1',	'Ghr',	'Adamts12',	'Npr3',	'Otulin',	'Ankrd33b',	'Trps1',	'Sntb1',	'Tmem65',	'Fam49b',	'St3gal1',	'Slc45a4',	'Gpr20',	'Mb',	'Tcf20',	'Rtl6',	
                         'Fam118a',	'Fam19a5',	'Syt10',	'Tmem117',	'Slc38a4',	'Smug1',	'Sec14l5',	'Eef2kmt',	'Rtl10',	'Klhl6',	'Il1rap',	'Gmnc',	'Fgf12',	'Opa1',	'Fam43a',	'Tfrc',	'Hacd2',	'Csta1',	'Igsf11',	'Lsamp',	'Bach1',	'Setd4',	'Ets2',	'B3galt5',	'Pcp4',	'Dact2',	'Cldn9',	'1520401A03Rik',	'Lemd2',	'Glp1r',	'Pde9a',	'H2-Q2',	'Spats1',	'Ptprm',	'Arhgap28',	'Dlgap1',	'Crim1',	'Thada',	'Slc3a1',	'Six2',	'Srbd1',	'Zfp521',	'B4galt6',	'Arap3',	'Pabpc2',	'Spink1',	'Sema6a',	'Eno1b',	'Cep120',	'Prrc1',	'St8sia3',	'Pmaip1',	'Tcf4',	'Ccdc68',	'Dcc',	'Timm21',	'Nrxn2',	'Gnaq',	'Rfk',	'Trpm3',	'Mamdc2',	'Dmrt3',	'Gm815',	'Pten',	'Gm340',	'Sorcs3',	'Sorcs1',	'Add3',	'Tectb',	'Tcf7l2',	'Habp2',	'Emx2',	'Cops5',	'Ncoa2',	'Tram1',	'Lactb2',	'Eya1',	'Msc',	'Terf1',	'Ly96',	'B3gat2',	'Ptp4a1',	'Ccdc115',	'Cnnm3',	'Lipt1',	'Lyg2',	'Rev1',	'Aff3',	'Pdcl3',	'Map4k4',	'Il1r2',	'Mrps9',	'AI597479',	'Uxs1',	'Dnah7b',	'Nabp1',	'Stat4',	'Hibch',	'Pgap1',	'Plcl1',	'Satb2',	'Fzd7',	'Gm973',	'Bmpr2',	'Fam117b',	'Fzd5',	'Idh1',	'Spag16',	'Mreg',	'Pecr',	'Xrcc5',	'Tns1',	'Catip',	'Ctdsp1',	'Cnot9',	'Zfp142',	'Wnt10a',	'Ihh',	'Ap1s3',	'Irs1',	'Rhbdd1',	'Daw1',	'A630001G21Rik',	'Armc9',	'Spp2',	'Agap1',	'Gbx2',	'Ackr3',	'Col6a3',	'Ube2f',	'Asb1',	'Hdac4',	'Crocc2',	'Stk25',	'Pdcd1',	'Phlpp1',	'Gli2',	'Steap3',	'Actr3',	'Ccnt2',	'Cd55b',	'Zp3r',	'Fcamr',	'Mapkapk2',	'Dyrk3',	'Rassf5',	'Ikbke',	'Slc45a3',	'Cdk18',	'Nuak2',	'Lax1',	'Optc',	'Cyb5r1',	'Syt2',	'Elf3',	'Lmod1',	'Ipo9',	'Csrp1',	'Tmem9',	'Kif14',	'Dennd1b',	'Trove2',	'Ivns1abp',	'Arpc5',	'Ncf2',	'Smg7',	'Shcbp1l',	'Dhx9',	'Npl',	'Rnasel',	'Cacna1e',	'BC034090',	'Qsox1',	'Tor1aip1',	'Abl2',	'Ralgps2',	'Pappa2',	'4930558K02Rik',	'Mettl11b',	'Xcl1',	'Uck2',	'Aldh9a1',	'Lrrc52',	'Pbx1',	'Fcgr4',	'Fcgr3',	'Cfap126',	'Tomm40l',	'Alyref2',	'Cfap45',	'Cadm3',	'Pld5',	'Rbm8a2',	'Akt3',	'Catspere2',	'Kif26b',	'Cnst',	'Nvl',	'Lbr',	'Srp9',	'Tlr5',	'43891',	'Bpnt1',	'Ptpn14',	'Rps6kc1',	'Nenf',	'Lpgat1',	'Sertad4',	'Cd46',	'Cr1l',	'Nmt2',	'Bend7',	'Proser2',	'Gata3',	'Taf3',	'Itih5',	'Sfmbt2',	'Trdmt1',	'Commd3',	'Pdss1',	'Abi1',	'Psd4',	'Dph7',	'Nrarp',	'Ccdc187',	'Sec16a',	'Vav2',	'Wdr5',	'Fcnb',	'Mrps2',	'Spaca9',	'Cfap77',	'Rapgef1',	'Sptan1',	'Ptpa',	'Ier5l',	'Cstad',	'1700001O22Rik',	'Ncs1',	'Exosc2',	'Slc25a25',	'Fam102a',	'Dpm2',	'Fam129b',	'Garnl3',	
                         'Angptl2',	'Zbtb34',	'Rab14',	'Stom',	'Ttll11',	'Ndufa8',	'Rabgap1',	'Nr6a1',	'Acvr2a',	'Mbd5',	'Epc2',	'Arl5a',	'Stam2',	'Prpf40a',	'Rprm',	'Cytip',	'Pkp4',	'Tanc1',	'Wdsub1',	'Rbms1',	'Ifih1',	'Kcnh7',	'Grb14',	'Slc38a11',	'Scn3a',	'Scn2a',	'Myo3b',	'Gad1',	'Cybrd1',	'Metap1d',	'Dlx2',	'Rapgef4',	'Sp9',	'Gpr155',	'Wipf1',	'Chrna1',	'Lnpk',	'Nfe2l2',	'Agps',	'Ttc30a1',	'Rbm45',	'Prkra',	'Cwc22',	'Zdhhc5',	'Gm19426',	'Prg3',	'Tnks1bp1',	'Olfr1274-ps',	'Ptprj',	'Madd',	'Arfgap2',	'1110051M20Rik',	'Lrp4',	'Phf21a',	'1700029I15Rik',	'Cry2',	'Cd82',	'Accs',	'Api5',	'B230118H07Rik',	'Traf6',	'Commd9',	'Abtb2',	'Nat10',	'Caprin1',	'Qser1',	'Kif18a',	'Lin7c',	'BC052040',	'Chst14',	'Rpusd2',	'Dnajc17',	'Chp1',	'Oip5',	'Itpka',	'Rpap1',	'Capn3',	'Zfp106',	'Ubr1',	'Zscan29',	'Pdia3',	'Mfap1b',	'Mfap1a',	'Shf',	'AA467197',	'Gabpb1',	'Trpm7',	'1810024B03Rik',	'Kcnip3',	'Mall',	'Anapc1',	'Mertk',	'Tmem87b',	'Zc3h8',	'Gm14025',	'Ptpra',	'Cenpb',	'Pank2',	'Lamp5',	'Btbd3',	'Macrod2',	'Kif16b',	'Rbbp9',	'Dtd1',	'Insm1',	'Nkx2-2',	'Nxt1',	'Zfp442',	'Acss1',	'Rspo4',	'Csnk2a1',	'Sox12',	'Zcchc3',	'Mcts2',	'Plagl2',	'Commd7',	'Dnmt3b',	'Snta1',	'Cbfa2t2',	'1700003F12Rik',	'Ahcy',	'Itch',	'Pigu',	'Edem2',	'Epb41l1',	'Soga1',	'Rpn2',	'Blcap',	'Vstm2l',	'Plcg1',	'Emilin3',	'Jph2',	'Tomm34',	'Sys1',	'Wfdc2',	'Ncoa5',	'Zfp663',	'Zmynd8',	'Trp53rkb',	'Znfx1',	'Slc9a8',	'Snai1',	'Tmem189',	'Ptpn1',	'Nfatc2',	'Zfp217',	'Bcas1',	'Rae1',	'Prelid3b',	'Zfp931',	'Cdh26',	'Taf4',	'Rps21',	'Rbbp8nl',	'Slco4a1',	'Mrgbp',	'Fndc11',	'Sox18',	'Shroom4',	'Usp27x',	'Bcor',	'Usp9x',	'Gpr82',	'Maoa',	'Jade3',	'Uba1',	'Slc25a43',	'Ube2a',	'Nkap',	'Ocrl',	'Bcorl1',	'Enox2',	'Mbnl3',	'Zfp449',	'Slc9a6',	'1700020N15Rik',	'Cd99l2',	'Flna',	'Rpl10',	'Tbl1x',	'Arx',	'Pcyt1b',	'Maged1',	'Yipf6',	'Stard8',	'Kif4',	'Slc7a3',	'Rtl5',	'Pin4',	'Ercc6l',	'Hdac8',	'Cstf2',	'Cenpi',	'Rpl36a',	'Armcx1',	'Armcx5',	'Bhlhb9',	'Tceal9',	'Tceal3',	'Tsc22d3',	'Kcne1l',	'Rps6ka3',	'Tmsb4x',	'Arhgap6',	'Gm21887',	'Ythdf3',	'Tnfsf10',	'Pld1',	'Mynn',	'Prkci',	'Skil',	'Ccdc39',	'Fxr1',	'Atp11b',	'Dcun1d1',	'D3Ertd254e',	'1810062G17Rik',	'Rab33b',	'Maml3',	'Cog6',	'Stoml3',	'Frem2',	'Smad9',	'Commd2',	'Tsc22d2',	'Rap2b',	'Shox2',	'Schip1',	'Ift80',	'Arl14',	'Nmd3',	'Etfdh',	'Tmem144',	'Ctso',	'Tlr2',	'Trim2',	'Gm6525',	'Fbxw7',	'Gatb',
                         'Glt28d2',	'Mab21l2',	'Prcc',	'Hdgf',	'Rrnad1',	'Nes',	'Mef2d',	'Tsacc',	'Tmem79',	'Arhgef2',	'Syt11',	'Clk2',	'Trim46',	'Krtcap2',	'Efna4',	'Flad1',	'Kcnn3',	'Hax1',	'Tpm3',	'Crtc2',	'Ints3',	'Rptn',	'Snx27',	'Rfx5',	'Psmd4',	'Vps72',	'Scnm1',	'Sema6c',	'Cdc42se1',	'Cers2',	'Setdb1',	'Hormad1',	'Mcl1',	'Tars2',	'Rprd2',	'Bola1',	'Hist2h2aa2',	'Hfe2',	'Bcl9',	'Notch2',	'Ttf2',	'Nras',	'Trim33',	'Rsbn1',	'Lrig2',	'Ddx20',
                         'Adora3',	'Kcnc4',	'Csf1',	'Gstm7',	'Atxn7l2',	'5330417C22Rik',	'Taf13',	'Rtca',	'Dbt',	'Alg14',	'Arhgap29',	'Fnbp1l',	'Synpo2',	'Egf',	'Ostc',	'Cyp2u1',	'Papss1',	'Aimp1',	'Nfkb1',	'Ppp3ca',	'Gm21962',	'Gtf2b',	'Lmo4',	'Sh3glb1',	'Znhit6',	'Ttll7',	'Adgrl2',	'Dnajb4',	'St6galnac3',	'Erich3',	'Gm11808',	'Penk',	'Nsmaf',	'Rab2a',	'Chd7',	'2610301B20Rik',	'Pdp1',	'Rbm12b2',	'Manea',	'Pnrc1',	'Akirin2',	'Aco1',	'B4galt1',	'Ubap2',	'Ubap1',	'Enho',	'Fam166b',	'Ccdc107',	'Trmt10b',	'Shb',	'Ncbp1',	'Anks6',	'Stx17',	'Tex10',	'Rnf20',	'Fsd1l',	'Ptpn3',	'Inip',	'Zfp37',	'Wdr31',	'4933430I17Rik',	'Kif12',	'Megf9',	'Tle1',	'Cer1',	'Bnc2',	'Adamtsl1',	'Hacd4',	'Klhl9',	'Nfia',	'Dock7',	'Pgm2',	'Ror1',	'Ube2u',	'Cachd1',	'Raver2',	'Ak4',	'Sgip1',	'Tctex1d1',	'Plpp3',	'Fam151a',	'Acot11',	'Ssbp3',	'Elavl4',	'Spata6',	'Cmpk1',	'Stil',	'Pdzk1ip1',	'Tex38',	'Nsun4',	'Uqcrh',	'Rad54l',	'Tmem69',	'Akr1a1',	'Tmem53',	'Tmem125',	'Cfap57',	'Slc2a1',	'Zfp691',	'Cldn19',	'Ccdc30',	'Scmh1',	'Pou3f1',	'9930104L06Rik',	'Dnali1',	'Snip1',	'Ago1',	'5730409E04Rik',	'Zmym4',	'Sfpq',	'Gjb3',	'Zfp362',	'Zbtb8os',	'Kpna6',	'Khdrbs1',	'Gm10300',	'Taf12',	'Rab42',	'Med18',	'Sesn2',	'Smpdl3b',	'Ppp1r8',	'Ahdc1',	'Wasf2',	'Tmem222',	'Kdf1',	'Arid1a',	'Lin28a',	'Cep85',	'Pdik1l',	'Pafah2',	'Man1c1',	'Ldlrap1',	'Rhd',	'Nipal3',	'Fuca1',	'E2f2',	'Asap3',	'Hnrnpr',	'Luzp1',	'Lactbl1',	'Zbtb40',	'Alpl',	'Eif4g3',	'Pla2g2d',	'Pla2g5',	'Capzb',	'Pax7',	'Arhgef10l',	'Padi6',	'Sdhb',	'Fbxo42',	'Rsg1',	'Arhgef19',	'Clcnka',	'Casp9',	'Efhd2',	'Tmem51',	'Kazn',	'Casz1',	'Rbp7',	'Gpr157',	'Vamp3',	'Tnfrsf25',	'Prdm16',	'Ttc34',	'Plch2',	'Ski',	'Faap20',	'Prkcz',	'C1qtnf12',	'Ttll10',	'9430015G10Rik',	'Cyp51',	'Pclo',	'Gnai1',	'Ptpn12',	'Fgl2',	'Kcnh2',	'Chpf2',	'Rheb',	'Prkag2',	'Actr3b',	'Rnf32',	'Dpysl5',	'Agbl5',	'Preb',	'Ift172',	'Ywhah',	'Spon2',	'Uvssa',	'Mxd4',	'Ablim2',	'Grpel1',	'D5Ertd579e',	'Rab28',	'Cpeb2',	'Bst1',	'Tapt1',	'Adgra3',	'Anapc4',	'Tbc1d19',	'0610040J01Rik',	'Klhl5',	'Ugdh',	'Smim14',	'Chrna9',	'Apbb2',	'Phox2b',	'Atp8a1',	'Gm5108',	'Nfxl1',	'Fryl',	'Lrrc66',	'Spata18',	'Usp46',	'Tmem165',	'Aasdh',	'Spink2',	'Mob1b',	'Slc4a4',	'Gc',	'Cox18',	'Afp',	'Rassf6',	'Mthfd2l',	'Sowahb',	'2010109A12Rik',	'4930524J08Rik',	'Enoph1',	'Cds1',	'Aff1',	'Klhl8',	'Lrrc8c',	'Zfp644',	'Gfi1',	'Ube2d2b',	'Mtf2',
                         'Pcgf3',	'Slc26a1',	'Crlf2',	'Gm15446',	'Zfp932',	'Plcxd1',	'Hscb',	'Pitpnb',	'2900026A02Rik',	'Sart3',	'Mmab',	'Fam222a',	'Gm10401',	'Sirt4',	'Pxn',	'Vsig10',	'Wsb2',	'Tesc',	'Rnft2',	'Tmem116',	'Acad12',	'Cux2',	'Kdm2b',	'Orai1',	'B3gnt4',	'Abcb9',	'Arl6ip4',	'Pitpnm2',	'Sbno1',	'Tmed2',	'Ncor2',	'Rcc1l',	'Cldn3',	'Abhd11',	'Mlxipl',	'Baz1b',	'Rhbdd2',	'Ssc4d',	'Dtx2',	'Rasa4',	'Polr2j',	'Col26a1',	'Fis1',	'Plod3',	'Ap1s1',	'Trim56',	'Ufsp1',	'Tsc22d4',	'Gm5294',	'Elfn1',	'Mrm2',	'Eif3b',	'Lfng',	'Gna12',	'Foxk1',	'Fam220a',	'Bri3',	'Nptx2',	'Usp12',	'Pomp',	'Mtus2',	'Medag',	'Wdr95',	'B3glct',	'N4bp2l2',	'Kl',	'Dync1i1',	'Dlx5',	'Mios',	'Ica1',	'Vwde',	'2610001J05Rik',	'Mdfic',	'Tes',	'St7',	'Wnt2',	'Asz1',	'Atp6v1f',	'Tnpo3',	'Nrf1',	'Zc3hc1',	'Klhdc10',	'Cpa4',	'Podxl',	'Exoc4',	'Wdr91',	'Creb3l2',	'Svopl',	'Ttc26',	'Tbxas1',	'Kdm7a',	'Rab19',	'Arhgef5',	'Pdia4',	'Skap2',	'Chn2',	'Mindy4',	'Ghrhr',	'Nap1l5',	'Il12rb2',	'Rpia',	'Mrpl35',	'Ptcd3',	'St3gal5',	'Atoh8',	'Usp39',	'Tmem150a',	'Ggcx',	'Capg',	'Tcf7l1',	'Kcmf1',	'Dnah6',	'Wbp1',	'Slc4a5',	'Zfp638',	'Cyp26b1',	'Noto',	'Nat8f4',	'Snrpg',	'Tia1',	'Anxa4',	'Nfu1',	'Tpra1',	'4930590J08Rik',	'Mrps25',	'Adamts9',	'Gm15737',	'Magi1',	'Slc25a26',	'Lrig1',	'Fam19a4',	'Lmod3',	'Itpr1',	'Sec13',	'Atg7',	'Tsen2',	'Plxnd1',	'Washc2',	'Csgalnact2',	'Ret',	'Zfp9',	'Dcp1b',	'Fbxl14',	'Ninj2',	'B4galnt3',	'D330020A13Rik',	'Hdhd5',	'Cecr2',	'Pex26',	'Phc1',	'Rimklb',	'Lpcat3',	'Phb2',	'Grcc10',	'Usp5',	'Ing4',	'Tnfrsf1a',	'Vwf',	'Parp11',	'Klrb1c',	'Klrb1b',	'Ybx3',	'Mansc1',	'Dusp16',	'Ddx47',	'Gsg1',	'Pbp2',	'Gm8994',	'Plbd1',	'Eps8',	'Mgst1',	'Slco1c1',	'Cmas',	'Arntl2',	'Caprin2',	'Oscar',	'Rdh13',	'Eps8l1',	'Hspbp1',	'Kmt5c',	'Zfp444',	'Zfp667',	'Zfp418',	'Rnf225',	'Zfp324',	'Zbtb45',	'Chmp2a',	'Bicra',	'Slc8a2',	'Prkd2',	'Calm3',	'Mypop',	'Irf2bp1',	'Six5',	'Exoc3l2',	'Ppp1r37',	'Relb',	'Clptm1',	'Nectin2',	'Bcam',	'Zfp61',	'Lypd3',	'Zfp526',	'Gsk3a',	'Erf',	'Cic',	'Pafah1b3',	'Megf8',	'Ccdc97',	'Rab4b',	'Map3k10',	'Zfp607b',	'Zfp626',	'Rinl',	'Hnrnpl',	'Ech1',	'Capn12',	'Actn4',	'Sipa1l3',	'Zfp790',	'Zfp566',	'Cox7a1',	'E130208F15Rik',	'Lrfn3',	'Nfkbid',	'U2af1l4',	'Zbtb32',	'Upk1a',	'Lgi4',	'Pdcd2l',	'Kctd15',	'Cebpg',	'Lrp3',	'Uri1',	'1600014C10Rik',	'Syt3',	'Emc10',	'Nr1h2',	'Cpt1c',	'Scaf1',	'Slc17a7',	'Ccdc155',	'Bax',	'Mamstr',	'Cyth2',	'Abcc6',	'Kcnc1',	
                         'Ptpn5',	'Atp10a',	'Mtmr10',	'Tm2d3',	'Selenos',	'Lrrk1',	'Asb7',	'Cers3',	'Adamts17',	'Ttc23',	'Fam169b',	'Arrdc4',	'Mctp2',	'Rgma',	'Chd2',	'Klhl25',	'Idh2',	'Blm',	'Iqgap1',	'3110040N11Rik',	'Mesd',	'Arnt2',	'Rab38',	'Eed',	'Myo7a',	'Thap12',	'Rnf169',	'Kcne3',	'C2cd3',	'Plekhb1',	'Stard10',	'Rnf121',	'Trim34a',	'Cyb5r2',	'Tub',	'St5',	'Ascl3',	'Zfp143',	'Sbf2',	'Ampd3',	'Usp47',	'Dkk3',	'Mical2',	'Parva',	'Arntl',	'Rras2',	'Sox6',	'1110004F10Rik',	'Knop1',	'Iqck',	'Rexo5',	'Mosmo',	'Rbbp6',	'Tnrc6a',	'Kdm8',	'Il21r',	'Mapk3',	'Pagr1a',	'Cd2bp2',	'E430018J23Rik',	'Bcl7c',	'Fbxl19',	'Orai3',	'Zfp646',	'Prss8',	'Sec23ip',	'Ate1',	'Tacc2',	'Btbd16',	'Plekha1',	'Lhpp',	'Edrf1',	'Dhx32',	'Adam12',	'Ebf3',	'Dpysl4',	'Inpp5a',	'Cfap46',	'Cox8b',	'Lrrc56',	'Pidd1',	'Ctsd',	'Syt8',	'Ins2',	'Tnfrsf22',	'Fadd',	'1810010D01Rik',	'Mrgprd',	'Armt1',	'Ust',	'Sash1',	'Phactr2',	'Aig1',	'Hivep2',	'Heca',	'Map7',	'1700020N01Rik',	'Slc18b1',	'L3mbtl3',	'Arhgap18',	'Ncoa7',	'Hddc2',	'Frk',	'Fam229b',	'Slc16a10',	'Gtf3c6',	'Cdk19',	'Mettl24',	'Cdc40',	'Fig4',	'Cd164',	'Atg5',	'Prep',	'Slc35f1',	'Pkib',	'Sh3rf3',	'Gm10273',	'Oit3',	'Anapc16',	'Gm17455',	'Col13a1',	'Dna2',	'Hnrnph3',	'Pbld1',	'Reep3',	'Zfp365',	'Cabcoco1',	'Rhobtb1',	'Rsph14',	'Rab36',	'Cabin1',	'Gstt2',	'Gm5134',	'Prmt2',	'Pttg1ip',	'Plpp2',	'Cnn2',	'Sbno2',	'Stk11',	'Ndufs7',	'Reep6',	'Btbd2',	'Slc39a3',	'Thop1',	'Eef2',	'Nmrk2',	'Matk',	'Tjp3',	'Tle2',	'Zfp873',	'Gm4924',	'Rfx4',	'Ric8b',	'Mterf2',	'Cry1',	'Ascl4',	'Fbxo7',	'Stab2',	'Gnptab',	'Sycp3',	'Spic',	'Anks1b',	'Cfap54',	'Cdk17',	'Lta4h',	'Plxnc1',	'Cradd',	'Socs2',	'4930430F08Rik',	'Nts',	'Ptprq',	'Pawr',	'E2f7',	'Krr1',	'Ptprb',	'Cnot2',	'Myrfl',	'Mdm2',	'Msrb3',	'Usp15',	'Lrig3',	'Atp23',	'Avil',	'43899',	'Kif5a',	'Gli1',	'Nab2',	'Atp5b',	'Rbms2',	'Spryd4',	'Pan2',	'Cnpy2',	'Cs',	'Zc3h10',	'Rpl41',	'Erbb3',	'Rps26',	'Ikzf4',	'Dgka',	'Pym1',	'Dnajc14',	'Gdf11',	'Mettl7b',	'Insr',	'Xab2',	'Ccl25',	'Cers4',	'Myo16',	'Irs2',	'Col4a2',	'Ankrd10',	'Tex29',	'Pcid2',	'Vps36',	'Polb',	'Ap3m2',	'Kat6a',	'Plekha2',	'Plpbp',	'Rnf122',	'Fut10',	'Tex15',	'Rbpms',	'Prag1',	'Micu3',	'Mtus1',	'Pcm1',	'Frg1',	'Fat1',	'F11',	'Lrp2bp',	'Primpol',	'Casp3',	'Cdkn2aip',	'Mfap3l',	'Clcn3',	'Nek1',	'Sh3rf1',	'Apela',	'Tma16',	'Naf1',	'Sh2d4a',	'Atp6v1b2',	'Zfp930',	'Yjefn3',	'Gatad2a',	'Ddx49',	'Crlf1',	'Kcnn1',	'Abhd8',	
                         'Fcho1',	'Large1',	'Arhgap10',	'Tmem184c',	'Zfp827',	'Smad1',	'Otud4',	'Smarca5',	'Gab1',	'Tbc1d9',	'Scoc',	'1700067K01Rik',	'Rad23a',	'Tnpo2',	'Neto2',	'Adcy7',	'Fto',	'Irx3',	'Irx6',	'Amfr',	'Arl2bp',	'Polr2c',	'Dok4',	'Adgrg1',	'Tepp',	'Csnk2a2',	'Gins3',	'Cklf',	'Nae1',	'Cdh16',	'B3gnt9',	'Slc9a5',	'Lrrc36',	'Zdhhc1',	'Atp6v0d1',	'Enkd1',	'Ranbp10',	'Cenpt',	'Psmb10',	'Slc12a4',	'Esrp2',	'Slc7a6',	'Smpd3',	'Gm10073',	'Cdh1',	'Tango6',	'Vps4a',	'Vac14',	'Mlkl',	'Cfdp1',	'Adat1',	'Gm6793',	'Mon1b',	'Wwox',	'Bco1',	'Gan',	'Plcg2',	'Fam92b',	'Klhdc4',	'Slc7a5',	'Mvd',	'Galnt2',	'Ntpcr',	'Slc35f3',	'Irf2bp2',	'Fam107a',	'4930452B06Rik',	'Thrb',	'Kcnk5',	'Nudt13',	'Adk',	'Kat6b',	'Samd8',	'Vdac2',	'Zfp503',	'Zmiz1',	'Arf4',	'Tmem110',	'Sh3bp5',	'Ankrd28',	'Dph3',	'Oxnad1',	'Bmpr1a',	'Sh2d4b',	'Fermt2',	'Gm15217',	'Fbxo34',	'Tmem260',	'Klhl33',	'Rnase2a',	'Ang4',	'Mettl17',	'Ndrg2',	'Sall2',	'Lrp10',	'Cdh24',	'Slc7a8',	'Cmtm5',	'Ngdn',	'Zfhx2',	'Thtpa',	'Nynrin',	'Mcpt8',	'Parp4',	'Pspc1',	'Zmym5',	'Zmym2',	'Il17d',	'Spata13',	'Mipep',	'Arl11',	'Wdfy2',	'Pinx1',	'Msra',	'Kif13b',	'Pnoc',	'Scara5',	'Ccdc25',	'Ppp2r2a',	'Adam7',	'Nkx3-1',	'R3hcc1',	'Pebp4',	'Reep4',	'Npm2',	'Dok2',	'Itm2b',	'Nufip1',	'Naa16',	'Kbtbd6',	'Wbp4',	'Tdrd3',	'Fbxl3',	'Slain1',	'Slitrk1',	'Abcc4',	'Dnajc3',	'Rap2a',	'Dock9',	'Timm8a2',	'Ggact',	'Tmtc4',	'Alkbh8',	'Gucy1a2',	'Dync2h1',	'Birc3',	'Trpc6',	'Endod1',	'Cep295',	'Ubl5',	'Mrpl4',	'Icam4',	'Icam5',	'Fdx1l',	'Ap1m2',	'Qtrt1',	'Ldlr',	'Rab3d',	'Glb1l2',	'Snx19',	'Aplp2',	'Kcnj5',	'Kirrel3',	'Cdon',	'Pus3',	'Stt3a',	'Slc37a2',	'Hspa8',	'Sc5d',	'Tecta',	'Arhgef12',	'Vps11',	'Phldb1',	'Atp5l',	'Ube4a',	'Jaml',	'Cep164',	'Pcsk7',	'Pafah1b2',	'Apoa1',	'Nnmt',	'Sik2',	'1810046K07Rik',	'Ddx10',	'Exph5',	'Kdelc2',	'4930550C14Rik',	'Cib2',	'Hykk',	'Rcn2',	'Lingo1',	'Snupn',	'Sin3a',	'Ppcdc',	'Edc3',	'Clk3',	'Arid3b',	'Sema7a',	'Pml',	'Cd276',	'Nptn',	'Senp8',	'Thsd4',	'Rplp1',	'Map2k5',	'Iqch',	'Aagab',	'Smad3',	'Snapc5',	'Dis3l',	'Rab11a',	'Parp16',	'Clpx',	'Oaz2',	'Snx22',	'Snx1',	'Herc1',	'Usp3',	'Rps27l',	'Lactb',	'Tln2',	'Ice2',	'Myo1e',	'Aldh1a2',	'Myzap',	'Zfp280d',	'Rfx7',	'Nedd4',	'Prtg',	'Pygo1',	'Wdr72',	'Myo5c',	'Bmp5',	'Hcrtr2',	'Fam83b',	'Mlip',	'Lrrc1',	'Elovl4',	'Ube2cbp',	'Cep162',	'Adamts7',	'Plscr1',	'Clstn2',	'Nme9',	'Dzip1l',	'Il20rb',	'Slco2a1',	'1300017J02Rik',	'Acpp',	'Pik3r4',	
                         'Col6a6',	'Manf',	'Cacna2d2',	'Hyal2',	'Sema3f',	'Mst1r',	'Traip',	'Amigo3',	'Nicn1',	'Rhoa',	'Qrich1',	'Dalrd3',	'Ip6k2',	'Cdc25a',	'Klhl18',	'Vill',	'Exog',	'Eif1b',	'Entpd3',	'Ctnnb1',	'Ulk4',	'Pomgnt2',	'Topaz1',	'Tcaim',	'Sacm1l',	'Pik3ip1',	'Zmat5',	'Nf2',	'Emid1',	'Znrf3',	'Polm',	'Gm11985',	'Igfbp1',	'Spata48',	'Cobl',	'Egfr',	'Wdr92',	'Peli1',	'Fam161a',	'4931440F15Rik',	'Kcnmb1',	'Clint1',	'Thg1l',	'Sox30',	'Irgm1',	'Olfr1396',	'Cby3',	'Zfp879',	'Zfp2',	'Zfp354b',	'Zfp354a',	'Phykpl',	'Jade2',	'Skp1a',	'Tcf7',	'Gm10447',	'Gm12216',	'Slc22a4',	'Pdlim4',	'Anxa6',	'Gm2a',	'Fat2',	'Atox1',	'Mfap3',	'Zfp692',	'Guk1',	'Wnt9a',	'Mprip',	'Cops3',	'Llgl1',	'Zswim7',	'Trpv2',	'Lrrc75a',	'F930015N05Rik',	'Tmem220',	'Pik3r6',	'Mfsd6l',	'Rpl26',	'Borcs6',	'Tmem107',	'Hes7',	'Aloxe3',	'Gucy2e',	'Chd3',	'Kdm6b',	'Dnah2',	'Wrap53',	'Cldn7',	'Dlg4',	'Pelp1',	'Mink1',	'Tekt1',	'Ggt6',	'Spns2',	'P2rx1',	'Ncbp3',	'Haspin',	'Rpa1',	'Ywhae',	'Rph3al',	'Rflnb',	'Glod4',	'Ssh2',	'Coro6',	'Ankrd13b',	'Git1',	'Myo18a',	'Gm10277',	'Phf12',	'Fam222b',	'Rpl23a',	'Aldoc',	'Gm9964',	'Suz12',	'Rnf135',	'Psmd11',	'Myo1d',	'Spaca3',	'Taf15',	'Wfdc21',	'Hnf1b',	'Ddx52',	'Lhx1',	'Gm11444',	'Brip1',	'Med13',	'Ypel2',	'Rad51c',	'Tex14',	'Supt4a',	'Dynll2',	'Cuedc1',	'Mrps23',	'Msi2',	'Akap1',	'Scpep1',	'Gm525',	'Tom1l1',	'Utp18',	'Epn3',	'Samd14',	'Dlx3',	'Fam117a',	'Spop',	'Nxph3',	'Abi3',	'Prr15l',	'Sp2',	'4933428G20Rik',	'Srcin1',	'Epop',	'Mllt6',	'Pcgf2',	'Fbxo47',	'Pgap3',	'Grb7',	'Wipf2',	'Rara',	'Tns4',	'Eif1',	'Hap1',	'Jup',	'P3h4',	'Fkbp10',	'Atp6v0a1',	'Coasy',	'Aarsd1',	'Rdm1',	'Sost',	'Dusp3',	'Nags',	'Gm11627',	'Adam11',	'Eftud2',	'Gfap',	'Nmt1',	'Fmnl1',	'Map3k14',	'Prr29',	'Tex2',	'Smurf2',	'Axin2',	'Arsg',	'Fam20a',	'Abca8b',	'Slc39a11',	'Sstr2',	'Slc9a3r1',	'Tmem104',	'Atp5h',	'Tmem94',	'Caskin2',	'Recql5',	'Itgb4',	'Wbp2',	'St6galnac2',	'Engase',	'Rbfox3',	'Cbx2',	'Cbx8',	'Nptx1',	'Rptor',	'Baiap2',	'Aatk',	'Slc38a10',	'Bahcc1',	'Slc25a10',	'P4hb',	'Arhgdia',	'Aspscr1',	'Rfng',	'Tbcd',	'Ptchd3',	'Calml3',	'Net1',	'Akr1e1',	'Ero1lb',	'Gpr137b',	'Lyst',	'Psma2',	'Mplkip',	'Nme8',	'Zscan12',	'Zkscan4',	'Hist1h4j',	'Zfp184',	'Abt1',	'Hist1h4h',	'Hist1h2af',	'Hist1h3g',	'Hist1h4f',	'Hist1h1d',	'Hist1h2be',	'Hist1h4c',	'Hist1h3c',	'Hist1h4b',	'Hist1h4a',	'Carmil1',	'Hus1b',	'Serpinb1b',	'Psmg4',	'Slc22a23',	'Cdyl',	'Ly86',	'Ssr1',	'Dsp',	'Bmp6',	'Ofcc1',	
                         'Tmem170b',	'Kif13a',	'Fam120a',	'Ninj1',	'Spin1',	'Shc3',	'Secisbp2',	'Sptlc1',	'Gprin1',	'Unc5a',	'Slc34a1',	'B4galt7',	'B230219D22Rik',	'Txndc15',	'Pcbd2',	'Idnk',	'Ntrk2',	'Zfp808',	'6720489N17Rik',	'Platr25',	'Gm5141',	'Fancc',	'Ptch1',	'Slc35d2',	'Zfp367',	'Uqcrb',	'Zfp429',	'Zfp738',	'Zfp85',	'Mtrr',	'1700001L19Rik',	'Adamts16',	'Ndufs6',	'Tert',	'Slc12a7',	'Erap1',	'Pou5f2',	'Edil3',	'Hapln1',	'Ssbp2',	'Fam151b',	'Serinc5',	'Cmya5',	'Bhmt',	'Wdr41',	'Aggf1',	'F2rl1',	'Ankdd1b',	'Fam169a',	'Enc1',	'Tmem171',	'Mccc2',	'Gtf2h2',	'Taf9',	'Mrps36',	'Pik3r1',	'Cd180',	'Nln',	'Adamts6',	'Ndufaf2',	'Smim15',	'Rab3c',	'Ankrd55',	'Plpp1',	'Ccno',	'Cdc20b',	'Gpx8',	'Ndufs4',	'Zfp131',	'Asxl2',	'Sdc1',	'Nbas',	'Lpin1',	'Ntsr2',	'E2f6',	'2410004P03Rik',	'Kcnf1',	'Atp6v1c2',	'Cmpk2',	'Adi1',	'Slc26a3',	'Pik3cg',	'Sypl',	'Atxn7l1',	'Agr3',	'Ifrd1',	'Dock4',	'Prkd1',	'Coch',	'Nubpl',	'Akap6',	'Eapp',	'Brms1l',	'Nkx2-1',	'Nkx2-9',	'Foxa1',	'Trappc6b',	'Gm9887',	'Trim9',	'Psma3',	'Tomm20l',	'Daam1',	'Lrrc9',	'Pcnx4',	'Trmt5',	'Hif1a',	'Gphb5',	'Ppp2r5e',	'Plekhg3',	'Rab15',	'Fut8',	'Fam71d',	'Tmem229b',	'Zfyve26',	'Rad51b',	'Zfp36l1',	'Slc39a9',	'Ccdc177',	'Susd6',	'Srsf5',	'Ttc9',	'Pcnx',	'Dcaf4',	'Zfyve1',	'Fam161b',	'Lin52',	'Isca2',	'Flvcr2',	'Irf2bpl',	'Cipc',	'Pomt2',	'Gtf2a1',	'Galc',	'Ptpn21',	'Ttc8',	'Foxn3',	'Efcab11',	'Psmc1',	'Calm1',	'Ttc7b',	'Rps6ka5',	'Slc24a4',	'Serpina6',	'Clmn',	'Setd3',	'Degs2',	'Gm266',	'Aspg',	'Inf2',	'Cep170b',	'Tedc1',	'Wdr60',	'Dab2',	'Tars',	'Zfr',	'Retreg1',	'Trio',	'Dnah5',	'43896',	'Cct5',	'Matn2',	'Cox6c',	'Ankrd46',	'Ywhaz',	'Grhl2',	'Ubr5',	'Klf10',	'Nudcd1',	'Rad21',	'Med30',	'Ext1',	'Taf2',	'Fam83a',	'Ndufb9',	'Mtss1',	'Asap1',	'Tmem71',	'Tg',	'Zfat',	'Jrk',	'Slurp1',	'Mafa',	'Puf60',	'Exosc4',	'Adck5',	'Zfp251',	'Txn2',	'Ift27',	'Mpst',	'Kctd17',	'Mfng',	'Gga1',	'Pdxp',	'Ddx17',	'Gtpbp1',	'Cbx7',	'Tnrc6b',	'Sgsm3',	'Mkl1',	'Dnajb7',	'Zc3h7b',	'Tob2',	'Xrcc6',	'Srebf2',	'Poldip3',	'Cyb5r3',	'Arfgap3',	'Arhgap8',	'Nup50',	'Upk3a',	'Trmu',	'Celsr1',	'Tbc1d22a',	'Rabl2',	'Pphln1',	'Scaf11',	'Adcy6',	'Ddn',	'Prph',	'Spats2',	'Kcnh3',	'Atf1',	'Slc11a2',	'Slc4a8',	'Atg101',	'Eif4b',	'Mfsd5',	'Espl1',	'Npff',	'Atp5g2',	'Hoxc4',	'Adcy9',	'Tfap4',	'Cdip1',	'Abat',	'1810013L24Rik',	'Tekt5',	'Ciita',	'Clec16a',	'Socs1',	'Mkl2',	'Fgd4',	'Top3b',	'Ccdc116',	'Hic2',	'Lrrc74b',	'Smpd4',	
                         'Klhl22',	'Trmt2a',	'Comt',	'Txnrd2',	'Abcc5',	'Ap2m1',	'Eif4g1',	'Fam131a',	'Ephb3',	'Vps8',	'Etv5',	'Adipoq',	'Bcl6',	'Lpp',	'Tprg',	'Atp13a5',	'Tmem44',	'Xxylt1',	'Bdh1',	'Zdhhc19',	'Tnk2',	'Iqcg',	'Lmln',	'Osbpl11',	'Snx4',	'Slc12a8',	'Kalrn',	'Ropn1',	'Fam162a',	'Stfa3',	'Slc15a2',	'Zbtb20',	'Qtrt2',	'Zdhhc23',	'Gramd1c',	'Sidt1',	'Cfap44',	'Cblb',	'Alcam',	'Nit2',	'Tbc1d23',	'Cmss1',	'Cggbp1',	'Chmp2b',	'Cxadr',	'N6amt1',	'Scaf4',	'4932438H23Rik',	'Rcan1',	'Runx1',	'Morc3',	'Igsf5',	'Dscam',	'Bace2',	'Tmprss2',	'A630089N07Rik',	'Cldn20',	'Arid1b',	'Ldhal6b',	'Rsph3a',	'Rgmb',	'Spaca6',	'Zfp53',	'Zfp13',	'Bicdl2',	'Dcpp3',	'Kctd5',	'Tpsg1',	'Sstr5',	'Mettl26',	'Mrpl28',	'Luc7l',	'Cuta',	'D17Wsu92e',	'Tead3',	'Brpf3',	'Rnf8',	'Zfand3',	'Btbd9',	'Dnah8',	'Tmprss3',	'Ndufv3',	'Hsf2bp',	'Notch3',	'Ephx3',	'Akap8l',	'Cyp4f17',	'Rab11b',	'Zbtb22',	'Rgl2',	'H2-DMb2',	'Vars',	'Abhd16a',	'Prrc2a',	'H2-Q10',	'Gtf2h4',	'H2-T23',	'Mog',	'9130008F23Rik',	'Adgrf2',	'Cd2ap',	'Rcan2',	'Enpp5',	'Cdc5l',	'Tmem151b',	'Slc35b2',	'Mrpl14',	'Ptk7',	'Ubr2',	'Mrps10',	'Guca1a',	'Taf8',	'Ccnd3',	'1700122O11Rik',	'1700067P10Rik',	'Nfya',	'Lrfn2',	'Daam2',	'Satb1',	'Mpnd',	'Fem1a',	'Ticam1',	'Gtf2f1',	'Slc25a23',	'Ralbp1',	'Lama1',	'Zbtb14',	'Lpin2',	'Yipf4',	'Fez2',	'Sult6b1',	'Ndufaf7',	'Rmdn2',	'Galm',	'Dhx57',	'Cdkl4',	'Kcng3',	'Mta3',	'Zfp36l2',	'Plekhh2',	'Camkmt',	'Gm10309',	'Mcfd2',	'Calm2',	'Msh2',	'Msh6',	'Foxn2',	'Gtf2a1l',	'Gm47283',	'Zfp438',	'Epc1',	'Gata6',	'Rbbp8',	'Cables1',	'Npc1',	'Ankrd29',	'Garem1',	'Mapre2',	'Zfp24',	'Rprd1a',	'AW554918',	'Ammecr1l',	'Bin1',	'Camk4',	'Epb41l4a',	'Apc',	'Brd8',	'Ctnna1',	'Lrrtm2',	'Matr3',	'Spata24',	'Dnajc18',	'Nrg2',	'Cystm1',	'Pcdhgc3',	'Pcdhgc5',	'Fgf1',	'Yipf5',	'Prelid2',	'Rbm27',	'Ccdc112',	'Dtwd2',	'Srfbp1',	'Ppic',	'Phax',	'Megf10',	'Ctxn3',	'Slc12a2',	'Chsy3',	'Slc6a7',	'Ppargc1b',	'Afap1l1',	'Sec11c',	'Seh1l',	'Fam210a',	'Mex3c',	'Lipg',	'Rpl17',	'Smad7',	'Smad2',	'Atp5a1',	'Gm6133',	'Setbp1',	'Pard6g',	'Txnl4a',	'Zfp236',	'Tshz1',	'Tmx3',	'Rps6kb2',	'Grk2',	'Rce1',	'Zdhhc24',	'Npas4',	'Banf1',	'Scyl1',	'Frmd8',	'Tigd3',	'Capn1',	'Sac3d1',	'Sf1',	'Macrod1',	'Otub1',	'Rcor2',	'Mark2',	'2700081O15Rik',	'Atl3',	'Bscl2',	'Ganab',	'Syt7',	'Tmem132a',	'Tmem109',	'Mrpl16',	'Mpeg1',	'Dtx4',	'Tle4',	'Gm3443',	'Fxn',	'Pip5k1b',	'Fam122a',	'Tmem252',	'Dock8',	'Kank1',	'Dmrt1',
                         'Plgrkt',	'Pdcd1lg2',	'Ric1',	'Ranbp6',	'Asah2',	'Sgms1',	'Rnls',	'Pank1',	'Btaf1',	'Cpeb3',	'Exoc6',	'Myof',	'Noc3l',	'Pdlim1',	'Sorbs1',	'Aldh18a1',	'Cc2d2b',	'Opalin',	'Slit1',	'Mms19',	'Cyp2c23',	'Cwf19l1',	'Scd2',	'Slf2',	'Sfxn3',	'Btrc',	'Poll',	'Pprc1',	'Cnnm2',	'Gm6970',	'Neurl1a',	'Stn1',	'Cfap58',	'Xpnpep1',	'Mxi1',	'Dusp5',	'Rbm20',	'Acsl5',	'Adrb1',	'Ablim1',	'Trub1',	'Hspa12a',	'Eno4',	'Rab11fip2',	'Csf2ra',	'Rb1cc1',	'Prdm14',	'Rdh10',	'Ube2w',	'Tmem70',	'Paqr8',	'Tram2',	'Kcnq5',	'Smap1',	'Phf3',	'Prim2',	'Fam168b',	'Hs6st1',	'Uggt1',	'Arid5a',	'Kansl3',	'Cnnm4',	'Ankrd23',	'Gm42417',	'Eif5b',	'Tbc1d8',	'Il1r1',	'Il18r1',	'Nab1',	'Nemp2',	'Mfsd6',	'Ankrd44',	'Rftn2',	'Aox1',	'Orc2',	'Tmem237',	'Ccnyl1',	'Kansl1l',	'Atic',	'Smarcal1',	'Cxcr2',	'Vil1',	'Plcd4',	'Cnppd1',	'Tuba4a',	'Asic4',	'Sgpp2',	'Mrpl44',	'Mff',	'A030014E15Rik',	'Dis3l2',	'Ngef',	'Dgkd',	'Ugt1a6b',	'Arl4c',	'Sh3bp4',	'Rab17',	'Lrrfip1',	'Ramp1',	'Ilkap',	'Hes6',	'Ndufa10',	'Kif1a',	'Sned1',	'Mterf4',	'D2hgdh',	'Gin1',	'Pign',	'Tnfrsf11a',	'Dsel',	'Clasp1',	'Tfcp2l1',	'Epb41l5',	'Cd55',	'Gm29427',	'Srgap2',	'Elk4',	'Mfsd4a',	'Ppp1r15b',	'Golt1a',	'Kiss1',	'Sox13',	'Mybph',	'Adora1',	'Rabif',	'Ppp1r12b',	'Rnpep',	'Camsap2',	'Zbtb41',	'Pla2g4a',	'Rnf2',	'Tsen15',	'Mr1',	'Xpr1',	'Cep350',	'Tnr',	'Rc3h1',	'Sft2d2',	'Dcaf6',	'Creg1',	'Mgst3',	'Nos1ap',	'Zbtb18',	'Psen2',	'Stum',	'Lin9',	'Acbd3',	'Lefty2',	'Tmem63a',	'Fbxo28',	'Mia3',	'43892',	'Iars2',	'Fam71a',	'Ints7',	'Dclre1c',	'Prpf18',	'Il15ra',	'Mindy3',	'Rsu1',	'Skida1',	'Spopl',	'Tor4a',	'Ssna1',	'Man1b1',	'Uap1l1',	'Tmem141',	'Camsap1',	'Nacc2',	'Gpsm1',	'Dnlz',	'Agpat2',	'Col5a1',	'Ralgds',	'Cel',	'Ttf1',	'Setx',	'Wdr34',	'Miga2',	'Dolpp1',	'Asb6',	'Abl1',	'Lamc3',	'Prrc2b',	'Swi5',	'Dnm1',	'St6galnac6',	'Cdk9',	'Ttc16',	'Stxbp1',	'Psmd5',	'Phf19',	'Traf1',	'Lhx6',	'Ptgs1',	'Nmi',	'Gpd2',	'Acvr1',	'Ly75',	'Bbs5',	'Ccdc173',	'Gm26558',	'Dcaf17',	'Map3k20',	'Cir1',	'Plekha3',	'Ccdc141',	'Nckap1',	'Rapsn',	'Mybpc3',	'Atg13',	'Pex16',	'Trim44',	'Cd44',	'Pdhx',	'Cat',	'Hipk3',	'Mettl15',	'Eif2ak4',	'Pak6',	'Bahd1',	'Rmdn3',	'Chac1',	'Exd1',	'Vps39',	'Hypk',	'Secisbp2l',	'Cops2',	'Usp50',	'Stard7',	'Slc20a1',	'Snrpb',	'Vps16',	'Spef1',	'Ap5s1',	'Rassf2',	'Slc23a2',	'Cds2',	'Jag1',	'Dstn',	'Rrbp1',	'Rin2',	'Ralgapa2',	'Kiz',	'Apmap',	'Fam110a',	'Defb20',	'H13',	'Asxl1',	'Bpifb1',	
                         'Pxmp4',	'Raly',	'Eif2s2',	'Dynlrb1',	'Trp53inp2',	'Acss2',	'Myh7b',	'Uqcc1',	'Spag4',	'Scand1',	'Cnbd2',	'Tti1',	'Rprd1b',	'Ada',	'Sdc4',	'Pigt',	'Snx21',	'Zfp335',	'Slc35c2',	'Elmo2',	'Eya2',	'Cse1l',	'Ube2v1',	'Ripor3',	'Adnp',	'Gm20721',	'Lsm14b',	'Osbpl2',	'Adrm1',	'Ythdf1',	'Ptk6',	'Helz2',	'Zfp512b',	'Myt1',	'Wdr13',	'Gm14862',	'1810030O07Rik',	'Cask',	'Fundc1',	'Kdm6a',	'Araf',	'Lonrf3',	'Lamp2',	'Rap2c',	'Fmr1',	'Gabrq',	'Trex2',	'Taz',	'Pls3',	'Pdk3',	'Ophn1',	'Efnb1',	'Phka1',	'Magee1',	'Atrx',	'Tbc1d8b',	'Prps1',	'Shroom2',	'Ap1s2',	'Impa1',	'Chmp4c',	'Mtfr1',	'Gyg',	'Tbl1xr1',	'Nceh1',	'Eif5a2',	'Gpr160',	'Zmat3',	'Ttc14',	'Acad9',	'Fgf2',	'Ankrd50',	'Larp1b',	'Noct',	'Spg20',	'Nbea',	'Tm4sf4',	'Igsf10',	'Ssr3',	'Mfsd1',	'Il12a',	'Fnip2',	'Plrg1',	'Gm37240',	'Rps3a1',	'Gm38392',	'Slc25a44',	'Sema4a',	'Rxfp4',	'Rit1',	'Gon4l',	'Msto1',	'Dap3',	'Slc50a1',	'Ube2q1',	'Il6ra',	'Rps27',	'Dennd4b',	'Snapin',	'Tdrkh',	'Cgn',	'Zfp687',	'Golph3l',	'Ensa',	'Ciart',	'Rnf115',	'Pdzk1',	'Chd1l',	'Fmo5',	'Adam30',	'Cd101',	'Atp1a1',	'Hipk1',	'Dclre1b',	'Slc16a1',	'St7l',	'Rap1a',	'I830077J02Rik',	'Dennd2d',	'Lamtor5',	'Gstm6',	'Sort1',	'Mybphl',	'Stxbp3',	'Fndc7',	'Prpf38b',	'Extl2',	'Trmt13',	'Sec24d',	'Gstcd',	'Tet2',	'Cisd2',	'Adh5',	'Rap1gds1',	'Clca2',	'Ctbs',	'Spata1',	'Usp33',	'Srsf11',	'Lrrc40',	'Tmem68',	'Lyn',	'Tox',	'Plekhf2',	'Dpy19l4',	'Osgin2',	'Atp6v0d2',	'Ggh',	'Tstd3',	'Fbxl4',	'Map3k7',	'Ube2j1',	'Ddx58',	'Smim27',	'Bag1',	'Aqp7',	'Ube2r2',	'Dcaf12',	'Fam219a',	'Fam214b',	'Car9',	'Tln1',	'Npr2',	'Olfr71',	'Spaar',	'Zcchc7',	'Slc25a51',	'Coro2a',	'Sec61b',	'Msantd3',	'Ppp3r2',	'Slc44a1',	'Fktn',	'Zfp462',	'Tmem245',	'Zkscan16',	'Ugcg',	'Snx30',	'Atp6v1g1',	'Tmem268',	'Tnfsf15',	'Tnc',	'Rasef',	'Kdm4c',	'Ttc39b',	'Snapc3',	'Focad',	'Atg4c',	'Jak1',	'Slc35d1',	'Oma1',	'Prkaa2',	'Gpx7',	'Btf3l4',	'Osbpl9',	'Ttc39a',	'Bend5',	'Gm12830',	'Efcab14',	'Mob3c',	'Lurap1',	'Plk3',	'Rps8',	'Atp6v0b',	'Dph2',	'Ipo13',	'Artn',	'St3gal3',	'Szt2',	'AU022252',	'Rimkla',	'Slfnl1',	'Ctps',	'Cap1',	'Macf1',	'Ndufs5',	'Inpp5b',	'Yrdc',	'Zc3h12a',	'Stk40',	'Thrap3',	'Tekt2',	'AU040320',	'Tmem35b',	'Azin2',	'Rnf19b',	'Zbtb8a',	'Dcdc2b',	'Ptp4a2',	'Pef1',	'Hcrtr1',	'Tmem200b',	'Ptafr',	'Slc9a1',	'Pigv',	'Sh3bgrl3',	'Slc30a2',	'Mtfr1l',	'Selenon',	'Clic4',	'Pnrc2',	'Gale',	'Cdc42',	'Ece1',	'Hp1bp3',	'Pink1',	'Camk2n1',	'Otud3',	
                         'Epha2',	'Spen',	'Tmem82',	'Plekhm2',	'Agmat',	'Tnfrsf1b',	'Fv1',	'Mfn2',	'2510039O18Rik',	'Mthfr',	'Fbxo6',	'Fbxo2',	'Ubiad1',	'Pgd',	'Clstn1',	'Slc25a33',	'Spsb1',	'H6pd',	'Rere',	'Park7',	'Dnajc11',	'Thap3',	'Phf13',	'Klhl21',	'Plekhg5',	'Kcnab2',	'Dffb',	'Lrrc47',	'Smim1',	'Tprgl',	'Tnfrsf14',	'Gabrd',	'Gnb1',	'Mxra8',	'B3galt6',	'Ankib1',	'Cldn12',	'Cfap69',	'Gnat3',	'Rsbn1l',	'Gsap',	'Napepld',	'Tomm7',	'Atg9b',	'Tmub1',	'Asb10',	'Nub1',	'Kmt2c',	'Paxip1',	'Rbm33',	'Ost4',	'Khk',	'Gm29609',	'Mrpl33',	'Rbks',	'Nelfa',	'Zfyve28',	'Fam193a',	'Mfsd10',	'Msantd1',	'Acox3',	'Sh3tc1',	'Psapl1',	'Tada2b',	'Man2b2',	'Zbtb49',	'Slc2a9',	'Wdr1',	'Cd38',	'Sel1l3',	'Rell1',	'Tlr1',	'Tmem156',	'Dcun1d4',	'Chic2',	'Clock',	'Rufy3',	'Ankrd17',	'Btc',	'Rchy1',	'Cdkl2',	'Paqr3',	'Nkx6-1',	'Wdfy3',	'1700016H13Rik',	'Nudt9',	'Gm17660',	'Zfp326',	'Fam69a',	'Cplx1',	'Gak',	'Idua',	'Chfr',	'Pgam5',	'Pxmp2',	'Galnt9',	'Noc4l',	'Ddx51',	'Ep400',	'Tpst2',	'Coro1c',	'Kctd10',	'Ube3b',	'Ankrd13a',	'Oasl1',	'Rnf10',	'Gatc',	'Rab35',	'Prkab1',	'Suds3',	'Taok3',	'Fbxo21',	'Tpcn1',	'Hectd4',	'Aldh2',	'Atxn2',	'Sh2b3',	'Myl2',	'Ift81',	'Rnf34',	'Diablo',	'Gm49027',	'Clip1',	'Zcchc8',	'Hip1r',	'Vps37b',	'Kmt5a',	'Snrnp35',	'Rilpl1',	'Zfp664',	'Scarb1',	'Glt1d1',	'Nupr1l',	'Tpst1',	'Kctd7',	'Rabgef1',	'Tmem248',	'Lat2',	'Eif4h',	'Stx1a',	'Tmem120a',	'Srrm3',	'Orai2',	'Gm20605',	'Sap25',	'Irs3',	'Cyp3a13',	'Mblac1',	'Sun1',	'Gpr146',	'Gper1',	'Mafk',	'Snx8',	'Ap5z1',	'Papolb',	'Rac1',	'Cyth3',	'Usp42',	'Ccz1',	'Ocm',	'Smurf1',	'Arpc1b',	'Zkscan14',	'Zfp655',	'Wasf3',	'Lnx2',	'Slc7a1',	'Ubl3',	'Gm42791',	'N4bp2l1',	'Casd1',	'C1galt1',	'Tspan12',	'Cadps2',	'Wasl',	'Rbm28',	'Calu',	'Kcp',	'Ube2h',	'Klf14',	'Akr1b7',	'Stra8',	'Fam180a',	'Zc3hav1',	'Ubn2',	'Clec2l',	'Tra2a',	'Fam221a',	'Osbpl3',	'Hibadh',	'Tax1bp1',	'Cpvl',	'Prr15',	'Plekha8',	'Znrf2',	'Nod1',	'Gng12',	'Krcc1',	'Rnf103',	'Kdm3a',	'Reep1',	'Rnf181',	'Vamp8',	'Mat2a',	'Sh2d6',	'Suclg1',	'Mrpl19',	'Dok1',	'Loxl3',	'Mogs',	'Dctn1',	'Mob1a',	'Bola3',	'Smyd5',	'Add2',	'Pcbp1',	'Asprv1',	'Mxd1',	'Snrnp27',	'Gmcl1',	'Aak1',	'Rab43',	'Cnbp',	'Rab7',	'Rpn1',	'Sec61a1',	'Abtb1',	'Chchd6',	'Txnrd3',	'Zxdc',	'Klf15',	'Aldh1l1',	'Slc41a3',	'Nup210',	'Hdac11',	'Fbln2',	'Tmem43',	'Xpc',	'Nr2c2',	'Suclg2',	'Gxylt2',	'Edem1',	'Thumpd3',	'Lhfpl4',	'Prrt3',	'Irak2',	'Tatdn2',	'Tamm41',	'Mkrn2',
                         'Raf1',	'Tmem40',	'Rpl32',	'43898',	'Rasgef1a',	'Adipor2',	'Il17ra',	'Foxj2',	'Chd4',	'Nop2',	'Ltbr',	'Plekhg6',	'Tspan9',	'Fkbp4',	'Lrp6',	'Crebl2',	'Strap',	'Recql',	'Ldhb',	'Lrmp',	'Bhlhe41',	'Ccdc91',	'Amn1',	'Cacng7',	'Leng9',	'Tnnt1',	'Ppp6r1',	'Shisa7',	'Zfp580',	'Zfp787',	'Selenow',	'Nop53',	'Ccdc9',	'Bbc3',	'Zc3h4',	'Slc1a5',	'Ppp5c',	'Opa3',	'Vasp',	'Ercc2',	'Mark4',	'Smg9',	'Arhgef1',	'Grik5',	'Dedd2',	'Ceacam1',	'Tmem91',	'Tgfb1',	'Egln2',	'Numbl',	'Sptbn4',	'Akt2',	'Med29',	'Catsperg1',	'Dpf1',	'Sdhaf1',	'Scn1b',	'Lsm14a',	'Faap24',	'Pdcd5',	'Dpy19l3',	'Vsig10l',	'Mybpc2',	'Spib',	'Kcnc3',	'Ap2a1',	'Fcgrt',	'Aldh16a1',	'Gys1',	'Bcat2',	'Rasip1',	'Kdelr1',	'Ccdc114',	'Sergef',	'Uevld',	'Gas2',	'Nipa2',	'Herc2',	'Klf13',	'E030018B13Rik',	'Fan1',	'Lysmd4',	'Aen',	'Hapln3',	'Pex11a',	'Zfp710',	'Fes',	'Slc28a1',	'Rps17',	'Cemip',	'Abhd17c',	'Fah',	'Picalm',	'Ccdc83',	'Kctd21',	'Alg8',	'Ndufc2',	'Ints4',	'Acer3',	'Mogat2',	'Arrb1',	'Tpbgl',	'Ucp3',	'Ucp2',	'Clpb',	'Il18bp',	'Nup98',	'Trim6',	'Smpd1',	'Apbb1',	'Hpx',	'Trim3',	'Rpl27a',	'Nrip3',	'Dennd5a',	'Tmem41b',	'Ipo7',	'Swap70',	'Mrvi1',	'Gm4353',	'Rps13',	'Pik3c2a',	'Arl6ip1',	'Smg1',	'Coq7',	'Gde1',	'Gprc5b',	'Lyrm1',	'Abca14',	'Usp31',	'Lcmt1',	'Il4ra',	'D430042O09Rik',	'Gsg1l',	'Atxn2l',	'Sult1a1',	'Tmem219',	'Kctd13',	'Zfp768',	'Ctf1',	'Kat8',	'Armc5',	'BC017158',	'Bag3',	'Inpp5f',	'Fgfr2',	'Nsmce4a',	'Fam53b',	'9430038I01Rik',	'Ppp2r2d',	'Zfp941',	'Irf7',	'Tmem80',	'Pnpla2',	'Mob2',	'Dusp8',	'Kcnq1',	'Slc22a18',	'Osbpl5',	'Ppfia1',	'Mrgprf',	'Cnksr3',	'Ginm1',	'Utrn',	'Ltv1',	'Reps1',	'Arfgef3',	'Il22ra2',	'Map3k5',	'Pde7b',	'Ahi1',	'Rnf146',	'Tspyl1',	'Tspyl4',	'Mfsd4b4',	'Mfsd4b5',	'Armc2',	'Sec63',	'Scml4',	'Bend3',	'Gm9803',	'Ascc3',	'Mcm9',	'Fam184a',	'Msl3l2',	'Lims1',	'Edar',	'44084',	'Mcu',	'Eif4ebp2',	'Lrrc20',	'Aifm2',	'Hk1',	'4930507D05Rik',	'Ccar1',	'Mypn',	'A930033H14Rik',	'Tfam',	'Bcr',	'Upb1',	'Susd2',	'Gstt1',	'Smarcb1',	'S100b',	'Pcnt',	'Lss',	'1810043G02Rik',	'Trappc10',	'Agpat3',	'Cdc34',	'Bsg',	'Misp',	'Tmem259',	'Abca7',	'Arhgap45',	'Polr2e',	'Gpx4',	'Dazap1',	'Rps15',	'2310011J03Rik',	'Adamtsl5',	'Mknk2',	'Ap3d1',	'Dot1l',	'Plekhj1',	'Gng7',	'4930442H23Rik',	'Hmg20b',	'Smim24',	'Nfic',	'Aes',	'Glt8d2',	'Tcp11l2',	'Polr3b',	'Nt5dc3',	'Parpbp',	'Scyl2',	'Uhrf1bp1l',	'Hal',	'Nr2c1',	'Tmcc3',	'Zdhhc17',	'Atxn7l3b',	'Tbc1d15',	'Rab21',
                         'Wif1',	'Gns',	'Tbk1',	'Agap2',	'B4galnt1',	'Itga7',	'Stxbp2',	'Lig4',	'Naxd',	'Cars2',	'Tubgcp3',	'Atp11a',	'Adprhl1',	'Dcun1d2',	'Tmem255b',	'Gas6',	'Rasa3',	'Champ1',	'Cln8',	'Angpt2',	'Atp7b',	'Nkx6-3',	'Adam32',	'Plpp5',	'Hook3',	'Brf2',	'Got1l1',	'Tti2',	'Mak16',	'Wrn',	'Smim18',	'Pdgfrl',	'Snx25',	'Cfap97',	'Wwc2',	'Lzts1',	'Nr2c2ap',	'Ell',	'Rab3a',	'Ccdc124',	'Use1',	'Pgls',	'Med26',	'Pkn1',	'Ptger1',	'Nanos3',	'Mri1',	'Dnase2a',	'Hook2',	'Vps35',	'Itfg1',	'Siah1a',	'N4bp1',	'Brd7',	'Chd9',	'Rpgrip1l',	'Herpud1',	'Nlrc5',	'Cpne2',	'Cx3cl1',	'Mmp15',	'Ccdc113',	'Terb1',	'Cbfb',	'D230025D16Rik',	'Nol3',	'4931428F04Rik',	'Exoc3l',	'Lrrc29',	'Fhod1',	'Hsd11b2',	'Ripor1',	'Nfatc3',	'Cyb5b',	'Ap1g1',	'Marveld3',	'Ldhd',	'Zfp1',	'Bcar1',	'Tmem231',	'Terf2ip',	'Syce1l',	'Dynlrb2',	'Cdyl2',	'Hsbp1',	'Mlycd',	'Hsdl1',	'Klhl36',	'Gse1',	'Map1lc3b',	'Zcchc14',	'Piezo1',	'Pabpn1l',	'Ankrd11',	'Spata2l',	'Abcb10',	'Capn9',	'Gnpat',	'Map10',	'Coa6',	'Itgb1',	'Pxk',	'Gm49355',	'Ngly1',	'Ube2e1',	'Ecd',	'Usp54',	'Fut11',	'Vcl',	'Lrmda',	'Ppif',	'Anxa11',	'Wnt5a',	'Cacna1d',	'Tkt',	'Prkcd',	'Rft1',	'Sfmbt1',	'Nisch',	'Phf7',	'Gprin2',	'Fam25c',	'Opn4',	'Wapl',	'Ccser2',	'Tspan14',	'Gmfb',	'Atg14',	'Ktn1',	'Ppp1r3e',	'Bcl2l2',	'Myh6',	'Nrl',	'Tm9sf1',	'Dhrs1',	'Khnyn',	'Cenpj',	'Gm6904',	'Cdadc1',	'Ebpl',	'Ctsb',	'Fdft1',	'Fam167a',	'Ints9',	'Nuggc',	'9930012K11Rik',	'Ppp3cc',	'Lgi3',	'Fam160b2',	'Fndc3a',	'Lrch1',	'Tsc22d1',	'Dgkh',	'Vwa8',	'Diaph3',	'Acod1',	'Cln5',	'Dzip1',	'Gpr18',	'Tm9sf2',	'Tmem123',	'Maml2',	'Amotl1',	'Fut4',	'Zfp426',	'S1pr2',	'Pde4a',	'Ilf3',	'Dnm2',	'Kank2',	'Zfp872',	'Zfp810',	'Herpud2',	'Adamts8',	'Zbtb44',	'Prdm10',	'Esam',	'Nectin1',	'Bcl9l',	'Ddx6',	'Usp28',	'Ncam1',	'Plet1',	'Dlat',	'Dixdc1',	'Fdx1',	'Rdx',	'Zc3h12c',	'Acat1',	'Dmxl2',	'Idh3a',	'Dnaja4',	'Wdr61',	'Snx33',	'Imp3',	'Ptpn9',	'Cox5a',	'Tmem202',	'Fem1b',	'Dennd4a',	'Hacd3',	'Dpp8',	'Rasl12',	'Mtfmt',	'Zfp609',	'Gcnt3',	'Ccpg1os',	'Ccpg1',	'Khdc3',	'Fam214a',	'Elovl5',	'Ick',	'Slc17a5',	'Myo6',	'Tbc1d2b',	'Plscr5',	'Xrn1',	'Tfdp2',	'Zbtb38',	'Ryk',	'Rab6b',	'Cdv3',	'Nudt16',	'Ppm1m',	'Alas1',	'Dusp7',	'Abhd14b',	'Parp3',	'Rrp9',	'Tex264',	'Mapkapk3',	'Cish',	'Hemk1',	'Zmynd10',	'Tusc2',	'Rbm6',	'Mst1',	'Apeh',	'Gpx1',	'P4htm',	'Prkar2a',	'Celsr3',	'Shisa5',	'Plxnb1',	'Dhx30',	'Cspg5',	'Scap',	'Ptpn23',	'Nradd',	'Ccdc12',	'Rtp3',	
                         'Lrrfip2',	'Clasp2',	'Dync1li1',	'Itga9',	'Acaa1a',	'Xylb',	'Gorasp1',	'Trak1',	'Higd1a',	'Ano10',	'Slc6a20a',	'Fyco1',	'Patz1',	'Limk2',	'Rnf185',	'Morc2a',	'Sec14l4',	'Mtfp1',	'Sec14l2',	'Ascc2',	'Uqcr10',	'Ap1b1',	'Gas2l1',	'Dbnl',	'Gck',	'Ykt6',	'Ogdh',	'Purb',	'Fbxo48',	'Cep68',	'Sertad2',	'Zrsr1',	'Efemp1',	'Psme4',	'Erlec1',	'Cpeb4',	'Mpg',	'Ccng1',	'Adra1b',	'Rnf145',	'Rack1',	'Gm12184',	'Olfr1393',	'Rnf130',	'Sqstm1',	'Mgat4b',	'Col23a1',	'Sec24a',	'Sar1b',	'Ube2b',	'Cdkl3',	'Ppp2ca',	'Hspa4',	'Aff4',	'Gm9945',	'Kif3a',	'Irf1',	'P4ha2',	'Acsl6',	'Fnip1',	'Cdc42se2',	'Tnip1',	'Fam114a2',	'Irgm2',	'4930438A08Rik',	'Zfp672',	'Trim11',	'Iba57',	'2310033P09Rik',	'Arf1',	'Myo15',	'Mief2',	'Tmem11',	'Natd1',	'Map2k3',	'Aldh3a1',	'Slc47a2',	'Epn2',	'Ulk2',	'Akap10',	'Tekt3',	'Hs3st3a1',	'Elac2',	'1700086D15Rik',	'Myocd',	'Zkscan6',	'Gas7',	'Stx8',	'Ndel1',	'Per1',	'Cntrob',	'Tmem102',	'Gps2',	'Slc16a11',	'Med11',	'Inca1',	'Zfp3',	'Pitpnm3',	'Mybbp1a',	'Cyb5d2',	'Atp2a3',	'Camkk1',	'Shpk',	'Mettl16',	'Mnt',	'Sgsm2',	'Hic1',	'Rtn4rl1',	'Wdr81',	'Rilp',	'Scarf1',	'Slc43a2',	'Doc2b',	'Vps53',	'Dbil5',	'Sdf2',	'Poldip2',	'Nos2',	'Lgals9',	'Ccl8',	'Mmp28',	'Ccl5',	'Heatr6',	'Ppm1d',	'Rnft1',	'Tubd1',	'Vmp1',	'Cltc',	'Gdpd1',	'Trim25',	'Gm45716',	'Stxbp4',	'Cox11',	'Spag9',	'Wfikkn2',	'Cacna1g',	'Spata20',	'Mycbpap',	'Rsad1',	'Slc35b1',	'B130006D01Rik',	'Phospho1',	'Snx11',	'Cdk5rap3',	'Pnpo',	'Tbkbp1',	'Npepps',	'Mrpl45',	'Gpr179',	'Arhgap23',	'Cwc25',	'B230217C12Rik',	'Fbxl20',	'Med1',	'Stard3',	'Tcap',	'Erbb2',	'Ormdl3',	'Med24',	'Thra',	'Krt42',	'Nkiras2',	'Stat5a',	'Stat3',	'Naglu',	'Retreg3',	'Dhx8',	'Hdac5',	'Slc25a39',	'Hexim1',	'Plekhm1',	'Nsf',	'Ace',	'Map3k3',	'Limd2',	'Gm10840',	'Strada',	'Ccdc47',	'Cd79b',	'Polg2',	'Helz',	'Cacng1',	'Gna13',	'9930022D16Rik',	'Cdc42ep4',	'Cdr2l',	'Mrpl58',	'Mif4gd',	'Llgl2',	'Myo15b',	'Smim5',	'Unk',	'Unc13d',	'Mrpl38',	'Srp68',	'Ube2o',	'Rhbdf2',	'Mettl23',	'Sec14l1',	'6030468B19Rik',	'Cant1',	'C1qtnf1',	'Sgsh',	'Rnf213',	'Endov',	'Chmp6',	'Ndufaf8',	'Faap100',	'Gcgr',	'Pcyt2',	'Ccdc57',	'Csnk1d',	'Metrnl',	'Gdi2',	'Pfkp',	'Larp4b',	'Lgals8',	'Ggps1',	'Cdk13',	'Yae1d1',	'Stard3nl',	'BC005537',	'Acot13',	'Fam50b',	'Prpf4b',	'Rpp40',	'Slc35b3',	'Gfod1',	'Nol7',	'Ranbp9',	'Mcur1',	'Gmpr',	'Phf2',	'Card19',	'Tes3-ps',	'Iars',	'Auh',	'Arl10',	'Tspan17',	'Grk6',	'H2afy',	'Tifab',	'Klhl3',
                         '2210016F16Rik',	'Ell2',	'Ttc37',	'Slf1',	'Cetn3',	'Rasa1',	'Gm20379',	'Zfyve16',	'Ap3b1',	'Pde8b',	'Col4a3bp',	'Tnpo1',	'Ptcd2',	'Serf1',	'Erbin',	'Kif2a',	'Il6st',	'BC067074',	'Emb',	'Paip1',	'Nim1k',	'Rab10',	'Kif3c',	'Ncoa1',	'Klhl29',	'Ldah',	'Hs1bp3',	'Pum2',	'Laptm4a',	'Ttc32',	'Smc6',	'Pqlc3',	'Cpsf3',	'Iah1',	'Adam17',	'Taf1b',	'Mboat2',	'Kidins220',	'Rnf144a',	'Tpo',	'Ccdc71l',	'Nampt',	'Snx13',	'Agmo',	'Arl4a',	'Zfp277',	'Dnajb9',	'Sptssa',	'Ppp2r3c',	'Rpl36al',	'Klhdc2',	'Vcpkmt',	'Sos2',	'Abhd12b',	'Actr10',	'Arid4a',	'Zbtb25',	'Gphn',	'Plekhh1',	'Psen1',	'Pnma1',	'Elmsan1',	'Rnf113a2',	'Aldh6a1',	'Zc2hc1c',	'Tmed10',	'Erg28',	'Adck1',	'Rin3',	'Lgmn',	'Chga',	'Itpk1',	'Bdkrb2',	'Gskip',	'Ccdc85c',	'Slc25a47',	'Wdr25',	'Ppp2r5c',	'Zfp839',	'Cinp',	'Tecpr2',	'Traf3',	'Mark3',	'Trmt61a',	'Bag5',	'Ppp1r13b',	'Zbtb42',	'Tex22',	'Mta1',	'Crip1',	'Fbxo4',	'Oxct1',	'Rictor',	'Egflam',	'Gdnf',	'Dnajc21',	'Zfp622',	'Fbxl7',	'Dap',	'Vps13b',	'Spag1',	'Rrm2b',	'Atp6v1c1',	'Eif3e',	'Eny2',	'Col14a1',	'Derl1',	'9130401M01Rik',	'Fam91a1',	'Rnf139',	'Washc5',	'Trib1',	'Efr3a',	'Ago2',	'Dennd3',	'Zc3h3',	'Tigd5',	'Pycrl',	'Scrib',	'Nrbp2',	'BC024139',	'Plec',	'Parp10',	'Smpd5',	'Hgh1',	'Bop1',	'Fbxl6',	'Kifc2',	'Ppp1r16a',	'1110038F14Rik',	'Apol6',	'Apol8',	'Triobp',	'Micall1',	'Polr2f',	'Pla2g6',	'Tmem184b',	'Kdelr3',	'Sun2',	'Dnal4',	'Apobec3',	'Polr3h',	'Desi1',	'Naga',	'Ndufa6',	'Cyp2d22',	'Pacsin2',	'Ttll1',	'Bik',	'Mcat',	'Ttll12',	'5031439G07Rik',	'Ppara',	'Alg12',	'Creld2',	'Plxnb2',	'C730034F03Rik',	'Mapk8ip2',	'Gxylt1',	'Pus7l',	'Twf1',	'Endou',	'Rapgef3',	'Slc48a1',	'Hdac7',	'Asb8',	'Cacnb3',	'Ccdc65',	'Kmt2d',	'Tmbim6',	'Faim2',	'Aqp2',	'Smarcd1',	'Gpd1',	'Gm5475',	'Fignl2',	'Krt80',	'Tns2',	'Zfp740',	'Itga5',	'Gtsf1',	'Dnaja3',	'Hmox2',	'4930562C15Rik',	'Ubald1',	'Mgrn1',	'Nudt16l1',	'Gm5767',	'Litaf',	'Zc3h7a',	'Rsl1d1',	'Cpped1',	'Parn',	'Pla2g10',	'Pdxdc1',	'Mapk1',	'Ppil2',	'Scarf2',	'Dgcr6',	'Prodh',	'Hira',	'Camk2n2',	'Ehhadh',	'Tmem41a',	'Liph',	'Atp13a4',	'Pak2',	'Pcyt1a',	'Fyttd1',	'Heg1',	'Adcy5',	'Sec22a',	'Iqcb1',	'Lrrc58',	'Atp6v1a',	'Usf3',	'Spice1',	'Slc35a5',	'Cd47',	'Cep97',	'Zbtb11',	'Olfr172',	'Riox2',	'Usp25',	'App',	'Map3k7cl',	'Tiam1',	'Sod1',	'Eva1c',	'Synj1',	'Paxbp1',	'Ifnar2',	'Ifnar1',	'Ifngr2',	'Tmem50b',	'Dnajc28',	'Itsn1',	'Clic6',	'Brwd1',	'Ripk4',	'Zdhhc14',	'Serac1',	'Sytl3',
                         'Ezr',	'AC122413.2',	'Sft2d1',	'Prr18',	'Igf2r',	'Acat3',	'Wtap',	'Gm3448',	'Chd1',	'Lnpep',	'Zfp948',	'Zfp943',	'Pdpk1',	'Amdhd2',	'Tbc1d24',	'Caskin1',	'Nthl1',	'Eme2',	'Tekt4',	'Rab40c',	'Capn15',	'Tmem8',	'Atp6v0e',	'Crebrf',	'Bnip1',	'AI413582',	'Taf11',	'Zfp523',	'Ppard',	'Fkbp5',	'Mapk14',	'Kctd20',	'Stk38',	'BC004004',	'Gm28043',	'Slc37a1',	'Wdr4',	'Rrp1b',	'Cyp4f13',	'Adamts10',	'Kank3',	'Ndufa7',	'H2-K1',	'Tap1',	'Gm15821',	'Notch4',	'Pbx2',	'Atf6b',	'Neu1',	'H2-D1',	'Atat1',	'Trim15',	'Rnf39',	'Cyp39a1',	'Enpp4',	'Tmem63b',	'Gtpbp2',	'Polr1c',	'Zfp318',	'Cul9',	'1700001C19Rik',	'Tomm6',	'Kif6',	'Tbc1d5',	'Zfp119b',	'Stap2',	'Sema6b',	'Dpp9',	'Nrtn',	'Prr22',	'Rfx2',	'1700061G19Rik',	'Mllt1',	'Vav1',	'Fbxl17',	'A930002H24Rik',	'Pja2',	'Ppp4r1',	'Twsg1',	'Ankrd12',	'Washc1',	'Mtcl1',	'Emilin2',	'Clip4',	'Ypel5',	'Xdh',	'Ttc27',	'Fam98a',	'Strn',	'Prkd3',	'Atl2',	'Hnrnpll',	'Sos1',	'Cox7a2l',	'Dync2li1',	'Prkce',	'Ttc7',	'Ston1',	'Cul2',	'Arhgap12',	'Kif5b',	'Mpp7',	'Wac',	'Usp14',	'Rock1',	'Mib1',	'Osbpl1a',	'Ss18',	'Kctd1',	'Rnf138',	'Sft2d3',	'Reep5',	'Cdc23',	'Paip2',	'Ube2d2a',	'Cxxc5',	'Pcdhga2',	'Pcdhgb5',	'Pcdhga10',	'Rnf14',	'Trim36',	'Snx2',	'Csnk1g3',	'Synpo',	'Rps14',	'Hmgxb3',	'Pde6a',	'Arhgef37',	'Pcyox1l',	'Grpel2',	'Ablim3',	'Fbxo38',	'Fech',	'Nars',	'Cidea',	'Afg3l2',	'Psmg2',	'Ldlrad4',	'Mbd2',	'Ctif',	'Hdhd2',	'Adnp2',	'Pqlc1',	'Ctdp1',	'Mbp',	'Cndp2',	'Cyb5a',	'Cpt1a',	'Tesmin',	'Lrp5',	'Chka',	'Unc93b1',	'Cdk2ap2',	'Coro1b',	'Pold4',	'Rhod',	'2010003K11Rik',	'Lrfn4',	'Rin1',	'Snx32',	'Ap5b1',	'Kcnk7',	'Ehbp1l1',	'Sssca1',	'Ltbp3',	'Majin',	'Cdc42bpg',	'Slc22a12',	'Gpr137',	'Cox8a',	'Slc3a2',	'Lrrn4cl',	'B3gat3',	'Rom1',	'Fth1',	'Best1',	'Myrf',	'Cpsf7',	'Vps37c',	'Ms4a8a',	'Stx3',	'Carnmt1',	'D030056L22Rik',	'Trpm6',	'Klf9',	'Smarca2',	'Vldlr',	'Glis3',	'Rcl1',	'Jak2',	'Mlana',	'Uhrf2',	'Pcgf5',	'43895',	'Lcor',	'Rrp12',	'Exosc1',	'Ankrd2',	'Pi4k2a',	'Sfrp5',	'R3hcc1l',	'Pyroxd2',	'Got1',	'Erlin1',	'Bloc1s2',	'Lzts2',	'Gm17018',	'Fbxw4',	'Mgea5',	'Hps6',	'Ldb1',	'Sufu',	'Trim8',	'Wbp1l',	'Cyp17a1',	'Calhm2',	'Sh3pxd2a',	'Vti1a',	'Casp7',	'Nhlrc2',	'Ccdc186',	'Vwa2',	'Fam160b1',	'Atrnl1',	'Shtn1',	'Pdzd8',	'Cacul1',	'Zfp950',	'Vcpip1',	'Wdr75',	'Tmeff2',	'Gls',	'Hecw2',	'Cdk15',	'Klf7',	'Plekhm3',	'Map2',	'Scg2',	'Sp100',	'Klhl30',	'Per2',	'Tmem185b',	'Sctr',	'Klhdc8a',	'Rgs13',	'Nmnat2',	
                         'Tnfsf18',	'Pigc',	'Tada1',	'Atf6',	'Tstd1',	'Atp1a4',	'Capn8',	'Celf2',	'Cubn',	'H2al2a',	'A930004D18Rik',	'Lcn4',	'Brd3',	'Slc2a8',	'Gtdc1',	'Lypd6',	'Gcg',	'Mtx2',	'Ttn',	'Syt13',	'Ext2',	'Rnf24',	'Lrrn4',	'Ism1',	'Slc24a3',	'Snph',	'Mroh8',	'D630003M21Rik',	'Ptprt',	'Wfdc13',	'Wfdc3',	'Ptgis',	'Fam210b',	'Fam209',	'Tpd52l2',	'Bmp15',	'Dgkk',	'Suv39h1',	'Xiap',	'Stag2',	'Plxnb3',	'Zmym3',	'Abcb7',	'Fgf16',	'Pcdh19',	'Map3k15',	'Pir',	'Lrrcc1',	'Car2',	'Gm9733',	'Intu',	'Golim4',	'Mnd1',	'Efna1',	'Tchhl1',	'Car14',	'Otud7b',	'Hsd3b2',	'Man1a2',	'Slc22a15',	'Mov10',	'Kcnd3',	'Fabp2',	'Gm5105',	'Bcl10',	'Sdcbp',	'Necab1',	'Ccnc',	'Epb41l4b',	'Pakap',	'Slc31a1',	'Zdhhc21',	'Mllt3',	'Gm10306',	'Patj',	'Kank4',	'Trabd2b',	'Olfr1342',	'Nfyc',	'Trit1',	'Zbtb8b',	'Phactr4',	'6030445D17Rik',	'Cda',	'Pqlc2',	'Iffo2',	'Tardbp',	'Mterf1b',	'Kmt2e',	'Tmem214',	'Tcf23',	'Depdc5',	'Rnf4',	'Rbpj',	'Wdr19',	'Bend4',	'Fip1l1',	'Kit',	'Nmu',	'Grsf1',	'Lin54',	'Mn1',	'Trpv4',	'Rasal1',	'Camkk2',	'Zbed5',	'Fam20c',	'Get4',	'Zfp853',	'Baiap2l1',	'Hsph1',	'Phf14',	'Tmem168',	'Asb15',	'Tmem209',	'Slc23a4',	'Gimap3',	'Crhr2',	'Ccser1',	'Prdm5',	'Il23r',	'Sfxn5',	'Prokr1',	'Eefsec',	'Rho',	'Ankrd26',	'Wnk1',	'Tuba3a',	'Hebp1',	'Spx',	'Rassf8',	'Ppfibp1',	'Rpl28',	'Arhgap35',	'Zfp658',	'Tjp1',	'Lins1',	'Sv2b',	'Anpep',	'Alpk3',	'Tm6sf1',	'Saxo2',	'Me3',	'Tenm4',	'Gdpd5',	'Neu3',	'Ctr9',	'Xylt1',	'Cdr2',	'Dmbt1',	'Acadsb',	'Ctbp2',	'6430531B16Rik',	'Lsp1',	'Tab2',	'Olig3',	'Stx7',	'Mical1',	'Ranbp2',	'Pald1',	'Gm10118',	'Fam13c',	'Gzmm',	'Igf1',	'Ccdc59',	'Best3',	'Cct2',	'9530003J23Rik',	'Slc35e3',	'Ppm1h',	'Tcim',	'Hgsnat',	'Dctn6',	'Dusp4',	'Mfhas1',	'Tusc3',	'Acsl1',	'Nwd1',	'Ednra',	'1700011L22Rik',	'Gm10645',	'Clgn',	'Palm3',	'4930432K21Rik',	'Zfp423',	'Aktip',	'Nup93',	'Sntb2',	'Znrf1',	'Atmin',	'Cdh15',	'Nup133',	'Agt',	'Ptprg',	'Rarb',	'Rps24',	'Erc2',	'Ptpn20',	'Zfp488',	'Cgrrf1',	'Olfr221',	'Mettl3',	'Ajuba',	'Atp12a',	'Xpo4',	'Rhobtb2',	'Xpo7',	'Ccdc122',	'Akap11',	'Cnmd',	'Arhgap42',	'Jrkl',	'Smco4',	'Arhgap32',	'Kcnj1',	'Fdxacb1',	'Atm',	'Ube2q2',	'Odf3l1',	'Neo1',	'Lctl',	'Megf11',	'Slc24a1',	'Gm10642',	'Cgnl1',	'Pls1',	'Iqcf5',	'Ccdc71',	'Eomes',	'Zfp651',	'Tbc1d10a',	'Adcy1',	'Tns3',	'Ppp3r1',	'Actr2',	'Eml6',	'Tenm2',	'Fstl4',	'Igtp',	'Prss38',	'Cenpv',	'Cdrt4',	'Map2k4',	'Usp43',	'Cfap52',	'Ccdc42',	'Slc46a1',	'Pex12',	
                         'Ccl6',	'Gm11492',	'Mpo',	'Mmd',	'Chad',	'Pdk2',	'Hoxb1',	'Skap1',	'Cdk12',	'Tubg2',	'C1ql1',	'Cacng5',	'Rgs9',	'Abca9',	'Asb13',	'Wdr37',	'Hecw1',	'Aoah',	'Hist1h4k',	'Hist1h2ak',	'Ripor2',	'Serpinb1c',	'Fars2',	'Adtrp',	'Fam8a1',	'Fgd3',	'Rab24',	'Jmy',	'S100z',	'Foxd1',	'Gm10320',	'Ocln',	'Ipo11',	'Gapt',	'Mocs2',	'Mfsd2b',	'Ispd',	'Arhgap5',	'Snx6',	'Cdkl1',	'Atl1',	'Ppp1r36',	'Numb',	'Riox1',	'Dnal1',	'Ltbp2',	'Ahsa1',	'Gm28051',	'Serpina16',	'Evl',	'Sp4',	'Lmbrd2',	'Ank',	'Sdc2',	'Ncald',	'Angpt1',	'Rspo2',	'Mtbp',	'Has2',	'Gm6569',	'Serhl',	'Adamts20',	'Rarg',	'Myh11',	'Cldn1',	'Zfp148',	'Ccdc14',	'Ildr1',	'Ndufb4',	'Cd96',	'Zpld1',	'Filip1l',	'Snx9',	'Lix1',	'Gm38655',	'Abcg1',	'Gm8909',	'2410137M14Rik',	'Ttbk1',	'Bysl',	'Mocs1',	'Lbh',	'Rasgrp3',	'Cabyr',	'Impact',	'Rnf125',	'2700062C07Rik',	'Tnfaip8',	'Hsd17b4',	'Zfp474',	'Smim3',	'Apcdd1',	'Chmp1b',	'Spire1',	'Hsbp1l1',	'Pcsk5',	'1700028P14Rik',	'Ptar1',	'Cd274',	'Mbl2',	'Prkg1',	'Cstf2t',	'Ifit2',	'Ide',	'Fra10ac1',	'Gbf1',	'Adra2a',	'Prex2',	'Efhc1',	'Lman2l',	'Mgat4a',	'Npas2',	'Stat1',	'Gtf3c3',	'Als2cr12',	'Nbeal1',	'Gpbar1',	'Kcne4',	'Nyap2',	'Agfg1',	'Rbm44',	'Traf3ip1',	'Fcmr',	'Tmcc2',	'Dstyk',	'Zfp281',	'Nek7',	'Axdnd1',	'Fmo6',	'Mael',	'Tagln2',	'Susd4',	'Cdc123',	'Gm13199',	'Il2ra',	'Pip4k2a',	'Gad2',	'Ehmt1',	'Sohlh1',	'Rpl12',	'Cntrl',	'Gpr21',	'Dennd1a',	'Fastkd1',	'Sestd1',	'Nup160',	'Psmc3',	'F2',	'Ambra1',	'Alkbh3',	'Ttc17',	'Srp14',	'Inafm2',	'Haus2',	'Ciao1',	'Ttl',	'Stk35',	'Tgm3',	'Slc52a3',	'Ccm2l',	'Tm9sf4',	'Cdk5rap1',	'Gss',	'Trpc4ap',	'Phf20',	'Manbal',	'Ctnnbl1',	'Arhgap40',	'Dnttip1',	'Slc2a10',	'Zfp64',	'Ss18l1',	'Tcfl5',	'Zgpat',	'Lime1',	'Plp2',	'Ftsj1',	'Il13ra1',	'Elf4',	'Gm648',	'Haus7',	'Slc6a8',	'Itgb1bp2',	'Phka2',	'1700064H15Rik',	'Kcnmb3',	'Nudt6',	'Eif2a',	'Gpr171',	'P2ry14',	'Dhx36',	'Gatad2b',	'Them4',	'Anxa9',	'Plekho1',	'Zfp697',	'Nr1h5',	'Capza1',	'Tmem167b',	'Dapp1',	'Acadm',	'Tnni3k',	'Tgs1',	'Gm12353',	'Slc7a13',	'Casp8ap2',	'Creb3',	'Frmpd1',	'Tstd2',	'Fam206a',	'Mrpl37',	'Cc2d1b',	'Dmap1',	'Kdm4a',	'Ppt1',	'Gnl2',	'Lsm10',	'Eva1b',	'Trappc3',	'Ago3',	'Yars',	'Dnajc8',	'Tmco4',	'Ubr4',	'Angptl7',	'Kif1b',	'Zbtb48',	'Mib2',	'Gatad1',	'Sri',	'Slc25a40',	'E130116L18Rik',	'Supt7l',	'Fosl2',	'Yes1',	'Cfap99',	'Sh3bp2',	'Bloc1s4',	'Wfs1',	'Dhx15',	'Srd5a3',	'Slc10a6',	'Tmem175',	'4930522L14Rik',	'Sppl3',	'Fbxw8',	'Oas3',	
                         'Ptpn11',	'Trafd1',	'Adam1b',	'Ccdc63',	'2810006K23Rik',	'Ccdc92',	'Sfswap',	'Ywhag',	'Zkscan1',	'Zscan21',	'Rnf216',	'Lrrc4',	'Akr1b3',	'Cnot4',	'Chrm2',	'Klrg2',	'Parp12',	'Braf',	'Chmp3',	'Pradc1',	'Pcyox1',	'Aplf',	'Srgap3',	'Cidec',	'Il17re',	'Rassf4',	'Bms1',	'Cacna1c',	'Slc25a18',	'Aebp2',	'Sbk3',	'Zfp579',	'Zscan22',	'Ercc1',	'Clasrp',	'Eif3k',	'Spint2',	'Grin2d',	'Hps5',	'Ntrk3',	'Zscan2',	'Ankrd42',	'Slco2b1',	'Spcs2',	'Lipt2',	'Arap1',	'Rhog',	'Rnf141',	'Cog7',	'Slc5a11',	'4930451I11Rik',	'Zfp747',	'Fbrs',	'Slc5a2',	'Stk32c',	'Rnh1',	'Muc6',	'Tpcn2',	'Vip',	'Stxbp5',	'Slc2a12',	'Moxd1',	'Themis',	'Gja1',	'Pla2g12b',	'Micu1',	'Ado',	'Derl3',	'Ybey',	'Ube2g2',	'2610008E11Rik',	'Celf5',	'Ttc41',	'Apaf1',	'Arhgef18',	'Pex11g',	'Fam155a',	'Arhgef10',	'Gm19410',	'Triml2',	'Aga',	'Spcs3',	'Wdr17',	'Tmem192',	'Slc18a1',	'Mau2',	'Haus8',	'Zfp961',	'Gm10282',	'Usp38',	'Rfx1',	'Cacna1a',	'Nkd1',	'Nod2',	'Ccl22',	'Usb1',	'Elmo3',	'Atxn1l',	'Tat',	'Mtss1l',	'Cmip',	'Mbtps1',	'Zfpm1',	'Vps9d1',	'Spire2',	'2310022B05Rik',	'Abhd6',	'Ndst2',	'Dennd6a',	'Fam208a',	'Olfr750',	'Mphosph8',	'Zdhhc20',	'Kcnrg',	'Blk',	'Gm21750',	'Esd',	'Dis3',	'Scel',	'Rbm26',	'Cep57',	'Panx1',	'Tmed1',	'Hspb2',	'Acsbg1',	'Mpi',	'Hcn4',	'Itga11',	'Ppib',	'Rnf111',	'Tex9',	'Mapk6',	'Lysmd2',	'Fbxo9',	'Gk5',	'Slc25a36',	'Gm1123',	'Pik3cb',	'9630041A04Rik',	'Acad11',	'Uba7',	'Bsn',	'Nckipsd',	'Setd2',	'Pdcd6ip',	'Xirp1',	'Etaa1',	'B3gnt2',	'Pus10',	'Rel',	'Cfap36',	'Rhbdf1',	'Nprl3',	'Cdkn2aipnl',	'44082',	'Sap30l',	'Sh3bp5l',	'Pld6',	'Prpsap2',	'Fbxw10',	'Rnf222',	'Asgr1',	'Gm43951',	'Inpp5k',	'Mrm3',	'Dhrs13',	'Atad5',	'Tada2a',	'Tob1',	'Gm11541',	'Osbpl7',	'Kpnb1',	'Krt31',	'Stat5b',	'Mapt',	'Ddx5',	'Cep95',	'Tmc8',	'Pgs1',	'Cep295nl',	'Lgals3bp',	'Aldh5a1',	'Dtnbp1',	'Stmnd1',	'A830005F24Rik',	'Bicd2',	'Ogn',	'Nxnl2',	'Nsd1',	'Prelid1',	'Habp4',	'Slc6a19',	'Ahrr',	'Fam172a',	'Ckmt2',	'Iqgap2',	'Mrps27',	'Fkbp1b',	'Matn3',	'Strn3',	'Sav1',	'Mpp5',	'Plekhd1',	'Ptgr2',	'Zfp410',	'Eif2b2',	'Ift43',	'Gpatch2l',	'Dglucy',	'Papola',	'Ccnk',	'Hhipl1',	'Rtl1',	'Amn',	'Cpq',	'Stk3',	'Snx31',	'Fzd6',	'Ebag9',	'Trappc9',	'Fam83h',	'Dgat1',	'Cpsf1',	'Apol9b',	'Tmprss6',	'Baiap2l2',	'Cbx6',	'Brd1',	'Cpne8',	'Prickle1',	'Fkbp11',	'Krt79',	'Amhr2',	'Calcoco1',	'Pam16',	'Anks3',	'Ppl',	'Serpind1',	'Aifm3',	'Slc7a4',	'Klhl24',	'Map6d1',	'Popdc2',	'Boc',	'Tmem30c',	'Zfp654',	'Usp16',	'Il10rb',
                         'Hlcs',	'Prdm15',	'Tcte2',	'Fam120b',	'Flywch1',	'E4f1',	'Mlst8',	'Slc9a3r2',	'Gm8225',	'Ergic1',	'Nudt3',	'Fgd2',	'Zfp763',	'Rnf5',	'Tnxb',	'2310061I04Rik',	'Ppp1r11',	'H2-M3',	'Ptchd4',	'Runx2',	'Nfkbie',	'Capn11',	'Rftn1',	'Ndufa11',	'Vapa',	'Ddx11',	'Myl12b',	'Abcg5',	'Stpg4',	'Riok3',	'Dsg2',	'Map3k2',	'Kdm3b',	'Etf1',	'Pfdn1',	'Diaph1',	'Dcp2',	'Ticam2',	'Arl14epl',	'Gm9949',	'Impa2',	'Elac1',	'Zadh2',	'Kmt5b',	'Tsga10ip',	'Rps6ka4',	'Lgals12',	'Cyb561a3',	'Zp1',	'Osbp',	'Zfand5',	'Abhd17b',	'Kcnv2',	'Ifit1bl1',	'Zfyve27',	'Golga7b',	'Hif1an',	'Kcnip2',	'Slk',	'Cfap43',	'Gsto2',	'Alkal1',	'Ogfrl1',	'Sdhaf4',	'Il18rap',	'Ndufb3',	'Cflar',	'Abca12',	'Arpc2',	'Resp18',	'Slc19a3',	'Cops7b',	'Mlph',	'Prlh',	'Erfe',	'Tsn',	'Inhbb',	'Dbi',	'Marco',	'Mgat5',	'Il10',	'Shisa4',	'Kif21b',	'1700025G04Rik',	'Tnfsf4',	'Myoc',	'Nme7',	'Atp1b1',	'Gpr161',	'Uap1',	'Apoa2',	'Fcer1g',	'Vsig8',	'Fh1',	'Kmo',	'Sdccag8',	'Sccpdh',	'Degs1',	'Rab3gap2',	'Hsd11b1',	'Stam',	'Hnmt',	'Il1f9',	'Rnf224',	'Fam69b',	'Stkld1',	'Tsc1',	'Cercam',	'Usp20',	'Ptges2',	'Rc3h2',	'Cobll1',	'Pdk1',	'Scrn3',	'Ttc30b',	'Spi1',	'Creb3l1',	'Pla2g4f',	'Galk2',	'Usp8',	'Gzf1',	'Entpd6',	'Map1lc3a',	'Actr5',	'Oser1',	'Gdap1l1',	'Ocstamp',	'Atp9a',	'Cass4',	'Rbm38',	'Pmepa1',	'Col20a1',	'Mtm1',	'Pnck',	'Las1l',	'Igbp1',	'Sat1',	'Ofd1',	'Trappc2',	'Jade1',	'Nhlrc3',	'Bcan',	'Pygo2',	'Ubap2l',	'S100a10',	'Pi4kb',	'Tmod4',	'Sv2a',	'Sec22b',	'Mab21l3',	'Ovgp1',	'Chil4',	'Amigo1',	'Ppa2',	'Nexn',	'Ttpa',	'Mob3b',	'Aptx',	'AI464131',	'Olfr70',	'Tdrd7',	'Tmeff1',	'Ctnnal1',	'Fkbp15',	'Hdhd3',	'Eqtn',	'Mysm1',	'Podn',	'Mknk1',	'Eif2b3',	'Ccdc24',	'Med8',	'Smap2',	'Zmpste24',	'A3galt2',	'Trim62',	'Ythdf2',	'Xkr8',	'Stx12',	'Necap2',	'Vps13d',	'Tas1r1',	'Hes2',	'Gpr153',	'Megf6',	'Nadk',	'Perm1',	'Gm8773',	'Steap2',	'Crygn',	'Adgrf3',	'Pisd',	'Rgs12',	'Hgfac',	'Hs3st1',	'Klb',	'Ccng2',	'Mrpl1',	'Antxr2',	'Bmp3',	'Tmem150c',	'Coq2',	'Lrrc8d',	'Rpl5',	'Pole',	'Grk3',	'Tchp',	'Git2',	'Rad9b',	'Limk1',	'Bcl7b',	'Upk3b',	'Gigyf1',	'Gnb2',	'Agfg2',	'Dnaaf5',	'Micall2',	'Trrap',	'Snd1',	'Flnc',	'Ndufb2',	'Agk',	'Gstk1',	'Zfp746',	'Cd207',	'Tprkb',	'D6Ertd527e',	'Arhgap25',	'Mtmr14',	'Vhl',	'Tmem121b',	'Atp6v1e1',	'Pzp',	'Borcs5',	'H2afj',	'Rerg',	'Pyroxd1',	'Abcc9',	'Stk38l',	'Far2',	'Lair1',	'Isoc2b',	'C5ar2',	'Gpr4',	'Gemin7',	'Bcl3',	'Ceacam16',	'Xrcc1',	'Zfp575',	'Bckdha',	'9530053A07Rik',	'Rps16',	'Gpi1',	'Akt1s1',
                         'Ntn5',	'Rpl18',	'Cyfip1',	'Mrps11',	'Rlbp1',	'Mesp1',	'Mesp2',	'Rccd1',	'Crtc3',	'Sytl2',	'Relt',	'P2ry2',	'Folr1',	'Ilk',	'Eif3f',	'Ccp110',	'Dcun1d3',	'Ubfd1',	'Il27',	'Ccdc189',	'Stx4a',	'Plpp4',	'Abraxas2',	'Dock1',	'Mapk1ip1',	'Cracr2b',	'Chid1',	'Zc3h12d',	'Sf3b5',	'Ccdc162',	'Ostm1',	'Sowahc',	'Tet1',	'Tmem26',	'Ank3',	'Gnaz',	'Dip2a',	'Kiss1r',	'Uqcr11',	'Rexo1',	'Cactin',	'Washc4',	'Utp20',	'Elk3',	'Gm33543',	'Rassf3',	'BC048403',	'Zbtb39',	'A430078G23Rik',	'Camsap3',	'Prr36',	'Tnfsf13b',	'F7',	'Cul4a',	'Lamp1',	'Fbxo25',	'Tpte',	'Bag4',	'Pomk',	'Slc7a2',	'Cyp4v3',	'Sugp1',	'Crtc1',	'Uba52',	'Myo9b',	'Nr2f6',	'Tpm4',	'Prkaca',	'Gpt2',	'Abcc12',	'Ces1h',	'Slc38a7',	'Hsf4',	'Plekhg4',	'Tsnaxip1',	'Txnl4b',	'Cmtr2',	'Il34',	'Hsd17b2',	'Fbxo31',	'Jph3',	'Tcf25',	'Afg3l1',	'Trim67',	'2810004N23Rik',	'Sprtn',	'Rpp14',	'Saysd1',	'Kcnma1',	'Itih1',	'Tnnc1',	'Ddhd1',	'Gm29776',	'Fitm1',	'Rabggta',	'Ltb4r2',	'Gm4491',	'Cryl1',	'Phf11d',	'Nudt18',	'Rcbtb2',	'Lrrc63',	'Lacc1',	'Tnfsf11',	'Pcca',	'1700012B09Rik',	'Zfp317',	'Zfp809',	'Acrv1',	'Tbrg1',	'Usp2',	'Cxcr5',	'Tmprss4',	'BC049352',	'1110032A03Rik',	'Tnfaip8l3',	'Spg21',	'Pigb',	'Unc13c',	'Me1',	'Mthfs',	'Pxylp1',	'Msl2',	'Anapc13',	'Twf2',	'Rad54l2',	'Rbm15b',	'Rassf1',	'Ifrd2',	'1700102P08Rik',	'Prss46',	'Golga4',	'Hhatl',	'Tmem42',	'Exosc7',	'Clec3b',	'Cdcp1',	'Drg1',	'Dusp18',	'Slc35e4',	'Xbp1',	'Camk2b',	'Spred2',	'Vps54',	'Pank3',	'Ttc1',	'Adam19',	'Nipal4',	'Mgat1',	'Tbc1d9b',	'Rmnd5b',	'Gm17334',	'Flcn',	'Top3a',	'Shmt1',	'Tnfsf13',	'Chrnb1',	'Mis12',	'P2rx5',	'Cluh',	'Tsr1',	'Proca1',	'Unc119',	'Slc13a2',	'Tnfaip1',	'Rhbdl3',	'Cdk5r1',	'Rffl',	'1700020L24Rik',	'Wfdc18',	'Synrg',	'Acaca',	'Aatf',	'1700125H20Rik',	'Mtmr4',	'Abcc3',	'Socs7',	'Nt5c3b',	'Ttc25',	'Cnp',	'Coa3',	'Tmem101',	'Asb16',	'Hexim2',	'Gh',	'Grb2',	'Fbf1',	'Tbc1d16',	'Hmga1b',	'Heatr1',	'Nid1',	'Txndc5',	'Faf2',	'Ddx41',	'Smad5',	'Golm1',	'Ice1',	'Rps23',	'Hexb',	'Srek1ip1',	'Depdc1b',	'Gzmk',	'Itga1',	'Klf11',	'Hdac9',	'Ahr',	'Cfl2',	'Slc25a21',	'Nemf',	'Nin',	'Abcd4',	'Ism2',	'Nrde2',	'Cpsf2',	'Fam181a',	'Syne3',	'Ankrd9',	'Selenop',	'Wdr70',	'Pop1',	'Dcstamp',	'Khdrbs3',	'2010109I03Rik',	'Cyc1',	'Gpt',	'Zfp647',	'Slc16a8',	'Csnk1e',	'Tomm22',	'Aco2',	'Prr5',	'Trabd',	'Sbf1',	'Lrrk2',	'Pdzrn4',	'Pced1b',	'Pfkm',	'Lmbr1l',	'Fam186b',	'Pou6f1',	'Krt88',	'Dnase1',	'Rogdi',	'Txndc11',	'Ntan1',	'Dgcr8',	'2510009E07Rik',	
                         'Ahsg',	'Fetub',	'Slc51a',	'Hcls1',	'Nr1i2',	'Tmem39a',	'Btla',	'Retnlb',	'Tmem45a2',	'Crybg3',	'Arl6',	'Cryzl1',	'Dyrk1a',	'Erg',	'Wrb',	'Slc22a1',	'Acat2',	'Prss22',	'Pkd1',	'Bak1',	'Ip6k3',	'Slc26a8',	'Mtch1',	'Pknox1',	'Angptl4',	'H2-Ob',	'Gm20481',	'Mea1',	'Cnpy3',	'Gm7334',	'Kat2b',	'Shd',	'Safb2',	'Togaram2',	'Memo1',	'Morn2',	'Map4k3',	'Crem',	'Trappc8',	'Galnt1',	'Sap130',	'Gpr17',	'Gypc',	'Pkd2l2',	'Gnpda1',	'Prr16',	'Prdm6',	'Dctn4',	'Ptpn2',	'Dym',	'Zfp407',	'Rttn',	'Cabp2',	'Rab1b',	'AI837181',	'Atg2a',	'Fads1',	'Cd5',	'Slc35g1',	'Avpi1',	'Pkd2l1',	'Wnt8b',	'Fgf8',	'Cuedc2',	'Actr1a',	'Shoc2',	'Gucy2g',	'Eif3a',	'Sema4c',	'Vwa3b',	'Abi2',	'43894',	'Cfap221',	'C1ql2',	'Rab7b',	'Plekha6',	'Kdm5b',	'Kcnt2',	'Swt1',	'Blzf1',	'Klhdc9',	'Vangl2',	'Atp1a2',	'Igsf9',	'Rgs7',	'Opn3',	'Efcab2',	'Traf5',	'Kcnh1',	'Irf6',	'Pomt1',	'Tnfaip6',	'Csrnp3',	'Galnt3',	'Erich2',	'4833423E24Rik',	'C1qtnf4',	'Trp53i11',	'D430041D05Rik',	'Grem1',	'Gjd2',	'Dph6',	'Dll4',	'Slc12a1',	'Smox',	'Top1',	'Fitm2',	'Cdh4',	'H2al1a',	'Rgn',	'Dock11',	'Apln',	'Ints6l',	'Prrg3',	'Cnga2',	'Gm7173',	'5430427O19Rik',	'Cxcr3',	'Gprasp2',	'Atg4a',	'Prps2',	'Zfhx4',	'Il7',	'Sucnr1',	'Ccnl1',	'Map9',	'Tmem154',	'Lrba',	'Selenbp2',	'Hist2h2ac',	'Hist2h3b',	'Hist2h2bb',	'Tmigd3',	'Pifo',	'Cept1',	'Cyb561d1',	'Aknad1',	'Fam102b',	'Etnppl',	'Ddit4l',	'Syde2',	'Adgrl4',	'Epha7',	'Rragd',	'Zfp292',	'Ifnk',	'1700009N14Rik',	'Nfx1',	'Dnajb5',	'Acnat1',	'Dnajc25',	'Zfp618',	'Cntln',	'Tek',	'Cyp2j6',	'Cyp2j9',	'Angptl3',	'Alg6',	'Dab1',	'Rnf220',	'Slc6a9',	'Hivep3',	'Gm8439',	'Utp11',	'Zscan20',	'Fgr',	'Vwa5b1',	'Tas1r2',	'Atp13a2',	'Spata21',	'Fhad1',	'Pex10',	'Lhfpl3',	'Cnpy1',	'Lmbr1',	'Letm1',	'Kcnip4',	'Lgi2',	'Sepsecs',	'Arap2',	'Kctd8',	'Corin',	'Cnga1',	'Ociad2',	'Adamts3',	'Bmp2k',	'Gbp6',	'Lrcol1',	'Ulk1',	'Sgsm1',	'1700069L16Rik',	'Acacb',	'Gm10382',	'Slc15a4',	'Ift22',	'Zan',	'Pdk4',	'Met',	'Capza2',	'Opn1sw',	'Strip2',	'Copg2',	'Tmem139',	'Cycs',	'Gars',	'Eif2ak3',	'Alms1',	'C87436',	'Oxtr',	'Ogg1',	'43898',	'Mical3',	'Lrrc23',	'Cracr2a',	'Dera',	'Tmc4',	'Zfp541',	'Ccdc61',	'Irgq',	'Krtdap',	'Zfp507',	'Vstm2b',	'Klk14',	'1700008O03Rik',	'Nell1',	'Pde8a',	'Gm2115',	'Gab2',	'Pak1',	'Omp',	'Tsku',	'Lrrc32',	'Syt17',	'Gga2',	'Oat',	'D7Ertd443e',	'Utf1',	'Ect2l',	'Fyn',	'Mfsd4b1',	'Ddo',	'Hkdc1',	'Jmjd1c',	'Mrln',	'Slc19a1',	'Gm5426',	'Slc6a15',	'Acss3',	'Myf5',	'Kcnc2',	'Yeats4',	'Mdm1',	'D930020B18Rik',
                         'Srgap1',	'Avpr1a',	'Pip4k2c',	'Nxph4',	'Smarcc2',	'Grtp1',	'Myom2',	'Htra4',	'Tacc1',	'Tll1',	'Tmem59l',	'Gipc1',	'Nfix',	'Adgrg3',	'Nudt7',	'Irf8',	'Banp',	'Urb2',	'Disc1',	'Nid2',	'Dupd1',	'Hesx1',	'Syt15',	'Efs',	'Nfatc4',	'Sdr39u1',	'Gja3',	'Eef1akmt1',	'Cab39l',	'Neil2',	'Slc39a14',	'Sftpc',	'Fgf17',	'Gm16485',	'Eepd1',	'Zfp202',	'Cbl',	'Gm32742',	'Arhgap20',	'Csk',	'Arpp19',	'Tinag',	'Bcl2a1d',	'Atr',	'Gnat1',	'Bcl2a1c',	'Crtap',	'Nacad',	'Abca13',	'Ikzf1',	'Ccnjl',	'Dhrs7b',	'Fam83g',	'Ncor1',	'Myh13',	'Cd68',	'Slc16a13',	'Emc6',	'Efcab5',	'Pipox',	'Ksr1',	'Cisd3',	'Acly',	'Plcd3',	'Cacng4',	'Gprc5c',	'Galk1',	'Rnf157',	'Mxra7',	'Gm11733',	'Afmid',	'Fscn2',	'Cd7',	'Calm4',	'Akr1c13',	'Gli3',	'Hist1h2ah',	'Hist1h2ag',	'Tfap2a',	'Rgs14',	'Tgfbi',	'Gm45623',	'Fbp1',	'Rnf180',	'Elovl7',	'Mcidas',	'Nnt',	'Dtnb',	'Trib2',	'Prkar2b',	'Etv1',	'Psma6',	'Mdga2',	'Acot1',	'Bcl11b',	'Dio3',	'Cdh6',	'Cmbl',	'Lrrc6',	'C1qtnf6',	'Mgat3',	'Nell2',	'Tmem106c',	'Acvr1b',	'Ppp1r1a',	'Rmi2',	'Dgkg',	'Muc4',	'Smbd1',	'Cadm2',	'Samsn1',	'Cldn8',	'Mrps6',	'Fndc1',	'Thbs2',	'Hs3st6',	'Cacna1h',	'Hmga1',	'Fance',	'Cpne5',	'Trim40',	'Gm20219',	'Tmem178',	'Abcg8',	'Adcyap1',	'Greb1l',	'Tslp',	'Stard4',	'Gm10549',	'Srp19',	'Fam13b',	'Tmem173',	'E230025N22Rik',	'Gramd3',	'43893',	'St8sia5',	'Znhit2',	'Rfx3',	'Plpp6',	'Tnks2',	'Blnk',	'Nkx2-3',	'Entpd7',	'Scd4',	'Sfxn2',	'Rpl13a-ps1')

VariableFeatures(integrate_epi4, assay = NULL) <- allgenesforanalysis
#A few gene names get mislabeled in transfer between files: 43900, 44083, 44085, 44080, 43891, 43899, 43896, 43892, 43898, 44084, 43895, 44082, 43894, 43898, 43893
ScaleData(integrate_epi4)


AT1_NKX21_binding_acquired_over_time <- list(c('Rb1cc1',	'Arfgef1',	'Prex2',	'Prdm14',	'Tram1',	'Eya1',	'Rdh10',	'Efhc1',	'Tram2',	'Gsta3',	'Smap1',	'Hs6st1',	'Uggt1',	'Arid5a',	'Kansl3',	'Lman2l',	'Mgat4a',	'2010300C02Rik',	'Eif5b',	'Npas2',	'Tbc1d8',	'Il1r1',	'AI597479',	'Cavin2',	'Myo1b',	'Stat4',	'Stat1',	'Stk17b',	'Gtf3c3',	'Rftn2',	'Als2cr12',	'Tmem237',	'Fzd7',	'Nbeal1',	'Nrp2',	'Fzd5',	'Mreg',	'Rpl37a',	'Tns1',	'Gpbar1',	'Stk36',	'Kcne4',	'Ap1s3',	'Fam124b',	'Nyap2',	'Rhbdd1',	'Col4a4',	'Col4a3',	'Agfg1',	'Sp100',	'Kcnj13',	'Hjurp',	'Agap1',	'Ackr3',	'Rbm44',	'Per2',	'Traf3ip1',	'Twist2',	'Hdac4',	'Mterf4',	'Clasp1',	'Tmem185b',	'Tmem37',	'Fcmr',	'Tmcc2',	'Dstyk',	'Mybph',	'Adora1',	'Ipo9',	'Zfp281',	'Ptprc',	'Nek7',	'Ivns1abp',	'Lamc2',	'Axdnd1',	'Tor3a',	'4930523C07Rik',	'Cacybp',	'Rabgap1l',	'Prdx6',	'Dnm3',	'Fmo6',	'Mael',	'Mgst3',	'Atf6',	'F11r',	'Tagln2',	'Grem2',	'1700016C15Rik',	'Smyd3',	'Stum',	'Parp1',	'Capn2',	'Susd4',	'1700056E22Rik',	'Iars2',	'Tgfb2',	'Spata17',	'Smyd2',	'G0s2',	'Prpf18',	'Cdc123',	'Gm13199',	'Celf2',	'Taf3',	'Il2ra',	'Mindy3',	'Cubn',	'Nebl',	'Pip4k2a',	'Arhgap21',	'Gad2',	'Ehmt1',	'Uap1l1',	'Sohlh1',	'Camsap1',	'Setx',	'Med27',	'Sptan1',	'Wdr34',	'Dolpp1',	'1700001O22Rik',	'Aif1l',	'Fam102a',	'Rpl12',	'Mvb12b',	'Cntrl',	'Rab14',	'Gsn',	'Dab2ip',	'Ttll11',	'Gpr21',	'Dennd1a',	'Gtdc1',	'Epc2',	'Ly75',	'Itgb6',	'Fign',	'Fastkd1',	'Gm26558',	'Rapgef4',	'Map3k20',	'Wipf1',	'Osbpl6',	'Prkra',	'Ccdc141',	'Sestd1',	'Zfp385b',	'Ctnnd1',	'Ptprj',	'Nup160',	'Psmc3',	'F2',	'Ambra1',	'Chst1',	'Prdm11',	'Cd82',	'Alkbh3',	'Ttc17',	'Commd9',	'Nat10',	'Bdnf',	'Lin7c',	'Aven',	'Thbs1',	'Srp14',	'Inafm2',	'Bahd1',	'Chac1',	'Ehd4',	'Haus2',	'Sema6d',	'Slc24a5',	'Dut',	'Cops2',	'Ciao1',	'Ttl',	'Slc20a1',	'Stk35',	'Tgm3',	'Cds2',	'Plcb4',	'Kif16b',	'Slc24a3',	'Rin2',	'Fam110a',	'Slc52a3',	'Cox4i2',	'Ccm2l',	'Hck',	'Tm9sf4',	'Cdk5rap1',	'Snta1',	'Chmp4b',	'Gss',	'Trpc4ap',	'Uqcc1',	'Phf20',	'Cnbd2',	'Soga1',	'Samhd1',	'Manbal',	'Ctnnbl1',	'Tgm2',	'Arhgap40',	'Pkig',	'Pigt',	'Dnttip1',	'Snx21',	'Slc2a10',	'Cse1l',	'Slc9a8',	'Ptpn1',	'Pard6b',	'Adnp',	'Zfp64',	'Bmp7',	'Edn3',	'Taf4',	'Ss18l1',	'Tcfl5',	'Ythdf1',	'Zgpat',	'Lime1',	'Myt1',	'Plp2',	'Ftsj1',	'Il13ra1',	'Pgrmc1',	'Elf4',	'Gm648',	'Fhl1',	'Haus7',	'Slc6a8',	'Pls3',	'Tbl1x',	'Efnb1',	'Itgb1bp2',	'Phka1',	'Morf4l2',	'Prps1',	'Tsc22d3',	'Phka2',	'Cdkl5',	'Dnajc5b',	'1700064H15Rik',	'Mecom',	'Lrrc31',	'Kcnmb3',	'Dcun1d1',	'Anxa5',	'Nudt6',	
                                               'Noct',	'Lhfp',	'Commd2',	'Eif2a',	'Gpr171',	'P2ry14',	'Arhgef26',	'Dhx36',	'Rapgef2',	'Tmem144',	'Fam198b',	'Tmem131l',	'Mnd1',	'Trim2',	'Gatb',	'Rusc1',	'Il6ra',	'Rps27',	'Dennd4b',	'Gatad2b',	'S100a14',	'Them4',	'Snx27',	'Cgn',	'Anxa9',	'Setdb1',	'Plekho1',	'Otud7b',	'Notch2',	'Adam30',	'Zfp697',	'Atp1a1',	'Nr1h5',	'Mov10',	'Capza1',	'Kcnd3',	'Sort1',	'Mybphl',	'Tmem167b',	'Stxbp3',	'Slc44a3',	'Bcar3',	'Fnbp1l',	'Aimp1',	'Ppp3ca',	'Dapp1',	'Tspan5',	'Cyr61',	'Bcl10',	'Ttll7',	'Gipc2',	'Acadm',	'Tnni3k',	'Tgs1',	'Rab2a',	'Chd7',	'Plekhf2',	'Gm12353',	'Slc7a13',	'Casp8ap2',	'Pnrc1',	'Akirin2',	'B4galt1',	'Spink4',	'Dcaf12',	'Rusc2',	'Fam166b',	'Creb3',	'Frmpd1',	'Slc25a51',	'Tmod1',	'Tstd2',	'Coro2a',	'Col15a1',	'Ppp3r2',	'Abca1',	'Fam206a',	'Epb41l4b',	'Akap2',	'D630039A03Rik',	'Inip',	'Rgs3',	'Pappa',	'Tle1',	'Lurap1l',	'Ttc39b',	'Dennd4c',	'Cdkn2b',	'Ror1',	'Prkaa2',	'Mrpl37',	'Cc2d1b',	'Osbpl9',	'Ttc39a',	'Cyp4b1',	'Lurap1',	'Plk3',	'Dmap1',	'Artn',	'Kdm4a',	'Szt2',	'Zfp691',	'Edn2',	'Foxo6',	'Ppt1',	'Inpp5b',	'Gnl2',	'Lsm10',	'Eva1b',	'Sh3d21',	'Thrap3',	'Map7d1',	'Trappc3',	'Ago3',	'Rnf19b',	'Yars',	'Pef1',	'Phactr4',	'Dnajc8',	'Sfn',	'Man1c1',	'Clic4',	'Ifnlr1',	'Id3',	'Cdc42',	'Hspg2',	'Alpl',	'Ece1',	'Eif4g3',	'Hp1bp3',	'Otud3',	'Tmco4',	'Nbl1',	'Capzb',	'Ubr4',	'Epha2',	'Spen',	'Kazn',	'Pdpn',	'Dhrs3',	'Angptl7',	'Casz1',	'Kif1b',	'Dnajc11',	'Klhl21',	'Zbtb48',	'Prdm16',	'Ski',	'Faap20',	'Mib2',	'Tnfrsf4',	'Fam133b',	'Gatad1',	'Cyp51',	'Mterf1b',	'Sri',	'Slc25a40',	'Rsbn1l',	'Gsap',	'Kmt2e',	'Rheb',	'E130116L18Rik',	'Rbm33',	'Shh',	'Supt7l',	'Babam2',	'Fosl2',	'Yes1',	'Depdc5',	'Mxd4',	'Cfap99',	'Sh3bp2',	'Msantd1',	'Psapl1',	'D5Ertd579e',	'Bloc1s4',	'Wfs1',	'Dhx15',	'Cckar',	'Chrna9',	'Limch1',	'Slain2',	'Usp46',	'Srd5a3',	'Hopx',	'Igfbp7',	'Shroom3',	'Fras1',	'Anxa3',	'Prkg2',	'Gpat3',	'Cds1',	'Ptpn13',	'Slc10a6',	'Aff1',	'Gak',	'Tmem175',	'4930522L14Rik',	'Ddx51',	'Coro1c',	'Ssh1',	'Kctd10',	'Sppl3',	'Gm10401',	'Taok3',	'Tesc',	'Fbxw8',	'Oas3',	'Ptpn11',	'Trafd1',	'Adam1b',	'Sh2b3',	'Myl2',	'Ccdc63',	'Orai1',	'Rhof',	'Clip1',	'Hcar2',	'Abcb9',	'2810006K23Rik',	'Ccdc92',	'Ncor2',	'Sfswap',	'Tpst1',	'Rfc2',	'Lat2',	'Tmem120a',	'Ywhag',	'Orai2',	'Sh2b2',	'Cux1',	'Azgp1',	'Zkscan1',	'Zscan21',	'Fam20c',	'Gm5294',	'Pdgfa',	'Prkar1b',	'Sun1',	'Get4',	'Gper1',	'Elfn1',	'Mad1l1',	'Chst12',	'Gna12',	'Rnf216',	'Rac1',	'Fam220a',	'Lmtk2',	'Usp12',	'Slc7a1',	'Ubl3',	'Fry',	'N4bp2l1',
                                               'Pds5b',	'Ppp1r9a',	'Mdfic',	'Tes',	'Cav2',	'Cav1',	'Tspan12',	'Cped1',	'Lmod2',	'Wasl',	'Lrrc4',	'Rbm28',	'Plxna4',	'Akr1b3',	'Cnot4',	'Chrm2',	'Klrg2',	'Clec2l',	'Parp12',	'Braf',	'Nfe2l3',	'Snx10',	'Tril',	'Cpvl',	'Wipf3',	'Mindy4',	'Aqp1',	'Chmp3',	'Sftpb',	'Usp39',	'Mat2a',	'Cyp26b1',	'Pradc1',	'Tgfa',	'Pcyox1',	'Anxa4',	'Aak1',	'Antxr1',	'Aplf',	'H1fx',	'Mgll',	'Tpra1',	'Plxna1',	'Chchd6',	'Hdac11',	'Fbln2',	'Tmem43',	'Magi1',	'Foxp1',	'Pdzrn3',	'Itpr1',	'Srgap3',	'Cidec',	'Il17re',	'Rassf4',	'Rasgef1a',	'Csgalnact2',	'Ret',	'Bms1',	'Cacna1c',	'Lrtm2',	'Ninj2',	'Slc25a18',	'Foxj2',	'Cd9',	'Tspan9',	'Gabarapl1',	'Lrp6',	'Crebl2',	'Ddx47',	'Atf7ip',	'Aebp2',	'Recql',	'Ldhb',	'Rassf8',	'Ccdc91',	'Sbk3',	'Zfp579',	'Zscan22',	'Nop53',	'Ehd2',	'Bbc3',	'Sae1',	'Fosb',	'Ercc1',	'Clasrp',	'Bcam',	'Plaur',	'Cyp2b10',	'Capn12',	'Eif3k',	'2200002D01Rik',	'Spint2',	'Dpf1',	'Zfp568',	'Myh14',	'Ap2a1',	'Grin2d',	'Hps5',	'Klf13',	'Tjp1',	'Pcsk6',	'Selenos',	'Igf1r',	'Pgpep1l',	'Fam169b',	'Fam174b',	'Slco3a1',	'Akap13',	'Ntrk3',	'Aen',	'Isg20',	'Mfge8',	'Idh2',	'Zscan2',	'Slc28a1',	'Tm6sf1',	'Saxo2',	'Mex3b',	'Abhd17c',	'Prss23',	'Ankrd42',	'Tenm4',	'Ints4',	'Map6',	'Gdpd5',	'Rps3',	'Arrb1',	'Tpbgl',	'Slco2b1',	'Spcs2',	'Lipt2',	'Ucp3',	'Arap1',	'Rhog',	'Cavin3',	'Rpl27a',	'St5',	'Wee1',	'Swap70',	'Ampd3',	'Rnf141',	'Mrvi1',	'Galnt18',	'Mical2',	'Parva',	'Tead1',	'Rassf10',	'Plekha7',	'Xylt1',	'Gprc5b',	'Scnn1g',	'Scnn1b',	'Cog7',	'Slc5a11',	'D430042O09Rik',	'4930451I11Rik',	'Hirip3',	'Tmem219',	'Zfp747',	'Fbrs',	'Slc5a2',	'Bag3',	'Wdr11',	'Htra1',	'Lhpp',	'Fam53b',	'Dhx32',	'Ptpre',	'Stk32c',	'Inpp5a',	'Rnh1',	'Irf7',	'Muc6',	'Mob2',	'Cttn',	'Ano1',	'Tpcn2',	'Vip',	'Ust',	'Sash1',	'Stxbp5',	'Utrn',	'Stx11',	'Aig1',	'Heca',	'Arfgef3',	'Map3k5',	'Map7',	'1700020N01Rik',	'Sgk1',	'Slc2a12',	'Moxd1',	'Ctgf',	'Themis',	'Marcks',	'Traf3ip2',	'Prdm1',	'Popdc3',	'Mcm9',	'Msl3l2',	'Gja1',	'P4ha1',	'Pla2g12b',	'Oit3',	'Mcu',	'Micu1',	'Spock2',	'Pcbd1',	'Lrrc20',	'H2afy2',	'Col13a1',	'Tspan15',	'Tacr2',	'Gm10118',	'Ado',	'Rtkn2',	'Arid5b',	'A930033H14Rik',	'Ccdc6',	'Susd2',	'Derl3',	'Smarcb1',	'2610028H24Rik',	'Ybey',	'Lss',	'Ube2g2',	'Trappc10',	'2610008E11Rik',	'Cnn2',	'Gpx4',	'Sbno2',	'Ap3d1',	'Celf5',	'Appl2',	'Timp3',	'Ttc41',	'Apaf1',	'Nr2c1',	'Tmcc3',	'Cradd',	'Atp2b1',	'Poc1b',	'Best3',	'Slc35e3',	'Rap1b',	'Baz2a',	'Cs',	'Arhgef18',	'Pex11g',	'Fam155a',	'Irs2',	'Col4a1',	'Col4a2',	'Cars2',	'Atp11a',	
                                               'Arhgef10',	'Kat6a',	'Adam32',	'Plpp5',	'Zfp703',	'Plpbp',	'Mak16',	'Smim18',	'Rbpms',	'Mfhas1',	'Gm19410',	'Mtus1',	'Triml2',	'Fat1',	'Sorbs2',	'Pdlim3',	'Ccdc110',	'Snx25',	'Cfap97',	'Irf2',	'Enpp6',	'Wwc2',	'Aga',	'Spcs3',	'Wdr17',	'Tmem192',	'Slc18a1',	'Gatad2a',	'Mau2',	'Lsm4',	'Haus8',	'Zfp961',	'Gm10282',	'Klf2',	'Large1',	'Arhgap10',	'Smad1',	'Gab1',	'Usp38',	'Gm10645',	'Tbc1d9',	'Rfx1',	'Cacna1a',	'Junb',	'Siah1a',	'Nkd1',	'Nod2',	'Irx6',	'Slc12a3',	'Ccl22',	'Usb1',	'B3gnt9',	'Elmo3',	'Ranbp10',	'Slc12a4',	'Terf2',	'Atxn1l',	'Ap1g1',	'Tat',	'Mtss1l',	'Znrf1',	'Bco1',	'Gan',	'Cmip',	'Cdh13',	'Osgin1',	'Mbtps1',	'Map1lc3b',	'Zfpm1',	'Piezo1',	'Pabpn1l',	'Cdh15',	'Vps9d1',	'Spire2',	'Acta1',	'2310022B05Rik',	'Kcnk1',	'Slc35f3',	'Itgb1',	'Abhd6',	'Pxk',	'Ndst2',	'Vcl',	'Vdac2',	'Zmiz1',	'Anxa11',	'Dennd6a',	'Fam208a',	'Wnt5a',	'Prkcd',	'Ccser2',	'Bmp4',	'Samd4',	'Ktn1',	'Olfr750',	'Mphosph8',	'Zdhhc20',	'Gm6904',	'Kcnrg',	'Blk',	'Fam167a',	'Pinx1',	'Pnoc',	'Dpysl2',	'Dock5',	'9930012K11Rik',	'Sorbs3',	'Gm21750',	'Esd',	'Lrch1',	'Rgcc',	'Dis3',	'Lmo7',	'Scel',	'Rbm26',	'Mbnl2',	'Rap2a',	'Ipo5',	'Farp1',	'Stk24',	'Dync2h1',	'Cep57',	'Panx1',	'Icam1',	'Slc44a2',	'Ilf3',	'Tmed1',	'Kank2',	'Herpud2',	'Ntm',	'Ets1',	'Cdon',	'Nectin1',	'Hspb2',	'Rdx',	'Zc3h12c',	'Kdelc2',	'4930550C14Rik',	'Acsbg1',	'Dnaja4',	'Ube2q2',	'Tspan3',	'Mpi',	'Arid3b',	'Sema7a',	'Hcn4',	'Gramd2',	'Paqr5',	'Anp32a',	'Itga11',	'Pias1',	'Map2k5',	'Aagab',	'Smad3',	'Smad6',	'Hacd3',	'Zfp609',	'Ppib',	'Dapk2',	'Tln2',	'Ice2',	'Rnf111',	'Gm10642',	'Adam10',	'Myzap',	'Tex9',	'Nedd4',	'Mapk6',	'Lysmd2',	'Lrrc1',	'Elovl5',	'Fbxo9',	'Nt5e',	'Plod2',	'Pls1',	'Gk5',	'Slc25a36',	'Gm1123',	'Pik3cb',	'9630041A04Rik',	'Cep63',	'Amotl2',	'Tmem108',	'Acad11',	'Poc1a',	'Hemk1',	'Hyal2',	'Sema3b',	'Sema3f',	'Uba7',	'Bsn',	'Rhoa',	'Prkar2a',	'Nckipsd',	'Cdc25a',	'Setd2',	'Rtp3',	'Lrrfip2',	'Pdcd6ip',	'Cmtm8',	'Tgfbr2',	'Itga9',	'Ctdspl',	'Exog',	'Xirp1',	'Ctnnb1',	'Trak1',	'Cck',	'Snrk',	'Limk2',	'Sec14l3',	'Tbc1d10a',	'Lif',	'Pgam2',	'Polm',	'Tns3',	'Cobl',	'Etaa1',	'B3gnt2',	'Zrsr1',	'Pus10',	'Rel',	'Cfap36',	'Bod1',	'Rhbdf1',	'Nprl3',	'Fgf18',	'Adra1b',	'Rnf145',	'Rnf130',	'Jade2',	'Cdkn2aipnl',	'Skp1a',	'Hspa4',	'44082',	'Slc22a4',	'P4ha2',	'Cdc42se2',	'Tnip1',	'Fat2',	'Sap30l',	'Sh3bp5l',	'Arf1',	'Mprip',	'Pld6',	'Tmem11',	'Prpsap2',	'Adora2b',	'Fbxw10',	'Cdrt4',	'Tekt3',	'Pmp22',	'F930015N05Rik',	'Myocd',	'Map2k4',	'Zkscan6',	
                                               'Ndel1',	'Rnf222',	'Dlg4',	'Asgr1',	'Pimreg',	'Gm43951',	'Cyb5d2',	'Atp2a3',	'Srr',	'Slc43a2',	'Inpp5k',	'Mrm3',	'Timm22',	'Coro6',	'Gm10277',	'Dhrs13',	'Crlf3',	'Atad5',	'Myo1d',	'Dusp14',	'Tada2a',	'Ypel2',	'Mrps23',	'Tob1',	'Wfikkn2',	'Gm11541',	'Pdk2',	'Itga3',	'Dlx3',	'Tac4',	'Slc35b1',	'Phb',	'Osbpl7',	'Kpnb1',	'Mrpl45',	'Stard3',	'Tcap',	'Krt31',	'Krt15',	'Krt42',	'Stat5b',	'Stat3',	'Cavin1',	'Arhgap27',	'Mapt',	'Itgb3',	'Ddx5',	'Cep95',	'Smurf2',	'Helz',	'Cacng5',	'Apoh',	'Slc39a11',	'Unc13d',	'Sphk1',	'Mettl23',	'Mgat5b',	'Tmc8',	'Pgs1',	'Cep295nl',	'Lgals3bp',	'Baiap2',	'Ndufaf8',	'Fasn',	'Gdi2',	'Klf6',	'Zmynd11',	'Chrm3',	'Lyst',	'Slc17a2',	'Carmil1',	'Acot13',	'Aldh5a1',	'Cdkal1',	'Gmds',	'Serpinb6b',	'Serpinb9',	'Pxdc1',	'Rpp40',	'Fars2',	'Gfod1',	'Jarid2',	'Dtnbp1',	'Atxn1',	'Stmnd1',	'Fam8a1',	'Kif13a',	'Rnf144b',	'Zfp169',	'A830005F24Rik',	'Fam120a',	'Card19',	'Fgd3',	'Bicd2',	'Tes3-ps',	'Ogn',	'Nxnl2',	'Gadd45g',	'Diras2',	'Nsd1',	'Prelid1',	'H2afy',	'Fancc',
                                               'Ercc6l2',	'Habp4',	'1700001L19Rik',	'Slc6a19',	'Ahrr',	'Fam172a',	'Ckmt2',	'Rasgrf2',	'Gm20379',	'Ap3b1',	'Pde8b',	'Aggf1',	'F2r',	'Iqgap2',	'F2rl2',	'Arhgef28',	'Utp15',	'Mrps27',	'Ocln',	'Cd180',	'Ndufaf2',	'Arl15',	'Fkbp1b',	'Laptm4a',	'Matn3',	'Ttc32',	'Fam49a',	'2410004P03Rik',	'Hpcal1',	'Cpsf3',	'Cys1',	'Id2',	'Pxdn',	'Sntg2',	'Agr3',	'Stxbp6',	'Prkd1',	'Strn3',	'Arhgap5',	'Nfkbia',	'Mbip',	'Sav1',	'Tmx1',	'Frmd6',	'Ppp2r5e',	'Akap5',	'Max',	'Mpp5',	'Actn1',	'Plekhd1',	'Sipa1l1',	'Ptgr2',	'Zfp410',	'Fcf1',	'Eif2b2',	'Zc2hc1c',	'Tmed10',	'Fos',	'Jdp2',	'Batf',	'Ift43',	'Gpatch2l',	'Cipc',	'Ston2',	'Ttc7b',	'Dglucy',	'Fbln5',	'Rin3',	'Chga',	'Itpk1',	'Papola',	'Ccnk',	'Ccdc85c',	'Hhipl1',	'Degs2',	'Slc25a47',	'Wdr25',	'Rtl1',	'Ppp2r5c',	'Hsp90aa1',	'Mok',	'Amn',	'Mark3',	'Adssl1',	'Cep170b',	'Ahnak2',	'Crip1',	'Wdr60',	'Ccdc152',	'Rictor',	'Rai14',	'1810049J17Rik',	'Zfp622',	'Sema5a',	'Cpq',	'Stk3',	'Snx31',	'Zfp706',	'Ubr5',	'Fzd6',	'Abra',	'Ebag9',	'Trps1',	'Eif3h',	'Mal2',	'Col14a1',	'Zhx2',	'Fam91a1',	'Nsmce2',	'Trib1',	'Fam49b',	'Trappc9',	'Ago2',	'Dennd3',	'Tigd5',	'Fam83h',	'Dgat1',	'Cpsf1',	'Rbfox2',	'Apol9b',	'Myh9',	'Tmprss6',	'Baiap2l2',	'Pla2g6',	'Maff',	'Cbx6',	'Sgsm3',	'Tef',	'Srebf2',	'Poldip3',	'Pacsin2',	'Brd1',	'Creld2',	'Pim3',	'Cpne8',	'Prickle1',	'Arid2',	'Scaf11',	'Endou',	'Ccdc65',	'Fkbp11',	'Aqp2',	'Aqp5',	'Dip2b',	'Atf1',	'6030408B16Rik',	'Krt80',	'Krt7',	'Krt79',	'Sp1',	'Amhr2',	'Calcoco1',	'Adcy9',	'Pam16',	'Cdip1',	'Anks3',	'Ppl',	'Sec14l5',	'Emp2',	'Tekt5',	'Socs1',	'Mkl2',	'Myh11',	'Serpind1',	'Aifm3',	'Slc7a4',	'Comt',	'Txnrd2',	'Hira',	'Klhl24',	'Map6d1',	'Vps8',	'Igf2bp2',	'Tprg',	'P3h2',	'Hes1',	'Xxylt1',	'Muc13',	'Umps',	'Hacd2',	'Adcy5',	'Popdc2',	'Zbtb20',	'Boc',	'Ccdc80',	'Abhd10',	'Cd47',	'Cblb',	'Tmem30c',	'4930453N24Rik',	'Zfp654',	'Usp25',	'Btg3',	'Usp16',	'Eva1c',	'Il10rb',	'Ifngr2',	'Kcne2',	'Rcan1',	'Hlcs',	'Kcnj15',	'Mx1',	'Ripk4',	'Prdm15',	'Arid1b',	'Zdhhc14',	'AC122413.2',	'Prr18',	'Agpat4',	'Tcte2',	'Fam120b',	'Flywch1',	'Kctd5',	'Pdpk1',	'E4f1',	'Mlst8',	'Slc9a3r2',	'Gm38655',	'Mslnl',	'Gm8225',	'Ergic1',	'Atp6v0e',	'Crebrf',	'Lemd2',	'Nudt3',	'Taf11',	'Fgd2',	'Pim1',	'Zfand3',	'Btbd9',	'Zfp763',	'Rab11b',	'Brd2',	'Ager',	'Rnf5',	'Tnxb',	'Cdsn',	'Ddr1',	'2310061I04Rik',	'Rnf39',	'Ppp1r11',	'H2-M3',	'Ptchd4',	'Tnfrsf21',	'Adgrf5',	'Rcan2',	'Runx2',	'Supt3',	'Nfkbie',	'Capn11',	'Tmem63b',	'1600014C23Rik',	'Vegfa',	'Ptk7',	'Taf8',	'Daam2',	'Rftn1',	
                                               'Ndufa11',	'Rfx2',	'Fbxl17',	'Pja2',	'Vapa',	'Ralbp1',	'Ddx11',	'Rab12',	'Arhgap28',	'Tgif1',	'Myl12b',	'Myl12a',	'Lbh',	'Vit',	'Strn',	'Cdkl4',	'Abcg5',	'Prkce',	'Gm10309',	'Stpg4',	'Epcam',	'Foxn2',	'Svil',	'Cables1',	'Riok3',	'Npc1',	'Kctd1',	'Dsg2',	'Dtna',	'Rprd1a',	'Polr2d',	'Lims2',	'Map3k2',	'Nrep',	'Kdm3b',	'Reep2',	'Etf1',	'Cxxc5',	'Pfdn1',	'Diaph1',	'Fgf1',	'Sh3rf2',	'Dcp2',	'Mcc',	'Ccdc112',	'Ticam2',	'Arl14epl',	'Lox',	'Csnk1g3',	'Zfp608',	'Fbn2',	'Smim3',	'Ndst1',	'Rps14',	'Arhgef37',	'Afap1l1',	'Adrb2',	'Gm9949',	'Nedd4l',	'Zfp532',	'Lman1',	'Impa2',	'Elac1',	'Mro',	'Myo5b',	'Acaa2',	'Smad7',	'Zbtb7c',	'Setbp1',	'Adnp2',	'Pqlc1',	'Mbp',	'Tshz1',	'Zadh2',	'Ppp6r3',	'Kmt5b',	'Aldh3b1',	'Lrfn4',	'Tsga10ip',	'Scyl1',	'Frmd8',	'Rps6ka4',	'Cox8a',	'Lgals12',	'Ahnak',	'Scgb1a1',	'Incenp',	'Fth1',	'Cyb561a3',	'Zp1',	'Osbp',	'Psat1',	'Carnmt1',	'Anxa1',	'Zfand5',	'Abhd17b',	'Klf9',	'Gm9493',	'Fam189a2',	'Pip5k1b',	'Dock8',	'Kank1',	'Kcnv2',	'Jak2',	'Ric1',	'Dkk1',	'Prkg1',	'Acta2',	'Ifit2',	'Ifit3',	'Ifit1bl1',	'Ifit1',	'Kif20b',	'Rpp30',	'Ide',	'Plce1',	'Noc3l',	'Pdlim1',	'Sorbs1',	'Mms19',	'Zfyve27',	'Golga7b',	'Scd2',	'Hif1an',	'Kcnip2',	'Trim8',	'Wbp1l',	'Cnnm2',	'Ina',	'Slk',	'Cfap43',	'Gsto1',	'Gsto2',	'Afap1l2',	'Ablim1',	'Trub1',	'Cacul1'))

integrate_epi4 <- AddModuleScore(object = integrate_epi4, features = AT1_NKX21_binding_acquired_over_time, ctrl = 500, name = "NKXAT1AcquiredGenes", search = TRUE, random.seed = 1)
#1 gene not accounted for: 44082
FeaturePlot(object = integrate_epi4, features = 'NKXAT1AcquiredGenes1', cols = c('lightgrey','red'))


AT2_NKX21_binding_retained_from_progenitor <- list(c('Vcpip1',	'Arfgef1',	'Prdm14',	'Ncoa2',	'Lactb2',	'Tram2',	'Phf3',	'Hs6st1',	'Rpl31',	'Nck2',	'1500015O10Rik',	'Wdr75',	'Slc39a10',	'Tmeff2',	'Cavin2',	'Myo1b',	'Stat4',	'Gls',	'Nab1',	'Dnah7a',	'Hecw2',	'Ankrd44',	'Plcl1',	'Sgo2a',	'Cdk15',	'Nrp2',	'Klf7',	'Plekhm3',	'Map2',	'Kansl1l',	'Erbb4',	'Ikzf2',	'Rpl37a',	'Igfbp2',	'Tns1',	'Tmbim1',	'Epha4',	'Scg2',	'Col4a4',	'Sp100',	'Spp2',	'Glrp1',	'Arl4c',	'Sh3bp4',	'Agap1',	'Gbx2',	'Ackr3',	'Lrrfip1',	'Klhl30',	'Per2',	'Hdac4',	'Pdcd1',	'Pam',	'Phlpp1',	'Tmem185b',	'Epb41l5',	'Sctr',	'Nckap5',	'Mapkapk2',	'Slc45a3',	'Lemd1',	'Klhdc8a',	'Cntn2',	'Lax1',	'Prelp',	'Syt2',	'Lmod1',	'Csrp1',	'Ptprc',	'Trove2',	'Rgs2',	'Rgs13',	'Pla2g4a',	'Hmcn1',	'Ncf2',	'Nmnat2',	'Lamc2',	'Xpr1',	'4930523C07Rik',	'Cacybp',	'Rabgap1l',	'Tnfsf18',	'4930558K02Rik',	'Pigc',	'Tada1',	'Uck2',	'Atf6',	'Fcgr3',	'Tstd1',	'Atp1a4',	'Fmn2',	'Pld5',	'Akt3',	'Zbtb18',	'1700016C15Rik',	'Smyd3',	'Lbr',	'Srp9',	'Capn2',	'Capn8',	'1700056E22Rik',	'Iars2',	'Lyplal1',	'Tgfb2',	'Esrrg',	'Kctd3',	'Kcnk2',	'Smyd2',	'Rps6kc1',	'Plxna2',	'Prpf18',	'Bend7',	'Phyh',	'Usp6nl',	'Celf2',	'Taf3',	'Cubn',	'Trdmt1',	'Hacd1',	'H2al2a',	'A930004D18Rik',	'Otud1',	'Arhgap21',	'Abi1',	'Spopl',	'Lcn4',	'Brd3',	'1700001O22Rik',	'Ncs1',	'Qrfp',	'Swi5',	'Fam102a',	'Stxbp1',	'Slc2a8',	'Garnl3',	'Mvb12b',	'Pbx3',	'Phf19',	'Dab2ip',	'Ttll11',	'Crb2',	'Gtdc1',	'Zeb2',	'Lypd6',	'Arl5a',	'Prpf40a',	'Acvr1',	'Wdsub1',	'Rbms1',	'Tank',	'Dpp4',	'Gcg',	'Grb14',	'Map3k20',	'Gpr155',	'Mtx2',	'Osbpl6',	'Ttn',	'Zfp804a',	'Ctnnd1',	'Zdhhc5',	'Ptprj',	'Arfgap2',	'1110051M20Rik',	'Pex16',	'Syt13',	'Ext2',	'Traf6',	'Prr5l',	'Commd9',	'Abtb2',	'BC052040',	'Thbs1',	'Chac1',	'Ehd4',	'Sema6d',	'Stard7',	'Bcl2l11',	'Gm14025',	'Rnf24',	'Lrrn4',	'Plcb4',	'Btbd3',	'Ism1',	'Flrt3',	'Slc24a3',	'Insm1',	'Foxa2',	'Cd93',	'Snph',	'Id1',	'Bcl2l1',	'Asxl1',	'Eif2s2',	'Myl9',	'Mroh8',	'Src',	'Tgm2',	'D630003M21Rik',	'Ppp1r16b',	'Chd6',	'Ptprt',	'Wfdc13',	'Wfdc3',	'Eya2',	'Zmynd8',	'Ptgis',	'Snai1',	'Ripor3',	'Adnp',	'Bcas1',	'Fam210b',	'Fam209',	'Edn3',	'Taf4',	'Lsm14b',	'Lama5',	'Rbbp8nl',	'Tpd52l2',	'Bmp15',	'Dgkk',	'Usp27x',	'Suv39h1',	'Bcor',	'Maob',	'Xiap',	'Stag2',	'Gpc4',	'Gpc3',	'Hprt',	'Plac1',	'Plxnb3',	'Msn',	'Zmym3',	'Hdac8',	'Abcb7',	'Fgf16',	'Pcdh19',	'Tceal9',	'Col4a5',	'Map3k15',	'Pir',	'Tmsb4x',	'Gm21887',	'Lrrcc1',	'Car2',	'Gm9733',	'Pde7a',	'Dnajc5b',	'Ect2',	'Fndc3b',	'Tmem212',	'Mecom',	'Prkci',	'Sox2',	'Atp11b',	'1810062G17Rik',
                                                     'Intu',	'Mgst2',	'Stoml3',	'Smad9',	'Tm4sf4',	'P2ry13',	'Mbnl1',	'Arhgef26',	'Lekr1',	'Schip1',	'Sptssb',	'Golim4',	'Rapgef2',	'Fam198b',	'Ctso',	'Sfrp2',	'Mnd1',	'Glt28d2',	'Kirrel',	'Etv3',	'Arhgef2',	'Gon4l',	'Efna1',	'Kcnn3',	'Tchhl1',	'Snx27',	'Mcl1',	'Car14',	'Otud7b',	'Notch2',	'Hsd3b2',	'Man1a2',	'Igsf3',	'Atp1a1',	'Slc22a15',	'Magi3',	'Mov10',	'Cttnbp2nl',	'Kcnd3',	'Cym',	'Sort1',	'Mybphl',	'Cdc14a',	'Cnn3',	'Bcar3',	'Fnbp1l',	'Fabp2',	'Sec24d',	'Camk2d',	'Enpep',	'Npnt',	'Bdh2',	'Ppp3ca',	'Gm5105',	'Pdlim5',	'Gtf2b',	'Hs2st1',	'Znhit6',	'Cyr61',	'Ddah1',	'Bcl10',	'Adgrl2',	'Erich3',	'Fam110b',	'Sdcbp',	'Nsmaf',	'Chd7',	'Triqk',	'Necab1',	'Ccnc',	'Bach2',	'Aqp3',	'Fam166b',	'Fbxo10',	'Shb',	'Igfbpl1',	'Col15a1',	'Slc44a1',	'Zfp462',	'Epb41l4b',	'Pakap',	'Akap2',	'D630039A03Rik',	'Slc31a1',	'Tnc',	'Pappa',	'Lurap1l',	'Nfib',	'Zdhhc21',	'Cer1',	'Adamtsl1',	'Mllt3',	'Cdkn2b',	'Gm10306',	'Jun',	'Hook1',	'E130114P18Rik',	'Patj',	'Kank4',	'Cachd1',	'Tctex1d1',	'Oma1',	'Acot11',	'Ssbp3',	'Gpx7',	'Dmrta2',	'Trabd2b',	'Olfr1342',	'Guca2a',	'Scmh1',	'Ctps',	'Nfyc',	'Mycl',	'Trit1',	'Macf1',	'AU040320',	'Rnf19b',	'Zbtb8b',	'Phactr4',	'Ahdc1',	'Wasf2',	'Nipal3',	'Id3',	'6030445D17Rik',	'Hspg2',	'Ece1',	'Cda',	'Nbl1',	'Pqlc2',	'Iffo2',	'Epha2',	'Kazn',	'Pdpn',	'Tardbp',	'Rbp7',	'Gpr157',	'Camta1',	'Kcnab2',	'Prdm16',	'B230104I21Rik',	'Morn1',	'Mmp23',	'Mterf1b',	'Cdk14',	'Sema3e',	'Sema3c',	'Kmt2e',	'Wdr86',	'Rnf32',	'Tmem214',	'Tcf23',	'Mrpl33',	'Babam2',	'Depdc5',	'Rnf4',	'Sorcs2',	'Cytl1',	'C1qtnf7',	'Fgfbp1',	'Qdpr',	'Rbpj',	'Cckar',	'Tbc1d19',	'Stim2',	'0610040J01Rik',	'Tlr1',	'Wdr19',	'Apbb2',	'Bend4',	'Gm5108',	'Spata18',	'Usp46',	'Rasl11b',	'Fip1l1',	'Kit',	'Kdr',	'Nmu',	'Igfbp7',	'Grsf1',	'Gc',	'Npffr2',	'Cxcl1',	'Shroom3',	'Sowahb',	'Fras1',	'Paqr3',	'Lin54',	'Arhgap24',	'Aff1',	'Gm17660',	'Lrrc8c',	'Tgfbr3',	'Fgfrl1',	'Zfp932',	'Mn1',	'Coro1c',	'Ssh1',	'Mmab',	'Trpv4',	'Pxn',	'Vsig10',	'Wsb2',	'Fbxo21',	'Rasal1',	'Camkk2',	'Kdm2b',	'Hip1r',	'Zbed5',	'Tmem248',	'Galnt17',	'Gtf2i',	'Cux1',	'Serpine1',	'Fam20c',	'Pdgfa',	'Prkar1b',	'Get4',	'Snx8',	'Chst12',	'Lfng',	'Tnrc18',	'Zfp853',	'Baiap2l1',	'Smurf1',	'Mtus2',	'Ubl3',	'Tex26',	'Wdr95',	'Hsph1',	'Stard13',	'Ppp1r9a',	'Pon3',	'Slc25a13',	'Glcci1',	'Phf14',	'Vwde',	'Tmem168',	'Foxp2',	'Mdfic',	'Tes',	'St7',	'Cped1',	'Asb15',	'Tmem209',	'Plxna4',	'Bpgm',	'Cald1',	'Slc23a4',	'Ptn',	'Dgki',	'Creb3l2',	'Tmem213',	'Hipk2',	'Tbxas1',	'Ezh2',	'Gimap3',	
                                                     'Tril',	'Cpvl',	'Chn2',	'Wipf3',	'Crhr2',	'Aqp1',	'Ccser1',	'Tnip3',	'Ndnf',	'Prdm5',	'Gng12',	'Il23r',	'Atoh8',	'Sh2d6',	'Eva1a',	'Pole4',	'Mthfd2',	'Sfxn5',	'Noto',	'Pcbp1',	'Prokr1',	'Eefsec',	'Plxna1',	'Fbln2',	'Wnt7a',	'Gm15737',	'Magi1',	'Lrig1',	'Fam19a4',	'Pdzrn3',	'Lrrn1',	'Lmcd1',	'Rad18',	'Vgll4',	'Tamm41',	'Rho',	'Csgalnact2',	'Ankrd26',	'Dcp1b',	'Fbxl14',	'Wnk1',	'Tuba8',	'Tuba3a',	'Ccnd2',	'Tspan11',	'Etv6',	'Bcl2l14',	'Lrp6',	'Ddx47',	'Hebp1',	'Gsg1',	'Pbp2',	'Gm8994',	'E330021D16Rik',	'Spx',	'Sox5',	'Rassf8',	'Smco2',	'Ppfibp1',	'Ccdc91',	'Rpl28',	'Usp29',	'Arhgap35',	'Bcam',	'Cyp2b10',	'Zfp626',	'Actn4',	'Sipa1l3',	'Zfp568',	'Fxyd3',	'Zfp658',	'Myh14',	'Sergef',	'Dbx1',	'Nipa2',	'Tjp1',	'Pcsk6',	'Lrrk1',	'Lins1',	'Igf1r',	'Pgpep1l',	'Fam169b',	'Arrdc4',	'Mctp2',	'Fam174b',	'St8sia2',	'Slco3a1',	'Sv2b',	'Anpep',	'Iqgap1',	'Alpk3',	'Tm6sf1',	'Saxo2',	'Mesd',	'Tmem135',	'Prss23',	'Me3',	'Tenm4',	'Nars2',	'Emsy',	'Uvrag',	'Serpinh1',	'Gdpd5',	'Neu3',	'P4ha3',	'Plekhb1',	'Scube2',	'Adm',	'Lyve1',	'Ctr9',	'Galnt18',	'Usp47',	'Mical2',	'Parva',	'Tead1',	'Rassf10',	'Psma1',	'Sox6',	'Plekha7',	'Xylt1',	'Smg1',	'Iqck',	'Cdr2',	'Scnn1b',	'Rbbp6',	'D430042O09Rik',	'Gsg1l',	'Zfp629',	'Fgfr2',	'Tacc2',	'Plekha1',	'Htra1',	'Dmbt1',	'Acadsb',	'Lhpp',	'Fam53b',	'Ctbp2',	'Dhx32',	'Adam12',	'Ptpre',	'Mki67',	'Tcerg1l',	'Dpysl4',	'6430531B16Rik',	'Lsp1',	'Ins2',	'Tspan32',	'Cdkn1c',	'Tnfrsf23',	'Shank2',	'Ano1',	'Ccnd1',	'1810010D01Rik',	'Cnksr3',	'Tab2',	'Ust',	'Utrn',	'Hivep2',	'Reps1',	'Olig3',	'Stx7',	'Ctgf',	'Marcks',	'Traf3ip2',	'Mical1',	'Sec63',	'Prep',	'Dcbld1',	'Msl3l2',	'Ranbp2',	'Sh3rf3',	'44084',	'Anapc16',	'Psap',	'Pald1',	'Tspan15',	'Mypn',	'Ctnna3',	'Gm10118',	'Lrrtm3',	'Rtkn2',	'Cabcoco1',	'Fam13c',	'Agpat3',	'Gzmm',	'Nuak1',	'Ric8b',	'Syn3',	'Igf1',	'Cfap54',	'Nudt4',	'Btg1',	'Atp2b1',	'Dusp6',	'Rassf9',	'Ccdc59',	'Nap1l1',	'Tbc1d15',	'Tspan8',	'Cnot2',	'Best3',	'Cct2',	'Frs2',	'9530003J23Rik',	'Mdm2',	'Slc35e3',	'Hmga2',	'Msrb3',	'Ppm1h',	'Agap2',	'Ndufa4l2',	'Col4a1',	'Ankrd10',	'Atp11a',	'Angpt2',	'Tcim',	'Hgsnat',	'Zfp703',	'Rbpms',	'Dctn6',	'Dusp4',	'Mfhas1',	'Dlc1',	'Tusc3',	'Pdgfrl',	'Fat1',	'Pdlim3',	'Acsl1',	'Irf2',	'Enpp6',	'Wwc2',	'Clcn3',	'Cbr4',	'Psd3',	'Sh2d4a',	'Nwd1',	'Ednra',	'Rbmxl1',	'Zfp827',	'1700011L22Rik',	'Gab1',	'Gm10645',	'Clgn',	'Palm3',	'4930432K21Rik',	'G430095P16Rik',	'Zfp423',	'Adcy7',	'Tox3',	'Aktip',	'Nup93',	'Slc12a3',	'Cdh1',	'Sntb2',	'Znrf1',	'Mon1b',
                                                     'Wwox',	'Atmin',	'Cdh15',	'Nup133',	'Agt',	'Capn9',	'Egln1',	'Kcnk1',	'Slc35f3',	'Irf2bp2',	'Nrp1',	'Flnb',	'Ptprg',	'Gm49355',	'Top2b',	'Rarb',	'Gng2',	'Usp54',	'Rps24',	'Zmiz1',	'Erc2',	'Wnt5a',	'Rft1',	'Arhgap22',	'Ptpn20',	'Zfp488',	'Opn4',	'Fam213a',	'Bmp4',	'Cgrrf1',	'Ndrg2',	'Olfr221',	'Mettl3',	'Ajuba',	'Tgm1',	'Atp12a',	'Xpo4',	'Msra',	'Kif13b',	'Zfp395',	'Rhobtb2',	'Xpo7',	'Siah3',	'Serp2',	'Ccdc122',	'Akap11',	'Naa16',	'Wbp4',	'Cnmd',	'Diaph3',	'Lmo7',	'Spry2',	'Dct',	'Abcc4',	'Mbnl2',	'Farp1',	'Tmem123',	'Arhgap42',	'Jrkl',	'Maml2',	'Amotl1',	'Smco4',	'Slc44a2',	'Arhgap32',	'Kcnj5',	'Kcnj1',	'Fli1',	'Ets1',	'Tbcel',	'Arhgef12',	'Phldb1',	'Dscaml1',	'Sik3',	'Cadm1',	'Nxpe4',	'Dixdc1',	'Fdxacb1',	'Ppp2r1b',	'Exph5',	'Atm',	'Idh3a',	'Ube2q2',	'Peak1',	'Odf3l1',	'1700017B05Rik',	'Arid3b',	'Neo1',	'Thsd4',	'Rplp1',	'Anp32a',	'Coro2b',	'Pias1',	'Map2k5',	'Smad6',	'Lctl',	'Megf11',	'Slc24a1',	'Snx22',	'Herc1',	'Lactb',	'Tpm1',	'Tln2',	'Rora',	'Ice2',	'Anxa2',	'Gm10642',	'Adam10',	'Cgnl1',	'Fam83b',	'Lrrc1',	'Slc17a5',	'Hmgn3',	'Ibtk',	'Plscr1',	'1190002N15Rik',	'Pls1',	'Tfdp2',	'Spsb4',	'Cldn18',	'Cep63',	'Amotl2',	'Slco2a1',	'Cdv3',	'Tmem108',	'Col6a6',	'Dusp7',	'Iqcf5',	'Dag1',	'Ccdc71',	'Prkar2a',	'Lrrfip2',	'Tgfbr2',	'Eomes',	'Ctdspl',	'Scn10a',	'Ctnnb1',	'Trak1',	'Lyzl4',	'Zfp651',	'Tcaim',	'Tbc1d10a',	'Adcy1',	'Gm11985',	'Tns3',	'Fignl1',	'Ddc',	'Grb10',	'Cobl',	'Egfr',	'Ppp3r1',	'Actr2',	'Bcl11a',	'Eml6',	'Sptbn1',	'Fbxw11',	
                                                     'Rars',	'Wwc1',	'Tenm2',	'Adra1b',	'Clint1',	'Fstl4',	'Sparc',	'Igtp',	'Wnt9a',	'Prss38',	'Rai1',	'Srebf1',	'Map2k3',	'Cenpv',	'Trpv2',	'Cdrt4',	'Pmp22',	'Cox10',	'Myocd',	'Map2k4',	'Gas7',	'Usp43',	'Cfap52',	'Stx8',	'Ccdc42',	'Atp2a3',	'Doc2b',	'1700016K19Rik',	'Coro6',	'Slc46a1',	'Pex12',	'Rasl10b',	'Ccl6',	'Hnf1b',	'Dusp14',	'Bcas3',	'Gm11492',	'Mpo',	'Nog',	'Ankfn1',	'Mmd',	'Tom1l1',	'Chad',	'Pdk2',	'Itga3',	'Dlx3',	'Igf2bp1',	'Hoxb1',	'Skap1',	'Srcin1',	'Lasp1',	'Cdk12',	'Grb7',	'Krt13',	'Tubg2',	'Fzd2',	'Higd1b',	'C1ql1',	'Arhgap27',	'Mrc2',	'43900',	'Pitpnc1',	'Cacng5',	'Rgs9',	'Abca9',	'Kcnj16',	'Sox9',	'Llgl2',	'Myo15b',	'Sphk1',	'Sec14l1',	'Engase',	'Rptor',	'Baiap2',	'Slc38a10',	'Sectm1a',	'Asb13',	'Klf6',	'Wdr37',	'Chrm3',	'Hecw1',	'Aoah',	'Hist1h4k',	'Hist1h2ak',	'Slc17a2',	'Carmil1',	'Ripor2',	'Sox4',	'Cdkal1',	'Foxf2',	'Gmds',	'Mylk4',	'Serpinb1c',	'Tubb2a',	'Cdyl',	'Fars2',	'Ly86',	'Rreb1',	'Dsp',	'Bmp6',	'Nedd9',	'Tmem170b',	'Adtrp',	'Tbc1d7',	'Mcur1',	'Gmpr',	'Atxn1',	'Fam8a1',	'Fgd3',	'Spin1',	'Gadd45g',	'Diras2',	'Rab24',	'Klhl3',	'2010111I01Rik',	'Fancc',	'Hsd17b3',	'1700001L19Rik',	'Irx2',	'Ndufs6',	'Rasgrf2',	'Cmya5',	'Jmy',	'S100z',	'F2rl2',	'Enc1',	'Gm10260',	'Arhgef28',	'Foxd1',	'Gm10320',	'Gtf2h2',	'Ocln',	'Pik3r1',	'Srek1',	'Erbin',	'Adamts6',	'Ipo11',	'Rab3c',	'Gapt',	'Plk2',	'Il6st',	'Plpp1',	'BC067074',	'Arl15',	'Ndufs4',	'Mocs2',	'Mfsd2b',	'Ldah',	'Laptm4a',	'Osr1',	'Fam49a',	'Rock2',	'Hpcal1',	'Kidins220',	'Sox11',	'Pxdn',	'Tpo',	'Snx13',	'Tspan13',	'Ispd',	'Stxbp6',	'Arhgap5',	'Snx6',	'Atp5s',	'Cdkl1',	'Atl1',	'Pygl',	'Tmx1',	'Frmd6',	'Six4',	'Ppp2r5e',	'Akap5',	'Ppp1r36',	'Plekhg3',	'Rab15',	'Rad51b',	'Actn1',	'Dcaf5',	'Susd6',	'Map3k9',	'Numb',	'Riox1',	'Dnal1',	'Ltbp2',	'Ttll5',	'Esrrb',	'Irf2bpl',	'Cipc',	'Ahsa1',	'Ptpn21',	'Calm1',	'Ttc7b',	'Gpr68',	'Fbln5',	'Gm28051',	'Ppp4r4',	'Serpina16',	'Evl',	'Traf3',	'Inf2',	'Dnah11',	'Sp4',	'Ghr',	'Slc1a3',	'Nadk2',	'Lmbrd2',	'Rai14',	'Basp1',	'Myo10',	'Fbxl7',	'Ank',	'Trio',	'Dnah5',	'Dap',	'Ankrd33b',	'Sdc2',	'Ywhaz',	'Zfp706',	'Ncald',	'Ubr5',	'Azin1',	'Abra',	'Angpt1',	'Rspo2',	'Trps1',	'Eif3h',	'Med30',	'Ext1',	'Mrpl13',	'Mtbp',	'Has2',	'Zhx2',	'Atad2',	'Fer1l6',	'Trib1',	'Fam84b',	'Fam49b',	'Asap1',	'Dennd3',	'Gm6569',	'1110038F14Rik',	'Mb',	'Apol6',	'Txn2',	'Ift27',	'Mkl1',	'Serhl',	'Cyb5r3',	'Arfgap3',	'Pacsin2',	'Rtl6',	'Ppara',	'Celsr1',	'Fam19a5',	'Adamts20',	'Tmem117',	'Ano6',	'Arid2',	'Slc38a4',	
                                                     'Slc48a1',	'Adcy6',	'Tuba1b',	'Lima1',	'Fignl2',	'Krt18',	'Rarg',	'Emp2',	'Mkl2',	'Parn',	'Myh11',	'Klhl6',	'Vps8',	'Tprg',	'Trp63',	'P3h2',	'Cldn1',	'Fgf12',	'Mb21d2',	'Atp13a5',	'Atp13a4',	'Opa1',	'Hes1',	'Fam43a',	'Dlg1',	'Snx4',	'Zfp148',	'Ccdc14',	'Adcy5',	'Ildr1',	'Ndufb4',	'Fstl1',	'Lsamp',	'Qtrt2',	'Cd96',	'Nectin3',	'Cd47',	'Alcam',	'Zpld1',	'Filip1l',	'Cxadr',	'App',	'Bach1',	'Eva1c',	'Rcan1',	'Clic6',	'Runx1',	'Pcp4',	'Ripk4',	'Arid1b',	'Snx9',	'Qk',	'Agpat4',	'Igf2r',	'Afdn',	'Lix1',	'Spaca6',	'Gm38655',	'D17Wsu92e',	'Ppard',	'Pim1',	'Zfand3',	'Btbd9',	'Abcg1',	'Cryaa',	'Akap8l',	'Brd2',	'Ddr1',	'Gm8909',	'Trim31',	'2410137M14Rik',	'Tnfrsf21',	'Rcan2',	'Supt3',	'Vegfa',	'Ttbk1',	'1700001C19Rik',	'Ccnd3',	'Bysl',	'Lrfn2',	'Mocs1',	'Daam2',	'Kif6',	'Satb1',	'Rfx2',	'Pja2',	'Themis3',	'Ptprm',	'Arhgap28',	'Epb41l3',	'Lbh',	'Rasgrp3',	'Rmdn2',	'Hnrnpll',	'Cox7a2l',	'Zfp36l2',	'Plekhh2',	'Dync2li1',	'Calm2',	'Gm47283',	'Svil',	'Gata6',	'Ankrd29',	'Lama3',	'Cabyr',	'Impact',	'Cdh2',	'Rnf125',	'Dtna',	'2700062C07Rik',	'AW554918',	'Celf4',	'Nrep',	'Epb41l4a',	'Ctnna1',	'Ecscr',	'Cxxc5',	'Cystm1',	'Pcdhgc5',	'Fgf1',	'Nr3c1',	'Pabpc2',	'Ccdc112',	'Tnfaip8',	'Hsd17b4',	'Zfp474',	'Snx2',	'Snx24',	'Cep120',	'Zfp608',	'Megf10',	'Prrc1',	'Ctxn3',	'Slc12a2',	'Fbn2',	'Smim3',	'Ablim3',	'Apcdd1',	'Nedd4l',	'Alpk2',	'Oacyl',	'Chmp1b',	'Spire1',	'Ldlrad4',	'Fam210a',	'Tcf4',	'Gm9925',	'Myo5b',	'Lipg',	'Smad7',	'Ctif',	'Gm6133',	'Setbp1',	'Hsbp1l1',	'Nfatc1',	'Tshz1',	'Cyb5a',	'Timm21',	'Chka',	'Atl3',	'Pla2g16',	'Ahnak',	'Scgb1a1',	'Fads3',	'Myrf',	'Tle4',	'Gnaq',	'Pcsk5',	'Rorb',	'Anxa1',	'Trpm3',	'1700028P14Rik',	'Ptar1',	'Kank1',	'Dmrt1',	'Dmrt3',	'Smarca2',	'Cd274',	'Mbl2',	'Prkg1',	'Cstf2t',	'Sgms1',	'Pten',	'Ifit2',	'Slc16a12',	'Ppp1r3c',	'Btaf1',	'Ide',	'Fra10ac1',	'Plce1',	'Noc3l',	'Tm9sf3',	'Gm340',	'Slit1',	'Frat2',	'Gbf1',	'Ina',	'Stn1',	'Sorcs3',	'Adra2a',	'Acsl5',	'Vti1a',	'Tcf7l2',	'Adrb1',	'Afap1l2',	'Ablim1',	'Rab11fip2',	'Cacul1'))

integrate_epi4 <- AddModuleScore(object = integrate_epi4, features = AT2_NKX21_binding_retained_from_progenitor, ctrl = 500, name = "NKXAT1RetainedGenes", search = TRUE, random.seed = 1)
#2 genes not accounted for 44084 and 43900
FeaturePlot(object = integrate_epi4, features = 'NKXAT1RetainedGenes1', cols = c('lightgrey','red'))


AT2_NKX21_binding_acquired_over_time <- list(c('Alkal1',	'Arfgef1',	'Ogfrl1',	'Sdhaf4',	'Fam168b',	'Uggt1',	'Tmem131',	'Il1r1',	'Il18rap',	'Mrps9',	'AI597479',	'Nabp1',	'Myo1b',	'Ankrd44',	'Aox3',	'Ndufb3',	'Cflar',	'Tmem237',	'Fzd7',	'Cps1',	'Erbb4',	'Abca12',	'Arpc2',	'Tmbim1',	'Ihh',	'Resp18',	'Slc19a3',	'Gm47969',	'Cops7b',	'Dgkd',	'Arl4c',	'Agap1',	'Cops8',	'Mlph',	'Prlh',	'Rbm44',	'Erfe',	'Sned1',	'Tsn',	'Inhbb',	'Epb41l5',	'Dbi',	'Marco',	'Mgat5',	'Tmem163',	'Cd55',	'Gm29427',	'Il10',	'Mapkapk2',	'Slc26a9',	'Pik3c2b',	'Ppp1r15b',	'Golt1a',	'Lax1',	'Chil1',	'Lmod1',	'Shisa4',	'Kif21b',	'Atp6v1g3',	'1700025G04Rik',	'Nmnat2',	'Lamc1',	'Ier5',	'Tor3a',	'Tnfsf4',	'Tnfsf18',	'Myoc',	'Nme7',	'Atp1b1',	'Gpr161',	'Pbx1',	'Uap1',	'Apoa2',	'Fcer1g',	'F11r',	'Alyref2',	'Vsig8',	'Fh1',	'Kmo',	'Sdccag8',	'1700016C15Rik',	'Sccpdh',	'Lbr',	'Degs1',	'Capn2',	'1700056E22Rik',	'Rab3gap2',	'Kcnk2',	'Nenf',	'Hhat',	'Hsd11b1',	'Prpf18',	'Usp6nl',	'Stam',	'Hnmt',	'Il1f9',	'Rnf224',	'Fam69b',	'Lcn4',	'Stkld1',	'Vav2',	'Wdr5',	'Rxra',	'Col5a1',	'Tsc1',	'Cercam',	'Ptpa',	'Usp20',	'Abl1',	'Prrc2b',	'Lcn2',	'Ptges2',	'St6galnac4',	'Cdk9',	'Zbtb43',	'Hc',	'Ptgs1',	'Rc3h2',	'Crb2',	'Tanc1',	'Wdsub1',	'Itgb6',	'Grb14',	'Cobll1',	'Lrp2',	'Gm26558',	'Dcaf17',	'Pdk1',	'Map3k20',	'Scrn3',	'Nfe2l2',	'Ttc30b',	'Ccdc141',	'Zfp804a',	'Spi1',	'Creb3l1',	'Pex16',	'Cd44',	'Elf5',	'Fsip1',	'Bmf',	'Bub1b',	'Ino80',	'Pla2g4f',	'Wdr76',	'Sord',	'Shf',	'Galk2',	'Hdc',	'Gabpb1',	'Usp8',	'Usp50',	'Stard7',	'Slc20a1',	'Ap5s1',	'Rassf2',	'Jag1',	'Btbd3',	'Rrbp1',	'Snx5',	'Ralgapa2',	'Kiz',	'Cd93',	'Gzf1',	'Apmap',	'Acss1',	'Entpd6',	'Slc52a3',	'H13',	'Cox4i2',	'Map1lc3a',	'Uqcc1',	'Phf20',	'Myl9',	'Ctnnbl1',	'Actr5',	'Fam83d',	'Tox2',	'Oser1',	'Gdap1l1',	'Slpi',	'Elmo2',	'Ocstamp',	'Sulf2',	'Ube2v1',	'Tmem189',	'Atp9a',	'Tshz2',	'Zfp217',	'Cass4',	'Rbm38',	'Pmepa1',	'Ctsz',	'Cdh26',	'Lsm14b',	'Cables2',	'Col20a1',	'Polr3k',	'Gata1',	'Cask',	'Fundc1',	'Slc9a7',	'Mtm1',	'Pnck',	'Tbl1x',	'Las1l',	'Igbp1',	'Sat1',	'Ofd1',	'Trappc2',	'Fabp5',	'Car13',	'Mtfr1',	'Nceh1',	'Tnik',	'Actl6a',	'Spry1',	'Jade1',	'Rab33b',	'Nhlrc3',	'Il12a',	'Ift80',	'Fnip2',	'Dchs2',	'Trim2',	'Fhdc1',	'Fbxw7',	'Fam160a1',	'Bcan',	'Mef2d',	'Sema4a',	'Efna1',	'Pygo2',	'Ubap2l',	'S100a10',	'Selenbp1',	'Rfx5',	'Pi4kb',	'Zfp687',	'Tmod4',	'Ciart',	'Sv2a',	'Bcl9',	'Sec22b',	'Notch2',	'Cd101',	'Igsf3',	'Atp1a1',	'Mab21l3',	'Slc22a15',	'Vangl1',	'Ddx20',	'I830077J02Rik',	'Ovgp1',	'Chia1',	'Chil4',	
                                               'Dennd2d',	'Amigo1',	'Sort1',	'Mybphl',	'Vav3',	'Cdc14a',	'Slc44a3',	'Ppa2',	'H2afz',	'Nexn',	'Sdcbp',	'Tox',	'2610301B20Rik',	'Plekhf2',	'Pdp1',	'Ttpa',	'Mob3b',	'Ddx58',	'Aptx',	'B4galt1',	'Aqp3',	'AI464131',	'Olfr70',	'Glipr2',	'Gne',	'Pax5',	'Fbxo10',	'Shb',	'Tdrd7',	'Ncbp1',	'Stx17',	'Msantd3',	'Tmeff1',	'Ppp3r2',	'Fsd1l',	'Klf4',	'Ctnnal1',	'Ptgr1',	'Fkbp15',	'Bspry',	'Hdhd3',	'Ambp',	'Orm1',	'Orm2',	'Plin2',	'Eqtn',	'Mysm1',	'E130114P18Rik',	'Dio1',	'Podn',	'Mknk1',	'Akr1a1',	'Eif2b3',	'Ccdc24',	'Artn',	'Szt2',	'Med8',	'Elovl1',	'Cldn19',	'Rimkla',	'Edn2',	'Slfnl1',	'Smap2',	'Zmpste24',	'Mfsd2a',	'Inpp5b',	'Yrdc',	'Zc3h12a',	'Thrap3',	'A3galt2',	'Trim62',	'Ak2',	'C77080',	'Gm10300',	'Ythdf2',	'Med18',	'Ptafr',	'Xkr8',	'Stx12',	'Sfn',	'Selenon',	'Man1c1',	'Ephb2',	'Cela3b',	'Alpl',	'Hp1bp3',	'Ubr4',	'Arhgef10l',	'Necap2',	'Plekhm2',	'Efhd2',	'Tmem51',	'Dhrs3',	'Vps13d',	'Tnfrsf1b',	'Nppa',	'Pgd',	'Lzic',	'Gpr157',	'Rere',	'Zbtb48',	'Tas1r1',	'Hes2',	'Acot7',	'Gpr153',	'Megf6',	'Prkcz',	'Nadk',	'Perm1',	'Fzd1',	'Gm8773',	'Steap2',	'Tmem243',	'Dmtf1',	'Gnat3',	'Atg9b',	'Crygn',	'Prkag2',	'Paxip1',	'Il6',	'Adgrf3',	'Khk',	'Mrpl33',	'Fosl2',	'Pisd',	'Mxd4',	'Sh3bp2',	'Rgs12',	'Hgfac',	'Psapl1',	'Tada2b',	'Tbc1d14',	'Hs3st1',	'Slc34a2',	'Smim20',	'Cckar',	'Stim2',	'Klb',	'Rbm47',	'Nipal1',	'Slain2',	'Lrrc66',	'Spata18',	'Lnx1',	'Hopx',	'Grsf1',	'Cxcl1',	'Scarb2',	'Ccng2',	'Mrpl1',	'Antxr2',	'Bmp3',	'4930524J08Rik',	'Tmem150c',	'Coq2',	'Cds1',	'Aff1',	'Gm17660',	'Lrrc8d',	'Rpl5',	'Cplx1',	'Pole',	'Tpst2',	'Grk3',	'Tchp',	'Git2',	'Cit',	'Prkab1',	'Taok3',	'Tesc',	'Med13l',	'Aldh2',	'Sh2b3',	'Rad9b',	'Camkk2',	'Kdm2b',	'Orai1',	'Vps37b',	'Rflna',	'Ncor2',	'Tmem248',	'Auts2',	'Limk1',	'Bcl7b',	'Upk3b',	'Myl10',	'Gigyf1',	'Gnb2',	'Agfg2',	'Dnaaf5',	'Micall2',	'Mad1l1',	'Chst12',	'Papolb',	'Usp42',	'Bri3',	'Trrap',	'Smurf1',	'Zfp655',	'N4bp2l1',	'Pds5b',	'Kl',	'Ppp1r9a',	'Pon1',	'Pon3',	'Slc25a13',	'Ica1',	'Mdfic',	'Snd1',	'Flnc',	'Klf14',	'Plxna4',	'Akr1b3',	'Akr1b8',	'Agbl3',	'Kdm7a',	'Ndufb2',	'Agk',	'Gstk1',	'Zfp746',	'Fam221a',	'Osbpl3',	'Skap2',	'Aqp1',	'Rnf103',	'Reep1',	'Sftpb',	'Ggcx',	'Tcf7l1',	'Suclg1',	'Eva1a',	'Cd207',	'Cyp26b1',	'Smyd5',	'Tprkb',	'Asprv1',	'D6Ertd527e',	'Antxr1',	'Arhgap25',	'Rpn1',	'Txnrd3',	'Klf15',	'Slc41a3',	'Gm15737',	'1700123L14Rik',	'Eif4e3',	'Edem1',	'Mtmr14',	'Cpne9',	'Vhl',	'Tatdn2',	'Atg7',	'Csgalnact2',	'Ret',	'Tmem121b',	'Atp6v1e1',	'Usp18',	'Prmt8',	
                                               'Tspan11',	'Pzp',	'Bcl2l14',	'Borcs5',	'Ddx47',	'Plbd1',	'H2afj',	'Rerg',	'Eps8',	'Pyroxd1',	'Abcc9',	'Etnk1',	'Bhlhe41',	'Stk38l',	'Far2',	'Lair1',	'Isoc2b',	'Zfp787',	'Zfp418',	'C5ar2',	'Sae1',	'Gpr4',	'Vasp',	'Gemin7',	'Relb',	'Tomm40',	'Bcl3',	'Ceacam16',	'Xrcc1',	'Zfp575',	'Cxcl17',	'Bckdha',	'Numbl',	'Map3k10',	'9530053A07Rik',	'Rps16',	'Lrfn3',	'Gpi1',	'Cebpa',	'Vsig10l',	'Napsa',	'Akt1s1',	'Ntn5',	'Rpl18',	'Hps5',	'Dbx1',	'Cyfip1',	'Klf13',	'Fan1',	'Selenos',	'Chd2',	'St8sia2',	'Klhl25',	'Mrps11',	'Rlbp1',	'Mesp1',	'Mesp2',	'Rccd1',	'Prc1',	'Crtc3',	'Iqgap1',	'Fah',	'Ctsc',	'Fzd4',	'Sytl2',	'Thrsp',	'Mogat2',	'Relt',	'P2ry2',	'Clpb',	'Folr1',	'Hpx',	'Ilk',	'Eif3f',	'Rpl27a',	'Rnf141',	'Mrvi1',	'Mical2',	'Arl6ip1',	'Ccp110',	'Gprc5b',	'Dcun1d3',	'Lyrm1',	'Ubfd1',	'Lcmt1',	'Il21r',	'Gsg1l',	'Il27',	'Nupr1',	'Sult1a1',	'Ccdc189',	'Stx4a',	'Cox6a2',	'Slc5a2',	'Plpp4',	'Wdr11',	'Fgfr2',	'Ate1',	'Plekha1',	'Fam53b',	'Abraxas2',	'Edrf1',	'Dhx32',	'Dock1',	'Mapk1ip1',	'Inpp5a',	'Zfp941',	'Rnh1',	'Cracr2b',	'Chid1',	'Cdkn1c',	'Slc22a18',	'Osbpl5',	'Shank2',	'Ppfia1',	'Fadd',	'Ccnd1',	'Zc3h12d',	'Stx11',	'Sf3b5',	'Aig1',	'Il22ra2',	'Map3k5',	'Map7',	'Pde7b',	'Arg1',	'Echdc1',	'Tspyl4',	'Frk',	'Slc16a10',	'Ccdc162',	'Ostm1',	'Gm9803',	'Fam184a',	'Sh3rf3',	'Sowahc',	'Oit3',	'Hk1',	'Tet1',	'Tmem26',	'Ank3',	'Ccdc6',	'Gnaz',	'Bcr',	'Dip2a',	'Pcnt',	'Col6a2',	'Col6a1',	'Trappc10',	'Kiss1r',	'Polr2e',	'Gpx4',	'Dazap1',	'Uqcr11',	'Rexo1',	'Cactin',	'Washc4',	'1500009L16Rik',	'Timp3',	'Gnptab',	'Utp20',	'Uhrf1bp1l',	'Cdk17',	'Elk3',	'Cradd',	'Gm33543',	'Poc1b',	'Myrfl',	'Lyz2',	'Lyz1',	'Cpm',	'Slc35e3',	'Gns',	'Rassf3',	'Tbk1',	'BC048403',	'Slc16a7',	'Zbtb39',	'Rbms2',	'A430078G23Rik',	'Camsap3',	'Prr36',	'Tnfsf13b',	'Myo16',	'Irs2',	'Naxd',	'Tex29',	'Atp11a',	'F7',	'Cul4a',	'Lamp1',	'Dcun1d2',	'Tmem255b',	'Gas6',	'Rasa3',	'Fbxo25',	'Cln8',	'Atp7b',	'Tpte',	'Tcim',	'Bag4',	'Pomk',	'Dusp4',	'Dlc1',	'Slc7a2',	'Mtus1',	'F11',	'Cyp4v3',	'Casp3',	'Slc18a1',	'Sugp1',	'Crtc1',	'Uba52',	'Ell',	'Isyna1',	'Myo9b',	'Nr2f6',	'Tpm4',	'Mcm5',	'Prkaca',	'Ier2',	'Lyl1',	'G430095P16Rik',	'Gpt2',	'Abcc12',	'Siah1a',	'Brd7',	'Nkd1',	'Sall1',	'Chd9',	'Rbl2',	'Ces1h',	'Amfr',	'Slc12a3',	'Herpud1',	'Nlrc5',	'Adgrg1',	'Mmp15',	'Slc38a7',	'B3gnt9',	'Hsf4',	'Plekhg4',	'Kctd19',	'Hsd11b2',	'Tsnaxip1',	'Cyb5b',	'Psmd7',	'Txnl4b',	'Cmtr2',	'Il34',	'Ldhd',	'Hsd17b2',	'Mlycd',	'Osgin1',	'Gse1',	'Fbxo31',	'Jph3',	'Slc7a5',	'Ankrd11',	
                                               'Tcf25',	'Afg3l1',	'Rhou',	'Galnt2',	'2310022B05Rik',	'Trim67',	'2810004N23Rik',	'Gnpat',	'Sprtn',	'Egln1',	'Sipa1l2',	'Irf2bp2',	'Tomm20',	'Pard3',	'Flnb',	'Rpp14',	'Ptprg',	'Saysd1',	'Kcnma1',	'Tkt',	'Rft1',	'Mustn1',	'Itih4',	'Itih1',	'Stab1',	'Tnnc1',	'Fam25c',	'Sh2d4b',	'Sftpa1',	'Psmc6',	'Ddhd1',	'Gmfb',	'Fbxo34',	'Tmem260',	'Klhl33',	'Lrp10',	'Gm29776',	'Myh7',	'Fitm1',	'Tgm1',	'Rabggta',	'Ltb4r2',	'Atp12a',	'Gjb2',	'Gm4491',	'Cryl1',	'Phf11d',	'Ctsb',	'Kif13b',	'Scara5',	'Pbk',	'Egr3',	'Nudt18',	'Rcbtb2',	'Lrrc63',	'Lacc1',	'Tnfsf11',	'Spry2',	'Abcc4',	'Dzip1',	'Gpr18',	'Pcca',	'Fut4',	'1700012B09Rik',	'Zfp317',	'Spc24',	'Angptl8',	'Zfp809',	'Snx19',	'St14',	'Aplp2',	'St3gal4',	'Acrv1',	'Tbrg1',	'Scn3b',	'Sorl1',	'Tbcel',	'Usp2',	'Cxcr5',	'Tmprss4',	'BC049352',	'Fxyd2',	'Cadm1',	'Nnmt',	'Zbtb16',	'Plet1',	'1110032A03Rik',	'Ppp2r1b',	'Fdx1',	'Tnfaip8l3',	'1700017B05Rik',	'Tmem202',	'Pkm',	'Tle3',	'Pias1',	'Snapc5',	'Rab11a',	'Slc24a1',	'Rasl12',	'Spg21',	'Snx1',	'Ice2',	'Pygo1',	'Pigb',	'Unc13c',	'Fam214a',	'Myo5c',	'Eef1a1',	'Me1',	'Mthfs',	'Tbc1d2b',	'Zbtb38',	'Pxylp1',	'Msl2',	'Anapc13',	'Rab6b',	'Twf2',	'Alas1',	'Rad54l2',	'Rbm15b',	'Rassf1',	'Ifrd2',	'Amigo3',	'1700102P08Rik',	'Plxnb1',	'Cspg5',	'Scap',	'Setd2',	'Prss46',	'Tgfbr2',	'Golga4',	'Ctdspl',	'Acaa1a',	'Exog',	'Ctnnb1',	'Trak1',	'Hhatl',	'Higd1a',	'Snrk',	'Ano10',	'Abhd5',	'Tmem42',	'Zdhhc3',	'Exosc7',	'Clec3b',	'Cdcp1',	'Limd1',	'Drg1',	'Morc2a',	'Dusp18',	'Slc35e4',	'Ascc2',	'Nf2',	'Gas2l1',	'Emid1',	'Xbp1',	'Camk2b',	'Igfbp3',	'Spred2',	'Vps54',	'Bod1',	'Cpeb4',	'Ubtd2',	'Fbxw11',	'Pank3',	'Ttc1',	'Adra1b',	'Rnf145',	'Adam19',	'Nipal4',	'Ifi47',	'Mgat1',	'Tbc1d9b',	'Col23a1',	'Rmnd5b',	'Fstl4',	'44082',	'Gm17334',	'Acsl6',	'Gpx3',	'Fat2',	'Trim11',	'Pld6',	'Flcn',	'Rasd1',	'Llgl1',	'Top3a',	'Shmt1',	'Natd1',	'Map2k3',	'B9d1',	'Epn2',	'Akap10',	'Adora2b',	'Tekt3',	'Shisa6',	'Gas7',	'Dhrs7c',	'Myh10',	'Dnah2',	'Tnfsf13',	'Chrnb1',	'Slc16a11',	'Mis12',	'Pimreg',	'Tekt1',	'Spns2',	'P2rx1',	'Haspin',	'P2rx5',	'Cluh',	'Mnt',	'Tsr1',	'Srr',	'Hic1',	'Rtn4rl1',	'Rpa1',	'Inpp5k',	'Doc2b',	'Rflnb',	'Dbil5',	'Coro6',	'Proca1',	'Unc119',	'Slc13a2',	'Poldip2',	'Tnfaip1',	'Lyrm9',	'Rhbdl3',	'Cdk5r1',	'Rffl',	'1700020L24Rik',	'Wfdc18',	'Synrg',	'Acaca',	'Aatf',	'1700125H20Rik',	'Tbx2',	'Rnft1',	'Tubd1',	'Vmp1',	'Mtmr4',	'Hsf5',	'Trim25',	'Nog',	'Tob1',	'Abcc3',	'Abi3',	'Snx11',	'Sp2',	'Osbpl7',	'Tbkbp1',	'Socs7',	'Srcin1',	'Ppp1r1b',	'Stard3',	
                                               'Pgap3',	'Nt5c3b',	'Ttc25',	'Cnp',	'Stat3',	'Coa3',	'Meox1',	'Tmem101',	'Asb16',	'Dcakd',	'Hexim2',	'Map3k14',	'Arhgap27',	'Tanc2',	'Cyb561',	'Gh',	'Tex2',	'Pitpnc1',	'Cacng1',	'Apoh',	'Cdc42ep4',	'Grb2',	'Tmem94',	'Wbp2',	'Fbf1',	'Sphk1',	'St6galnac2',	'Pgs1',	'Tbc1d16',	'Endov',	'Chmp6',	'Faap100',	'Cbr2',	'Hmga1b',	'Ccdc57',	'Ptchd3',	'Larp4b',	'Heatr1',	'Nid1',	'Lyst',	'Rpp40',	'Ly86',	'Rreb1',	'Ssr1',	'Bmp6',	'Txndc5',	'Cd83',	'Stmnd1',	'Fam8a1',	'Rnf144b',	'Phf2',	'Gadd45g',	'Auh',	'Nfil3',	'Faf2',	'Ddx41',	'Smad5',	'Agtpbp1',	'Golm1',	'2010111I01Rik',	'Aaed1',	'Ice1',	'Rps23',	'Ssbp2',	'Lhfpl2',	'Hexb',	'Pik3r1',	'Cd180',	'Nln',	'Srek1ip1',	'Ipo11',	'Depdc1b',	'Rab3c',	'Cdc20b',	'Gzmk',	'Itga1',	'Parp8',	'Gm17509',	'Hmgcs1',	'Klhl29',	'Hs1bp3',	'Sdc1',	'Fam49a',	'Ntsr2',	'Klf11',	'Rnf144a',	'Adi1',	'Tmem18',	'Hdac9',	'Ahr',	'Ifrd1',	'Cfl2',	'Slc25a21',	'Nemf',	'Nin',	'Psma3',	'Slc38a6',	'Gphb5',	'Fut8',	'Plekhh1',	'Zfp36l1',	'Ccdc177',	'Ttc9',	'Pcnx',	'Dpf3',	'Dcaf4',	'Abcd4',	'Npc2',	'Ltbp2',	'Tmed10',	'Fos',	'Cipc',	'Pomt2',	'Ahsa1',	'Ism2',	'Nrxn3',	'Nrde2',	'Cpsf2',	'Chga',	'Itpk1',	'Fam181a',	'Syne3',	'Mok',	'Ankrd9',	'Amn',	'Siva1',	'Selenop',	'Fbxo4',	'Lifr',	'Egflam',	'Wdr70',	'Prlr',	'Npr3',	'Dap',	'Pop1',	'Ubr5',	'Dcstamp',	'Deptor',	'Derl1',	'Ndufb9',	'Mtss1',
                                               'Trib1',	'Fam49b',	'Tg',	'Ndrg1',	'St3gal1',	'Khdrbs3',	'Ly6d',	'2010109I03Rik',	'Ly6e',	'Cyc1',	'Scx',	'Gpt',	'Zfp647',	'Rbfox2',	'Slc16a8',	'Tmem184b',	'Csnk1e',	'Ddx17',	'Tomm22',	'Rpl3',	'Sgsm3',	'Aco2',	'Desi1',	'Mcat',	'Prr5',	'Fbln1',	'Wnt7b',	'Ppara',	'Fam19a5',	'Brd1',	'Pim3',	'Trabd',	'Plxnb2',	'Sbf1',	'Lrrk2',	'Pdzrn4',	'Pced1b',	'Pfkm',	'Lmbr1l',	'Fam186b',	'Slc11a2',	'Gm5475',	'Pou6f1',	'Cela1',	'Nr4a1',	'Krt80',	'Krt88',	'Tns2',	'Amhr2',	'Nfe2',	'Dnase1',	'Mgrn1',	'Nudt16l1',	'Rogdi',	'Gm5767',	'Clec16a',	'Litaf',	'Txndc11',	'Snx29',	'Cpped1',	'Ercc4',	'Mkl2',	'Ntan1',	'Fgd4',	'Slc7a4',	'Dgcr8',	'Comt',	'Lamp3',	'Ap2m1',	'2510009E07Rik',	'Etv5',	'Ahsg',	'Fetub',	'Rfc4',	'Bcl6',	'Lpp',	'Tprg',	'Il1rap',	'Slc51a',	'Tfrc',	'Snx4',	'Kalrn',	'Hacd2',	'Hcls1',	'Gsk3b',	'Nr1i2',	'Tmem39a',	'Qtrt2',	'Btla',	'Retnlb',	'Tmem45a2',	'Cmss1',	'Crybg3',	'Arl6',	'Btg3',	'Map3k7cl',	'Bach1',	'Tmem50b',	'Cryzl1',	'Clic6',	'Chaf1b',	'Dyrk1a',	'Kcnj15',	'Erg',	'Wrb',	'Ezr',	'Slc22a3',	'Slc22a1',	'Mrpl18',	'Acat2',	'Prss22',	'Kctd5',	'Rnps1',	'Pkd1',	'Dusp1',	'Bak1',	'Ip6k3',	'Taf11',	'Slc26a8',	'Stk38',	'BC004004',	'Mtch1',	'Fgd2',	'Pim1',	'Pknox1',	'Cryaa',	'Sik1',	'Angptl4',	'H2-Ob',	'H2-Aa',	'Gm20481',	'Ddx39b',	'1600014C23Rik',	'Vegfa',	'Polr1c',	'Ptk7',	'Mea1',	'Cnpy3',	'Mocs1',	'Gm7334',	'Tbc1d5',	'Kat2b',	'Shd',	'Dpp9',	'Safb2',	'Rfx2',	'C3',	'Ptprm',	'Togaram2',	'Xdh',	'Memo1',	'Rasgrp3',	'Cdc42ep3',	'Rmdn2',	'Morn2',	'Map4k3',	'Kcng3',	'Zfp36l2',	'Thada',	'Dync2li1',	'Gm10309',	'Epas1',	'Mcfd2',	'Ttc7',	'Calm2',	'Crem',	'Svil',	'Zfp438',	'Cables1',	'Npc1',	'Cabyr',	'Trappc8',	'Rnf125',	'Galnt1',	'Sap130',	'Gpr17',	'Gypc',	'Nrep',	'Reep5',	'Pkd2l2',	'Sil1',	'Pcdhgc5',	'Diaph1',	'Gnpda1',	'Pabpc2',	'Rbm27',	'Trim36',	'Ticam2',	'Prr16',	'Snx24',	'Prdm6',	'Dctn4',	'Synpo',	'Cd74',	'Hmgxb3',	'Pde6a',	'Ppargc1b',	'Nedd4l',	'Impa2',	'Ptpn2',	'Ldlrad4',	'Fam210a',	'Mex3c',	'Acaa2',	'Dym',	'Ctif',	'Zfp236',	'Zfp516',	'Tshz1',	'Zadh2',	'Zfp407',	'Rttn',	'Lrp5',	'Gstp3',	'Cabp2',	'Pcx',	'Mrpl11',	'Rab1b',	'AI837181',	'Ap5b1',	'Atg2a',	'Rasgrp2',	'Otub1',	'Fads1',	'Cd5',	'Cd6',	'Gna14',	'Rfk',	'Anxa1',	'Klf9',	'Pip5k1b',	'Fam122a',	'Gm815',	'Vldlr',	'Kcnv2',	'Rcl1',	'Il33',	'Acta2',	'Lipa',	'Kif20b',	'Ppp1r3c',	'Slc35g1',	'Sorbs1',	'Ccnj',	'Ankrd2',	'Avpi1',	'Pyroxd2',	'Cwf19l1',	'Pkd2l1',	'Scd1',	'Wnt8b',	'Fgf8',	'Cuedc2',	'Actr1a',	'Cnnm2',	'Sh3pxd2a',	'Sorcs3',	'Xpnpep1',	'Dusp5',	'Shoc2',	'Gpam',	'Gucy2g',	'Shtn1',	'Eif3a',	'Zfp950'))


integrate_epi4 <- AddModuleScore(object = integrate_epi4, features = AT2_NKX21_binding_acquired_over_time, ctrl = 500, name = "NKXAT2AcquiredGenes", search = TRUE, random.seed = 1)
#1 gene unaccounted for 44082
FeaturePlot(object = integrate_epi4, features = 'NKXAT2AcquiredGenes1', cols = c('lightgrey','red'))


AT2_NKX21_binding_retained_from_progenitor <- list(c('Ncoa2',	'Sema4c',	'Tmem131',	'Vwa3b',	'Inpp4a',	'Pdcl3',	'Slc39a10',	'Nabp1',	'Myo1b',	'Mfsd6',	'1700019D03Rik',	'Dnah7a',	'Satb2',	'Fzd7',	'Gm973',	'Fam117b',	'Abi2',	'Fzd5',	'Kansl1l',	'Cps1',	'Abca12',	'43894',	'Igfbp5',	'Epha4',	'Acsl3',	'Serpine2',	'Kcnj13',	'Cops8',	'Asb1',	'Bok',	'Gli2',	'Cfap221',	'Dbi',	'C1ql2',	'Slc35f5',	'Gpr39',	'Lypd1',	'Mgat5',	'Ccnt2',	'Cxcr4',	'Cd55b',	'Il10',	'Mapkapk2',	'Rab7b',	'Slc26a9',	'Cntn2',	'Plekha6',	'Golt1a',	'Snrpe',	'Prelp',	'Kdm5b',	'Syt2',	'Ube2t',	'Csrp1',	'Zfp281',	'Atp6v1g3',	'Nek7',	'Kcnt2',	'Ivns1abp',	'Swt1',	'Edem3',	'1700025G04Rik',	'Shcbp1l',	'Ier5',	'Soat1',	'Tnfsf18',	'Blzf1',	'Dpt',	'Gpr161',	'Lrrc52',	'Pbx1',	'Cfap126',	'Klhdc9',	'Vangl2',	'Atp1a2',	'Igsf9',	'Tagln2',	'Cfap45',	'Rgs7',	'Opn3',	'1700016C15Rik',	'Efcab2',	'Acbd3',	'Tmem63a',	'Lpgat1',	'Traf5',	'Kcnh1',	'Hhat',	'Irf6',	'A130010J15Rik',	'Proser2',	'Celf2',	'Notch1',	'Vav2',	'Rxra',	'Ier5l',	'Cstad',	'1700001O22Rik',	'Ass1',	'Pomt1',	'Lmx1b',	'Mvb12b',	'Hc',	'Cntrl',	'Lypd6b',	'Rnd3',	'Tnfaip6',	'Fmnl2',	'Rprm',	'Itgb6',	'Tank',	'Grb14',	'Cobll1',	'Csrnp3',	'Galnt3',	'Sp5',	'Erich2',	'Cybrd1',	'Map3k20',	'Cdca7',	'Osbpl6',	'Zfp385b',	'Prdx6b',	'4833423E24Rik',	'Ptprj',	'C1qtnf4',	'Trp53i11',	'Cd44',	'Ehf',	'D430041D05Rik',	'Rcn1',	'Mpped2',	'Grem1',	'Gjd2',	'Dph6',	'Fsip1',	'Eif2ak4',	'Bmf',	'Dll4',	'Sqor',	'Slc12a1',	'Mal',	'Acoxl',	'Bcl2l11',	'Tmem87b',	'Cenpb',	'Smox',	'Bmp2',	'Plcb4',	'Btbd3',	'Rrbp1',	'Ralgapa2',	'Foxa2',	'Fam110a',	'Id1',	'Dnmt3b',	'Bpifb1',	'Bpifb5',	'Epb41l1',	'Bpi',	'Dhx35',	'Top1',	'Ptprt',	'Fitm2',	'Sdc4',	'Eya2',	'Sulf2',	'Cse1l',	'Slc9a8',	'Ptpn1',	'Zfp217',	'Rbm38',	'Pmepa1',	'Cdh4',	'Taf4',	'Shroom4',	'Clcn5',	'Mid1ip1',	'H2al1a',	'Rgn',	'Dock11',	'Apln',	'Mbnl3',	'Gpc4',	'Ints6l',	'Prrg3',	'Cnga2',	'Gm7173',	'5430427O19Rik',	'Amer1',	'Yipf6',	'Pja1',	'Cxcr3',	'Nexmif',	'Gprasp2',	'Bex2',	'Atg4a',	'Smpx',	'Rps6ka3',	'Tceanc',	'Egfl6',	'Prps2',	'Zfhx4',	'Il7',	'Zbtb10',	'Zfp704',	'Fabp5',	'Fabp12',	'Lrrcc1',	'Dnajc5b',	'Fndc3b',	'Tmem212',	'Pld1',	'Skil',	'Ccdc39',	'Atp11b',	'Pgrmc2',	'Noct',	'Mgst2',	'Maml3',	'Frem2',	'Mab21l1',	'Sucnr1',	'Ccnl1',	'Sptssb',	'Map9',	'Dchs2',	'Trim2',	'Fhdc1',	'Tigd4',	'Tmem154',	'Fbxw7',	'Lrba',	'Bcan',	'Arhgef2',	'Efna1',	'Pmvk',	'Gatad2b',	'Selenbp2',	'Hist2h2ac',	'Hist2h3b',	'Hist2h2bb',	'Chd1l',	'Vtcn1',	'Ptgfrn',	'Igsf3',	'Atp1a1',	'Vangl1',	'Slc16a1',	'Ddx20',
                                                     'Tmigd3',	'Pifo',	'Cept1',	'Cyb561d1',	'Aknad1',	'Fam102b',	'Cdc14a',	'Snx7',	'Dpyd',	'Camk2d',	'Enpep',	'Elovl6',	'Etnppl',	'Dkk2',	'Arhgef38',	'Slc39a8',	'Ddit4l',	'Bmpr1b',	'Syde2',	'Adgrl2',	'Adgrl4',	'St6galnac3',	'Nsmaf',	'Tox',	'Car8',	'Chd7',	'Gem',	'Epha7',	'Bach2',	'Rragd',	'Zfp292',	'Mob3b',	'Ifnk',	'1700009N14Rik',	'Nfx1',	'Aqp3',	'Ubap2',	'Dnajb5',	'Tgfbr1',	'Tex10',	'Acnat1',	'Rnf20',	'Rad23b',	'D630039A03Rik',	'Dnajc25',	'Susd1',	'Slc31a2',	'Zfp618',	'Ambp',	'Cntln',	'Adamtsl1',	'Mllt3',	'Tek',	'Jun',	'Cyp2j6',	'Cyp2j9',	'E130114P18Rik',	'Nfia',	'Angptl3',	'Alg6',	'Ror1',	'Cachd1',	'Jak1',	'Oma1',	'Dab1',	'Bend5',	'Pdzk1ip1',	'Kif2c',	'Rnf220',	'Slc6a9',	'Hivep3',	'Gm8439',	'Kcnq4',	'Mfsd2a',	'Pou3f1',	'Utp11',	'Zscan20',	'Yars',	'C77080',	'Marcksl1',	'Pef1',	'Hcrtr1',	'Nkain1',	'Fgr',	'Fuca1',	'Hnrnpr',	'Lactbl1',	'Rap1gap',	'Vwa5b1',	'Tmco4',	'Tas1r2',	'Pax7',	'Arhgef10l',	'Atp13a2',	'Spata21',	'Arhgef19',	'Epha2',	'Spen',	'Fhad1',	'Tmem51',	'Vps13d',	'Cenps',	'Kif1b',	'Hes2',	'Acot7',	'Prdm16',	'Pex10',	'Fzd1',	'Sri',	'Sema3c',	'Cd36',	'Gnai1',	'Lhfpl3',	'Srpk2',	'Insig1',	'Cnpy1',	'Shh',	'Lmbr1',	'Letm1',	'Cpeb2',	'Cc2d2a',	'Kcnip4',	'Ppargc1a',	'Lgi2',	'Sepsecs',	'Pi4k2b',	'Cckar',	'Arap2',	'0610040J01Rik',	'Klf3',	'Klhl5',	'Wdr19',	'9130230L23Rik',	'Rbm47',	'Atp8a1',	'Kctd8',	'Corin',	'Cnga1',	'Ociad2',	'Spata18',	'Rasl11b',	'Fip1l1',	'Lnx1',	'Rufy3',	'Gc',	'Npffr2',	'Adamts3',	'Cox18',	'Cxcl15',	'Areg',	'Btc',	'Parm1',	'Mrpl1',	'Bmp2k',	'Prkg2',	'Tmem150c',	'Coq2',	'Arhgap24',	'Ptpn13',	'Aff1',	'Gbp6',	'Lrrc8c',	'Lrrc8d',	'Lrcol1',	'Ulk1',	'Sez6l',	'Myo18b',	'Sgsm1',	'1700069L16Rik',	'Acacb',	'Wsb2',	'2410131K14Rik',	'Med13l',	'Rhof',	'Arl6ip4',	'Gm10382',	'Tmem132c',	'Slc15a4',	'Tmem132d',	'Galnt17',	'Auts2',	'Cldn4',	'Cldn3',	'Mlxipl',	'Myl10',	'Col26a1',	'Ift22',	'Fis1',	'Ephb4',	'Zan',	'Cyp3a13',	'Mad1l1',	'Smurf1',	'Usp12',	'Gm42791',	'Medag',	'Tex26',	'Fry',	'Pdk4',	'Glcci1',	'Ppp1r3a',	'Met',	'Capza2',	'Wnt2',	'Tspan12',	'Aass',	'Cadps2',	'Snd1',	'Opn1sw',	'Strip2',	'Tmem209',	'Cep41',	'Copg2',	'Klf14',	'Plxna4',	'Chchd3',	'Chrm2',	'Tmem213',	'Tmem178b',	'Tmem139',	'Gsdme',	'Cycs',	'Nfe2l3',	'Snx10',	'Hibadh',	'Znrf2',	'Gars',	'Fam13a',	'Eif2ak3',	'Tgoln1',	'Tcf7l1',	'Hk2',	'Cyp26b1',	'Alms1',	'C87436',	'Pcbp1',	'Antxr1',	'Prokr1',	'Iqsec1',	'Hdac11',	'Gm15737',	'Magi1',	'Mitf',	'Foxp1',	'Rybp',	'Oxtr',	'Ogg1',	'Tatdn2',	'Vgll4',	'Tsen2',	'43898',
                                                     'Dcp1b',	'Tmem121b',	'Cecr2',	'Mical3',	'Lrrc23',	'Cracr2a',	'Bcl2l14',	'Mansc1',	'Emp1',	'Plbd1',	'H2afj',	'Eps8',	'Dera',	'Mgst1',	'Pik3c2g',	'Etnk1',	'Sox5',	'Tmc4',	'Zfp541',	'Zc3h4',	'Hif3a',	'Ccdc61',	'Irf2bp1',	'Vasp',	'Bcl3',	'Irgq',	'Ceacam1',	'Cyp2s1',	'Cyp2f2',	'Hnrnpl',	'Catsperg1',	'Krtdap',	'Zfp507',	'Zfp536',	'Uri1',	'Vstm2b',	'Klk14',	'Klk13',	'1700008O03Rik',	'Abcc6',	'Dbx1',	'Nell1',	'Cers3',	'Arrdc4',	'Slco3a1',	'Zscan2',	'Pde8a',	'Mesd',	'Gm2115',	'Ctsc',	'Tmem135',	'Eed',	'Sytl2',	'Nars2',	'Gab2',	'Pak1',	'Myo7a',	'Omp',	'Tsku',	'Lrrc32',	'Thap12',	'Serpinh1',	'Lmo1',	'Ascl3',	'Wee1',	'Adm',	'Rnf141',	'Mical2',	'Parva',	'Rassf10',	'Arntl',	'Spon1',	'Xylt1',	'Syt17',	'Gga2',	'BC017158',	'Rgs10',	'Wdr11',	'Nsmce4a',	'Oat',	'D7Ertd443e',	'Mgmt',	'Tcerg1l',	'Inpp5a',	'Utf1',	'Eps8l2',	'Kcnq1',	'Cdkn1c',	'Tnfrsf23',	'Ccnd1',	'Sash1',	'Plagl1',	'Cited2',	'Ect2l',	'Olig3',	'Il22ra2',	'Map7',	'Arg1',	'Ptprk',	'Fyn',	'Mfsd4b1',	'Slc16a10',	'Ddo',	'Foxo3',	'Prdm1',	'Fam184a',	'Pkib',	'Sowahc',	'P4ha1',	'Hkdc1',	'Gm10118',	'Jmjd1c',	'Nrbf2',	'Arid5b',	'Mrln',	'Gstt2',	'Slc19a1',	'Mbd3',	'Tjp3',	'Lta4h',	'Plxnc1',	'Socs2',	'Ube2n',	'Gm5426',	'Btg1',	'Atp2b1',	'Kitl',	'Slc6a15',	'Acss3',	'Myf5',	'Phlda1',	'Kcnc2',	'Yeats4',	'Cpsf6',	'Mdm1',	'Grip1',	'Msrb3',	'D930020B18Rik',	'BC048403',	'Srgap1',	'Avpr1a',	'Slc16a7',	'Pip4k2c',	'Nxph4',	'Smarcc2',	'A430078G23Rik',	'Efnb2',	'Irs2',	'Col4a2',	'Grtp1',	'Kbtbd11',	'Myom2',	'Angpt2',	'Zmat4',	'Tcim',	'Htra4',	'Tacc1',	'Unc5d',	'Tex15',	'Dusp4',	'Eri1',	'Cldn23',	'Prag1',	'Sorbs2',	'Galnt7',	'Sh3rf1',	'Cbr4',	'Tll1',	'Lzts1',	'Cers1',	'Tmem59l',	'Jund',	'Arhgap10',	'1700011L22Rik',	'Mmaa',	'Smad1',	'Gypa',	'Inpp4b',	'Il15',	'Gipc1',	'Adgrl1',	'Ier2',	'Nfix',	'Siah1a',	'Irx5',	'Ces1a',	'Ces1d',	'Amfr',	'Adgrg1',	'Adgrg3',	'Mmp15',	'Fhod1',	'Smpd3',	'Gm10073',	'Cdh1',	'Tango6',	'Zfhx3',	'Hp',	'Il34',	'Tmem170',	'Adamts18',	'Nudt7',	'Wwox',	'Cmip',	'Tldc1',	'Cotl1',	'Gins2',	'Irf8',	'Slc7a5',	'Banp',	'Ankrd11',	'Urb2',	'Galnt2',	'Disc1',	'Sipa1l2',	'Map3k21',	'Kcnk1',	'Pard3',	'Nrp1',	'Fam107a',	'Ptprg',	'Gm49355',	'Nid2',	'Dupd1',	'Lrmda',	'Dlg5',	'Zmiz1',	'Hesx1',	'Tkt',	'Tmem110',	'Mustn1',	'Syt15',	'Gm15217',	'Slc35f4',	'Efs',	'Nfatc4',	'Sdr39u1',	'Zmym5',	'Gja3',	'Gjb2',	'Eef1akmt1',	'Cab39l',	'Cdadc1',	'Spata13',	'Neil2',	'Zfp395',	'Ccdc25',	'Ebf2',	'Adam7',	'Adam28',	'Egr3',	'Slc39a14',	'Sftpc',	'Fgf17',	'Rcbtb2',	
                                                     'Itm2b',	'Ccdc122',	'Dnajc15',	'Rgcc',	'Pcdh9',	'Slain1',	'Gpc5',	'Dct',	'Abcc4',	'Cldn10',	'Dock9',	'Birc3',	'Gm16485',	'Sesn3',	'Eepd1',	'Ntm',	'Zbtb44',	'St3gal4',	'Fez1',	'Zfp202',	'Hspa8',	'Tbcel',	'Nectin1',	'Cbl',	'Fxyd6',	'Pafah1b2',	'Sik3',	'Ppp2r1b',	'Sik2',	'Gm32742',	'Colca2',	'Arhgap20',	'4930550C14Rik',	'Cib2',	'Wdr61',	'Lingo1',	'Csk',	'Neo1',	'Nr2e3',	'Uaca',	'Tle3',	'Smad3',	'Smad6',	'Parp16',	'Dapk2',	'Usp3',	'Tpm1',	'C2cd4b',	'Rora',	'Ice2',	'Myo1e',	'Cgnl1',	'Arpp19',	'Myo5c',	'Hcrtr2',	'Fam83b',	'Tinag',	'Ibtk',	'Me1',	'Bcl2a1d',	'Bcl2a1b',	'Atr',	'Clstn2',	'Rbp2',	'Cpne4',	'Mapkapk3',	'Gnat1',	'Bcl2a1c',	'Crtap',	'Cmtm8',	'Tgfbr2',	'Entpd3',	'Cck',	'Snrk',	'Topaz1',	'Ykt6',	'Ppia',	'Purb',	'Nacad',	'Abca13',	'Ikzf1',	'Fignl1',	'Spred2',	'Kcnmb1',	'Pank3',	'Ccnjl',	'Rnf145',	'Cby3',	'Csf2',	'Anxa6',	'Fat2',	'Zfp692',	'Srebf1',	'Shmt1',	'Dhrs7b',	'Mfap4',	'Fam83g',	'Ncor1',	'Hs3st3b1',	'Hs3st3a1',	'Myh13',	'Cfap52',	'Ntn1',	'Cd68',	'Slc16a13',	'Alox12',	'Atp2a3',	'Emc6',	'Rph3al',	'Efcab5',	'Pipox',	'Rab34',	'Slc13a2',
                                                     'Ksr1',	'Myo1d',	'Spaca3',	'Ccl8',	'Pex12',	'Hnf1b',	'1700125H20Rik',	'Tbx2',	'Pctp',	'Skap1',	'Epop',	'Cisd3',	'B230217C12Rik',	'Grb7',	'Csf3',	'Igfbp4',	'Acly',	'Arl4d',	'Meox1',	'Plcd3',	'Mrc2',	'Tanc2',	'Cyb561',	'Tex2',	'Cacng4',	'Kcnj2',	'Sox9',	'Sstr2',	'Gprc5c',	'Galk1',	'Rnf157',	'Mxra7',	'Sec14l1',	'Gm11733',	'Afmid',	'Rnf213',	'Baiap2',	'Fscn2',	'Fasn',	'Cd7',	'Calml3',	'Calm4',	'Akr1c13',	'Akr1c19',	'Zmynd11',	'Lyst',	'Gli3',	'Stard3nl',	'Trim27',	'Hist1h2ah',	'Hist1h2ag',	'Sox4',	'Bphl',	'Rpp40',	'Ssr1',	'Ofcc1',	'Tfap2a',	'Hivep1',	'Edn1',	'Phactr1',	'Cd83',	'Jarid2',	'Dtnbp1',	'Atxn1',	'Rnf144b',	'Spin1',	'Gadd45g',	'Syk',	'Auh',	'Ror2',	'Rgs14',	'Tgfbi',	'Gm45623',	'Gas1',	'Dapk1',	'Fbp1',	'2010111I01Rik',	'Ptch1',	'Ice1',	'Irx2',	'Slc12a7',	'Vcan',	'Lhfpl2',	'Ap3b1',	'Zbed3',	'Enc1',	'Gm10260',	'Cartpt',	'Cd180',	'Srek1ip1',	'Rnf180',	'Smim15',	'Elovl7',	'Il6st',	'Mcidas',	'Parp8',	'Nnt',	'Kif3c',	'Dtnb',	'Rhob',	'Sdc1',	'Fam84a',	'Trib2',	'Greb1',	'Rock2',	'Hpcal1',	'Mboat2',	'Id2',	'Rnf144a',	'Lamb1',	'Prkar2b',	'Atxn7l1',	'Etv1',	'Lrrn3',	'Egln3',	'Snx6',	'Psma6',	'Brms1l',	'Nkx2-1',	'Slc25a21',	'Mdga2',	'Gm9887',	'Pygl',	'Tmx1',	'Frmd6',	'Six1',	'Prkch',	'Esr2',	'Plekhg3',	'Rab15',	'Fut8',	'Rad51b',	'Srsf5',	'Ttc9',	'Pcnx',	'Rgs6',	'Dpf3',	'Acot1',	'Npc2',	'Isca2',	'Tgfb3',	'Nrxn3',	'Calm1',	'Ccdc88c',	'Slc24a4',	'Fam181a',	'Clmn',	'Vrk1',	'Bcl11b',	'Setd3',	'Dlk1',	'Rtl1',	'Dio3',	'Rapgef5',	'Selenop',	'Osmr',	'Lifr',	'Rai14',	'Cdh6',	'Ctnnd2',	'Dap',	'Cmbl',	'Cthrc1',	'Abra',	'Ext1',	'Mal2',	'Taf2',	'Col14a1',	'Derl1',	'Mtss1',	'Lrrc6',	'St3gal1',	'Khdrbs3',	'Ly6g',	'Gpihbp1',	'C1qtnf6',	'Gcat',	'Mgat3',	'Tnrc6b',	'Atxn10',	'Trmu',	'Celsr1',	'Gramd4',	'Lrrk2',	'Twf1',	'Nell2',	'Tmem106c',	'Col2a1',	'Tuba1c',	'Acvr1b',	'Eif4b',	'Ppp1r1a',	'Crebbp',	'4930562C15Rik',	'Emp2',	'Rmi2',	'Efcab1',	'Cldn5',	'Lamp3',	'Etv5',	'Dgkg',	'St6gal1',	'Bcl6',	'Opa1',	'Hes1',	'Bdh1',	'Zdhhc19',	'Tfrc',	'Muc4',	'Smbd1',	'Kalrn',	'Hacd2',	'Slc35a5',	'Phldb2',	'Cblb',	'Alcam',	'Nfkbiz',	'Stx19',	'Cadm2',	'Samsn1',	'Grik1',	'Cldn8',	'Sod1',	'Mis18a',	'Mrps6',	'Clic6',	'Runx1',	'Setd4',	'Ripk4',	'Fndc1',	'Dact2',	'Smoc2',	'Thbs2',	'Noxo1',	'Hs3st6',	'Cacna1h',	'Neurl1b',	'Lemd2',	'Hmga1',	'Fance',	'Clps',	'Cdkn1a',	'Cpne5',	'Mtch1',	'Rsph1',	'Slc37a1',	'H2-DMb2',	'Ddah2',	'Mpig6b',	'Prrc2a',	'Trim40',	'Adgrf5',	'1600014C23Rik',	'Tomm6',	'Mdfi',	'Shd',	'Gm20219',	'Man2a1',	
                                                     'Tgif1',	'Rasgrp3',	'Crim1',	'Atl2',	'Galm',	'Tmem178',	'Pkdcc',	'Mta3',	'Zfp36l2',	'Abcg8',	'Camkmt',	'Six2',	'Adcyap1',	'Zfp438',	'Greb1l',	'Galnt1',	'Celf4',	'Tslp',	'Stard4',	'Gm10549',	'Srp19',	'Reep5',	'Fam13b',	'Tmem173',	'E230025N22Rik',	'Pcdh1',	'Fgf1',	'Pabpc2',	'Stk32a',	'Ticam2',	'Dtwd2',	'Snx24',	'Zfp608',	'Gramd3',	'Lmnb1',	'43893',	'Slc12a2',	'Slc27a6',	'Arhgef37',	'Grpel2',	'Afap1l1',	'Fbxo38',	'Nedd4l',	'Pmaip1',	'Seh1l',	'Rab27b',	'Mbd2',	'Rpl17',	'Smad2',	'St8sia5',	'Setbp1',	'Kcng2',	'Ctdp1',	'Zfp516',	'Timm21',	'Znhit2',	'Rps6ka4',	'Mark2',	'Cd5',	'Cd6',	'Fam111a',	'Tle4',	'Gnaq',	'Aldh1a7',	'Fam189a2',	'Pip5k1b',	'Gm815',	'Vldlr',	'Rfx3',	'Plpp6',	'Ranbp6',	'Sgms1',	'Ppp1r3c',	'Tnks2',	'Myof',	'Ffar4',	'Blnk',	'Tm9sf3',	'Frat1',	'Avpi1',	'Nkx2-3',	'Entpd7',	'Scd2',	'Scd4',	'Scd1',	'Sfxn2',	'Neurl1a',	'Sh3pxd2a',	'Sorcs3',	'Rpl13a-ps1',	'Add3',	'Gpam',	'Gucy2g',	'Vwa2',	'Pnlip',	'Zfp950'))

#one gene unaccounted for 44082
integrate_epi4 <- AddModuleScore(object = integrate_epi4, features = AT2_NKX21_binding_retained_from_progenitor , ctrl = 500, name = "NKXAT2RetainedGenes", search = TRUE, random.seed = 1)
#1 gene unaccounted for 43894, 43898, 43893
FeaturePlot(object = integrate_epi4, features = 'NKXAT2RetainedGenes1', cols = c('lightgrey','red'))


Both_NKX21_binding_acquired_over_time <- list(c('Rb1cc1',	'Prdm14',	'Rdh10',	'Ube2w',	'Tmem70',	'Paqr8',	'Tram2',	'Kcnq5',	'B3gat2',	'Smap1',	'Phf3',	'Prim2',	'Fam168b',	'Hs6st1',	'Uggt1',	'Arid5a',	'Kansl3',	'Cnnm4',	'Cnnm3',	'Ankrd23',	'Gm42417',	'Tmem131',	'Eif5b',	'Rpl31',	'Tbc1d8',	'Il1r2',	'Il1r1',	'Il18r1',	'Nabp1',	'Myo1b',	'Nab1',	'Nemp2',	'Mfsd6',	'Stk17b',	'Ankrd44',	'Hspe1',	'Rftn2',	'Aox1',	'Orc2',	'Tmem237',	'Fzd7',	'Sumo1',	'Bmpr2',	'Ccnyl1',	'Fzd5',	'Kansl1l',	'Acadl',	'Atic',	'Mreg',	'Pecr',	'Smarcal1',	'Tns1',	'Cxcr2',	'Tmbim1',	'Ctdsp1',	'Vil1',	'Plcd4',	'Cnppd1',	'Tuba4a',	'Asic4',	'Sgpp2',	'Ap1s3',	'Mrpl44',	'Col4a4',	'Col4a3',	'Mff',	'A030014E15Rik',	'Cab39',	'Dis3l2',	'Ngef',	'Dgkd',	'Ugt1a6b',	'Arl4c',	'Sh3bp4',	'Agap1',	'Rab17',	'Lrrfip1',	'Ramp1',	'Ilkap',	'Hes6',	'Twist2',	'Ndufa10',	'Kif1a',	'Sned1',	'Mterf4',	'D2hgdh',	'Gin1',	'Pign',	'Tnfrsf11a',	'Dsel',	'Clasp1',	'Tfcp2l1',	'Epb41l5',	'Tmem37',	'Slc35f5',	'Lypd1',	'Nckap5',	'Tmem163',	'Ccnt2',	'Cd55',	'Gm29427',	'Fcamr',	'Mapkapk2',	'Srgap2',	'Fam72a',	'Slc45a3',	'Elk4',	'Mfsd4a',	'Cdk18',	'Nuak2',	'Ppp1r15b',	'Golt1a',	'Kiss1',	'Sox13',	'Mybph',	'Adora1',	'Rabif',	'Syt2',	'Ppp1r12b',	'Elf3',	'Rnpep',	'Ipo9',	'Csrp1',	'Phlda3',	'Camsap2',	'Nr5a2',	'Zbtb41',	'Pla2g4a',	'Rnf2',	'Edem3',	'Tsen15',	'Arpc5',	'Lamc2',	'Teddm2',	'Mr1',	'Xpr1',	'Qsox1',	'Cep350',	'Tor1aip1',	'Soat1',	'Tor3a',	'Tnr',	'4930523C07Rik',	'Cacybp',	'Rabgap1l',	'Rc3h1',	'Prdx6',	'Dnm3',	'Fmo1',	'Sft2d2',	'Dcaf6',	'Creg1',	'Mgst3',	'Pbx1',	'Nos1ap',	'Cfap126',	'Zbtb18',	'Catspere2',	'Cnst',	'Psen2',	'Stum',	'Lin9',	'Acbd3',	'Lefty2',	'Tmem63a',	'Fbxo28',	'Tlr5',	'Mia3',	'43891',	'43892',	'Iars2',	'Lyplal1',	'Tgfb2',	'Kctd3',	'Rps6kc1',	'Fam71a',	'Atf3',	'Nenf',	'Ints7',	'Lpgat1',	'Plxna2',	'Dclre1c',	'Prpf18',	'Bend7',	'Mcm10',	'Optn',	'Usp6nl',	'Il15ra',	'Mindy3',	'Rsu1',	'Nebl',	'Skida1',	'Commd3',	'Arhgap21',	'Abi1',	'Spopl',	'Noxa1',	'Tor4a',	'Ssna1',	'Man1b1',	'Uap1l1',	'Clic3',	'Tmem141',	'Camsap1',	'Nacc2',	'Gpsm1',	'Dnlz',	'Agpat2',	'Vav2',	'Rxra',	'Col5a1',	'Ralgds',	'Cel',	'Barhl1',	'Ttf1',	'Setx',	'Rapgef1',	'Sptan1',	'Wdr34',	'Miga2',	'Dolpp1',	'1700001O22Rik',	'Asb6',	'Abl1',	'Qrfp',	'Lamc3',	'Prrc2b',	'Swi5',	'Dnm1',	'Lcn2',	'Slc25a25',	'St6galnac6',	'Cdk9',	'Ttc16',	'Stxbp1',	'Fam129b',	'Garnl3',	'Angptl2',	'Zbtb34',	'Mvb12b',	'Psmd5',	'Phf19',	'Traf1',	'Hc',	'Rab14',	'Gsn',	'Dab2ip',	'Ttll11',	'Lhx6',	'Ptgs1',	'Rabgap1',	'Crb2',	'Rbm43',	'Nmi',	'Gpd2',	'Cytip',	'Acvr1',	'Pkp4',	'Tanc1',	'Wdsub1',
                                                'Ly75',	'Itgb6',	'Rbms1',	'Grb14',	'Bbs5',	'Ccdc173',	'Gm26558',	'Tlk1',	'Dcaf17',	'Map3k20',	'Cir1',	'Gpr155',	'Chrna1',	'Agps',	'Osbpl6',	'Prkra',	'Plekha3',	'Ccdc141',	'Nckap1',	'Ctnnd1',	'Tnks1bp1',	'Aplnr',	'Olfr1274-ps',	'Rapsn',	'Mybpc3',	'Madd',	'Atg13',	'Pex16',	'1700029I15Rik',	'Cry2',	'Prdm11',	'Cd82',	'Traf6',	'Trim44',	'Slc1a2',	'Cd44',	'Pdhx',	'Cat',	'Nat10',	'Hipk3',	'Mettl15',	'Aven',	'BC052040',	'Thbs1',	'Eif2ak4',	'Pak6',	'Bahd1',	'Rmdn3',	'Chac1',	'Exd1',	'Ehd4',	'Vps39',	'Tgm5',	'Ckmt1',	'Hypk',	'Secisbp2l',	'Cops2',	'Hdc',	'Gabpb1',	'Usp50',	'Trpm7',	'Stard7',	'Acoxl',	'Bcl2l11',	'Tmem87b',	'Slc20a1',	'Snrpb',	'Vps16',	'Spef1',	'Ap5s1',	'Rassf2',	'Slc23a2',	'Cds2',	'Trmt6',	'Plcb4',	'Lamp5',	'Jag1',	'Kif16b',	'Dstn',	'Rrbp1',	'Rin2',	'Ralgapa2',	'Kiz',	'Nxt1',	'Apmap',	'Pygb',	'Fam110a',	'Srxn1',	'Sox12',	'Defb20',	'H13',	'Mcts2',	'Bcl2l1',	'Asxl1',	'Commd7',	'Bpifb1',	'1700003F12Rik',	'Pxmp4',	'Chmp4b',	'Raly',	'Eif2s2',	'Dynlrb1',	'Pigu',	'Trp53inp2',	'Acss2',	'Myh7b',	'Uqcc1',	'Spag4',	'Scand1',	'Cnbd2',	'Epb41l1',	'Myl9',	'Src',	'Tti1',	'Rprd1b',	'Tgm2',	'Plcg1',	'Mybl2',	'Ada',	'Wfdc12',	'Sdc4',	'Pigt',	'Wfdc2',	'Snx21',	'Zfp335',	'Slc35c2',	'Elmo2',	'Eya2',	'Sulf2',	'Arfgef2',	'Cse1l',	'Snai1',	'Ube2v1',	'Tmem189',	'Ptpn1',	'Ripor3',	'Adnp',	'Nfatc2',	'Bmp7',	'Gm20721',	'Prelid3b',	'Taf4',	'Lsm14b',	'Osbpl2',	'Adrm1',	'Lama5',	'Cables2',	'Ythdf1',	'Ptk6',	'Helz2',	'Zfp512b',	'Myt1',	'Wdr13',	'Gm14862',	'Bcor',	'1810030O07Rik',	'Cask',	'Fundc1',	'Kdm6a',	'Araf',	'Lonrf3',	'Slc25a43',	'Lamp2',	'Rap2c',	'Slc9a6',	'Fhl1',	'Fmr1',	'Gabrq',	'Trex2',	'Bgn',	'Dusp9',	'Taz',	'Pls3',	'Tbl1x',	'Prkx',	'Pdk3',	'Ophn1',	'Efnb1',	'Pja1',	'Phka1',	'Magee1',	'Atrx',	'Cenpi',	'Morf4l2',	'Tbc1d8b',	'Prps1',	'Tsc22d3',	'Il13ra2',	'Shroom2',	'Ap1s2',	'Msl3',	'Tpd52',	'Zbtb10',	'Zfp704',	'Impa1',	'Chmp4c',	'Mtfr1',	'Dnajc5b',	'Gyg',	'Tbl1xr1',	'Ect2',	'Nceh1',	'Tmem212',	'Eif5a2',	'Mecom',	'Gpr160',	'Prkci',	'Skil',	'Zmat3',	'Gnb4',	'Ttc14',	'Dcun1d1',	'Acad9',	'Anxa5',	'1810062G17Rik',	'Fgf2',	'Spry1',	'Ankrd50',	'Larp1b',	'Noct',	'Rab33b',	'Mgst2',	'Smad9',	'Spg20',	'Nbea',	'Tm4sf4',	'Igsf10',	'Arhgef26',	'Ssr3',	'Ptx3',	'Mfsd1',	'Schip1',	'Il12a',	'Rapgef2',	'Fnip2',	'Ppid',	'Plrg1',	'Tlr2',	'Tmem131l',	'Gm37240',	'Fbxw7',	'Gatb',	'Fam160a1',	'Rps3a1',	'Mab21l2',	'Mef2d',	'Gm38392',	'Tsacc',	'Tmem79',	'Slc25a44',	'Sema4a',	'Mex3a',	'Rxfp4',	'Rit1',	'Gon4l',	'Msto1',	'Dap3',	'Slc50a1',	'Ube2q1',	
                                                'Il6ra',	'Rps27',	'Dennd4b',	'Snapin',	'Tdrkh',	'Snx27',	'Cgn',	'Selenbp1',	'Zfp687',	'Sema6c',	'Cers2',	'Golph3l',	'Ensa',	'Mcl1',	'Adamtsl4',	'Ciart',	'Anp32e',	'Rnf115',	'Pdzk1',	'Acp6',	'Bcl9',	'Chd1l',	'Fmo5',	'Notch2',	'Adam30',	'Reg4',	'Hmgcs2',	'Cd101',	'Ptgfrn',	'Atp1a1',	'Tspan2',	'Bcas2',	'Syt6',	'Olfml3',	'Hipk1',	'Dclre1b',	'Lrig2',	'Slc16a1',	'St7l',	'Wnt2b',	'Rap1a',	'I830077J02Rik',	'Dennd2d',	'Lamtor5',	'Csf1',	'Gstm6',	'Sort1',	'Mybphl',	'Stxbp3',	'Fndc7',	'Prpf38b',	'Extl2',	'Trmt13',	'Slc44a3',	'F3',	'Abca4',	'Bcar3',	'Fnbp1l',	'Sec24d',	'Camk2d',	'Npnt',	'Gstcd',	'Tet2',	'Cisd2',	'Nfkb1',	'Slc39a8',	'Ppp3ca',	'Adh5',	'Eif4e',	'Tspan5',	'Rap1gds1',	'Hs2st1',	'Clca2',	'Ctbs',	'Spata1',	'Ttll7',	'Usp33',	'Srsf11',	'Lrrc40',	'Tmem68',	'Lyn',	'Nsmaf',	'Tox',	'Rab2a',	'Chd7',	'2610301B20Rik',	'Plekhf2',	'Dpy19l4',	'Pdp1',	'Tmem64',	'Osgin2',	'Atp6v0d2',	'Ggh',	'Tstd3',	'Fbxl4',	'Map3k7',	'Bach2',	'Ube2j1',	'Pnrc1',	'Ddx58',	'Smim27',	'B4galt1',	'Bag1',	'Aqp7',	'Ube2r2',	'Dcaf12',	'Fam219a',	'Fam214b',	'Unc13b',	'Rusc2',	'Ccdc107',	'Car9',	'Tln1',	'Npr2',	'Olfr71',	'Spaar',	'Glipr2',	'Zcchc7',	'Slc25a51',	'Shb',	'Xpa',	'Anp32b',	'Coro2a',	'Sec61b',	'Nr4a3',	'Stx17',	'Msantd3',	'Ppp3r2',	'Abca1',	'Slc44a1',	'Fktn',	'Zfp462',	'Klf4',	'Tmem245',	'Ptpn3',	'Zkscan16',	'Ugcg',	'Snx30',	'Slc46a2',	'Atp6v1g1',	'Tmem268',	'Tnfsf15',	'Tnc',	'Trim32',	'Rasef',	'Kdm4c',	'Cer1',	'Ttc39b',	'Snapc3',	'Focad',	'Jun',	'Atg4c',	'Ror1',	'Jak1',	'Slc35d1',	'Oma1',	'Prkaa2',	'Gpx7',	'Btf3l4',	'Osbpl9',	'Ttc39a',	'Dmrta2',	'Bend5',	'Gm12830',	'Stil',	'Pdzk1ip1',	'Cyp4b1',	'Efcab14',	'Mob3c',	'Lurap1',	'Tspan1',	'Akr1a1',	'Plk3',	'Rps8',	'Atp6v0b',	'Dph2',	'Ipo13',	'Artn',	'St3gal3',	'Ptprf',	'Szt2',	'Elovl1',	'AU022252',	'Rimkla',	'Guca2a',	'Slfnl1',	'Ctps',	'Kcnq4',	'Cap1',	'Macf1',	'Ndufs5',	'Inpp5b',	'Yrdc',	'Zc3h12a',	'Stk40',	'Sh3d21',	'Thrap3',	'Tekt2',	'AU040320',	'Tmem35b',	'Zfp362',	'Azin2',	'Ak2',	'Rnf19b',	'Zbtb8a',	'Dcdc2b',	'Ptp4a2',	'Pef1',	'Hcrtr1',	'Tinagl1',	'Fabp3',	'Sdc3',	'Tmem200b',	'Sesn2',	'Ptafr',	'Slc9a1',	'Sfn',	'Pigv',	'Arid1a',	'Cd52',	'Sh3bgrl3',	'Cep85',	'Slc30a2',	'Stmn1',	'Mtfr1l',	'Selenon',	'Man1c1',	'Clic4',	'Ifnlr1',	'Pnrc2',	'Gale',	'Asap3',	'Luzp1',	'Lactbl1',	'Wnt4',	'Cdc42',	'Hspg2',	'Rap1gap',	'Ece1',	'Eif4g3',	'Hp1bp3',	'Pink1',	'Camk2n1',	'Otud3',	'Capzb',	'Szrd1',	'Fbxo42',	'Epha2',	'Spen',	'Tmem82',	'Plekhm2',	'Agmat',	'Efhd2',	'Dhrs3',	'Tnfrsf1b',	'Fv1',	'Mfn2',	'Plod1',	'2510039O18Rik',
                                                'Mthfr',	'Fbxo6',	'Fbxo2',	'Ubiad1',	'Casz1',	'Pgd',	'Ctnnbip1',	'Clstn1',	'Slc25a33',	'Spsb1',	'H6pd',	'Gpr157',	'Rere',	'Errfi1',	'Park7',	'Dnajc11',	'Thap3',	'Phf13',	'Klhl21',	'Plekhg5',	'Tnfrsf25',	'Acot7',	'Kcnab2',	'Dffb',	'Lrrc47',	'Smim1',	'Tprgl',	'Prdm16',	'B230104I21Rik',	'Tnfrsf14',	'Morn1',	'Faap20',	'Gabrd',	'Gnb1',	'Mxra8',	'B3galt6',	'Isg15',	'Ankib1',	'Fzd1',	'Cdk14',	'Cldn12',	'Cfap69',	'Tmem243',	'Gnat3',	'Rsbn1l',	'Gsap',	'Napepld',	'Srpk2',	'Tomm7',	'Atg9b',	'Tmub1',	'Asb10',	'Nub1',	'Rheb',	'Prkag2',	'Kmt2c',	'Paxip1',	'Insig1',	'Rbm33',	'Shh',	'Ost4',	'Khk',	'Preb',	'Gm29609',	'Mrpl33',	'Rbks',	'Nelfa',	'Mxd4',	'Zfyve28',	'Fam193a',	'Mfsd10',	'Msantd1',	'Lrpap1',	'Acox3',	'Sh3tc1',	'Psapl1',	'Tada2b',	'Tbc1d14',	'D5Ertd579e',	'Man2b2',	'Zbtb49',	'Slc2a9',	'Wdr1',	'Zfp518b',	'Rab28',	'Cd38',	'Anapc4',	'Sel1l3',	'Smim20',	'Cckar',	'Stim2',	'Rell1',	'Tlr1',	'Tmem156',	'N4bp2',	'Rbm47',	'Apbb2',	'Atp8a1',	'Nipal1',	'Fryl',	'Dcun1d4',	'Usp46',	'Chic2',	'Clock',	'Hopx',	'Spink2',	'Rufy3',	'Mob1b',	'Cox18',	'Ankrd17',	'Btc',	'Rchy1',	'Cdkl2',	'Cxcl10',	'Scarb2',	'Stbd1',	'Anxa3',	'Paqr3',	'Prkg2',	'Enoph1',	'Nkx6-1',	'Cds1',	'Wdfy3',	'Ptpn13',	'1700016H13Rik',	'Aff1',	'Klhl8',	'Nudt9',	'Gm17660',	'Zfp326',	'Fam69a',	'Pcgf3',	'Cplx1',	'Gak',	'Idua',	'Chfr',	'Pgam5',	'Pxmp2',	'Galnt9',	'Noc4l',	'Ddx51',	'Ep400',	'Pitpnb',	'Tpst2',	'2900026A02Rik',	'Coro1c',	'Kctd10',	'Ube3b',	'Fam222a',	'Ankrd13a',	'Oasl1',	'Rnf10',	'Gatc',	'Rab35',	'Cit',	'Prkab1',	'Suds3',	'Taok3',	'Fbxo21',	'Tesc',	'Tpcn1',	'Hectd4',	'Aldh2',	'Atxn2',	'Sh2b3',	'Myl2',	'Atp2a2',	'Ift81',	'Rnf34',	'Kdm2b',	'Orai1',	'Rhof',	'Diablo',	'Gm49027',	'Clip1',	'Zcchc8',	'Hcar2',	'Hip1r',	'Vps37b',	'Abcb9',	'Kmt5a',	'Snrnp35',	'Rilpl1',	'Zfp664',	'Rflna',	'Ncor2',	'Scarb1',	'Ubc',	'Glt1d1',	'Ran',	'Nupr1l',	'Tpst1',	'Kctd7',	'Rabgef1',	'Tmem248',	'Auts2',	'Ncf1',	'Gtf2i',	'Lat2',	'Eif4h',	'Stx1a',	'Tmem120a',	'Srrm3',	'Hspb1',	'Dtx2',	'Orai2',	'Sh2b2',	'Cux1',	'Gm20605',	'Sap25',	'Irs3',	'Cyp3a13',	'Mblac1',	'Sun1',	'Gpr146',	'Gper1',	'Mafk',	'Elfn1',	'Mrm2',	'Snx8',	'Eif3b',	'Ap5z1',	'Papolb',	'Actb',	'Rac1',	'Cyth3',	'Usp42',	'Ccz1',	'Ocm',	'Bhlha15',	'Bri3',	'Smurf1',	'Arpc1b',	'Zkscan14',	'Zfp655',	'Wasf3',	'Usp12',	'Rasl11a',	'Lnx2',	'Slc7a1',	'Ubl3',	'Gm42791',	'N4bp2l1',	'Stard13',	'Casd1',	'Ppp1r9a',	'Pon1',	'Slc25a13',	'C1galt1',	'Ppp1r3a',	'Foxp2',	'Mdfic',	'Tes',	'Tspan12',	'Cadps2',	'Wasl',	'Rbm28',	'Calu',	'Kcp',
                                                'Nrf1',	'Ube2h',	'Klf14',	'Mkln1',	'Podxl',	'Plxna4',	'Akr1b7',	'Agbl3',	'Tmem140',	'Wdr91',	'Stra8',	'Fam180a',	'Svopl',	'Zc3hav1',	'Ubn2',	'Clec2l',	'Hipk2',	'Tbxas1',	'Kdm7a',	'Tra2a',	'Fam221a',	'Osbpl3',	'Hibadh',	'Tax1bp1',	'Cpvl',	'Prr15',	'Wipf3',	'Plekha8',	'Znrf2',	'Nod1',	'Mindy4',	'Aqp1',	'Tnip3',	'Ndnf',	'Gng12',	'Gadd45a',	'Krcc1',	'Rnf103',	'Kdm3a',	'Reep1',	'Atoh8',	'Sftpb',	'Rnf181',	'Vamp8',	'Mat2a',	'Sh2d6',	'Tgoln1',	'Suclg1',	'Mrpl19',	'Eva1a',	'Pole4',	'Hk2',	'Dok1',	'Loxl3',	'Mogs',	'Dctn1',	'Slc4a5',	'Mthfd2',	'Mob1a',	'Bola3',	'Tet3',	'Smyd5',	'Add2',	'Pcbp1',	'Asprv1',	'Mxd1',	'Snrnp27',	'Gmcl1',	'Aak1',	'Antxr1',	'Rab43',	'Cnbp',	'Rab7',	'Rpn1',	'Sec61a1',	'Abtb1',	'Tpra1',	'Plxna1',	'Chchd6',	'Txnrd3',	'Zxdc',	'Klf15',	'Aldh1l1',	'Slc41a3',	'Nup210',	'Hdac11',	'Fbln2',	'Tmem43',	'Xpc',	'Slc6a6',	'Nr2c2',	'Mrps25',	'Prickle2',	'Adamts9',	'Gm15737',	'Magi1',	'Suclg2',	'Mitf',	'Rybp',	'Gxylt2',	'Itpr1',	'Bhlhe40',	'Edem1',	'Rad18',	'Thumpd3',	'Lhfpl4',	'Prrt3',	'Irak2',	'Tatdn2',	'Atp2b2',	'Atg7',	'Vgll4',	'Tamm41',	'Mkrn2',	'Raf1',	'Tmem40',	'Rpl32',	'43898',	'Hnrnpf',	'Fxyd4',	'Rasgef1a',	'Csgalnact2',	'Adipor2',	'Ninj2',	'Il17ra',	'Cecr2',	'Pex26',	'Foxj2',	'Lpcat3',	'Chd4',	'Nop2',	'Gapdh',	'Ltbr',	'Scnn1a',	'Plekhg6',	'Cd9',	'Tspan9',	'Fkbp4',	'Cd69',	'Gabarapl1',	'Lrp6',	'Dusp16',	'Crebl2',	'Ddx47',	'Gprc5a',	'Gsg1',	'Strap',	'Plekha5',	'Pde3a',	'Recql',	'Ldhb',	'Etnk1',	'Lrmp',	'Bhlhe41',	'Ccdc91',	'Amn1',	'Cacng7',	'Oscar',	'Leng9',	'Rdh13',	'Tnnt1',	'Ppp6r1',	'Shisa7',	'Zfp580',	'Zfp787',	'Zfp444',	'Selenow',	'Nop53',	'Bicra',	'Ccdc9',	'Bbc3',	'Sae1',	'Zc3h4',	'Slc1a5',	'Ppp5c',	'Opa3',	'Vasp',	'Ercc2',	'Mark4',	'Ppp1r37',	'Nectin2',	'Smg9',	'Arhgef1',	'Grik5',	'Pou2f2',	'Dedd2',	'Gsk3a',	'Ceacam1',	'Tmem91',	'Tgfb1',	'Cyp2b10',	'Egln2',	'Numbl',	'Sptbn4',	'Akt2',	'Zfp36',	'Med29',	'Rinl',	'Actn4',	'Catsperg1',	'2200002D01Rik',	'Dpf1',	'Sipa1l3',	'Sdhaf1',	'Scn1b',	'Lsm14a',	'Cebpg',	'Faap24',	'Pdcd5',	'Dpy19l3',	'Vsig10l',	'Mybpc2',	'Spib',	'Kcnc3',	'Myh14',	'Ap2a1',	'Fcgrt',	'Aldh16a1',	'Gys1',	'Bcat2',	'Rasip1',	'Kdelr1',	'Ccdc114',	'Sergef',	'Uevld',	'Dbx1',	'Gas2',	'Nipa2',	'Herc2',	'Atp10a',	'Klf13',	'E030018B13Rik',	'Fan1',	'Selenos',	'Lysmd4',	'Ttc23',	'Synm',	'Igf1r',	'Nr2f2',	'Rgma',	'Chd2',	'Fam174b',	'Slco3a1',	'Akap13',	'Klhl25',	'Aen',	'Hapln3',	'Abhd2',	'Pex11a',	'Zfp710',	'Idh2',	'Fes',	'Iqgap1',	'Slc28a1',	'Rps17',	'Cemip',	'Abhd17c',	'Fah',	'Tmem135',
                                                'Prss23',	'Picalm',	'Ccdc83',	'Kctd21',	'Alg8',	'Ndufc2',	'Ints4',	'Acer3',	'Mogat2',	'Map6',	'Arrb1',	'Tpbgl',	'Rnf169',	'Ucp3',	'Ucp2',	'Arhgef17',	'Pde2a',	'Clpb',	'Il18bp',	'Nup98',	'Trim6',	'Smpd1',	'Apbb1',	'Hpx',	'Trim3',	'Cyb5r2',	'Rpl27a',	'St5',	'Nrip3',	'Dennd5a',	'Tmem41b',	'Ipo7',	'Wee1',	'Swap70',	'Ampd3',	'Mrvi1',	'Mical2',	'Parva',	'Rassf10',	'Arntl',	'Gm4353',	'Plekha7',	'Rps13',	'Pik3c2a',	'Nucb2',	'Arl6ip1',	'Smg1',	'Coq7',	'Gde1',	'Knop1',	'Gprc5b',	'Lyrm1',	'Abca14',	'Usp31',	'Scnn1b',	'Lcmt1',	'Kdm8',	'Il4ra',	'D430042O09Rik',	'Gsg1l',	'Atxn2l',	'Sult1a1',	'Tmem219',	'Kctd13',	'Kif22',	'Zfp768',	'Ctf1',	'Fbxl19',	'Kat8',	'Prss8',	'Armc5',	'BC017158',	'Rgs10',	'Bag3',	'Inpp5f',	'Wdr11',	'Fgfr2',	'Nsmce4a',	'Btbd16',	'Lhpp',	'Fam53b',	'Dhx32',	'Nps',	'9430038I01Rik',	'Glrx3',	'Ppp2r2d',	'Dpysl4',	'Pwwp2b',	'Zfp941',	'Scgb1c1',	'Cox8b',	'Irf7',	'Tmem80',	'Pnpla2',	'Mob2',	'Dusp8',	'Tspan32',	'Kcnq1',	'Slc22a18',	'Osbpl5',	'Shank2',	'Ppfia1',	'Fadd',	'Ano1',	'Ccnd1',	'1810010D01Rik',	'Mrgprf',	'Ipcef1',	'Cnksr3',	'Ginm1',	'Sash1',	'Utrn',	'Ltv1',	'Hivep2',	'Heca',	'Reps1',	'Arfgef3',	'Tnfaip3',	'Il22ra2',	'Map3k5',	'Pde7b',	'Ahi1',	'Slc18b1',	'Akap7',	'Rnf146',	'Tspyl1',	'Tspyl4',	'Mfsd4b4',	'Mfsd4b5',	'Slc16a10',	'Fig4',	'Armc2',	'Foxo3',	'Sec63',	'Scml4',	'Bend3',	'Gm9803',	'Atg5',	'Prep',	'Ascc3',	'Mcm9',	'Fam184a',	'Msl3l2',	'Lims1',	'Edar',	'Sh3rf3',	'44084',	'P4ha1',	'Gm10273',	'Oit3',	'Mcu',	'Ddit4',	'Eif4ebp2',	'Lrrc20',	'Aifm2',	'Tspan15',	'Tacr2',	'Hk1',	'4930507D05Rik',	'Ccar1',	'Mypn',	'Lrrtm3',	'Nrbf2',	'Rtkn2',	'Arid5b',	'Rhobtb1',	'A930033H14Rik',	'Tfam',	'Bcr',	'Upb1',	'Susd2',	'Cabin1',	'Gstt1',	'Gstt2',	'Smarcb1',	'S100b',	'Pcnt',	'2610028H24Rik',	'Lss',	'Pttg1ip',	'1810043G02Rik',	'Trappc10',	'Agpat3',	'Cdc34',	'Bsg',	'Misp',	'Tmem259',	'Abca7',	'Arhgap45',	'Polr2e',	'Gpx4',	'Dazap1',	'Rps15',	'2310011J03Rik',	'Adamtsl5',	'Btbd2',	'Mknk2',	'Ap3d1',	'Dot1l',	'Plekhj1',	'Gng7',	'Thop1',	'4930442H23Rik',	'Hmg20b',	'Smim24',	'Nfic',	'Aes',	'Glt8d2',	'Appl2',	'Tcp11l2',	'Polr3b',	'Rfx4',	'Syn3',	'Timp3',	'Nt5dc3',	'Parpbp',	'Scyl2',	'Uhrf1bp1l',	'Cdk17',	'Hal',	'Nr2c1',	'Tmcc3',	'Plxnc1',	'Socs2',	'Nudt4',	'Atp2b1',	'Poc1b',	'Dusp6',	'Zdhhc17',	'Krr1',	'Atxn7l3b',	'Tbc1d15',	'Rab21',	'Frs2',	'Mdm2',	'Rap1b',	'Msrb3',	'Wif1',	'Gns',	'Tbk1',	'Slc16a7',	'Atp23',	'Agap2',	'B4galnt1',	'Rbms2',	'Spryd4',	'Gdf11',	'Itga7',	'Stxbp2',	'Lig4',	'Irs2',	'Col4a1',	'Col4a2',	'Naxd',	'Cars2',	
                                                'Tex29',	'Tubgcp3',	'Atp11a',	'Mcf2l',	'Adprhl1',	'Dcun1d2',	'Tmem255b',	'Gas6',	'Rasa3',	'Champ1',	'Cln8',	'Angpt2',	'Atp7b',	'Nkx6-3',	'Adam32',	'Adam9',	'Plpp5',	'Hook3',	'Brf2',	'Rab11fip1',	'Got1l1',	'Tti2',	'Mak16',	'Wrn',	'Smim18',	'Rbpms',	'Eri1',	'Dlc1',	'Pdgfrl',	'Asah1',	'Fat1',	'Ccdc110',	'Ankrd37',	'Snx25',	'Cfap97',	'Irf2',	'Enpp6',	'Cdkn2aip',	'Wwc2',	'Neil3',	'Sh3rf1',	'Tma16',	'Lzts1',	'Yjefn3',	'Gatad2a',	'Nr2c2ap',	'Ell',	'Rab3a',	'Ccdc124',	'Use1',	'Pgls',	'Klf2',	'Med26',	'Arhgap10',	'Otud4',	'Gab1',	'Dnajb1',	'Pkn1',	'Ptger1',	'Nanos3',	'Mri1',	'Dnase2a',	'Junb',	'Hook2',	'Vps35',	'Orc6',	'Dnaja2',	'Itfg1',	'Siah1a',	'N4bp1',	'Brd7',	'Chd9',	'Rpgrip1l',	'Irx5',	'Irx6',	'Slc12a3',	'Herpud1',	'Nlrc5',	'Cpne2',	'Cx3cl1',	'Ccdc102a',	'Mmp15',	'Csnk2a2',	'Ccdc113',	'Cmtm4',	'Terb1',	'Cbfb',	'D230025D16Rik',	'Nol3',	'4931428F04Rik',	'Exoc3l',	'Lrrc29',	'Fhod1',	'Kctd19',	'Lrrc36',	'Tppp3',	'Hsd11b2',	'Atp6v0d1',	'Ripor1',	'Ranbp10',	'Nfatc3',	'Gm10073',	'Tango6',	'Cyb5b',	'Hp',	'Ap1g1',	'Marveld3',	'Ldhd',	'Zfp1',	'Bcar1',	'Tmem231',	'Terf2ip',	'Syce1l',	'Dynlrb2',	'Cdyl2',	'Bco1',	'Gan',	'Plcg2',	'Sdr42e1',	'Hsbp1',	'Mlycd',	'Osgin1',	'Hsdl1',	'Wfdc1',	'Cotl1',	'Klhl36',	'Gse1',	'Gins2',	'Map1lc3b',	'Zcchc14',	'Piezo1',	'Pabpn1l',	'Ankrd11',	'Spata2l',	'Rhou',	'Abcb10',	'Capn9',	'Gnpat',	'Sipa1l2',	'Map10',	'Ntpcr',	'Coa6',	'Irf2bp2',	'Tomm20',	'Nrp1',	'Itgb1',	'Pxk',	'Gm49355',	'Psmd6',	'Ngly1',	'Top2b',	'Rpl15',	'Ube2e1',	'Ecd',	'Usp54',	'Fut11',	'Vcl',	'Adk',	'Kat6b',	'Samd8',	'Lrmda',	'Zmiz1',	'Ppif',	'Anxa11',	'Wnt5a',	'Cacna1d',	'Tkt',	'Prkcd',	'Rft1',	'Sfmbt1',	'Tmem110',	'Nisch',	'Phf7',	'Gprin2',	'Fam25c',	'Bmpr1a',	'Opn4',	'Wapl',	'Ccser2',	'Sh2d4b',	'Tspan14',	'Fam213a',	'Sftpd',	'Fermt2',	'Bmp4',	'Gmfb',	'Samd4',	'Lgals3',	'Dlgap5',	'Fbxo34',	'Atg14',	'Ktn1',	'Eddm3b',	'Hnrnpc',	'Ppp1r3e',	'Bcl2l2',	'Myh6',	'Nrl',	'Tm9sf1',	'Tgm1',	'Dhrs1',	'Khnyn',	'Cenpj',	'Il17d',	'Gm6904',	'Cdadc1',	'Ebpl',	'Ctsb',	'Fdft1',	'Fam167a',	'Ints9',	'Zfp395',	'Nuggc',	'Dpysl2',	'Ppp2r2a',	'Dock5',	'Chmp7',	'9930012K11Rik',	'Pdlim2',	'Sorbs3',	'Ppp3cc',	'Lgi3',	'Reep4',	'Fam160b2',	'Fndc3a',	'Itm2b',	'Lrch1',	'Siah3',	'Tpt1',	'Nufip1',	'Tsc22d1',	'Dnajc15',	'Dgkh',	'Vwa8',	'Rgcc',	'Wbp4',	'Diaph3',	'Tdrd3',	'Lmo7',	'Acod1',	'Cln5',	'Fbxl3',	'Abcc4',	'Dzip1',	'Dnajc3',	'Mbnl2',	'Farp1',	'Stk24',	'Gpr18',	'Tm9sf2',	'Ggact',	'Tmem123',	'Birc3',	'Maml2',	'Amotl1',	'Piwil4',	'Fut4',	'Zfp426',	'S1pr2',	'Icam1',
                                                'Icam5',	'Pde4a',	'Ilf3',	'Dnm2',	'Spc24',	'Kank2',	'Zfp872',	'Zfp810',	'Herpud2',	'Adamts8',	'Zbtb44',	'Prdm10',	'Ets1',	'St3gal4',	'Esam',	'Sorl1',	'Nectin1',	'Rnf26',	'Bcl9l',	'Ddx6',	'Phldb1',	'Cadm1',	'Nnmt',	'Zbtb16',	'Usp28',	'Ncam1',	'Plet1',	'Dlat',	'Dixdc1',	'Cryab',	'Ppp2r1b',	'Fdx1',	'Rdx',	'Zc3h12c',	'Acat1',	'Dmxl2',	'Cib2',	'Idh3a',	'Dnaja4',	'Wdr61',	'Hykk',	'Chrnb4',	'Snx33',	'Imp3',	'Ptpn9',	'Cox5a',	'Clk3',	'Sema7a',	'Tmem202',	'Gramd2',	'Nr2e3',	'Anp32a',	'Fem1b',	'Smad3',	'Smad6',	'Rab11a',	'Dennd4a',	'Hacd3',	'Dpp8',	'Parp16',	'Rasl12',	'Mtfmt',	'Plekho2',	'Zfp609',	'Snx22',	'Herc1',	'Lactb',	'Tpm1',	'Tln2',	'Ice2',	'Gcnt3',	'Adam10',	'Mns1',	'Nedd4',	'Ccpg1os',	'Ccpg1',	'Khdc3',	'Fam214a',	'Tmod3',	'Hmgcll1',	'Mlip',	'Lrrc1',	'Gclc',	'Elovl5',	'Ick',	'Gsta4',	'Slc17a5',	'Myo6',	'Ctsh',	'Tbc1d2b',	'Plscr5',	'Xrn1',	'Tfdp2',	'Zbtb38',	'Il20rb',	'Pccb',	'Cep63',	'Amotl2',	'Ryk',	'Rab6b',	'Srprb',	'1300017J02Rik',	'Cdv3',	'Nudt16',	'Atp2c1',	'Ppm1m',	'Alas1',	'Dusp7',	'Abhd14b',	'Pcbp4',	'Parp3',	'Rrp9',	'Tex264',	'Mapkapk3',	'Cish',	'Hemk1',	'Zmynd10',	'Tusc2',	'Sema3b',	'Rbm6',	'Mst1',	'Apeh',	'Dag1',	'Nicn1',	'Gpx1',	'P4htm',	'Prkar2a',	'Ip6k2',	'Celsr3',	'Shisa5',	'Plxnb1',	'Cdc25a',	'Dhx30',	'Cspg5',	'Scap',	'Ptpn23',	'Nradd',	'Ccdc12',	'Rtp3',	'Lrrfip2',	'Clasp2',	'Dync1li1',	'Cmtm8',	'Tgfbr2',	'Itga9',	'Ctdspl',	'Vill',	'Acaa1a',	'Xylb',	'Exog',	'Gorasp1',	'Ctnnb1',	'Trak1',	'Cck',	'Higd1a',	'Snrk',	'Ano10',	'Abhd5',	'Zdhhc3',	'Slc6a20a',	'Fyco1',	'Patz1',	'Pik3ip1',	'Limk2',	'Rnf185',	'Morc2a',	'Sec14l4',	'Mtfp1',	'Sec14l2',	'Lif',	'Ascc2',	'Uqcr10',	'Ap1b1',	'Gas2l1',	'Dbnl',	'Pgam2',	'Gck',	'Ykt6',	'Ogdh',	'Ppia',	'Purb',	'Grb10',	'Cobl',	'Egfr',	'Fbxo48',	'Pno1',	'Cep68',	'Sertad2',	'Peli1',	'Zrsr1',	'Efemp1',	'Psme4',	'Erlec1',	'Bod1',	'Cpeb4',	'Mpg',	'Fbxw11',	'Slit3',	'Rars',	'Wwc1',	'Mat2b',	'Nudcd2',	'Ccng1',	'Adra1b',	'Rnf145',	'Clint1',	'Rack1',	'Gm12184',	'Olfr1393',	'Rnf130',	'Sqstm1',	'Mgat4b',	'Col23a1',	'Phykpl',	'Sec24a',	'Sar1b',	'Jade2',	'Ube2b',	'Cdkl3',	'Ppp2ca',	'Tcf7',	'Hspa4',	'Aff4',	'Gm9945',	'Kif3a',	'Irf1',	'Gm12216',	'Slc22a4',	'P4ha2',	'Acsl6',	'Fnip1',	'Cdc42se2',	'Tnip1',	'Anxa6',	'Fam114a2',	'Mfap3',	'Larp1',	'Irgm2',	'4930438A08Rik',	'Zfp672',	'Trim11',	'Iba57',	'2310033P09Rik',	'Arf1',	'Wnt3a',	'Mprip',	'Srebf1',	'Myo15',	'Mief2',	'Tmem11',	'Natd1',	'Map2k3',	'Aldh3a1',	'Slc47a2',	'B9d1',	'Epn2',	'Ulk2',	'Akap10',	'Adora2b',	'Trim16',	'Tekt3',	'Pmp22',	
                                                'Hs3st3b1',	'Cox10',	'Hs3st3a1',	'Elac2',	'1700086D15Rik',	'Myocd',	'Zkscan6',	'Gas7',	'Stx8',	'Ndel1',	'Per1',	'Cntrob',	'Tmem102',	'Gps2',	'Slc16a11',	'Med11',	'Mink1',	'Inca1',	'Zfp3',	'Pimreg',	'Pitpnm3',	'Med31',	'Mybbp1a',	'Cyb5d2',	'Atp2a3',	'Camkk1',	'Ncbp3',	'Shpk',	'Mettl16',	'Mnt',	'Sgsm2',	'Hic1',	'Rtn4rl1',	'Rpa1',	'Serpinf1',	'Wdr81',	'Rilp',	'Scarf1',	'Slc43a2',	'Myo1c',	'Ywhae',	'Doc2b',	'Rph3al',	'Vps53',	'Dbil5',	'Myo18a',	'Gm10277',	'Phf12',	'Sdf2',	'Poldip2',	'Nos2',	'Lgals9',	'Myo1d',	'Ccl8',	'Rasl10b',	'Mmp28',	'Ccl5',	'Heatr6',	'Dusp14',	'Lhx1',	'Ppm1d',	'Rnft1',	'Tubd1',	'Vmp1',	'Cltc',	'Ypel2',	'Gdpd1',	'Trim25',	'Gm45716',	'Stxbp4',	'Cox11',	'Spag9',	'Wfikkn2',	'Cacna1g',	'Spata20',	'Mycbpap',	'Rsad1',	'Fam117a',	'Slc35b1',	'B130006D01Rik',	'Phospho1',	'Abi3',	'Snx11',	'Cdk5rap3',	'Pnpo',	'Sp2',	'Tbkbp1',	'Npepps',	'Mrpl45',	'Gpr179',	'Arhgap23',	'Cwc25',	'B230217C12Rik',	'Fbxl20',	'Med1',	'Ppp1r1b',	'Stard3',	'Tcap',	'Erbb2',	'Ormdl3',	'Csf3',	'Med24',	'Thra',	'Nr1d1',	'Wipf2',	'Krt42',	'Eif1',	'Jup',	'Fkbp10',	'Nkiras2',	'Stat5a',	'Stat3',	'Cavin1',	'Naglu',	'Retreg3',	'Rdm1',	'Dhx8',	'Sost',	'Hdac5',	'Slc25a39',	'Dcakd',	'Hexim1',	'Arhgap27',	'Plekhm1',	'Nsf',	'Mettl2',	'Mrc2',	'Cyb561',	'Ace',	'Map3k3',	'Limd2',	'Gm10840',	'Strada',	'Ccdc47',	'Cd79b',	'Ern1',	'Tex2',	'Polg2',	'Smurf2',	'Helz',	'Cacng1',	'Apoh',	'Gna13',	'9930022D16Rik',	'Cdc42ep4',	'Slc9a3r1',	'Nat9',	'Tmem104',	'Cdr2l',	'Mrpl58',	'Mif4gd',	'Caskin2',	'Llgl2',	'Myo15b',	'Smim5',	'Unk',	'Unc13d',	'Mrpl38',	'Srp68',	'Ube2o',	'Rhbdf2',	'Cygb',	'Prcd',	'Mettl23',	'Sec14l1',	'44083',	'6030468B19Rik',	'Tk1',	'Socs3',	'Cant1',	'C1qtnf1',	'Cbx8',	'Cbx4',	'Sgsh',	'Rnf213',	'Endov',	'Chmp6',	'Baiap2',	'Aatk',	'Ndufaf8',	'Slc38a10',	'Bahcc1',	'Actg1',	'Faap100',	'Gcgr',	'Pcyt2',	'Dcxr',	'Ccdc57',	'Csnk1d',	'Metrnl',	'Ptchd3',	'Gdi2',	'Akr1c14',	'Klf6',	'Pfkp',	'Gtpbp4',	'Larp4b',	'Zmynd11',	'Lgals8',	'Gpr137b',	'Lyst',	'Ggps1',	'Cdk13',	'Yae1d1',	'Stard3nl',	'Carmil1',	'BC005537',	'Acot13',	'Tubb2a',	'Psmg4',	'Fam50b',	'Prpf4b',	'Rpp40',	'Ly86',	'Rreb1',	'Ssr1',	'Slc35b3',	'Nedd9',	'Tmem170b',	'Edn1',	'Tbc1d7',	'Gfod1',	'Nol7',	'Ranbp9',	'Mcur1',	'Gmpr',	'Nup153',	'Kif13a',	'Phf2',	'Ninj1',	'Card19',	'Tes3-ps',	'Iars',	'Spin1',	'Sema4d',	'Gadd45g',	'Auh',	'Nfil3',	'Arl10',	'Tspan17',	'Grk6',	'Ddx46',	'Pcbd2',	'H2afy',	'Tifab',	'Klhl3',	'2210016F16Rik',	'Fbp2',	'2010111I01Rik',	'Fancc',	'Lpcat1',	'Slc12a7',	'Erap1',	'Ell2',	'Ttc37',	'Slf1',
                                                'Cetn3',	'Rasa1',	'Ssbp2',	'Gm20379',	'Zfyve16',	'Ap3b1',	'Pde8b',	'F2rl1',	'F2r',	'Col4a3bp',	'Enc1',	'Arhgef28',	'Tnpo1',	'Ptcd2',	'Serf1',	'Gtf2h2',	'Pik3r1',	'Erbin',	'Nln',	'Kif2a',	'Smim15',	'Rab3c',	'Plk2',	'Ankrd55',	'Il6st',	'Plpp1',	'BC067074',	'Arl15',	'Ndufs4',	'Parp8',	'Emb',	'Paip1',	'Hmgcs1',	'Nim1k',	'Rab10',	'Kif3c',	'Dnmt3a',	'Ncoa1',	'Klhl29',	'Apob',	'Ldah',	'Hs1bp3',	'Pum2',	'Laptm4a',	'Ttc32',	'Smc6',	'Lpin1',	'Rock2',	'Pqlc3',	'Hpcal1',	'Cpsf3',	'Iah1',	'Adam17',	'Taf1b',	'Cys1',	'Mboat2',	'Kidins220',	'Rnf144a',	'Rsad2',	'Tpo',	'Ccdc71l',	'Nampt',	'Sypl',	'Atxn7l1',	'Snx13',	'Agmo',	'Arl4a',	'Lsmem1',	'Ifrd1',	'Zfp277',	'Immp2l',	'Dnajb9',	'Stxbp6',	'Sptssa',	'Ppp2r3c',	'Mbip',	'Fkbp3',	'Rpl36al',	'Klhdc2',	'Gm9887',	'Vcpkmt',	'Sos2',	'Abhd12b',	'Pygl',	'Tmx1',	'Frmd6',	'Actr10',	'Arid4a',	'Tmem30b',	'Gphb5',	'Ppp2r5e',	'Syne2',	'Mthfd1',	'Zbtb25',	'Max',	'Fut8',	'Gphn',	'Fam71d',	'Plekhh1',	'Arg2',	'Rad51b',	'Zfp36l1',	'Actn1',	'Ttc9',	'Pcnx',	'Psen1',	'Pnma1',	'Elmsan1',	'Rnf113a2',	'Aldh6a1',	'Zc2hc1c',	'Tmed10',	'Fos',	'Batf',	'Flvcr2',	'Erg28',	'Cipc',	'Adck1',	'Gtf2a1',	'Ston2',	'Ptpn21',	'Ttc7b',	'Ccdc88c',	'Fbln5',	'Rin3',	'Lgmn',	'Chga',	'Itpk1',	'Ifi27',	'Ppp4r4',	'Bdkrb2',	'Gskip',	'Ccdc85c',	'Degs2',	'Slc25a29',	'Slc25a47',	'Wdr25',	'Ppp2r5c',	'Hsp90aa1',	'Zfp839',	'Cinp',	'Tecpr2',	'Traf3',	'Mark3',	'Trmt61a',	'Bag5',	'Ppp1r13b',	'Inf2',	'Adssl1',	'Siva1',	'Zbtb42',	'Cdca4',	'Tex22',	'Mta1',	'Crip2',	'Crip1',	'Ccdc152',	'Fbxo4',	'Oxct1',	'Rictor',	'Egflam',	'Gdnf',	'Slc1a3',	'Dnajc21',	'Rai14',	'Zfr',	'1810049J17Rik',	'Myo10',	'Retreg1',	'Zfp622',	'Fbxl7',	'Otulin',	'Trio',	'Dap',	'Osr2',	'Vps13b',	'Cox6c',	'Spag1',	'Ywhaz',	'Zfp706',	'Rrm2b',	'Klf10',	'Azin1',	'Atp6v1c1',	'Abra',	'Eif3e',	'Eny2',	'Col14a1',	'Sntb1',	'Zhx2',	'Derl1',	'9130401M01Rik',	'Fam91a1',	'Rnf139',	'Washc5',	'Trib1',	'Myc',	'Fam49b',	'Asap1',	'Efr3a',	'Tg',	'Zfat',	'Ago2',	'Dennd3',	'Slc45a4',	'Gpr20',	'Zc3h3',	'Tigd5',	'Pycrl',	'Scrib',	'Puf60',	'Nrbp2',	'Eppk1',	'BC024139',	'Plec',	'Parp10',	'Smpd5',	'Hgh1',	'Bop1',	'Scx',	'Fbxl6',	'Kifc2',	'Ppp1r16a',	'Zfp251',	'1110038F14Rik',	'Mb',	'Apol6',	'Apol8',	'Myh9',	'Txn2',	'Sstr3',	'Mfng',	'Triobp',	'Micall1',	'Polr2f',	'Pla2g6',	'Maff',	'Tmem184b',	'Kdelr3',	'Sun2',	'Dnal4',	'Apobec3',	'Pdgfb',	'Tnrc6b',	'Sgsm3',	'Chadl',	'Tef',	'Polr3h',	'Desi1',	'Naga',	'Ndufa6',	'Cyp2d22',	'Pacsin2',	'Ttll1',	'Bik',	'Mcat',	'Ttll12',	'Rtl6',	'5031439G07Rik',	'Atxn10',	
                                                'Wnt7b',	'Ppara',	'Tbc1d22a',	'Alg12',	'Creld2',	'Pim3',	'Plxnb2',	'C730034F03Rik',	'Mapk8ip2',	'Gxylt1',	'Pus7l',	'Twf1',	'Ano6',	'Slc38a4',	'Endou',	'Rapgef3',	'Slc48a1',	'Hdac7',	'Asb8',	'Cacnb3',	'Ccdc65',	'Ddn',	'Kmt2d',	'Tuba1c',	'Tmbim6',	'Nckap5l',	'Faim2',	'Aqp2',	'Aqp5',	'Racgap1',
                                                'Smarcd1',	'Gpd1',	'Atf1',	'Mettl7a1',	'Gm5475',	'Fignl2',	'Nr4a1',	'Krt80',	'Krt8',	'Tns2',	'Soat2',	'Zfp740',	'Sp1',	'Npff',	'Atp5g2',	'Itga5',	'Gtsf1',	'Adcy9',	'Tfap4',	'Dnaja3',	'Nmral1',	'Hmox2',	'Cdip1',	'4930562C15Rik',	'Ubald1',	'Mgrn1',	'Nudt16l1',	'AU021092',	'Gm5767',	'1810013L24Rik',	'Emp2',	'Clec16a',	'Socs1',	'Litaf',	'Zc3h7a',	'Rsl1d1',	'Cpped1',	'Mkl2',	'Parn',	'Pla2g10',	'Pdxdc1',	'Nde1',	'Cebpd',	'Mapk1',	'Ppil2',	'Scarf2',	'Dgcr6',	'Prodh',	'Txnrd2',	'Hira',	'Klhl6',	'Camk2n2',	'Eif4g1',	'Ephb3',	'Ehhadh',	'Tmem41a',	'Liph',	'Igf2bp2',	'Atp13a4',	'Opa1',	'Hes1',	'Tmem44',	'Xxylt1',	'Pak2',	'Pcyt1a',	'Tnk2',	'Fyttd1',	'Osbpl11',	'Snx4',	'Slc12a8',	'Heg1',	'Muc13',	'Kalrn',	'Adcy5',	'Sec22a',	'Iqcb1',	'Lrrc58',	'Gsk3b',	'Zbtb20',	'Qtrt2',	'Atp6v1a',	'Usf3',	'Spice1',	'Cfap44',	'Slc35a5',	'Abhd10',	'Phldb2',	'Cd47',	'Cblb',	'Alcam',	'Cep97',	'Zbtb11',	'Tbc1d23',	'Olfr172',	'Riox2',	'4930453N24Rik',	'Cggbp1',	'Chmp2b',	'Usp25',	'Btg3',	'App',	'Adamts1',	'Map3k7cl',	'Bach1',	'Tiam1',	'Sod1',	'Eva1c',	'Synj1',	'Paxbp1',	'Ifnar2',	'Ifnar1',	'Ifngr2',	'Tmem50b',	'Dnajc28',	'Itsn1',	'Kcne2',	'Clic6',	'Cbr3',	'Kcnj15',	'Ets2',	'Brwd1',	'Tmprss2',	'Ripk4',	'Arid1b',	'Ldhal6b',	'Zdhhc14',	'Serac1',	'Sytl3',	'Ezr',	'AC122413.2',	'Sft2d1',	'Prr18',	'Qk',	'Agpat4',	'Igf2r',	'Acat3',	'Wtap',	'Afdn',	'Smoc2',	'Gm3448',	'Chd1',	'Rgmb',	'Lnpep',	'Zfp948',	'Zfp943',	'Pdpk1',	'Amdhd2',	'Tbc1d24',	'Caskin1',	'Nthl1',	'Npw',	'Eme2',	'Tekt4',	'Rab40c',	'Capn15',	'Tmem8',	'Dusp1',	'Atp6v0e',	'Crebrf',	'Bnip1',	'Lemd2',	'AI413582',	'Taf11',	'Zfp523',	'Ppard',	'Tulp1',	'Fkbp5',	'Mapk14',	'Kctd20',	'Stk38',	'Srsf3',	'Cdkn1a',	'BC004004',	'Rnf8',	'Gm28043',	'Zfand3',	'Btbd9',	'Slc37a1',	'Wdr4',	'Cryaa',	'Sik1',	'Rrp1b',	'Notch3',	'Cyp4f13',	'Adamts10',	'Rab11b',	'Kank3',	'Ndufa7',	'H2-K1',	'Tap1',	'Gm15821',	'H2-Ab1',	'Notch4',	'Pbx2',	'Atf6b',	'Neu1',	'H2-D1',	'H2-Q2',	'Ddr1',	'Ier3',	'Atat1',	'Ppp1r10',	'Trim15',	'Rnf39',	'Adgrf2',	'Cyp39a1',	'Rcan2',	'Enpp4',	'Clic5',	'Tmem63b',	'1600014C23Rik',	'Vegfa',	'Gtpbp2',	'Polr1c',	'Zfp318',	'Cul9',	'Ptk7',	'Gnmt',	'1700001C19Rik',	'Ccnd3',	'Tomm6',	'Pgc',	'Daam2',	'Kif6',	'Tbc1d5',	'Zfp119b',	'Stap2',	'Sema6b',	'Dpp9',	'Nrtn',	'Prr22',	'Rfx2',	'1700061G19Rik',	'Mllt1',	'Vav1',	'Efna5',	'Fbxl17',	'A930002H24Rik',	'Pja2',	'Ppp4r1',	'Ralbp1',	'Twsg1',	'Ankrd12',	'Washc1',	'Mtcl1',	'Ptprm',	'Arhgap28',	'Myl12a',	'Lpin2',	'Emilin2',	'Clip4',	'Ypel5',	'Xdh',	'Ttc27',	'Fam98a',
                                                'Strn',	'Prkd3',	'Atl2',	'Hnrnpll',	'Sos1',	'Cox7a2l',	'Mta3',	'Haao',	'Zfp36l2',	'Thada',	'Dync2li1',	'Lrpprc',	'Slc3a1',	'Srbd1',	'Prkce',	'Gm10309',	'Ttc7',	'Calm2',	'Foxn2',	'Ston1',	'Cul2',	'Svil',	'Zfp438',	'Arhgap12',	'Kif5b',	'Epc1',	'Mpp7',	'Wac',	'Usp14',	'Rock1',	'Mib1',	'Gata6',	'Cables1',	'Npc1',	'Osbpl1a',	'Ss18',	'Kctd1',	'Rnf138',	'Garem1',	'Mapre2',	'Rprd1a',	'Sft2d3',	'Nrep',	'Reep5',	'Cdc23',	'Egr1',	'Ctnna1',	'Paip2',	'Ecscr',	'Ube2d2a',	'Cxxc5',	'Hbegf',	'Pcdhga2',	'Pcdhgb5',	'Pcdhga10',	'Pcdh1',	'Rnf14',	'Fgf1',	'Nr3c1',	'Pabpc2',	'Trim36',	'Snx2',	'Snx24',	'Csnk1g3',	'Zfp608',	'Tex43',	'Slc12a2',	'Iigp1',	'Synpo',	'Ndst1',	'Rps14',	'Cd74',	'Csf1r',	'Hmgxb3',	'Pde6a',	'Arhgef37',	'Pcyox1l',	'Grpel2',	'Ablim3',	'Adrb2',	'Fbxo38',	'Fech',	'Nars',	'Nedd4l',	'Alpk2',	'Zfp532',	'Sec11c',	'Cidea',	'Afg3l2',	'Psmg2',	'Ldlrad4',	'Fam210a',	'Mbd2',	'Mex3c',	'Myo5b',	'Acaa2',	'Smad7',	'Ctif',	'Zbtb7c',	'Smad2',	'Hdhd2',	'Setbp1',	'Adnp2',	'Pqlc1',	'Ctdp1',	'Nfatc1',	'Mbp',	'Zfp236',	'Zfp516',	'Tshz1',	'Cndp2',	'Cyb5a',	'Tmx3',	'Cpt1a',	'Tesmin',	'Ppp6r3',	'Lrp5',	'Chka',	'Unc93b1',	'Cdk2ap2',	'Coro1b',	'Pold4',	'Grk2',	'Rhod',	'2010003K11Rik',	'Pcx',	'Lrfn4',	'Rce1',	'Rin1',	'Cd248',	'Mus81',	'Snx32',	'Ap5b1',	'Kcnk7',	'Ehbp1l1',	'Sssca1',	'Ltbp3',	'Scyl1',	'Frmd8',	'Majin',	'Cdc42bpg',	'Sf1',	'Slc22a12',	'Ccdc88b',	'Gpr137',	'Cox8a',	'Rcor2',	'Slc3a2',	'Lrrn4cl',	'B3gat3',	'Rom1',	'Scgb1a1',	'Fth1',	'Best1',	'Fads2',	'Myrf',	'Cpsf7',	'Vps37c',	'Ms4a8a',	'Stx3',	'Mpeg1',	'Carnmt1',	'D030056L22Rik',	'Trpm6',	'Rorb',	'Anxa1',	'Klf9',	'Fam189a2',	'Fxn',	'Pip5k1b',	'Fam122a',	'Dock8',	'Dmrt1',	'Smarca2',	'Gm815',	'Vldlr',	'Pum3',	'Glis3',	'Rcl1',	'Jak2',	'Plgrkt',	'Ric1',	'Mlana',	'Uhrf2',	'Asah2',	'Sgms1',	'Pten',	'Pcgf5',	'43895',	'Exoc6',	'Myof',	'Pdlim1',	'Sorbs1',	'Cc2d2b',	'Ccnj',	'Tm9sf3',	'Lcor',	'Rrp12',	'Exosc1',	'Ankrd2',	'Pi4k2a',	'Sfrp5',	'R3hcc1l',	'Pyroxd2',	'Got1',	'Erlin1',	'Bloc1s2',	'Scd2',	'Scd1',	'Lzts2',	'Gm17018',	'Fbxw4',	'Mgea5',	'Hps6',	'Ldb1',	'Sufu',	'Trim8',	'Wbp1l',	'Cyp17a1',	'Cnnm2',	'Ina',	'Calhm2',	'Neurl1a',	'Sh3pxd2a',	'Sfr1',	'Rbm20',	'Vti1a',	'Casp7',	'Nhlrc2',	'Adrb1',	'Ccdc186',	'Vwa2',	'Afap1l2',	'Ablim1',	'Fam160b1',	'Trub1',	'Atrnl1',	'Shtn1',	'Pdzd8',	'Rab11fip2',	'Cacul1',	'Zfp950'))

integrate_epi4 <- AddModuleScore(object = integrate_epi4, features = Both_NKX21_binding_acquired_over_time, ctrl = 500, name = "NKXBAcquiredGenes", search = TRUE, random.seed = 1)
#6 genes not accounted for 43891, 43892, 43898, 44084, 44083, 43895
FeaturePlot(object = integrate_epi4, features = 'NKXBAcquiredGenes1', cols = c('lightgrey','red'))

Both_NKX21_binding_retained_from_progenitor <- list(c('Cops5',	'Arfgef1',	'Ncoa2',	'Tram1',	'Lactb2',	'Eya1',	'Msc',	'Terf1',	'Ly96',	'B3gat2',	'Col9a1',	'Ptp4a1',	'Ccdc115',	'Cnnm3',	'Fam178b',	'Inpp4a',	'2010300C02Rik',	'Lipt1',	'Lyg2',	'Rev1',	'Aff3',	'Pdcl3',	'Map4k4',	'Il1r2',	'Mrps9',	'AI597479',	'Nck2',	'Uxs1',	'Dnah7b',	'Nabp1',	'Stat4',	'Hibch',	'Stk17b',	'Pgap1',	'Plcl1',	'Satb2',	'Fzd7',	'Gm973',	'Bmpr2',	'Fam117b',	'Nrp2',	'Fzd5',	'Idh1',	'Lancl1',	'Spag16',	'Mreg',	'Pecr',	'Xrcc5',	'Rpl37a',	'Igfbp5',	'Tns1',	'Tmbim1',	'Catip',	'Ctdsp1',	'Cnot9',	'Zfp142',	'Wnt10a',	'Ihh',	'Epha4',	'Acsl3',	'Ap1s3',	'Irs1',	'Rhbdd1',	'Col4a4',	'Col4a3',	'Daw1',	'A630001G21Rik',	'Armc9',	'Ptma',	'Spp2',	'Glrp1',	'Agap1',	'Gbx2',	'Ackr3',	'Col6a3',	'Ube2f',	'Asb1',	'Twist2',	'Hdac4',	'Gpc1',	'Crocc2',	'Stk25',	'Bok',	'Pdcd1',	'Slco4c1',	'Pam',	'Phlpp1',	'Gli2',	'Steap3',	'Actr3',	'Slc35f5',	'Nckap5',	'Ccnt2',	'Mcm6',	'Cd55b',	'Zp3r',	'Fcamr',	'Pigr',	'Mapkapk2',	'Dyrk3',	'Rassf5',	'Ikbke',	'Slc26a9',	'Slc45a3',	'Cdk18',	'Nuak2',	'Lax1',	'Optc',	'Fmod',	'Btg2',	'Cyb5r1',	'Syt2',	'Ube2t',	'Elf3',	'Lmod1',	'Ipo9',	'Csrp1',	'Tmem9',	'Cacna1s',	'Kif14',	'Nr5a2',	'Ptprc',	'Dennd1b',	'Trove2',	'Hmcn1',	'Ivns1abp',	'Edem3',	'Arpc5',	'Ncf2',	'Smg7',	'Lamc2',	'Shcbp1l',	'Dhx9',	'Npl',	'Rnasel',	'Cacna1e',	'Ier5',	'BC034090',	'Qsox1',	'Tor1aip1',	'Abl2',	'Tor3a',	'Fam20b',	'Ralgps2',	'Pappa2',	'Cacybp',	'4930558K02Rik',	'Mettl11b',	'Xcl1',	'Uck2',	'Aldh9a1',	'Lrrc52',	'Pbx1',	'Fcgr4',	'Fcgr3',	'Cfap126',	'Tomm40l',	'Alyref2',	'Cfap45',	'Cadm3',	'Fmn2',	'Grem2',	'Pld5',	'Rbm8a2',	'Akt3',	'1700016C15Rik',	'Catspere2',	'Kif26b',	'Smyd3',	'Cnst',	'Nvl',	'Lbr',	'Srp9',	'Tlr5',	'43891',	'Bpnt1',	'Lyplal1',	'Tgfb2',	'Kcnk2',	'Cenpf',	'Ptpn14',	'Smyd2',	'Prox1',	'Rps6kc1',	'Nenf',	'Lpgat1',	'Sertad4',	'G0s2',	'Camk1g',	'Cd46',	'Cr1l',	'Nmt2',	'Acbd7',	'Bend7',	'Mcm10',	'Camk1d',	'Proser2',	'Usp6nl',	'Gata3',	'Taf3',	'Itih5',	'Sfmbt2',	'Trdmt1',	'Commd3',	'Arhgap21',	'Pdss1',	'Abi1',	'Psd4',	'Dph7',	'Nrarp',	'Ccdc187',	'Sec16a',	'Vav2',	'Wdr5',	'Rxra',	'Fcnb',	'Mrps2',	'Spaca9',	'Barhl1',	'Cfap77',	'Med27',	'Rapgef1',	'Sptan1',	'Set',	'Ptpa',	'Ier5l',	'Cstad',	'1700001O22Rik',	'Ncs1',	'Ass1',	'Exosc2',	'Qrfp',	'Slc25a25',	'Fam102a',	'Dpm2',	'Ak1',	'Fam129b',	'Garnl3',	'Angptl2',	'Zbtb34',	'Mvb12b',	'Rab14',	'Stom',	'Dab2ip',	'Ttll11',	'Ndufa8',	'Rabgap1',	'Crb2',	'Nr6a1',	'Arhgap15',	'Acvr2a',	'Mbd5',	'Epc2',	'Rnd3',	'Arl5a',	'Stam2',	'Fmnl2',	'Prpf40a',	'Rprm',	'Cytip',	'Pkp4',	'Tanc1',	'Wdsub1',	'Itgb6',
                                                      'Rbms1',	'Dpp4',	'Ifih1',	'Kcnh7',	'Fign',	'Grb14',	'Slc38a11',	'Scn3a',	'Scn2a',	'Stk39',	'Myo3b',	'Gad1',	'Cybrd1',	'Metap1d',	'Dlx2',	'Rapgef4',	'Sp9',	'Gpr155',	'Wipf1',	'Chrna1',	'Lnpk',	'Nfe2l2',	'Agps',	'Ttc30a1',	'Rbm45',	'Prkra',	'Cwc22',	'Ube2e3',	'Prdx6b',	'Zfp804a',	'Tfpi',	'Ctnnd1',	'Zdhhc5',	'Gm19426',	'Prg3',	'Tnks1bp1',	'Aplnr',	'Olfr1274-ps',	'Ptprj',	'Madd',	'Arfgap2',	'1110051M20Rik',	'Lrp4',	'Phf21a',	'1700029I15Rik',	'Cry2',	'Cd82',	'Accs',	'Api5',	'B230118H07Rik',	'Traf6',	'Prr5l',	'Commd9',	'Abtb2',	'Nat10',	'Caprin1',	'A930018P22Rik',	'Qser1',	'Them7',	'Rcn1',	'Pax6',	'Mpped2',	'Arl14ep',	'Kif18a',	'Lin7c',	'Arhgap11a',	'BC052040',	'Rasgrp1',	'Thbs1',	'Fsip1',	'Bmf',	'Bub1b',	'Chst14',	'Ccdc32',	'Rpusd2',	'Knl1',	'Rad51',	'Dnajc17',	'Chp1',	'Oip5',	'Itpka',	'Rpap1',	'Ehd4',	'Capn3',	'Zfp106',	'Ubr1',	'Zscan29',	'Pdia3',	'Mfap1b',	'Mfap1a',	'Wdr76',	'Shf',	'AA467197',	'Gabpb1',	'Trpm7',	'1810024B03Rik',	'Kcnip3',	'Mall',	'Bcl2l11',	'Anapc1',	'Mertk',	'Tmem87b',	'Zc3h8',	'Gm14025',	'Ptpra',	'Cenpb',	'Pank2',	'Pcna',	'Chgb',	'Bmp2',	'Plcb4',	'Lamp5',	'Btbd3',	'Macrod2',	'Kif16b',	'Rbbp9',	'Dtd1',	'Insm1',	'Nkx2-2',	'Foxa2',	'Nxt1',	'Zfp442',	'Acss1',	'Rspo4',	'Csnk2a1',	'Sox12',	'Zcchc3',	'Mcts2',	'Id1',	'Tpx2',	'Plagl2',	'Commd7',	'Dnmt3b',	'Snta1',	'Cbfa2t2',	'1700003F12Rik',	'E2f1',	'Gm14226',	'Ahcy',	'Itch',	'Pigu',	'Edem2',	'Epb41l1',	'Myl9',	'Soga1',	'Rpn2',	'Blcap',	'Vstm2l',	'Tgm2',	'Lbp',	'Snhg11',	'Plcg1',	'Emilin3',	'Chd6',	'Mybl2',	'Jph2',	'Tomm34',	'Sys1',	'Wfdc2',	'Ncoa5',	'Zfp663',	'Zmynd8',	'Ncoa3',	'Sulf2',	'Trp53rkb',	'Znfx1',	'Slc9a8',	'Snai1',	'Tmem189',	'Ptpn1',	'Pard6b',	'Nfatc2',	'Zfp217',	'Bcas1',	'Rae1',	'Ankrd60',	'Prelid3b',	'Zfp831',	'Edn3',	'Zfp931',	'Cdh26',	'Taf4',	'Lama5',	'Rps21',	'Rbbp8nl',	'Slco4a1',	'Mrgbp',	'Fndc11',	'Sox18',	'Shroom4',	'Usp27x',	'Bcor',	'Usp9x',	'Gpr82',	'Maoa',	'Jade3',	'Uba1',	'Slc25a43',	'Ube2a',	'Nkap',	'Ocrl',	'Bcorl1',	'Enox2',	'Mbnl3',	'Gpc4',	'Gpc3',	'Hprt',	'Plac1',	'Zfp449',	'Slc9a6',	'1700020N15Rik',	'Cd99l2',	'Dusp9',	'Flna',	'Rpl10',	'Tbl1x',	'Arx',	'Pcyt1b',	'Maged1',	'Yipf6',	'Stard8',	'Kif4',	'Gdpd2',	'Slc7a3',	'Snx12',	'Ogt',	'Rtl5',	'Pin4',	'Ercc6l',	'Hdac8',	'Tspan6',	'Cstf2',	'Cenpi',	'Rpl36a',	'Armcx1',	'Armcx5',	'Bhlhb9',	'Bex4',	'Tceal8',	'Tceal9',	'Tceal3',	'Rnf128',	'Tsc22d3',	'Kcne1l',	'Pak3',	'Capn6',	'Rps6ka3',	'Tceanc',	'Egfl6',	'Tmsb4x',	'Arhgap6',	'Gm21887',	'Gm8797',	'Mrps28',	'Zfp704',	'Pag1',	'Ythdf3',	'Pde7a',	'Dnajc5b',
                                                      'Tnfsf10',	'Fndc3b',	'Pld1',	'Mecom',	'Mynn',	'Prkci',	'Skil',	'Actl6a',	'Ccdc39',	'Fxr1',	'Sox2',	'Atp11b',	'Dcun1d1',	'D3Ertd254e',	'1810062G17Rik',	'Slc7a11',	'Rab33b',	'Mgst2',	'Maml3',	'Cog6',	'Stoml3',	'Frem2',	'Smad9',	'Commd2',	'Pfn2',	'Tsc22d2',	'Mbnl1',	'Rap2b',	'Plch1',	'Shox2',	'Schip1',	'Ift80',	'Arl14',	'Nmd3',	'Rapgef2',	'Etfdh',	'Tmem144',	'Fam198b',	'Pdgfc',	'Ctso',	'Gucy1b1',	'Tlr2',	'Tmem131l',	'Trim2',	'Gm6525',	'Fhdc1',	'Fbxw7',	'Gatb',	'Glt28d2',	'Mab21l2',	'Kirrel',	'Etv3',	'Prcc',	'Hdgf',	'Rrnad1',	'Nes',	'Mef2d',	'Tsacc',	'Tmem79',	'Arhgef2',	'Syt11',	'Clk2',	'Thbs3',	'Trim46',	'Krtcap2',	'Efna4',	'Flad1',	'Kcnn3',	'Hax1',	'Tpm3',	'Crtc2',	'Ints3',	'Ilf2',	'Rptn',	'Snx27',	'Rfx5',	'Psmd4',	'Vps72',	'Scnm1',	'Sema6c',	'Cdc42se1',	'Cers2',	'Setdb1',	'Hormad1',	'Mcl1',	'Tars2',	'Rprd2',	'Bola1',	'Hist2h2aa2',	'Hist2h4',	'Fcgr1',	'Hfe2',	'Bcl9',	'Notch2',	'Hmgcs2',	'Ttf2',	'Cd2',	'Igsf3',	'Vangl1',	'Sycp1',	'Nras',	'Trim33',	'Rsbn1',	'Magi3',	'Lrig2',	'Wnt2b',	'Ddx20',	'Adora3',	'Kcnc4',	'Csf1',	'Gstm7',	'Atxn7l2',	'Psrc1',	'5330417C22Rik',	'Taf13',	'Cdc14a',	'Rtca',	'Dbt',	'Agl',	'Alg14',	'Cnn3',	'Slc44a3',	'F3',	'Arhgap29',	'Abca4',	'Fnbp1l',	'Synpo2',	'Camk2d',	'Elovl6',	'Egf',	'Ostc',	'Cyp2u1',	'Papss1',	'Aimp1',	'Cenpe',	'Bdh2',	'Nfkb1',	'Slc39a8',	'Ppp3ca',	'Gm21962',	'Bmpr1b',	'Pdlim5',	'Gtf2b',	'Lmo4',	'Hs2st1',	'Sh3glb1',	'Znhit6',	'Cyr61',	'Ssx2ip',	'Ttll7',	'Adgrl2',	'Dnajb4',	'Miga1',	'St6galnac3',	'Erich3',	'Wls',	'Gm11808',	'Penk',	'Fam110b',	'Nsmaf',	'Car8',	'Rab2a',	'Chd7',	'2610301B20Rik',	'Pdp1',	'Rbm12b2',	'Tmem64',	'Manea',	'Bach2',	'Pnrc1',	'Akirin2',	'Aco1',	'B4galt1',	'Ubap2',	'Ubap1',	'Enho',	'Fam166b',	'Ccdc107',	'Trmt10b',	'Shb',	'Ncbp1',	'Anks6',	'Stx17',	'Tex10',	'Rnf20',	'Fsd1l',	'Rad23b',	'Ptpn3',	'D630039A03Rik',	'Inip',	'Slc46a2',	'Zfp37',	'Cdc26',	'Wdr31',	'Bspry',	'4933430I17Rik',	'Rgs3',	'Kif12',	'Megf9',	'Tle1',	'Lurap1l',	'Mpdz',	'Nfib',	'Cer1',	'Bnc2',	'Adamtsl1',	'Hacd4',	'Klhl9',	'Mtap',	'Jun',	'Fggy',	'E130114P18Rik',	'Nfia',	'Dock7',	'Pgm2',	'Ror1',	'Ube2u',	'Cachd1',	'Raver2',	'Ak4',	'Sgip1',	'Tctex1d1',	'Plpp3',	'Fam151a',	'Acot11',	'Ssbp3',	'Lrrc42',	'Elavl4',	'Spata6',	'Cmpk1',	'Stil',	'Pdzk1ip1',	'Tex38',	'Nsun4',	'Uqcrh',	'Rad54l',	'Tmem69',	'Nasp',	'Akr1a1',	'Kif2c',	'Tmem53',	'Ptprf',	'Cdc20',	'Tmem125',	'Cfap57',	'Slc2a1',	'Zfp691',	'Cldn19',	'Ccdc30',	'Scmh1',	'Mycl',	'Pou3f1',	'9930104L06Rik',	'Dnali1',	'Snip1',	'Ago1',	'Clspn',	'5730409E04Rik',
                                                      'Zmym4',	'Sfpq',	'Gja4',	'Gjb3',	'Zfp362',	'Ak2',	'Tmem54',	'Zbtb8os',	'Kpna6',	'Khdrbs1',	'Tinagl1',	'Gm10300',	'Taf12',	'Rab42',	'Med18',	'Sesn2',	'Smpdl3b',	'Rpa2',	'Ppp1r8',	'Ahdc1',	'Wasf2',	'Cd164l2',	'Tmem222',	'Kdf1',	'Sfn',	'Arid1a',	'Hmgn2',	'Lin28a',	'Cd52',	'Cep85',	'Pdik1l',	'Pafah2',	'Stmn1',	'Man1c1',	'Ldlrap1',	'Rhd',	'Nipal3',	'Fuca1',	'E2f2',	'Asap3',	'Hnrnpr',	'Luzp1',	'Lactbl1',	'C1qa',	'Zbtb40',	'Wnt4',	'Hspg2',	'Alpl',	'Eif4g3',	'Pla2g2d',	'Pla2g5',	'Nbl1',	'Capzb',	'Pax7',	'Arhgef10l',	'Rcc2',	'Padi6',	'Sdhb',	'Szrd1',	'Fbxo42',	'Rsg1',	'Arhgef19',	'Clcnka',	'Casp9',	'Efhd2',	'Tmem51',	'Kazn',	'Casz1',	'Cenps',	'Rbp7',	'Gpr157',	'Eno1',	'Errfi1',	'Vamp3',	'Tnfrsf25',	'Acot7',	'Prdm16',	'B230104I21Rik',	'Ttc34',	'Mmel1',	'Plch2',	'Morn1',	'Ski',	'Faap20',	'Prkcz',	'C1qtnf12',	'Ttll10',	'9430015G10Rik',	'Cdk6',	'Cyp51',	'Cdk14',	'Sema3e',	'Pclo',	'Sema3c',	'Gnai1',	'Ptpn12',	'Fgl2',	'Srpk2',	'Kcnh2',	'Chpf2',	'Rheb',	'Prkag2',	'Actr3b',	'Shh',	'Rnf32',	'Dpysl5',	'Agbl5',	'Preb',	'Cad',	'Ift172',	'Ywhah',	'Spon2',	'Uvssa',	'Slbp',	'Mxd4',	'Ablim2',	'Sorcs2',	'Grpel1',	'Tbc1d14',	'D5Ertd579e',	'Rab28',	'Cpeb2',	'Bst1',	'Fgfbp1',	'Tapt1',	'Slit2',	'Adgra3',	'Anapc4',	'Slc34a2',	'Smim20',	'Cckar',	'Tbc1d19',	'Stim2',	'0610040J01Rik',	'Klf3',	'Klhl5',	'Ugdh',	'Smim14',	'N4bp2',	'Chrna9',	'9130230L23Rik',	'Apbb2',	'Limch1',	'Phox2b',	'Atp8a1',	'Gm5108',	'Nfxl1',	'Fryl',	'Lrrc66',	'Spata18',	'Usp46',	'Rasl11b',	'Lnx1',	'Tmem165',	'Aasdh',	'Spink2',	'Mob1b',	'Slc4a4',	'Gc',	'Npffr2',	'Cox18',	'Afp',	'Rassf6',	'Mthfd2l',	'Epgn',	'Areg',	'Stbd1',	'Shroom3',	'Sowahb',	'2010109A12Rik',	'Fras1',	'4930524J08Rik',	'Hnrnpdl',	'Enoph1',	'Gpat3',	'Cds1',	'Ptpn13',	'Aff1',	'Klhl8',	'Lrrc8c',	'Zfp644',	'Cdc7',	'Tgfbr3',	'Gfi1',	'Ube2d2b',	'Mtf2',	'Pcgf3',	'Slc26a1',	'Crlf2',	'Gm15446',	'Zfp932',	'Plcxd1',	'Hscb',	'Pitpnb',	'Sez6l',	'Myo18b',	'2900026A02Rik',	'Sart3',	'Ssh1',	'Mmab',	'Fam222a',	'Gm10401',	'Dynll1',	'Pla2g1b',	'Sirt4',	'Pxn',	'Rplp0',	'Vsig10',	'Wsb2',	'Rfc5',	'Tesc',	'Rnft2',	'2410131K14Rik',	'Med13l',	'Tmem116',	'Acad12',	'Cux2',	'Kdm2b',	'Orai1',	'B3gnt4',	'Kntc1',	'Abcb9',	'Arl6ip4',	'Pitpnm2',	'Cdk2ap1',	'Sbno1',	'Tmed2',	'Rflna',	'Ncor2',	'Ubc',	'Tmem132c',	'Galnt17',	'Auts2',	'Rcc1l',	'Gtf2i',	'Rfc2',	'Eln',	'Cldn3',	'Abhd11',	'Mlxipl',	'Baz1b',	'Rhbdd2',	'Ssc4d',	'Dtx2',	'Rasa4',	'Polr2j',	'Sh2b2',	'Cux1',	'Myl10',	'Col26a1',	'Fis1',	'Plod3',	'Ap1s1',	'Trim56',	'Ufsp1',	'Ephb4',	'Tsc22d4',	'Gm5294',	
                                                      'Elfn1',	'Mrm2',	'Eif3b',	'Lfng',	'Gna12',	'Foxk1',	'Slc29a4',	'Actb',	'Fscn1',	'Fam220a',	'Bri3',	'Nptx2',	'Usp12',	'Pomp',	'Mtus2',	'Alox5ap',	'Medag',	'Tex26',	'Wdr95',	'B3glct',	'N4bp2l2',	'Kl',	'Stard13',	'Ppp1r9a',	'Asb4',	'Dync1i1',	'Dlx5',	'Asns',	'Mios',	'Glcci1',	'Ica1',	'Vwde',	'2610001J05Rik',	'Ppp1r3a',	'Foxp2',	'Mdfic',	'Tes',	'St7',	'Wnt2',	'Asz1',	'Fam3c',	'Atp6v1f',	'Tnpo3',	'Tspan33',	'Nrf1',	'Zc3hc1',	'Klhdc10',	'Cpa4',	'Podxl',	'Exoc4',	'Agbl3',	'Wdr91',	'Dgki',	'Creb3l2',	'Svopl',	'Atp6v0a4',	'Tmem213',	'Ttc26',	'Hipk2',	'Tbxas1',	'Kdm7a',	'Rab19',	'Arhgef5',	'Ezh2',	'Pdia4',	'Igf2bp3',	'Mpp6',	'Gsdme',	'Nfe2l3',	'Snx10',	'Skap2',	'Creb5',	'Tril',	'Chn2',	'Wipf3',	'Mindy4',	'Ghrhr',	'Nap1l5',	'Fam13a',	'Tnip3',	'Ndnf',	'Il12rb2',	'Rpia',	'Mrpl35',	'Ptcd3',	'St3gal5',	'Atoh8',	'Usp39',	'Tmem150a',	'Ggcx',	'Capg',	'Tgoln1',	'Tcf7l1',	'Kcmf1',	'Dnah6',	'Eva1a',	'Sema4f',	'Wbp1',	'Slc4a5',	'Tet3',	'Zfp638',	'Cyp26b1',	'Noto',	'Nat8f4',	'Fam136a',	'Snrpg',	'Tia1',	'Anxa4',	'Nfu1',	'Gfpt1',	'Antxr1',	'Gkn3',	'Mgll',	'Tpra1',	'Plxna1',	'Iqsec1',	'Slc6a6',	'4930590J08Rik',	'Mrps25',	'Adamts9',	'Gm15737',	'Magi1',	'Slc25a26',	'Lrig1',	'Fam19a4',	'Lmod3',	'Foxp1',	'Itpr1',	'Bhlhe40',	'Lmcd1',	'Rad18',	'Sec13',	'Atg7',	'Vgll4',	'Tsen2',	'Plxnd1',	'Washc2',	'Zfp422',	'Hnrnpf',	'Csgalnact2',	'Ret',	'Zfp9',	'Dcp1b',	'Fbxl14',	'Ninj2',	'B4galnt3',	'D330020A13Rik',	'Hdhd5',	'Cecr2',	'Pex26',	'Phc1',	'Rimklb',	'Lpcat3',	'Emg1',	'Phb2',	'Grcc10',	'Usp5',	'Ing4',	'Tnfrsf1a',	'Vwf',	'Ntf3',	'Ccnd2',	'Parp11',	'Klrb1c',	'Klrb1b',	'Ybx3',	'Etv6',	'Mansc1',	'Dusp16',	'Ddx47',	'Gprc5a',	'Gsg1',	'Pbp2',	'Emp1',	'Grin2b',	'Gm8994',	'Atf7ip',	'Plbd1',	'Hist4h4',	'Eps8',	'Mgst1',	'Pik3c2g',	'Plekha5',	'Pde3a',	'Slco1c1',	'Cmas',	'Etnk1',	'Sox5',	'Arntl2',	'Klhl42',	'Pthlh',	'Caprin2',	'Oscar',	'Rdh13',	'Eps8l1',	'Hspbp1',	'Kmt5c',	'Zfp444',	'Zfp667',	'Zfp418',	'Rnf225',	'Zfp324',	'Zbtb45',	'Chmp2a',	'Bicra',	'Slc8a2',	'Prkd2',	'Calm3',	'Mypop',	'Irf2bp1',	'Six5',	'Fosb',	'Exoc3l2',	'Ppp1r37',	'Relb',	'Clptm1',	'Nectin2',	'Bcam',	'Zfp61',	'Lypd3',	'Zfp526',	'Gsk3a',	'Erf',	'Cic',	'Pafah1b3',	'Megf8',	'Ccdc97',	'Rab4b',	'Mia',	'Map3k10',	'Zfp607b',	'Zfp626',	'Rinl',	'Hnrnpl',	'Ech1',	'Capn12',	'Actn4',	'Sipa1l3',	'Zfp790',	'Zfp566',	'Cox7a1',	'E130208F15Rik',	'Lrfn3',	'Nfkbid',	'U2af1l4',	'Zbtb32',	'Upk1a',	'Lgi4',	'Pdcd2l',	'Kctd15',	'Cebpg',	'Lrp3',	'Zfp536',	'Uri1',	'Ccne1',	'1600014C10Rik',	'Syt3',	'Emc10',	'Nr1h2',	'Cpt1c',	'Prmt1',
                                                      'Scaf1',	'Slc17a7',	'Ccdc155',	'Bax',	'Ppp1r15a',	'Mamstr',	'Cyth2',	'Abcc6',	'Kcnj11',	'Kcnc1',	'Spty2d1',	'Ptpn5',	'E2f8',	'Dbx1',	'Atp10a',	'Mtmr10',	'Tm2d3',	'Selenos',	'Lrrk1',	'Asb7',	'Cers3',	'Adamts17',	'Ttc23',	'Igf1r',	'Fam169b',	'Arrdc4',	'Nr2f2',	'Gm10295',	'Mctp2',	'Rgma',	'Chd2',	'Fam174b',	'St8sia2',	'Slco3a1',	'Klhl25',	'Abhd2',	'Idh2',	'Blm',	'Iqgap1',	'3110040N11Rik',	'Mesd',	'Arnt2',	'Rab38',	'Tmem135',	'Eed',	'Ccdc90b',	'Nars2',	'Kctd14',	'Myo7a',	'Emsy',	'Thap12',	'Wnt11',	'Uvrag',	'Serpinh1',	'Rnf169',	'Kcne3',	'C2cd3',	'Plekhb1',	'Stard10',	'Phox2a',	'Rnf121',	'Trim34a',	'Cyb5r2',	'Tub',	'St5',	'Ascl3',	'Zfp143',	'Wee1',	'Sbf2',	'Ampd3',	'Usp47',	'Dkk3',	'Mical2',	'Parva',	'Rassf10',	'Arntl',	'Spon1',	'Rras2',	'Sox6',	'1110004F10Rik',	'Plekha7',	'Nucb2',	'Knop1',	'Iqck',	'Acsm4',	'Rexo5',	'Mosmo',	'Scnn1g',	'Rbbp6',	'Tnrc6a',	'Kdm8',	'Il21r',	'Mapk3',	'Hirip3',	'Cdipt',	'Pagr1a',	'Cd2bp2',	'E430018J23Rik',	'Bcl7c',	'Fbxl19',	'Orai3',	'Zfp646',	'Prss8',	'Cox6a2',	'Sec23ip',	'Wdr11',	'Ate1',	'Tacc2',	'Btbd16',	'Plekha1',	'Lhpp',	'Edrf1',	'Dhx32',	'Adam12',	'Ptpre',	'Mki67',	'Ebf3',	'Tcerg1l',	'Dpysl4',	'Pwwp2b',	'Inpp5a',	'Cfap46',	'Cox8b',	'Lrrc56',	'Pidd1',	'Ctsd',	'Syt8',	'Ins2',	'Tspan32',	'Tnfrsf22',	'Fadd',	'Ano1',	'Ccnd1',	'1810010D01Rik',	'Mrgprd',	'Iyd',	'Armt1',	'Ust',	'Sash1',	'Stx11',	'Phactr2',	'Aig1',	'Hivep2',	'Cited2',	'Heca',	'Perp',	'Map7',	'1700020N01Rik',	'Slc18b1',	'L3mbtl3',	'Arhgap18',	'Cenpw',	'Ncoa7',	'Hddc2',	'Frk',	'Hdac2',	'Marcks',	'Fam229b',	'Traf3ip2',	'Slc16a10',	'Rpf2',	'Gtf3c6',	'Cdk19',	'Mettl24',	'Cdc40',	'Fig4',	'Cd164',	'Afg1l',	'Cd24a',	'Atg5',	'Prdm1',	'Prep',	'Dcbld1',	'Slc35f1',	'Pkib',	'Sh3rf3',	'P4ha1',	'Gm10273',	'Oit3',	'Ddit4',	'Anapc16',	'Chst3',	'Psap',	'Gm17455',	'Unc5b',	'Ppa1',	'H2afy2',	'Col13a1',	'Dna2',	'Hnrnph3',	'Pbld2',	'Pbld1',	'Reep3',	'Nrbf2',	'Zfp365',	'Rtkn2',	'Arid5b',	'Cabcoco1',	'Rhobtb1',	'Cdk1',	'Ccdc6',	'Rsph14',	'Rab36',	'Cabin1',	'Gstt2',	'Mif',	'Gm5134',	'Zfp280b',	'Prmt2',	'Pofut2',	'Pttg1ip',	'Plpp2',	'Cnn2',	'Sbno2',	'Stk11',	'Ndufs7',	'Reep6',	'Tcf3',	'Btbd2',	'Slc39a3',	'Thop1',	'Eef2',	'Nmrk2',	'Matk',	'Tjp3',	'Tle2',	'Zfp873',	'Gm4924',	'Nfyb',	'Nuak1',	'Rfx4',	'Ric8b',	'Mterf2',	'Cry1',	'Ascl4',	'Fbxo7',	'Syn3',	'Stab2',	'Gnptab',	'Sycp3',	'Spic',	'Gas2l3',	'Anks1b',	'Tmpo',	'Cfap54',	'Cdk17',	'Lta4h',	'Snrpf',	'Plxnc1',	'Cradd',	'Socs2',	'Dcn',	'Atp2b1',	'4930430F08Rik',	'Nts',	'Ptprq',	'Otogl',	'Pawr',	'E2f7',	
                                                      'Bbs10',	'Nap1l1',	'Phlda1',	'Krr1',	'Ptprb',	'Cnot2',	'Myrfl',	'Mdm2',	'Rap1b',	'Grip1',	'Hmga2',	'Msrb3',	'Usp15',	'Lrig3',	'Atp23',	'Avil',	'43899',	'Kif5a',	'Gli1',	'Ndufa4l2',	'Nab2',	'Atp5b',	'Rbms2',	'Spryd4',	'Timeless',	'Pan2',	'Cnpy2',	'Cs',	'Zc3h10',	'Rpl41',	'Pa2g4',	'Erbb3',	'Rps26',	'Ikzf4',	'Dgka',	'Pym1',	'Dnajc14',	'Gdf11',	'Mettl7b',	'Insr',	'Xab2',	'Ccl25',	'Cers4',	'Efnb2',	'Myo16',	'Irs2',	'Col4a2',	'Rab20',	'Ankrd10',	'Tex29',	'Mcf2l',	'Pcid2',	'Cdc16',	'Kbtbd11',	'Ckap2',	'Vps36',	'Polb',	'Ap3m2',	'Kat6a',	'Zmat4',	'Plekha2',	'Plpbp',	'Rnf122',	'Fut10',	'Tex15',	'Gtf2e2',	'Rbpms',	'Ppp1r3b',	'Eri1',	'Prag1',	'Micu3',	'Mtus1',	'Pcm1',	'Frg1',	'Fat1',	'Mtnr1a',	'F11',	'Tlr3',	'Sorbs2',	'Pdlim3',	'Lrp2bp',	'Primpol',	'Casp3',	'Irf2',	'Cdkn2aip',	'Neil3',	'Sap30',	'Mfap3l',	'Clcn3',	'Nek1',	'Sh3rf1',	'Cbr4',	'Apela',	'Tma16',	'Naf1',	'Psd3',	'Sh2d4a',	'Atp6v1b2',	'Zfp930',	'Yjefn3',	'Gatad2a',	'Ddx49',	'Crlf1',	'Jund',	'Kcnn1',	'Abhd8',	'Fcho1',	'Large1',	'Mcm5',	'Arhgap10',	'Tmem184c',	'Zfp827',	'Smad1',	'Otud4',	'Smarca5',	'Gab1',	'Il15',	'Tbc1d9',	'Scoc',	'Dnajb1',	'Adgrl1',	'1700067K01Rik',	'Ier2',	'Lyl1',	'Rad23a',	'Tnpo2',	'Neto2',	'Adcy7',	'Rbl2',	'Fto',	'Irx3',	'Irx5',	'Irx6',	'Amfr',	'Slc12a3',	'Arl2bp',	'Pllp',	'Polr2c',	'Dok4',	'Ccdc102a',	'Adgrg1',	'Cngb1',	'Tepp',	'Csnk2a2',	'Gins3',	'Cklf',	'Nae1',	'Cdh16',	'B3gnt9',	'Slc9a5',	'Kctd19',	'Lrrc36',	'Zdhhc1',	'Atp6v0d1',	'Enkd1',	'Ranbp10',	'Cenpt',	'Psmb10',	'Slc12a4',	'Esrp2',	'Slc7a6',	'Smpd3',	'Cdh3',	'Gm10073',	'Cdh1',	'Tango6',	'Vps4a',	'Psmd7',	'Zfhx3',	'Vac14',	'Mlkl',	'Fa2h',	'Cfdp1',	'Adat1',	'Gm6793',	'Mon1b',	'Wwox',	'Bco1',	'Gan',	'Plcg2',	'Cdh13',	'Wfdc1',	'Fam92b',	'Gins2',	'Klhdc4',	'Slc7a5',	'Mvd',	'Galnt2',	'Ntpcr',	'Pcnx2',	'Kcnk1',	'Slc35f3',	'Irf2bp2',	'Tomm20',	'Pard3',	'Nrp1',	'Fam107a',	'4930452B06Rik',	'Top2b',	'Thrb',	'Kcnk5',	'Nudt13',	'Adk',	'Kat6b',	'Samd8',	'Vdac2',	'Zfp503',	'Dlg5',	'Zmiz1',	'Arf4',	'Tmem110',	'Nt5dc2',	'Sh3bp5',	'Ankrd28',	'Dph3',	'Oxnad1',	'Arhgap22',	'Bmpr1a',	'Sh2d4b',	'Sftpa1',	'Fermt2',	'Gm15217',	'Bmp4',	'Cdkn3',	'Samd4',	'Lgals3',	'Fbxo34',	'Tmem260',	'Klhl33',	'Rnase12',	'Rnase2a',	'Ang4',	'Mettl17',	'Ndrg2',	'Sall2',	'Lrp10',	'Cdh24',	'Slc7a8',	'Cmtm5',	'Ngdn',	'Zfhx2',	'Thtpa',	'Nynrin',	'Mcpt8',	'Parp4',	'Pspc1',	'Zmym5',	'Zmym2',	'Il17d',	'Fgf9',	'Spata13',	'Mipep',	'Tnfrsf19',	'Arl11',	'Wdfy2',	'Pinx1',	'Msra',	'Kif13b',	'Zfp395',	'Pnoc',	'Scara5',	'Pbk',	'Ccdc25',	'Ppp2r2a',	
                                                      'Nefm',	'Nefl',	'Adam7',	'Nkx3-1',	'R3hcc1',	'Pebp4',	'Reep4',	'Npm2',	'Dok2',	'Lpar6',	'Itm2b',	'Lcp1',	'Cog3',	'Tpt1',	'Nufip1',	'Serp2',	'Naa16',	'Kbtbd6',	'Wbp4',	'Tdrd3',	'Lmo7',	'Fbxl3',	'Slain1',	'Ednrb',	'Spry2',	'Slitrk1',	'Gpc5',	'Sox21',	'Abcc4',	'Cldn10',	'Dnajc3',	'Mbnl2',	'Rap2a',	'Farp1',	'Dock9',	'Timm8a2',	'Ggact',	'Tmtc4',	'Alkbh8',	'Gucy1a2',	'Dync2h1',	'Birc3',	'Trpc6',	'Sesn3',	'Endod1',	'Cep295',	'Ubl5',	'Dnmt1',	'Mrpl4',	'Icam4',	'Icam5',	'Fdx1l',	'Ap1m2',	'Qtrt1',	'Ldlr',	'Rab3d',	'Glb1l2',	'Jam3',	'Igsf9b',	'Opcml',	'Ntm',	'Snx19',	'Aplp2',	'Kcnj5',	'Ets1',	'Kirrel3',	'St3gal4',	'Cdon',	'Pus3',	'Stt3a',	'Slc37a2',	'Nrgn',	'Scn3b',	'Hspa8',	'Sorl1',	'Sc5d',	'Tecta',	'Arhgef12',	'Pou2f3',	'Vps11',	'Phldb1',	'Atp5l',	'Ube4a',	'Jaml',	'Cep164',	'Pcsk7',	'Tagln',	'Pafah1b2',	'Sik3',	'Apoa1',	'Cadm1',	'Nnmt',	'Sik2',	'Colca2',	'1810046K07Rik',	'Ddx10',	'Exph5',	'Kdelc2',	'4930550C14Rik',	'Cib2',	'Hykk',	'Rcn2',	'Tspan3',	'Lingo1',	'Snupn',	'Sin3a',	'Ppcdc',	'Edc3',	'Clk3',	'Arid3b',	'Sema7a',	'Pml',	'Cd276',	'Nptn',	'Pkm',	'Senp8',	'Nr2e3',	'Thsd4',	'Uaca',	'Tle3',	'Rplp1',	'Kif23',	'Paqr5',	'Anp32a',	'Coro2b',	'Map2k5',	'Iqch',	'Aagab',	'Smad3',	'Smad6',	'Snapc5',	'Tipin',	'Dis3l',	'Rab11a',	'Parp16',	'Clpx',	'Oaz2',	'Snx22',	'Snx1',	'Dapk2',	'Herc1',	'Usp3',	'Rps27l',	'Lactb',	'Tpm1',	'Tln2',	'Ice2',	'Anxa2',	'Myo1e',	'Aldh1a2',	'Polr2m',	'Myzap',	'Zfp280d',	'Mns1',	'Rfx7',	'Nedd4',	'Prtg',	'Pygo1',	'Wdr72',	'Myo5c',	'Bmp5',	'Hmgcll1',	'Hcrtr2',	'Fam83b',	'Mlip',	'Lrrc1',	'Gclc',	'Gsta4',	'Col12a1',	'Irak1bp1',	'Sh3bgrl2',	'Elovl4',	'Ube2cbp',	'Cep162',	'Ctsh',	'Adamts7',	'Plscr1',	'Spsb4',	'Clstn2',	'Nme9',	'Dzip1l',	'Il20rb',	'Cep63',	'Amotl2',	'Slco2a1',	'Srprb',	'1300017J02Rik',	'Tmem108',	'Acpp',	'Atp2c1',	'Pik3r4',	'Col6a6',	'Poc1a',	'Rpl29',	'Manf',	'Cacna2d2',	'Hyal2',	'Sema3f',	'Mst1r',	'Traip',	'Amigo3',	'Dag1',	'Nicn1',	'Rhoa',	'Lamb2',	'Qrich1',	'Dalrd3',	'Ip6k2',	'Cdc25a',	'Klhl18',	'Cmtm8',	'Stt3b',	'Tgfbr2',	'Vill',	'Exog',	'Scn5a',	'Eif1b',	'Entpd3',	'Ctnnb1',	'Ulk4',	'Pomgnt2',	'Topaz1',	'Tcaim',	'Limd1',	'Sacm1l',	'Pik3ip1',	'Lif',	'Zmat5',	'Nf2',	'Emid1',	'Kremen1',	'Znrf3',	'Polm',	'H2afv',	'Gm11985',	'Igfbp1',	'Spata48',	'Fignl1',	'Ddc',	'Cobl',	'Egfr',	'Wdr92',	'Peli1',	'Ugp2',	'Fam161a',	'4931440F15Rik',	'Sptbn1',	'Ubtd2',	'Kcnmb1',	'Wwc1',	'Clint1',	'Thg1l',	'Sox30',	'Irgm1',	'Olfr1396',	'Cby3',	'Zfp879',	'Zfp2',	'Zfp354b',	'Zfp354a',	'Phykpl',	'Jade2',	'Skp1a',	'Tcf7',
                                                      'Gm10447',	'Gm12216',	'Slc22a4',	'Pdlim4',	'Anxa6',	'Gm2a',	'Fat2',	'Sparc',	'Atox1',	'G3bp1',	'Mfap3',	'Larp1',	'Zfp692',	'Guk1',	'Wnt9a',	'Mprip',	'Cops3',	'Rai1',	'Srebf1',	'Llgl1',	'Zswim7',	'Trpv2',	'Lrrc75a',	'Cox10',	'F930015N05Rik',	'Dnah9',	'Tmem220',	'Dhrs7c',	'Ntn1',	'Pik3r6',	'Mfsd6l',	'Rpl26',	'Aurkb',	'Borcs6',	'Tmem107',	'Hes7',	'Aloxe3',	'Gucy2e',	'Chd3',	'Kdm6b',	'Dnah2',	'Wrap53',	'Eif5a',	'Cldn7',	'Dlg4',	'Pelp1',	'Mink1',	'Tekt1',	'Ggt6',	'Spns2',	'P2rx1',	'Ncbp3',	'Haspin',	'Rpa1',	'Ywhae',	'Rph3al',	'Rflnb',	'Glod4',	'Timm22',	'Ssh2',	'Coro6',	'Ankrd13b',	'Git1',	'Myo18a',	'Gm10277',	'Phf12',	'Fam222b',	'Traf4',	'Rpl23a',	'Aldoc',	'Wsb1',	'Gm9964',	'Suz12',	'Crlf3',	'Rnf135',	'Psmd11',	'Myo1d',	'Spaca3',	'Rasl10b',	'Taf15',	'Wfdc21',	'Hnf1b',	'Ddx52',	'Dusp14',	'Lhx1',	'Car4',	'Tbx2',	'Gm11444',	'Brip1',	'Med13',	'Ypel2',	'Ska2',	'Rad51c',	'Tex14',	'Rnf43',	'Supt4a',	'Dynll2',	'Cuedc1',	'Mrps23',	'Msi2',	'Akap1',	'Scpep1',	'Gm525',	'Ankfn1',	'Pctp',	'Tmem100',	'Hlf',	'Tom1l1',	'Utp18',	'Epn3',	'Samd14',	'Dlx3',	'Tac4',	'Fam117a',	'Spop',	'Nxph3',	'Phb',	'Abi3',	'Prr15l',	'Sp2',	'4933428G20Rik',	'Srcin1',	'Epop',	'Mllt6',	'Pcgf2',	'Lasp1',	'Fbxo47',	'Plxdc1',	'Pgap3',	'Grb7',	'Wipf2',	'Cdc6',	'Rara',	'Top2a',	'Tns4',	'Krt20',	'Krt23',	'Krtap1-5',	'Krt19',	'Eif1',	'Hap1',	'Jup',	'P3h4',	'Fkbp10',	'Atp6v0a1',	'Coasy',	'Plekhh3',	'Ccr10',	'Aarsd1',	'Rdm1',	'Arl4d',	'Etv4',	'Sost',	'Dusp3',	'Nags',	'Gm11627',	'Fzd2',	'Adam11',	'Eftud2',	'Gfap',	'Nmt1',	'Fmnl1',	'Map3k14',	'Arhgap27',	'Mettl2',	'Mrc2',	'43900',	'Prr29',	'Icam2',	'Ern1',	'Tex2',	'Smurf2',	'1810010H24Rik',	'Pitpnc1',	'Axin2',	'Arsg',	'Wipi1',	'Fam20a',	'Abca8b',	'Map2k6',	'Slc39a11',	'Sstr2',	'Slc9a3r1',	'Tmem104',	'Atp5h',	'Jpt1',	'Tmem94',	'Caskin2',	'Recql5',	'Itgb4',	'Wbp2',	'St6galnac2',	'Srsf2',	'44083',	'Engase',	'Rbfox3',	'Cbx2',	'Cbx8',	'Nptx1',	'Rptor',	'Baiap2',	'Aatk',	'Slc38a10',	'Bahcc1',	'Actg1',	'Slc25a10',	'P4hb',	'Arhgdia',	'Aspscr1',	'Rfng',	'Tbcd',	'Ptchd3',	'Calml3',	'Net1',	'Akr1e1',	'Klf6',	'Ero1lb',	'Gpr137b',	'Lyst',	'Psma2',	'Mplkip',	'Nme8',	'Elmo1',	'Trim27',	'Zscan12',	'Zkscan4',	'Hist1h2bl',	'Hist1h4j',	'Zfp184',	'Abt1',	'Btn1a1',	'Hist1h4h',	'Hist1h2af',	'Hist1h3g',	'Hist1h2bh',	'Hist1h4f',	'Hist1h1d',	'Hist1h3e',	'Hist1h2ae',	'Hist1h4d',	'Hist1h2be',	'Hist1h2ac',	'Hist1h2bc',	'Hist1h4c',	'Hist1h3c',	'Hist1h4b',	'Hist1h4a',	'Carmil1',	'Sox4',	'Cdkal1',	'Agtr1a',	'Hus1b',	'Gmds',	'Serpinb6b',	'Serpinb1b',	'Bphl',	'Psmg4',	'Slc22a23',	
                                                      'Pxdc1',	'Cdyl',	'Ly86',	'Rreb1',	'Ssr1',	'Dsp',	'Bmp6',	'Ofcc1',	'Nedd9',	'Tmem170b',	'Edn1',	'Phactr1',	'Jarid2',	'Atxn1',	'Kif13a',	'Rnf144b',	'Id4',	'Fam120a',	'Ninj1',	'Spin1',	'S1pr3',	'Shc3',	'Secisbp2',	'Gadd45g',	'Diras2',	'Nfil3',	'Sptlc1',	'Thoc3',	'Gprin1',	'Unc5a',	'Slc34a1',	'Dbn1',	'Pdlim7',	'B4galt7',	'B230219D22Rik',	'Txndc15',	'Pcbd2',	'Spock1',	'Idnk',	'Ntrk2',	'Dapk1',	'Zfp808',	'6720489N17Rik',	'Platr25',	'Gm5141',	'Fancc',	'Ptch1',	'Hsd17b3',	'Slc35d2',	'Zfp367',	'Aaed1',	'Uqcrb',	'Zfp429',	'Zfp738',	'Zfp85',	'Mtrr',	'1700001L19Rik',	'Adamts16',	'Irx1',	'Irx2',	'Ndufs6',	'Lpcat1',	'Tert',	'Slc12a7',	'Erap1',	'Pou5f2',	'Edil3',	'Hapln1',	'Ssbp2',	'Fam151b',	'Serinc5',	'Cmya5',	'Bhmt',	'Wdr41',	'Aggf1',	'F2rl1',	'F2r',	'Poc5',	'Ankdd1b',	'Fam169a',	'Enc1',	'Gm10260',	'Arhgef28',	'Tmem171',	'Fcho2',	'Cartpt',	'Mccc2',	'Gtf2h2',	'Taf9',	'Mrps36',	'Pik3r1',	'Cd180',	'Srek1',	'Nln',	'Adamts6',	'Ndufaf2',	'Smim15',	'Rab3c',	'Plk2',	'Ankrd55',	'Plpp1',	'Ccno',	'Cdc20b',	'Gpx8',	'Snx18',	'Arl15',	'Ndufs4',	'Fst',	'Zfp131',	'Asxl2',	'Dnmt3a',	'Sdc1',	'Fam49a',	'Mycn',	'Nbas',	'Lpin1',	'Ntsr2',	'E2f6',	'2410004P03Rik',	'Kcnf1',	'Atp6v1c2',	'Hpcal1',	'Rrm2',	'Id2',	'Cmpk2',	'Adi1',	'Tmem18',	'Slc26a3',	'Pik3cg',	'Sypl',	'Atxn7l1',	'Agr3',	'Tspan13',	'Ifrd1',	'Dock4',	'Prkd1',	'Coch',	'Nubpl',	'Akap6',	'Eapp',	'Brms1l',	'Mbip',	'Nkx2-1',	'Nkx2-9',	'Foxa1',	'Trappc6b',	'Gm9887',	'Pygl',	'Trim9',	'Tmx1',	'Frmd6',	'Psma3',	'Tomm20l',	'Dact1',	'Daam1',	'Lrrc9',	'Pcnx4',	'Trmt5',	'Tmem30b',	'Hif1a',	'Gphb5',	'Ppp2r5e',	'Wdr89',	'Syne2',	'Esr2',	'Plekhg3',	'Rab15',	'Fut8',	'Gm6657',	'Fam71d',	'Tmem229b',	'Zfyve26',	'Rad51b',	'Zfp36l1',	'Actn1',	'Dcaf5',	'Slc39a9',	'Ccdc177',	'Susd6',	'Srsf5',	'Ttc9',	'Map3k9',	'Pcnx',	'Sipa1l1',	'Rgs6',	'Dpf3',	'Dcaf4',	'Zfyve1',	'Fam161b',	'Lin52',	'Npc2',	'Isca2',	'Flvcr2',	'Ttll5',	'Tgfb3',	'Irf2bpl',	'Cipc',	'Pomt2',	'Gtf2a1',	'Galc',	'Ptpn21',	'Ttc8',	'Foxn3',	'Efcab11',	'Psmc1',	'Calm1',	'Ttc7b',	'Rps6ka5',	'Ccdc88c',	'Slc24a4',	'Serpina6',	'Clmn',	'Setd3',	'Degs2',	'Mok',	'Lbhd2',	'Gm266',	'Ckb',	'Aspg',	'Inf2',	'Akt1',	'Cep170b',	'Tedc1',	'Wdr60',	'Cdca7l',	'Dab2',	'Slc1a3',	'Nadk2',	'Rai14',	'Tars',	'Zfr',	'Basp1',	'Myo10',	'Retreg1',	'Otulin',	'Trio',	'Dnah5',	'Ctnnd2',	'43896',	'Cct5',	'Sema5a',	'Matn2',	'Osr2',	'Cox6c',	'Ankrd46',	'Ywhaz',	'Zfp706',	'Grhl2',	'Ubr5',	'Klf10',	'Nudcd1',	'Eif3h',	'Rad21',	'Med30',	'Ext1',	'Taf2',	'Fam83a',	'Atad2',	'Ndufb9',	'Mtss1',
                                                      'Nsmce2',	'Myc',	'Fam49b',	'Asap1',	'Tmem71',	'Tg',	'St3gal1',	'Zfat',	'Slc45a4',	'Jrk',	'Slurp1',	'Mafa',	'Puf60',	'Exosc4',	'Scx',	'Adck5',	'Slc39a4',	'Zfp251',	'Mb',	'Rbfox2',	'Myh9',	'Txn2',	'Ift27',	'Mpst',	'Kctd17',	'Mfng',	'Gga1',	'Pdxp',	'Ddx17',	'Gtpbp1',	'Cbx7',	'Syngr1',	'Tnrc6b',	'Sgsm3',	'Mkl1',	'Dnajb7',	'Zc3h7b',	'Tob2',	'Xrcc6',	'Srebf2',	'Cenpm',	'Poldip3',	'Cyb5r3',	'Arfgap3',	'Arhgap8',	'Nup50',	'Upk3a',	'Atxn10',	'Trmu',	'Celsr1',	'Tbc1d22a',	'Rabl2',	'Pphln1',	'Ano6',	'Arid2',	'Scaf11',	'Slc38a1',	'Slc38a4',	'Adcy6',	'Ddn',	'Tuba1b',	'Tuba1c',	'Prph',	'Spats2',	'Kcnh3',	'Lima1',	'Dip2b',	'Atf1',	'Slc11a2',	'Slc4a8',	'Atg101',	'6030408B16Rik',	'Krt8',	'Eif4b',	'Mfsd5',	'Espl1',	'Npff',	'Atp5g2',	'Hoxc4',	'Cbx5',	'Hnrnpa1',	'Nfe2',	'Gpr84',	'Adcy9',	'Tfap4',	'Nmral1',	'Cdip1',	'Eef2kmt',	'Abat',	'1810013L24Rik',	'Tekt5',	'Ciita',	'Clec16a',	'Socs1',	'Mkl2',	'Fgd4',	'Top3b',	'Ccdc116',	'Hic2',	'Lrrc74b',	'Smpd4',	'Klhl22',	'Trmt2a',	'Comt',	'Txnrd2',	'Gp1bb',	'Klhl6',	'Abcc5',	'Ap2m1',	'Eif4g1',	'Fam131a',	'Ephb3',	'Vps8',	'Etv5',	'Crygs',	'Adipoq',	'St6gal1',	'Bcl6',	'Lpp',	'Tprg',	'P3h2',	'Mb21d2',	'Atp13a5',	'Opa1',	'Hes1',	'Tmem44',
                                                      'Fam43a',	'Xxylt1',	'Bdh1',	'Dlg1',	'Zdhhc19',	'Tfrc',	'Tnk2',	'Iqcg',	'Lmln',	'Osbpl11',	'Snx4',	'Slc12a8',	'Umps',	'Kalrn',	'Ropn1',	'Hacd2',	'Fam162a',	'Stfa3',	'Slc15a2',	'Gpr156',	'Zbtb20',	'Qtrt2',	'Zdhhc23',	'Gramd1c',	'Sidt1',	'Cfap44',	'Ccdc80',	'Phldb2',	'Nectin3',	'Morc1',	'Cblb',	'Alcam',	'Nit2',	'Tbc1d23',	'Cmss1',	'Nsun3',	'Cggbp1',	'Chmp2b',	'Cxadr',	'Adamts1',	'N6amt1',	'Bach1',	'Scaf4',	'Mis18a',	'4932438H23Rik',	'Rcan1',	'Runx1',	'Setd4',	'Cbr3',	'Morc3',	'Chaf1b',	'Kcnj15',	'Ets2',	'Igsf5',	'Pcp4',	'Dscam',	'Bace2',	'Mx1',	'Tmprss2',	'A630089N07Rik',	'Cldn20',	'Arid1b',	'Ldhal6b',	'Tagap',	'Rsph3a',	'Qk',	'Pacrg',	'Dact2',	'Smoc2',	'Rgmb',	'Spaca6',	'Zfp53',	'Zfp13',	'Bicdl2',	'Tnfrsf12a',	'Dcpp3',	'Kctd5',	'Noxo1',	'Tpsg1',	'Sstr5',	'Mettl26',	'Mrpl28',	'Luc7l',	'Cuta',	'D17Wsu92e',	'Tead3',	'Tulp1',	'Brpf3',	'Srsf3',	'Ppil1',	'Pim1',	'Rnf8',	'Zfand3',	'Btbd9',	'Dnah8',	'Tmprss3',	'Pde9a',	'Ndufv3',	'Cryaa',	'Sik1',	'Hsf2bp',	'Notch3',	'Ephx3',	'Akap8l',	'Cyp4f17',	'Rab11b',	'Zbtb22',	'Rgl2',	'H2-DMb2',	'Lsm2',	'Vars',	'Abhd16a',	'Prrc2a',	'H2-Q10',	'Gtf2h4',	'Tubb5',	'Ppp1r10',	'H2-T23',	'Mog',	'9130008F23Rik',	'Adgrf2',	'Cd2ap',	'Rcan2',	'Enpp5',	'Clic5',	'Supt3',	'Cdc5l',	'Tmem151b',	'Slc35b2',	'Hsp90ab1',	'Mrpl14',	'1600014C23Rik',	'Vegfa',	'Mad2l1bp',	'Ptk7',	'Ubr2',	'Mrps10',	'Guca1a',	'Taf8',	'Ccnd3',	'1700122O11Rik',	'1700067P10Rik',	'Nfya',	'Lrfn2',	'Daam2',	'Satb1',	'Mpnd',	'Fem1a',	'Ticam1',	'Gtf2f1',	'Slc25a23',	'Efna5',	'Ralbp1',	'Ptprm',	'Lama1',	'Epb41l3',	'Zbtb14',	'Tgif1',	'Lpin2',	'Yipf4',	'Fez2',	'Sult6b1',	'Ndufaf7',	'Rmdn2',	'Galm',	'Gemin6',	'Dhx57',	'Cdkl4',	'Pkdcc',	'Eml4',	'Kcng3',	'Mta3',	'Haao',	'Zfp36l2',	'Thada',	'Plekhh2',	'Slc3a1',	'Camkmt',	'Srbd1',	'Gm10309',	'Mcfd2',	'Calm2',	'Epcam',	'Msh2',	'Msh6',	'Foxn2',	'Gtf2a1l',	'Gm47283',	'Bambi',	'Svil',	'Zfp438',	'Epc1',	'Gata6',	'Rbbp8',	'Cables1',	'Npc1',	'Ankrd29',	'Zfp521',	'Aqp4',	'Garem1',	'Dtna',	'Mapre2',	'Zfp24',	'Rprd1a',	'AW554918',	'Celf4',	'Ammecr1l',	'Bin1',	'Camk4',	'Nrep',	'Epb41l4a',	'Apc',	'Brd8',	'Ctnna1',	'Lrrtm2',	'Matr3',	'Spata24',	'Dnajc18',	'Psd2',	'Nrg2',	'Cystm1',	'Hbegf',	'Pcdhgc3',	'Pcdhgc5',	'Pcdh1',	'Fgf1',	'Pabpc2',	'Yipf5',	'Prelid2',	'Rbm27',	'Ccdc112',	'Sema6a',	'Dtwd2',	'Srfbp1',	'Lox',	'Ppic',	'Cep120',	'Zfp608',	'Phax',	'Megf10',	'Ctxn3',	'Slc12a2',	'Fbn2',	'Chsy3',	'Slc6a7',	'Ppargc1b',	'Afap1l1',	'Sh3tc2',	'Alpk2',	'Sec11c',	'Pmaip1',	'Seh1l',	'Fam210a',	'Tcf4',	'Rab27b',	'Mex3c',	'Smad4',
                                                      'Myo5b',	'Acaa2',	'Lipg',	'Rpl17',	'Smad7',	'Smad2',	'Atp5a1',	'Gm6133',	'Setbp1',	'Pard6g',	'Txnl4a',	'Nfatc1',	'Zfp236',	'Zfp516',	'Tshz1',	'Timm21',	'Tmx3',	'Ppp6r3',	'Acy3',	'Rps6kb2',	'Grk2',	'Rce1',	'Zdhhc24',	'Mrpl11',	'Npas4',	'Banf1',	'Scyl1',	'Frmd8',	'Tigd3',	'Capn1',	'Sac3d1',	'Sf1',	'Macrod1',	'Otub1',	'Rcor2',	'Mark2',	'2700081O15Rik',	'Atl3',	'Pla2g16',	'Bscl2',	'Ganab',	'Eef1g',	'Scgb1a1',	'Fads3',	'Fads2',	'Syt7',	'Cd6',	'Tmem132a',	'Tmem109',	'Mrpl16',	'Mpeg1',	'Dtx4',	'Tle4',	'Cep78',	'Prune2',	'Anxa1',	'Gm3443',	'Trpm3',	'Fxn',	'Pip5k1b',	'Fam122a',	'Tmem252',	'Dock8',	'Kank1',	'Dmrt1',	'Gm815',	'4430402I18Rik',	'Plgrkt',	'Pdcd1lg2',	'Ric1',	'Ranbp6',	'Dkk1',	'Asah2',	'Sgms1',	'Pten',	'Rnls',	'Slc16a12',	'Pank1',	'Ppp1r3c',	'Btaf1',	'Cpeb3',	'Exoc6',	'Myof',	'Noc3l',	'Pdlim1',	'Sorbs1',	'Aldh18a1',	'Cc2d2b',	'Opalin',	'Tm9sf3',	'Slit1',	'Mms19',	'Cyp2c23',	'Cwf19l1',	'Scd2',	'Slf2',	'Sfxn3',	'Btrc',	'Poll',	'Npm3',	'Pprc1',	'Cnnm2',	'Gm6970',	'Neurl1a',	'Stn1',	'Cfap58',	'Xpnpep1',	'Add3',	'Mxi1',	'Dusp5',	'Smc3',	'Rbm20',	'Tectb',	'Acsl5',	'Tcf7l2',	'Habp2',	'Adrb1',	'Afap1l2',	'Ablim1',	'Trub1',	'Gfra1',	'Hspa12a',	'Eno4',	'Emx2',	'Rab11fip2',	'Csf2ra'))

integrate_epi4 <- AddModuleScore(object = integrate_epi4, features = Both_NKX21_binding_retained_from_progenitor, ctrl = 500, name = "NKXBothRetainedGenes", search = TRUE, random.seed = 1)
#not accounted for 43891, 43899, 43900, 44083, 43896
FeaturePlot(object = integrate_epi4, features = 'NKXBothRetainedGenes1', cols = c('lightgrey','red'))


Pr_more <- list(c('Xkr4',	'Jph1',	'Fam178b',	'Gm3646',	'Pou3f3',	'Nck2',	'Nrp2',	'Rpe',	'Lancl1',	'Cps1',	'Erbb4',	'Igfbp5',	'Epha4',	'Acsl3',	'Pid1',	'Glrp1',	'Cops8',	'Twist2',	'Serpinb8',	'Cxcr4',	'Lemd1',	'Ube2t',	'Lgr6',	'Ptprc',	'Cdc73',	'Hmcn1',	'Teddm2',	'Fam163a',	'Brinp2',	'Cd247',	'Rxrg',	'Smyd3',	'1700056E22Rik',	'Lyplal1',	'Tgfb2',	'Esrrg',	'Kctd3',	'Smyd2',	'Prox1',	'Camk1g',	'Plxna2',	'Camk1d',	'St8sia6',	'Hacd1',	'Nebl',	'Notch1',	'Sardh',	'Rxra',	'Ppp1r26',	'Barhl1',	'Med27',	'Zbtb43',	'Crb2',	'Nr5a1',	'Arhgap15',	'Fmnl2',	'Nr4a2',	'Tank',	'Dpp4',	'Zfp385b',	'Prdx6b',	'Zfp804a',	'Slc43a3',	'Slc1a2',	'Wt1',	'Them7',	'Pax6',	'Mpped2',	'Arl14ep',	'Ryr3',	'Rasgrp1',	'Thbs1',	'Frmd5',	'Sord',	'Gatm',	'Sqor',	'Gm9913',	'Ncaph',	'Bcl2l11',	'Mrps26',	'Bmp2',	'Plcb4',	'Snap25',	'Ovol2',	'Syndig1',	'Gm14226',	'Src',	'Bpi',	'Dhx35',	'Mafb',	'Tox2',	'Sulf2',	'B4galt5',	'Kcng1',	'Sall4',	'Tfap2c',	'Bmp7',	'Ankrd60',	'Stx16',	'Ccdc160',	'Asb12',	'Snx12',	'Pak3',	'Zfp704',	'Pag1',	'Dnajc5b',	'Fndc3b',	'Tmem212',	'Egfem1',	'Slc7a14',	'Sox2',	'Mab21l1',	'Gpr87',	'Plch1',	'Iqcj',	'Rxfp1',	'Pdgfc',	'Sfrp2',	'Tmem131l',	'Kirrel',	'Fcgr1',	'Vangl1',	'Cym',	'A630076J17Rik',	'Vav3',	'Col11a1',	'Cdc14a',	'Plppr5',	'Dpyd',	'Ptbp2',	'Slc44a3',	'Ugt8a',	'Elovl6',	'Lef1',	'Dkk2',	'Adh6a',	'Unc5c',	'Bmpr1b',	'Pkn2',	'Col24a1',	'Cyr61',	'St6galnac5',	'Triqk',	'Tmem64',	'Fut9',	'Bach2',	'Lingo2',	'Pax5',	'Igfbpl1',	'Xpa',	'Tgfbr1',	'Rad23b',	'Ambp',	'Astn2',	'Trim32',	'Brinp1',	'Gm11232',	'Slc24a2',	'Mtap',	'Jun',	'Fggy',	'Hook1',	'E130114P18Rik',	'Lepr',	'Pde4b',	'Dmrta2',	'3100002H09Rik',	'Nkain1',	'Sdc3',	'Matn1',	'Ptpru',	'Ephb2',	'Wnt4',	'Igsf21',	'Camta1',	'A430005L14Rik',	'Fam133b',	'Steap4',	'Speer4c',	'Mnx1',	'Garem2',	'Cad',	'Babam2',	'Cpz',	'Sorcs2',	'Zfp518b',	'Gm16223',	'Fgfbp1',	'Prom1',	'Slit2',	'Ppargc1a',	'Smim20',	'Pcdh7',	'Klf3',	'9130230L23Rik',	'Lnx1',	'Adgrl3',	'Npffr2',	'Fras1',	'Prkg2',	'Arhgap24',	'Mapk10',	'Asphd2',	'Sez6l',	'Myo18b',	'Aym1',	'Gcn1l1',	'Hspb8',	'Ksr2',	'Nos1',	'2410131K14Rik',	'Rbm19',	'Rflna',	'Tmem132c',	'Fzd10',	'Galnt17',	'Auts2',	'Gtf2ird1',	'Cldn4',	'Ephb4',	'Mad1l1',	'Tex26',	'Slc25a13',	'Glcci1',	'Fam3c',	'Aass',	'Mest',	'Plxna4',	'Chchd3',	'Hipk2',	'Tmem178b',	'Tpk1',	'Cntnap2',	'Gimap7',	'Igf2bp3',	'Snx10',	'Creb5',	'Herc6',	'Atoh1',	'Mad2l1',	'Hk2',	'Gfpt1',	'Iqsec1',	'1700123L14Rik',	'Frmd4b',	'Rybp',	'Lrrn1',	'Lmcd1',	'Hrh1',	'Zfp637',	'Ntf3',	'Etv6',	'Bcl2l14',	'Grin2b',	'Gucy2c',	'Arhgdib',	'Ptpro',
                  'Lmo3',	'Pde3a',	'Etnk1',	'Sox5',	'Usp29',	'Ccdc8',	'Tshz3',	'Zfp536',	'Kcnj11',	'Htatip2',	'Chrna7',	'Gm10295',	'St8sia2',	'Tmem135',	'Ccdc90b',	'Rab30',	'Emsy',	'Wnt11',	'Adm',	'Rassf10',	'Spon1',	'Insc',	'Plekha7',	'Acsm1',	'Acsm4',	'Dnah3',	'Eef2k',	'Zkscan2',	'Wdr11',	'Nps',	'Mgmt',	'Kndc1',	'Ifitm10',	'Mrpl23',	'Cttn',	'Plekhg1',	'Akap12',	'Myb',	'Echdc1',	'Amd2',	'Crybg1',	'Chst3',	'Ctnna3',	'Lrrtm3',	'Arid5b',	'Phyhipl',	'Pofut2',	'Chst11',	'Appl2',	'1700113H08Rik',	'Cep83',	'Btg1',	'Lum',	'Kitl',	'Otogl',	'Bbs10',	'Ptprr',	'Cpm',	'Hmga2',	'Efnb2',	'Mcf2l',	'Csmd1',	'Unc5d',	'Ppp1r3b',	'Cldn23',	'Cenpu',	'Enpp6',	'Dctd',	'Tenm3',	'Galnt7',	'Cbr4',	'Psd3',	'Cers1',	'Comp',	'Plvap',	'Klf2',	'Hhip',	'Il15',	'Ndufb7',	'Sall1',	'Tox3',	'Rbl2',	'Irx5',	'Ces1a',	'Drc7',	'Setd6',	'Cdh8',	'Cdh11',	'Cdh3',	'Psmd7',	'Zfhx3',	'Fa2h',	'Adamts18',	'Maf',	'Tldc1',	'Foxl1',	'Map3k21',	'Tomm20',	'Nrp1',	'Top2b',	'Gng2',	'Slc35f4',	'Tnfrsf19',	'Gata4',	'Gfra2',	'Htr2a',	'Olfm4',	'Pcdh20',	'Pcdh9',	'Klf12',	'Ipo5',	'Stk24',	'Nalcn',	'Fgf14',	'Sesn3',	'Jam3',	'Igsf9b',	'Opcml',	'Ntm',	'St14',	'Barx2',	'St3gal4',	'Pknox2',	'Sorl1',	'Tbcel',	'Grik4',	'Pou2f3',	'Nxpe4',	'Elmod1',	'Cspg4',	'1700017B05Rik',	'Loxl1',	'Uaca',	'Tle3',	'Igdcc3',	'Tpm1',	'Rora',	'Impg1',	'Sh3bgrl2',	'Ibtk',	'Slc9a9',	'Rasa2',	'Mrps22',	'Mrpl3',	'Arpp21',	'Trim71',	'Scn5a',	'Mobp',	'Pkd1l1',	'Hus1',	'Vwc2',	'Meis1',	'Bcl11a',	'Ccdc85a',	'Sptbn1',	'Srebf1',	'Dnah9',	'Glp2r',	'Ntn1',	'Smtnl2',	'1700016K19Rik',	'Wsb1',	'Mrm1',	'Bcas3',	'Tbx2',	'Nog',	'Ankfn1',	'Pctp',	'Hils1',	'Tac4',	'Phb',
                  'Igf2bp1',	'Ikzf3',	'Krt20',	'Krt13',	'G6pc3',	'Wnt9b',	'1700012B07Rik',	'Map2k6',	'Kcnj16',	'Sox9',	'Sdk2',	'Tnrc6c',	'Cyth1',	'Cbx4',	'Zfp750',	'Klf6',	'Chrm3',	'Elmo1',	'Gpld1',	'Nrsn1',	'Cdkal1',	'Gmds',	'Mylk4',	'Hivep1',	'Phactr1',	'Rnf144b',	'Id4',	'Sema4d',	'Diras2',	'Ror2',	'Msx2',	'Pitx1',	'Spock1',	'Agtpbp1',	'Gas1',	'Dapk1',	'Fbp2',	'2010111I01Rik',	'Ercc6l2',	'Hsd17b3',	'Irx2',	'Vcan',	'Zbed3',	'Poc5',	'Cartpt',	'Naip6',	'Srek1',	'Isl1',	'B430218F22Rik',	'Dnmt3a',	'Pomc',	'Dnajc27',	'Fam49a',	'Sox11',	'Sntg2',	'Twist1',	'Sostdc1',	'Dgkb',	'Npas3',	'Egln3',	'Dact1',	'Six1',	'Slc38a6',	'Actn1',	'Sipa1l1',	'Dpf3',	'Jdp2',	'Tgfb3',	'Esrrb',	'Sel1l',	'Dlk1',	'Akt1',	'Dnah11',	'Ghr',	'Slc1a3',	'Adamts12',	'Npr3',	'Basp1',	'Otulin',	'Ctnnd2',	'Ankrd33b',	'Trps1',	'Sntb1',	'Tmem65',	'Myc',	'Fam49b',	'St3gal1',	'Slc45a4',	'Gpr20',	'Ly6h',	'Mb',	'Tcf20',	'Rtl6',	'Fam118a',	'Fam19a5',	'Syt10',	'Tmem117',	'Slc38a4',	'Col2a1',	'Smug1',	'Sec14l5',	'Eef2kmt',	'Rtl10',	'Klhl6',	'St6gal1',	'Il1rap',	'Gmnc',	'Fgf12',	'Opa1',	'Fam43a',	'Tfrc',	'Hacd2',	'Csta1',	'Gpr156',	'Igsf11',	'Lsamp',	'Bach1',	'Setd4',	'Ets2',	'B3galt5',	'Pcp4',	'Dact2',	'Cldn9',	'1520401A03Rik',	'Lemd2',	'Glp1r',	'Pde9a',	'H2-Q2',	'Supt3',	'Spats1',	'1600014C23Rik',	'Efna5',	'Ptprm',	'Arhgap28',	'Epb41l3',	'Dlgap1',	'Tgif1',	'Crim1',	'Pkdcc',	'Haao',	'Thada',	'Slc3a1',	'Six2',	'Srbd1',	'Zfp521',	'Cdh2',	'B4galt6',	'Arap3',	'Pabpc2',	'Ppp2r2b',	'Spink1',	'Sema6a',	'Eno1b',	'Cep120',	'Zfp608',	'Prrc1',	'Slc27a6',	'St8sia3',	'Zfp532',	'Pmaip1',	'Tcf4',	'Ccdc68',	'Dcc',	'Timm21',	'Nrxn2',	'Gnaq',	'Rfk',	'Trpm3',	'Mamdc2',	'Dmrt3',	'Gm815',	'Dkk1',	'Pten',	'Kif20b',	'Gm340',	'Sorcs3',	'Sorcs1',	'Add3',	'Tectb',	'Tcf7l2',	'Habp2',	'Gfra1',	'Pnlip',	'Emx2'))

integrate_epi4 <- AddModuleScore(object = integrate_epi4, features = Pr_more, ctrl = 500, name = "NKXProgMoreGenes", search = TRUE, random.seed = 1)
FeaturePlot(object = integrate_epi4, features = 'NKXProgMoreGenes1', cols = c('lightgrey','red'))

Pr_less <- list(c('Arfgef1',	'Slco5a1',	'Stau2',	'Pkhd1',	'Col9a1',	'Tmem131',	'Inpp4a',	'2010300C02Rik',	'Rpl31',	'Il1rl2',	'1500015O10Rik',	'Slc39a10',	'1700019D03Rik',	'Stk17b',	'Spats2l',	'Nrp2',	'Acadl',	'Lancl1',	'Cps1',	'Ikzf2',	'Rpl37a',	'Tmbim1',	'Speg',	'Epha4',	'Fam124b',	'Dner',	'Cab39',	'Kcnj13',	'Gpc1',	'Slc35f5',	'Gpr39',	'Nckap5',	'Tmem163',	'Pigr',	'Fam72a',	'Slc26a9',	'Cntn2',	'Nfasc',	'Mdm4',	'Pik3c2b',	'Prelp',	'Arl8a',	'Ptprc',	'Fam129a',	'Edem3',	'Lamc1',	'Gm10031',	'Fam20b',	'Angptl1',	'4930523C07Rik',	'Kifap3',	'Dpt',	'Mpzl1',	'Gpa33',	'Rgs5',	'B4galt3',	'Fmn2',	'Grem2',	'1700016C15Rik',	'Smyd3',	'Rrp15',	'Spata17',	'Kctd3',	'Dtl',	'Hhat',	'A130010J15Rik',	'Camk1g',	'Cd34',	'Phyh',	'Optn',	'Usp6nl',	'Dnajc1',	'Msrb2',	'Arhgap21',	'Zdhhc12',	'Hmcn2',	'Qrfp',	'Lmx1b',	'Mvb12b',	'Pbx3',	'Dab2ip',	'Strbp',	'Nek6',	'Arhgap15',	'Lypd6b',	'Rbm43',	'Tank',	'Fign',	'Scn7a',	'Stk39',	'Tlk1',	'Cdca7',	'Itgav',	'Tfpi',	'Ctnnd1',	'Ypel4',	'Ube2l6',	'Gm9821',	'Chst1',	'Prdm11',	'Ehf',	'Depdc7',	'Prrg4',	'Mpped2',	'Aven',	'Meis2',	'Thbs1',	'Fsip1',	'Bmf',	'Bub1b',	'Ccdc32',	'Ino80',	'Ehd4',	'Eid1',	'Ap4e1',	'Mrps5',	'Gpcpd1',	'Plcb4',	'Foxa2',	'Pygb',	'Id1',	'Bcl2l1',	'Efcab8',	'Bpifb5',	'Chmp4b',	'Nnat',	'Bpi',	'Adig',	'Chd6',	'Ncoa3',	'Sulf2',	'Arfgef2',	'B4galt5',	'Tshz2',	'Pck1',	'Rab22a',	'Zfp831',	'Cables2',	'Clcn5',	'Gripap1',	'Maob',	'Slc9a7',	'44080',	'Xpnpep2',	'Gpc4',	'Hprt',	'Abcd1',	'Prkx',	'Arhgef9',	'Amer1',	'Pja1',	'Ogt',	'Nexmif',	'Gm379',	'Gm6377',	'Pof1b',	'Chm',	'Hnrnph2',	'Gprasp1',	'Bex4',	'Tceal8',	'Rnf128',	'Huwe1',	'Spin2c',	'Eif1ax',	'Gemin8',	'Msl3',	'Mrps28',	'Tpd52',	'Zbtb10',	'Car13',	'Tnik',	'Mecom',	'Actl6a',	'Gm11549',	'Bbs12',	'Spry1',	'Pgrmc2',	'P2ry13',	'Mbnl1',	'Lekr1',	'Rapgef2',	'Gucy1b1',	'Rbm46',	'Dchs2',	'Fhdc1',	'Arfip1',	'Tigd4',	'Fam160a1',	'Etv3',	'Mrpl24',	'Rhbg',	'Atp8b2',	'Vps45',	'Mtmr11',	'Acp6',	'Reg4',	'Hmgcs2',	'Ptgfrn',	'Igsf3',	'Casq2',	'Syt6',	'Olfml3',	'Magi3',	'Wnt2b',	'Vav3',	'S1pr1',	'Slc44a3',	'Abca4',	'Gclm',	'Bcar3',	'1810037I17Rik',	'Camk2d',	'Rrh',	'Dkk2',	'Bdh2',	'Slc39a8',	'Bank1',	'Mttp',	'4930579F01Rik',	'Bmpr1b',	'Pdlim5',	'Ssx2ip',	'Miga1',	'Zzz3',	'Wls',	'Fam110b',	'Asph',	'Ccne2',	'Tmem64',	'Gpr63',	'Bach2',	'Unc13b',	'Atp8b5',	'Glipr2',	'Gne',	'Fbxo10',	'Tmod1',	'Trmo',	'Nans',	'Tgfbr1',	'Tal2',	'Klf4',	'D630039A03Rik',	'Susd1',	'Slc46a2',	'Slc31a2',	'Rgs3',	'Col27a1',	'Pappa',	'Tlr4',	'Lurap1l',	'Mpdz',	'Nfib',	'Plin2',	'Dennd4c',	'Fggy',	'Ttc22',	'Lrrc42',	
                  'Foxd2',	'Ptprf',	'Foxo6',	'Kcnq4',	'Mfsd2a',	'Ak2',	'C77080',	'Bsdc1',	'Epb41',	'Trim63',	'Srrm1',	'Rcan3',	'Ifnlr1',	'Rap1gap',	'Szrd1',	'Dnajc16',	'Pdpn',	'Dhrs3',	'Lzic',	'Car6',	'Acot7',	'B230104I21Rik',	'Mmel1',	'Fam213b',	'Morn1',	'Cdk14',	'Dmtf1',	'Sema3e',	'Srpk2',	'Shh',	'Garem2',	'Tbc1d14',	'C1qtnf7',	'Cc2d2a',	'Qdpr',	'Slit2',	'Ppargc1a',	'Stim2',	'Pcdh7',	'Klf3',	'N4bp2',	'Rbm47',	'Limch1',	'Slain2',	'Cwh43',	'Rasl11b',	'Thegl',	'Cxcl15',	'Parm1',	'Scarb2',	'Shroom3',	'Ccni',	'Gpat3',	'Arhgap24',	'Ptpn13',	'Cdc7',	'Tgfbr3',	'Fgfrl1',	'Crybb3',	'Ssh1',	'Nos1',	'Med13l',	'Atp2a2',	'Tmem120b',	'Tmem132d',	'Auts2',	'Gtf2i',	'Sh2b2',	'Cux1',	'Myl10',	'Prkar1b',	'Mad1l1',	'Slc29a4',	'Tnrc18',	'Rbak',	'Lmtk2',	'Bhlha15',	'Alox5ap',	'Tex26',	'Fry',	'Pds5b',	'Stard13',	'Ppp1r9a',	'Asb4',	'Cftr',	'Fam3c',	'Lmod2',	'Tspan33',	'Cep41',	'Mkln1',	'Lrguk',	'Bpgm',	'Agbl3',	'Dgki',	'Ephb6',	'Gsdme',	'Nfe2l3',	'Adcyap1r1',	'Thnsl2',	'Sema4f',	'Rtkn',	'Tet3',	'Ankrd53',	'Tgfa',	'Antxr1',	'Plxna1',	'Slc6a6',	'Prickle2',	'Kbtbd8',	'Mitf',	'Foxp1',	'Eif4e3',	'Shq1',	'Rad18',	'Setd5',	'Atp2b2',	'Vgll4',	'Cand2',	'Wnt5b',	'C1s2',	'Acrbp',	'Iffo1',	'A2ml1',	'Styk1',	'Bcl2l14',	'E330021D16Rik',	'Atf7ip',	'Arhgdib',	'Pik3c2g',	'Plekha5',	'St8sia1',	'Sox5',	'Smco2',	'Klhl42',	'Bicd1',	'Zfp773',	'Dact3',	'Zfp60',	'Zfp568',	'Zfp536',	'Plekhf1',	'Med25',	'Irf3',	'Emp3',	'Spty2d1',	'Dbx1',	'Mkrn3',	'Igf1r',	'Nr2f2',	'Gm10295',	'Fam174b',	'Slco3a1',	'Akap13',	'Isg20',	'Wdr73',	'Mex3b',	'Ctsc',	'Fzd4',	'Prss23',	'Ccdc90b',	'Nars2',	'Uvrag',	'Serpinh1',	'P4ha3',	'Arhgef17',	'Pde2a',	'Olfr571',	'Scube2',	'Adm',	'Galnt18',	'Plekha7',	'Rps15a',	'Dnah3',	'Ndufab1',	'Plk1',	'Tbx6',	'Zfp629',	'Uros',	'Tcerg1l',	'Pwwp2b',	'Tspan32',	'Cdkn1c',	'Shank2',	'Ano1',	'Ccnd1',	'Iyd',	'Plekhg1',	'Vnn3',	'Ctgf',	'Ptprk',	'Marcks',	'Wisp3',	'Traf3ip2',	'Foxo3',	'Afg1l',	'Cd24a',	'Rtn4ip1',	'Prdm1',	'Dcbld1',	'Pln',	'Asf1a',	'Psap',	'Unc5b',	'Pcbd1',	'Tacr2',	'Srgn',	'Nrbf2',	'Arid5b',	'Ccdc6',	'Specc1l',	'Mmp11',	'Zfp280b',	'Dnmt3l',	'AU041133',	'Nfyb',	'Nuak1',	'Chpt1',	'Nr1h4',	'Tmpo',	'Nudt4',	'Atp2b1',	'Poc1b',	'Lin7a',	'Tph2',	'Zfc3h1',	'Frs2',	'Cpsf6',	'Cpm',	'Rap1b',	'Grip1',	'Tbc1d30',	'Slc16a7',	'Shmt2',	'Baz2a',	'Rab20',	'Tfdp1',	'Cdc16',	'Kbtbd11',	'Xkr5',	'Smim19',	'Ppp1r3b',	'Eri1',	'Mtnr1a',	'Tlr3',	'Ccdc110',	'Irf2',	'Ing2',	'Cldn22',	'Sap30',	'Cpe',	'Psd3',	'Pbx4',	'Mmaa',	'Adgrl1',	'Ier2',	'G430095P16Rik',	'Dand5',	'Dnaja2',	'Ces1a',	
                  'Ces1g',	'Ogfod1',	'Pllp',	'Ccdc102a',	'Cfap20',	'Cmtm4',	'Terf2',	'Tmem170',	'Sdr42e1',	'Gins2',	'Rhou',	'Rab4a',	'Sipa1l2',	'Pcnx2',	'Pard3',	'Flnb',	'Kctd6',	'Dlg5',	'Il17rd',	'Hacl1',	'Gm8113',	'Bmp4',	'Cdkn3',	'Prmt5',	'Rnaseh2b',	'Zfp395',	'Trim35',	'Bnip3l',	'Dock5',	'Chmp7',	'Sorbs3',	'Lpar6',	'Siah3',
                  'Cog3',	'Serp2',	'Dnajc15',	'Kctd12',	'Pou4f1',	'Spry2',	'Dct',	'Mbnl2',	'Farp1',	'Gm11168',	'Gm10718',	'Ntm',	'St14',	'Ets1',	'Fez1',	'Scn3b',	'Sorl1',	'Mpzl3',	'Scn2b',	'Sik3',	'Cadm1',	'Zbtb16',	'Ppp2r1b',	'Colca2',	'Tmem266',	'Isl2',	'Peak1',	'1700017B05Rik',	'Stra6',	'Adpgk',	'Lrrc49',	'Tle3',	'Anp32a',	'Coro2b',	'Calml4',	'Pias1',	'Smad6',	'Plekho2',	'Pclaf',	'Tpm1',	'Gtf2a2',	'Adam10',	'Hmgcll1',	'Klhl31',	'Gcm1',	'Gsta2',	'Eef1a1',	'Irak1bp1',	'Ibtk',	'Spsb4',	'Dbr1',	'Slc35g2',	'Pccb',	'Cep63',	'Amotl2',	'Srprb',	'Atp2c1',	'Wdr82',	'Cmtm8',	'Stt3b',	'Tgfbr2',	'Ctdspl',	'Scn10a',	'Lyzl4',	'Snrk',	'Zfp105',	'Limd1',	'Ccr5',	'Kremen1',	'Rtn4',	'Bod1',	'Sh3pxd2b',	'Ubtd2',	'Fbxw11',	'Ranbp17',	'Slit3',	'Rars',	'Wwc1',	'Mat2b',	'Trim7',	'Csf2',	'Ccdc69',	'Galnt10',	'Larp1',	'Btnl10',	'Rai1',	'Zfp286',	'Hs3st3b1',	'Cox10',	'Shisa6',	'Dhrs7c',	'Myh10',	'Pfas',	'Scimp',	'Med31',	'Rap1gap2',	'Ccdc92b',	'Srr',	'Timm22',	'Rab34',	'Lyrm9',	'Rasl10b',	'Car4',	'Tbx2',	'Hsf5',	'Rnf43',	'Kat7',	'B4galnt2',	'Lasp1',	'Csf3',	'Krt25',	'Krt36',	'Plekhh3',	'Ccr10',	'Meox1',	'Cd300lg',	'Mpp2',	'BC030867',	'Ubtf',	'Arhgap27',	'Mettl2',	'Tanc2',	'Cyb561',	'Ern1',	'Kpna2',	'Bptf',	'Pitpnc1',	'Apoh',	'Wipi1',	'Nat9',	'Mgat5b',	'Slc16a3',	'Zfp750',	'Zmynd11',	'Arid4b',	'Inhba',	'Trim27',	'Gpx6',	'Hist1h2bl',	'Hist1h2an',	'Btn1a1',	'Hist1h2bh',	'Hist1h3e',	'Hist1h2bf',	'Hist1h4d',	'Hist1h3a',	'Gpld1',	'Sox4',	'Agtr1a',	'Uqcrfs1',	'Foxq1',	'Gmds',	'Bphl',	'Rreb1',	'Sycp2l',	'Nedd9',	'Edn1',	'Tbc1d7',	'Rnf182',	'Jarid2',	'Mylip',	'Atxn1',	'Nup153',	'Zfp169',	'Gadd45g',	'Diras2',	'Syk',	'Cplx2',	'Ddx46',	'2010111I01Rik',	'Aaed1',	'Gm28041',	'Irx1',	'Glrx',	'Vcan',	'Lhfpl2',	'Zbed3',	'Nsa2',	'Arhgef28',	'Utp15',	'Fcho2',	'Srek1',	'Plk2',	'Snx18',	'Hspb3',	'Arl15',	'Parp8',	'Gm17509',	'Apob',	'Osr1',	'Rock2',	'Cys1',	'Rsad2',	'Tmem18',	'Alkal2',	'Immp2l',	'Lrrn3',	'Stxbp6',	'Mbip',	'Pole2',	'Klhdc1',	'Atp5s',	'Tmx1',	'Frmd6',	'Six4',	'Tmem30b',	'Prkch',	'Syne2',	'Max',	'Dcaf5',	'Map3k9',	'Sipa1l1',	'Rgs6',	'Fcf1',	'Ttll5',	'Ston2',	'Gpr68',	'Ccdc88c',	'Golga5',	'Ppp4r4',	'Dicer1',	'Bdkrb1',	'Hsp90aa1',	'Dnah11',	'Ccdc152',	'Osmr',	'Nadk2',	'Rai14',	'Myo10',	'Ctnnd2',	'Nipal2',	'Zfp706',	'Azin1',	'Abra',	'Eif3h',	'Deptor',	'Mrpl13',	'Zhx2',	'Fer1l6',	'Nsmce2',	'Fam84b',	'Ndrg1',	'Adgrb1',	'Eppk1',	'Hsf1',	'Rbfox2',	'Myh9',	'Sstr3',	'Cdc42ep1',	'Syngr1',	'Chadl',	'Atxn10',	'Gramd4',	'Ano6',	'Arid2',	'Slc38a1',	'Nckap5l',	'Lima1',	'Dip2b',	'Sp1',	'Gpr84',	
                  'Zfp385a',	'Crebbp',	'Ercc4',	'Fopnl',	'Map3k13',	'Igf2bp2',	'St6gal1',	'P3h2',	'Mb21d2',	'Dlg1',	'Mylk',	'Stxbp5l',	'Gpr156',	'Gsk3b',	'Upk1b',	'Abhd10',	'Phldb2',	'Morc1',	'Nsun3',	'4930453N24Rik',	'Robo1',	'Robo2',	'Btg3',	'Adamts1',	'Grik1',	'Mis18a',	'Mx1',	'Tmem181a',	'Dynlt1a',	'Rsph3b',	'Pacrg',	'Agpat4',	'Afdn',	'Zfp820',	'Zfp947',	'Zfp945',	'Traf7',	'Neurl1b',	'Tulp1',	'Rsph1',	'Cryaa',	'Brd2',	'C2',	'Ddr1',	'Ier3',	'Ppp1r10',	'Adgrf5',	'Supt3',	'Dnph1',	'Unc5cl',	'Pp2d1',	'Ptprs',	'Efna5',	'Man2a1',	'Themis3',	'Epb41l3',	'Vit',	'Qpct',	'Cdc42ep3',	'Pkdcc',	'Eml4',	'Haao',	'Lrpprc',	'Svil',	'Ccny',	'Dtna',	'Nrep',	'Reep2',	'Sil1',	'Psd2',	'Hbegf',	'Pcdh1',	'Nr3c1',	'Gm5689',	'Stk32a',	'Snx24',	'Zfp608',	'Tex43',	'Slc27a6',	'Sh3tc2',	'Wdr7',	'Nedd4l',	'Zfp532',	'Oacyl',	'Grp',	'Lman1',	'Rab27b',	'Smad4',	'Mro',	'Gm9925',	'Myo5b',	'Acaa2',	'Zbtb7c',	'Nfatc1',	'Zfp516',	'Cndp1',	'Ppp6r3',	'Kdm2a',	'Mus81',	'Pcnx3',	'Pla2g16',	'Chrm1',	'Fads2',	'Anxa1',	'1110059E24Rik',	'Fam189a2',	'Pum3',	'Slc16a12',	'Hells',	'Ccnj',	'Tm9sf3',	'Frat1',	'Ina',	'Sfr1',	'Gsto1',	'Afap1l2',	'Gfra1',	'Pnlip',	'Pnliprp1',	'Prdx3'))

integrate_epi4 <- AddModuleScore(object = integrate_epi4, features = Pr_less, ctrl = 500, name = "NKXProgLessGenes", search = TRUE, random.seed = 1)
#1 gene not accounted for: 44080
FeaturePlot(object = integrate_epi4, features = 'NKXProgLessGenes1', cols = c('lightgrey','red'))

#save file
saveRDS(integrate_epi4, file = "/Users/drlittle/Desktop/Agg12_Sox9AT1AT2noprolif_mod_score.rds")


AT2ACClean <- list(c('Ncoa2',	'Sema4c',	'Vwa3b',	'Inpp4a',	'Pdcl3',	'Slc39a10',	'Mfsd6',	'1700019D03Rik',	'Dnah7a',	'Satb2',	'Gm973',	'Fam117b',	'Abi2',	'Fzd5',	'Kansl1l',	'43894',	'Igfbp5',	'Epha4',	'Acsl3',	'Serpine2',	'Kcnj13',	'Asb1',	'Bok',	'Gli2',	'Cfap221',	'C1ql2',	'Slc35f5',	'Gpr39',	'Lypd1',	'Ccnt2',	'Cxcr4',	'Cd55b',	'Rab7b',	'Cntn2',	'Plekha6',	'Snrpe',	'Prelp',	'Kdm5b',	'Syt2',	'Ube2t',	'Csrp1',	'Zfp281',	'Nek7',	'Kcnt2',	'Ivns1abp',	'Swt1',	'Edem3',	'Shcbp1l',	'Soat1',	'Blzf1',	'Dpt',	'Lrrc52',	'Cfap126',	'Klhdc9',	'Vangl2',	'Atp1a2',	'Igsf9',	'Tagln2',	'Cfap45',	'Rgs7',	'Opn3',	'Efcab2',	'Acbd3',	'Tmem63a',	'Lpgat1',	'Traf5',	'Kcnh1',	'Irf6',	'A130010J15Rik',	'Proser2',	'Celf2',	'Notch1',	'Ier5l',	'Cstad',	'1700001O22Rik',	'Ass1',	'Pomt1',	'Lmx1b',	'Mvb12b',	'Cntrl',	'Lypd6b',	'Rnd3',	'Tnfaip6',	'Fmnl2',	'Rprm',	'Tank',	'Csrnp3',	'Galnt3',	'Sp5',	'Erich2',	'Cybrd1',	'Cdca7',	'Osbpl6',	'Zfp385b',	'Prdx6b',	'4833423E24Rik',	'Ptprj',	'C1qtnf4',	'Trp53i11',	'Ehf',	'D430041D05Rik',	'Rcn1',	'Mpped2',	'Grem1',	'Gjd2',	'Dph6',	'Eif2ak4',	'Dll4',	'Sqor',	'Slc12a1',	'Mal',	'Acoxl',	'Bcl2l11',	'Tmem87b',	'Cenpb',	'Smox',	'Bmp2',	'Plcb4',	'Foxa2',	'Fam110a',	'Id1',	'Dnmt3b',	'Bpifb1',	'Bpifb5',	'Epb41l1',	'Bpi',	'Dhx35',	'Top1',	'Ptprt',	'Fitm2',	'Sdc4',	'Eya2',	'Cse1l',	'Slc9a8',	'Ptpn1',	'Cdh4',	'Taf4',	'Shroom4',	'Clcn5',	'Mid1ip1',	'H2al1a',	'Rgn',	'Dock11',	'Apln',	'Mbnl3',	'Gpc4',	'Ints6l',	'Prrg3',	'Cnga2',	'Gm7173',	'5430427O19Rik',	'Amer1',	'Yipf6',	'Pja1',	'Cxcr3',	'Nexmif',	'Gprasp2',	'Bex2',	'Atg4a',	'Smpx',	'Rps6ka3',	'Tceanc',	'Egfl6',	'Prps2',	'Zfhx4',	'Il7',	'Zbtb10',	'Zfp704',	'Fabp12',	'Lrrcc1',	'Dnajc5b',	'Fndc3b',	'Tmem212',	'Pld1',	'Skil',	'Ccdc39',	'Atp11b',	'Pgrmc2',	'Noct',	'Mgst2',	'Maml3',	'Frem2',	'Mab21l1',	'Sucnr1',	'Ccnl1',	'Sptssb',	'Map9',	'Tigd4',	'Tmem154',	'Lrba',	'Arhgef2',	'Pmvk',	'Gatad2b',	'Selenbp2',	'Hist2h2ac',	'Hist2h3b',	'Hist2h2bb',	'Chd1l',	'Vtcn1',	'Ptgfrn',	'Slc16a1',
                     'Tmigd3',	'Pifo',	'Cept1',	'Cyb561d1',	'Aknad1',	'Fam102b',	'Snx7',	'Dpyd',	'Camk2d',	'Enpep',	'Elovl6',	'Etnppl',	'Dkk2',	'Arhgef38',	'Slc39a8',	'Ddit4l',	'Bmpr1b',	'Syde2',	'Adgrl2',	'Adgrl4',	'St6galnac3',	'Nsmaf',	'Car8',	'Chd7',	'Gem',	'Epha7',	'Bach2',	'Rragd',	'Zfp292',	'Ifnk',	'1700009N14Rik',	'Nfx1',	'Ubap2',	'Dnajb5',	'Tgfbr1',	'Tex10',	'Acnat1',	'Rnf20',	'Rad23b',	'D630039A03Rik',	'Dnajc25',	'Susd1',	'Slc31a2',	'Zfp618',	'Cntln',	'Adamtsl1',	'Mllt3',	'Tek',	'Jun',	'Cyp2j6',	'Cyp2j9',	'Nfia',	'Angptl3',	'Alg6',	'Ror1',	'Cachd1',	'Jak1',	'Oma1',	'Dab1',	'Bend5',	'Pdzk1ip1',	'Kif2c',	'Rnf220',	'Slc6a9',	'Hivep3',	'Gm8439',	'Kcnq4',	'Pou3f1',	'Utp11',	'Zscan20',	'Yars',	'Marcksl1',	'Pef1',	'Hcrtr1',	'Nkain1',	'Fgr',	'Fuca1',	'Hnrnpr',	'Lactbl1',	'Rap1gap',	'Vwa5b1',	'Tmco4',	'Tas1r2',	'Pax7',	'Atp13a2',	'Spata21',	'Arhgef19',	'Epha2',	'Spen',	'Fhad1',	'Cenps',	'Kif1b',	'Prdm16',	'Pex10',	'Sri',	'Sema3c',	'Cd36',	'Gnai1',	'Lhfpl3',	'Srpk2',	'Insig1',	'Cnpy1',	'Shh',	'Lmbr1',	'Letm1',	'Cpeb2',	'Cc2d2a',	'Kcnip4',	'Ppargc1a',	'Lgi2',	'Sepsecs',	'Pi4k2b',	'Arap2',	'0610040J01Rik',	'Klf3',	'Klhl5',	'Wdr19',	'9130230L23Rik',	'Atp8a1',	'Kctd8',	'Corin',	'Cnga1',	'Ociad2',	'Rasl11b',	'Fip1l1',	'Rufy3',	'Gc',	'Npffr2',	'Adamts3',	'Cox18',	'Cxcl15',	'Areg',	'Btc',	'Parm1',	'Bmp2k',	'Prkg2',	'Arhgap24',	'Ptpn13',	'Gbp6',	'Lrrc8c',	'Lrcol1',	'Ulk1',	'Sez6l',	'Myo18b',	'Sgsm1',	'1700069L16Rik',	'Acacb',	'Wsb2',	'2410131K14Rik',	'Rhof',	'Arl6ip4',	'Gm10382',	'Tmem132c',	'Slc15a4',	'Tmem132d',	'Galnt17',	'Cldn4',	'Cldn3',	'Mlxipl',	'Col26a1',	'Ift22',	'Fis1',	'Ephb4',	'Zan',	'Cyp3a13',	'Usp12',	'Gm42791',	'Medag',	'Tex26',	'Fry',	'Pdk4',	'Glcci1',	'Ppp1r3a',	'Met',	'Capza2',	'Wnt2',	'Tspan12',	'Aass',	'Cadps2',	'Opn1sw',	'Strip2',	'Tmem209',	'Cep41',	'Copg2',	'Chchd3',	'Chrm2',	'Tmem213',	'Tmem178b',	'Tmem139',	'Gsdme',	'Cycs',	'Nfe2l3',	'Snx10',	'Hibadh',	'Znrf2',	'Gars',	'Fam13a',	'Eif2ak3',	'Tgoln1',	'Hk2',	'Alms1',
                     'C87436',	'Pcbp1',	'Prokr1',	'Iqsec1',	'Hdac11',	'Magi1',	'Mitf',	'Foxp1',	'Rybp',	'Oxtr',	'Ogg1',	'Vgll4',	'Tsen2',	'43898',	'Dcp1b',	'Cecr2',	'Mical3',	'Lrrc23',	'Cracr2a',	'Mansc1',	'Emp1',	'Dera',	'Mgst1',	'Pik3c2g',	'Sox5',	'Tmc4',	'Zfp541',	'Zc3h4',	'Hif3a',	'Ccdc61',	'Irf2bp1',	'Irgq',	'Ceacam1',	'Cyp2s1',	'Cyp2f2',	'Hnrnpl',	'Catsperg1',	'Krtdap',	'Zfp507',	'Zfp536',	'Uri1',	'Vstm2b',	'Klk14',	'Klk13',	'1700008O03Rik',	'Abcc6',	'Nell1',	'Cers3',	'Arrdc4',	'Slco3a1',	'Zscan2',	'Pde8a',	'Mesd',	'Gm2115',	'Tmem135',	'Eed',	'Nars2',	'Gab2',	'Pak1',	'Myo7a',	'Omp',	'Tsku',	'Lrrc32',	'Thap12',	'Serpinh1',	'Lmo1',	'Ascl3',	'Wee1',	'Adm',	'Parva',	'Rassf10',	'Arntl',	'Spon1',	'Xylt1',	'Syt17',	'Gga2',	'BC017158',	'Rgs10',	'Nsmce4a',	'Oat',	'D7Ertd443e',	'Mgmt',	'Tcerg1l',	'Utf1',	'Eps8l2',	'Kcnq1',	'Tnfrsf23',	'Sash1',	'Plagl1',	'Cited2',	'Ect2l',	'Olig3',	'Ptprk',	'Fyn',	'Mfsd4b1',	'Ddo',	'Foxo3',	'Prdm1',	'Pkib',	'P4ha1',	'Hkdc1',	'Gm10118',	'Jmjd1c',	'Nrbf2',	'Arid5b',	'Mrln',	'Gstt2',	'Slc19a1',	'Mbd3',	'Tjp3',	'Lta4h',	'Plxnc1',	'Socs2',	'Ube2n',	'Gm5426',	'Btg1',	'Atp2b1',	'Kitl',	'Slc6a15',	'Acss3',	'Myf5',	'Phlda1',	'Kcnc2',	'Yeats4',	'Cpsf6',	'Mdm1',	'Grip1',	'Msrb3',	'D930020B18Rik',	'Srgap1',	'Avpr1a',	'Pip4k2c',	'Nxph4',	'Smarcc2',	'Efnb2',	'Col4a2',	'Grtp1',	'Kbtbd11',	'Myom2',	'Angpt2',	'Zmat4',	'Htra4',	'Tacc1',	'Unc5d',	'Tex15',	'Eri1',	'Cldn23',	'Prag1',	'Sorbs2',	'Galnt7',	'Sh3rf1',	'Cbr4',	'Tll1',	'Lzts1',	'Cers1',	'Tmem59l',	'Jund',	'Arhgap10',	'1700011L22Rik',	'Mmaa',	'Smad1',	'Gypa',	'Inpp4b',	'Il15',	'Gipc1',	'Adgrl1',	'Nfix',	'Irx5',	'Ces1a',	'Ces1d',	'Adgrg3',	'Fhod1',	'Smpd3',	'Gm10073',	'Cdh1',	'Tango6',	'Zfhx3',	'Hp',	'Tmem170',	'Adamts18',	'Nudt7',	'Wwox',	'Cmip',	'Tldc1',	'Cotl1',	'Gins2',	'Irf8',	'Banp',	'Urb2',	'Disc1',	'Map3k21',	'Kcnk1',	'Nrp1',	'Fam107a',	'Gm49355',	'Nid2',	'Dupd1',	'Lrmda',	'Dlg5',	'Zmiz1',	'Hesx1',	'Tmem110',	'Syt15',	'Gm15217',	'Slc35f4',	'Efs',
                     'Nfatc4',	'Sdr39u1',	'Zmym5',	'Gja3',	'Eef1akmt1',	'Cab39l',	'Cdadc1',	'Spata13',	'Neil2',	'Zfp395',	'Ccdc25',	'Ebf2',	'Adam7',	'Adam28',	'Slc39a14',	'Sftpc',	'Fgf17',	'Itm2b',	'Ccdc122',	'Dnajc15',	'Rgcc',	'Pcdh9',	'Slain1',	'Gpc5',	'Dct',	'Cldn10',	'Dock9',	'Birc3',	'Gm16485',	'Sesn3',	'Eepd1',	'Ntm',	'Zbtb44',	'Fez1',	'Zfp202',	'Hspa8',	'Nectin1',	'Cbl',	'Fxyd6',	'Pafah1b2',	'Sik3',	'Sik2',	'Gm32742',	'Colca2',	'Arhgap20',	'4930550C14Rik',	'Cib2',	'Wdr61',	'Lingo1',	'Csk',	'Neo1',	'Nr2e3',	'Uaca',	'Smad3',	'Smad6',	'Parp16',	'Dapk2',	'Usp3',	'Tpm1',	'C2cd4b',	'Rora',	'Myo1e',	'Cgnl1',	'Arpp19',	'Hcrtr2',	'Fam83b',	'Tinag',	'Ibtk',	'Bcl2a1d',	'Bcl2a1b',	'Atr',	'Clstn2',	'Rbp2',	'Cpne4',	'Mapkapk3',	'Gnat1',	'Bcl2a1c',	'Crtap',	'Cmtm8',	'Entpd3',	'Cck',	'Topaz1',	'Ykt6',	'Ppia',	'Purb',	'Nacad',	'Abca13',	'Ikzf1',	'Fignl1',	'Kcnmb1',	'Ccnjl',	'Cby3',	'Csf2',	'Anxa6',	'Zfp692',	'Srebf1',	'Dhrs7b',	'Mfap4',	'Fam83g',	'Ncor1',	'Hs3st3b1',	'Hs3st3a1',	'Myh13',	'Cfap52',	'Ntn1',	'Cd68',	'Slc16a13',	'Alox12',	'Atp2a3',	'Emc6',	'Rph3al',	'Efcab5',	'Pipox',	'Rab34',	'Ksr1',	'Myo1d',	'Spaca3',	'Ccl8',	'Pex12',	'Hnf1b',	'Pctp',	'Skap1',	'Epop',	'Cisd3',	'B230217C12Rik',	'Grb7',	'Csf3',	'Igfbp4',	'Acly',	'Arl4d',	'Plcd3',	'Mrc2',	'Cacng4',	'Kcnj2',	'Sox9',	'Sstr2',	'Gprc5c',	'Galk1',	'Rnf157',	'Mxra7',	'Sec14l1',	'Gm11733',	'Afmid',	'Rnf213',	'Baiap2',	'Fscn2',	'Fasn',	'Cd7',	'Calml3',	'Calm4',	'Akr1c13',	'Akr1c19',	'Zmynd11',	'Gli3',	'Stard3nl',	'Trim27',	'Hist1h2ah',	'Hist1h2ag',	'Sox4',	'Bphl',	'Ofcc1',	'Tfap2a',	'Hivep1',	'Edn1',	'Phactr1',	'Jarid2',	'Dtnbp1',	'Atxn1',	'Spin1',	'Syk',	'Ror2',	'Rgs14',	'Tgfbi',	'Gm45623',	'Gas1',	'Dapk1',	'Fbp1',	'Ptch1',	'Irx2',	'Slc12a7',	'Vcan',	'Ap3b1',	'Zbed3',	'Enc1',	'Gm10260',	'Cartpt',	'Rnf180',	'Smim15',	'Elovl7',	'Il6st',	'Mcidas',	'Nnt',
                     'Kif3c',	'Dtnb',	'Rhob',	'Fam84a',	'Trib2',	'Greb1',	'Rock2',	'Hpcal1',	'Mboat2',	'Id2',	'Lamb1',	'Prkar2b',	'Atxn7l1',	'Etv1',	'Lrrn3',	'Egln3',	'Snx6',	'Psma6',	'Brms1l',	'Nkx2-1',	'Mdga2',	'Gm9887',	'Pygl',	'Tmx1',	'Frmd6',	'Six1',	'Prkch',	'Esr2',	'Plekhg3',	'Rab15',	'Rad51b',	'Srsf5',	'Rgs6',	'Acot1',	'Isca2',	'Tgfb3',	'Calm1',	'Ccdc88c',	'Slc24a4',	'Clmn',	'Vrk1',	'Bcl11b',	'Setd3',	'Dlk1',	'Rtl1',	'Dio3',	'Rapgef5',	'Osmr',	'Rai14',	'Cdh6',	'Ctnnd2',	'Cmbl',	'Cthrc1',	'Abra',	'Ext1',	'Mal2',	'Taf2',	'Col14a1',	'Lrrc6',	'Ly6g',	'Gpihbp1',	'C1qtnf6',	'Gcat',	'Mgat3',	'Tnrc6b',	'Atxn10',	'Trmu',	'Celsr1',	'Gramd4',	'Twf1',	'Nell2',	'Tmem106c',	'Col2a1',	'Tuba1c',	'Acvr1b',	'Eif4b',	'Ppp1r1a',	'Crebbp',	'4930562C15Rik',	'Emp2',	'Rmi2',	'Efcab1',	'Cldn5',	'Dgkg',	'St6gal1',	'Opa1',	'Hes1',	'Bdh1',	'Zdhhc19',	'Muc4',	'Smbd1',	'Slc35a5',	'Phldb2',	'Cblb',	'Alcam',	'Nfkbiz',	'Stx19',	'Cadm2',	'Samsn1',	'Grik1',	'Cldn8',	'Sod1',	'Mis18a',	'Mrps6',	'Runx1',	'Setd4',	'Ripk4',	'Fndc1',	'Dact2',	'Smoc2',	'Thbs2',	'Noxo1',	'Hs3st6',	'Cacna1h',	'Neurl1b',	'Lemd2',	'Hmga1',	'Fance',	'Clps',	'Cdkn1a',	'Cpne5',	'Rsph1',	'Slc37a1',	'H2-DMb2',	'Ddah2',	'Mpig6b',	'Prrc2a',	'Trim40',	'Adgrf5',	'Tomm6',	'Mdfi',	'Gm20219',	'Man2a1',	'Tgif1',	'Crim1',	'Atl2',	'Galm',	'Tmem178',	'Pkdcc',	'Mta3',	'Abcg8',	'Camkmt',	'Six2',	'Adcyap1',	'Greb1l',	'Celf4',	'Tslp',	'Stard4',	'Gm10549',	'Srp19',	'Fam13b',	'Tmem173',	'E230025N22Rik',	'Pcdh1',	'Fgf1',	'Stk32a',	'Dtwd2',	'Zfp608',	'Gramd3',	'Lmnb1',	'43893',	'Slc12a2',	'Slc27a6',	'Arhgef37',	'Grpel2',	'Afap1l1',	'Fbxo38',	'Pmaip1',	'Seh1l',	'Rab27b',	'Mbd2',	'Rpl17',	'Smad2',	'St8sia5',	'Setbp1',	'Kcng2',	'Ctdp1',	'Timm21',	'Znhit2',	'Rps6ka4',	'Mark2',	'Fam111a',	'Tle4',	'Gnaq',	'Aldh1a7',	'Fam189a2',	'Rfx3',	'Plpp6',	'Ranbp6',	'Sgms1',	'Tnks2',	'Myof',	'Ffar4',	'Blnk',	'Tm9sf3',	'Frat1',	'Nkx2-3',	'Entpd7',	'Scd2',	'Scd4',	'Sfxn2',	'Neurl1a',	'Rpl13a-ps1',	'Add3',	'Vwa2',	'Pnlip'))
#not accounted  43894, 43898, 43893
integrate <- AddModuleScore(object = integrate, features = AT2ACClean, ctrl = 500, name = "NKXAT2AcquiredCleanGenes", search = TRUE, random.seed = 1)
FeaturePlot(object = integrate, features = 'NKXAT2AcquiredCleanGenes1', split.by = "ID", cols = c('lightgrey','red'))
FeaturePlot(object = integrate, features = 'NKXAT2AcquiredCleanGenes1', cols = c('lightgrey','red'))


AT2RClean <- list(c('Alkal1',	'Arfgef1',	'Ogfrl1',	'Sdhaf4',	'Fam168b',	'Uggt1',	'Il1r1',	'Il18rap',	'Mrps9',	'AI597479',	'Ankrd44',	'Aox3',	'Ndufb3',	'Cflar',	'Tmem237',	'Erbb4',	'Arpc2',	'Tmbim1',	'Ihh',	'Resp18',	'Slc19a3',	'Gm47969',	'Cops7b',	'Dgkd',	'Arl4c',	'Agap1',	'Mlph',	'Prlh',	'Rbm44',	'Erfe',	'Sned1',	'Tsn',	'Inhbb',	'Epb41l5',	'Marco',	'Tmem163',	'Cd55',	'Gm29427',	'Pik3c2b',	'Ppp1r15b',	'Lax1',	'Chil1',	'Lmod1',	'Shisa4',	'Kif21b',	'Nmnat2',	'Lamc1',	'Tor3a',	'Tnfsf4',	'Myoc',	'Nme7',	'Atp1b1',	'Uap1',	'Apoa2',	'Fcer1g',	'F11r',	'Alyref2',	'Vsig8',	'Fh1',	'Kmo',	'Sdccag8',	'Sccpdh',	'Lbr',	'Degs1',	'Capn2',	'1700056E22Rik',	'Rab3gap2',	'Kcnk2',	'Nenf',	'Hsd11b1',	'Prpf18',	'Usp6nl',	'Stam',	'Hnmt',	'Il1f9',	'Rnf224',	'Fam69b',	'Lcn4',	'Stkld1',	'Wdr5',	'Col5a1',	'Tsc1',	'Cercam',	'Ptpa',	'Usp20',	'Abl1',	'Prrc2b',	'Lcn2',	'Ptges2',	'St6galnac4',	'Cdk9',	'Zbtb43',	'Ptgs1',	'Rc3h2',	'Crb2',	'Tanc1',	'Wdsub1',	'Lrp2',	'Gm26558',	'Dcaf17',	'Pdk1',	'Scrn3',	'Nfe2l2',	'Ttc30b',	'Ccdc141',	'Zfp804a',	'Spi1',	'Creb3l1',	'Pex16',	'Elf5',	'Bub1b',	'Ino80',	'Pla2g4f',	'Wdr76',	'Sord',	'Shf',	'Galk2',	'Hdc',	'Gabpb1',	'Usp8',	'Usp50',	'Stard7',	'Slc20a1',	'Ap5s1',	'Rassf2',	'Jag1',	'Snx5',	'Kiz',	'Cd93',	'Gzf1',	'Apmap',	'Acss1',	'Entpd6',	'Slc52a3',	'H13',	'Cox4i2',	'Map1lc3a',	'Uqcc1',	'Phf20',	'Myl9',	'Ctnnbl1',	'Actr5',	'Fam83d',	'Tox2',	'Oser1',	'Gdap1l1',	'Slpi',	'Elmo2',	'Ocstamp',	'Ube2v1',	'Tmem189',	'Atp9a',	'Tshz2',	'Cass4',	'Ctsz',	'Cdh26',	'Lsm14b',	'Cables2',	'Col20a1',	'Polr3k',	'Gata1',	'Cask',	'Fundc1',	'Slc9a7',	'Mtm1',	'Pnck',	'Tbl1x',	'Las1l',	'Igbp1',	'Sat1',	'Ofd1',	'Trappc2',	'Car13',	'Mtfr1',	'Nceh1',	'Tnik',	'Actl6a',	'Spry1',	'Jade1',	'Rab33b',	'Nhlrc3',	'Il12a',	'Ift80',	'Fnip2',	'Fam160a1',	'Mef2d',	'Sema4a',	'Pygo2',	'Ubap2l',	'S100a10',	'Selenbp1',	'Rfx5',	'Pi4kb',	'Zfp687',	'Tmod4',	'Ciart',	'Sv2a',	'Bcl9',	'Sec22b',	'Notch2',	'Cd101',	'Mab21l3',	'Slc22a15',	'I830077J02Rik',	'Ovgp1',	'Chia1',	'Chil4',	'Dennd2d',	'Amigo1',	'Sort1',	'Mybphl',	'Vav3',	'Slc44a3',	'Ppa2',	'H2afz',	'Nexn',	'Sdcbp',	'2610301B20Rik',	'Plekhf2',	'Pdp1',	'Ttpa',	'Ddx58',	'Aptx',	'B4galt1',	'AI464131',	'Olfr70',	'Glipr2',	'Gne',	'Pax5',	'Fbxo10',	'Shb',	'Tdrd7',	'Ncbp1',	'Stx17',	'Msantd3',	'Tmeff1',	'Ppp3r2',	'Fsd1l',	'Klf4',	'Ctnnal1',	'Ptgr1',	'Fkbp15',	'Bspry',	'Hdhd3',	'Orm1',	'Orm2',	'Plin2',	'Eqtn',	'Mysm1',	'Dio1',	'Podn',	'Mknk1',	'Akr1a1',	'Eif2b3',	'Ccdc24',	'Artn',	'Szt2',	'Med8',	'Elovl1',	'Cldn19',	'Rimkla',	'Edn2',	'Slfnl1',	'Smap2',	'Zmpste24',	'Inpp5b',	'Yrdc',	'Zc3h12a',	'Thrap3',	'A3galt2',	'Trim62',	'Ak2',	'Gm10300',	'Ythdf2',	'Med18',	'Ptafr',	'Xkr8',	'Stx12',	'Sfn',	'Selenon',	'Man1c1',	'Ephb2',	'Cela3b',	'Alpl',	'Hp1bp3',	'Ubr4',	'Necap2',	'Plekhm2',	'Efhd2',	'Dhrs3',	'Tnfrsf1b',	'Nppa',	'Pgd',	'Lzic',	'Gpr157',	'Rere',	'Zbtb48',	'Tas1r1',	'Gpr153',	'Megf6',	'Prkcz',
                    'Nadk',	'Perm1',	'Gm8773',	'Steap2',	'Tmem243',	'Dmtf1',	'Gnat3',	'Atg9b',	'Crygn',	'Prkag2',	'Paxip1',	'Il6',	'Adgrf3',	'Khk',	'Mrpl33',	'Fosl2',	'Pisd',	'Mxd4',	'Sh3bp2',	'Rgs12',	'Hgfac',	'Psapl1',	'Tada2b',	'Tbc1d14',	'Hs3st1',	'Slc34a2',	'Smim20',	'Stim2',	'Klb',	'Nipal1',	'Slain2',	'Lrrc66',	'Hopx',	'Grsf1',	'Cxcl1',	'Scarb2',	'Ccng2',	'Antxr2',	'Bmp3',	'4930524J08Rik',	'Cds1',	'Gm17660',	'Rpl5',	'Cplx1',	'Pole',	'Tpst2',	'Grk3',	'Tchp',	'Git2',	'Cit',	'Prkab1',	'Taok3',	'Tesc',	'Aldh2',	'Sh2b3',	'Rad9b',	'Camkk2',	'Kdm2b',	'Orai1',	'Vps37b',	'Rflna',	'Ncor2',	'Tmem248',	'Limk1',	'Bcl7b',	'Upk3b',	'Gigyf1',	'Gnb2',	'Agfg2',	'Dnaaf5',	'Micall2',	'Chst12',	'Papolb',	'Usp42',	'Bri3',	'Trrap',	'Zfp655',	'N4bp2l1',	'Pds5b',	'Kl',	'Ppp1r9a',	'Pon1',	'Pon3',	'Slc25a13',	'Ica1',	'Mdfic',	'Flnc',	'Akr1b3',	'Akr1b8',	'Agbl3',	'Kdm7a',	'Ndufb2',	'Agk',	'Gstk1',	'Zfp746',	'Fam221a',	'Osbpl3',	'Skap2',	'Aqp1',	'Rnf103',	'Reep1',	'Sftpb',	'Ggcx',	'Suclg1',	'Eva1a',	'Cd207',	'Smyd5',	'Tprkb',	'Asprv1',	'D6Ertd527e',	'Arhgap25',	'Rpn1',	'Txnrd3',	'Klf15',	'Slc41a3',	'1700123L14Rik',	'Eif4e3',	'Edem1',	'Mtmr14',	'Cpne9',	'Vhl',	'Atg7',	'Csgalnact2',	'Ret',	'Atp6v1e1',	'Usp18',	'Prmt8',	'Tspan11',	'Pzp',	'Borcs5',	'Ddx47',	'Rerg',	'Pyroxd1',	'Abcc9',	'Bhlhe41',	'Stk38l',	'Far2',	'Lair1',	'Isoc2b',	'Zfp787',	'Zfp418',	'C5ar2',	'Sae1',	'Gpr4',	'Gemin7',	'Relb',	'Tomm40',	'Ceacam16',	'Xrcc1',	'Zfp575',	'Cxcl17',	'Bckdha',	'Numbl',	'Map3k10',	'9530053A07Rik',	'Rps16',	'Lrfn3',	'Gpi1',	'Cebpa',	'Vsig10l',	'Napsa',	'Akt1s1',	'Ntn5',	'Rpl18',	'Hps5',	'Cyfip1',	'Klf13',	'Fan1',	'Selenos',	'Chd2',	'St8sia2',	'Klhl25',	'Mrps11',	'Rlbp1',	'Mesp1',	'Mesp2',	'Rccd1',	'Prc1',	'Crtc3',	'Iqgap1',	'Fah',	'Fzd4',	'Thrsp',	'Mogat2',	'Relt',	'P2ry2',	'Clpb',	'Folr1',	'Hpx',	'Ilk',	'Eif3f',	'Rpl27a',	'Mrvi1',	'Arl6ip1',	'Ccp110',	'Gprc5b',	'Dcun1d3',	'Lyrm1',	'Ubfd1',	'Lcmt1',	'Il21r',	'Gsg1l',	'Il27',	'Nupr1',	'Sult1a1',	'Ccdc189',	'Stx4a',	'Cox6a2',	'Slc5a2',	'Plpp4',	'Fgfr2',	'Ate1',	'Plekha1',	'Fam53b',	'Abraxas2',	'Edrf1',	'Dhx32',	'Dock1',	'Mapk1ip1',	'Zfp941',	'Rnh1',	'Cracr2b',	'Chid1',	'Slc22a18',	'Osbpl5',	'Shank2',	'Ppfia1',	'Fadd',	'Zc3h12d',	'Stx11',	'Sf3b5',	'Aig1',	'Map3k5',	'Pde7b',	'Echdc1',	'Tspyl4',	'Frk',	'Ccdc162',	'Ostm1',	'Gm9803',	'Sh3rf3',	'Oit3',	'Hk1',	'Tet1',	'Tmem26',	'Ank3',	'Ccdc6',	'Gnaz',	'Bcr',	'Dip2a',	'Pcnt',	'Col6a2',	'Col6a1',	'Trappc10',	'Kiss1r',	'Polr2e',	'Gpx4',	'Dazap1',	'Uqcr11',	'Rexo1',	'Cactin',	'Washc4',	'1500009L16Rik',	'Timp3',	'Gnptab',	'Utp20',	'Uhrf1bp1l',	'Cdk17',	'Elk3',	'Cradd',	'Gm33543',	'Poc1b',	'Myrfl',	'Lyz2',	'Lyz1',	'Cpm',	'Slc35e3',	'Gns',	'Rassf3',	'Tbk1',	'Zbtb39',	'Rbms2',	'Camsap3',	'Prr36',	'Tnfsf13b',	'Myo16',	'Naxd',	'Tex29',	'Atp11a',	'F7',	'Cul4a',	'Lamp1',	'Dcun1d2',	'Tmem255b',	'Gas6',	'Rasa3',	'Fbxo25',	'Cln8',	'Atp7b',	'Tpte',
                    'Bag4',	'Pomk',	'Dlc1',	'Slc7a2',	'Mtus1',	'F11',	'Cyp4v3',	'Casp3',	'Slc18a1',	'Sugp1',	'Crtc1',	'Uba52',	'Ell',	'Isyna1',	'Myo9b',	'Nr2f6',	'Tpm4',	'Mcm5',	'Prkaca',	'Lyl1',	'G430095P16Rik',	'Gpt2',	'Abcc12',	'Brd7',	'Nkd1',	'Sall1',	'Chd9',	'Rbl2',	'Ces1h',	'Slc12a3',	'Herpud1',	'Nlrc5',	'Slc38a7',	'B3gnt9',	'Hsf4',	'Plekhg4',	'Kctd19',	'Hsd11b2',	'Tsnaxip1',	'Cyb5b',	'Psmd7',	'Txnl4b',	'Cmtr2',	'Ldhd',	'Hsd17b2',	'Mlycd',	'Osgin1',	'Gse1',	'Fbxo31',	'Jph3',	'Tcf25',	'Afg3l1',	'Rhou',	'2310022B05Rik',	'Trim67',	'2810004N23Rik',	'Gnpat',	'Sprtn',	'Egln1',	'Irf2bp2',	'Tomm20',	'Flnb',	'Rpp14',	'Saysd1',	'Kcnma1',	'Rft1',	'Itih4',	'Itih1',	'Stab1',	'Tnnc1',	'Fam25c',	'Sh2d4b',	'Sftpa1',	'Psmc6',	'Ddhd1',	'Gmfb',	'Fbxo34',	'Tmem260',	'Klhl33',	'Lrp10',	'Gm29776',	'Myh7',	'Fitm1',	'Tgm1',	'Rabggta',	'Ltb4r2',	'Atp12a',	'Gm4491',	'Cryl1',	'Phf11d',	'Ctsb',	'Kif13b',	'Scara5',	'Pbk',	'Nudt18',	'Lrrc63',	'Lacc1',	'Tnfsf11',	'Spry2',	'Dzip1',	'Gpr18',	'Pcca',	'Fut4',	'1700012B09Rik',	'Zfp317',	'Spc24',	'Angptl8',	'Zfp809',	'Snx19',	'St14',	'Aplp2',	'Acrv1',	'Tbrg1',	'Scn3b',	'Sorl1',	'Usp2',	'Cxcr5',	'Tmprss4',	'BC049352',	'Fxyd2',	'Cadm1',	'Nnmt',	'Zbtb16',	'Plet1',	'1110032A03Rik',	'Fdx1',	'Tnfaip8l3',	'1700017B05Rik',	'Tmem202',	'Pkm',	'Pias1',	'Snapc5',	'Rab11a',	'Slc24a1',	'Rasl12',	'Spg21',	'Snx1',	'Pygo1',	'Pigb',	'Unc13c',	'Fam214a',	'Eef1a1',	'Mthfs',	'Tbc1d2b',	'Zbtb38',	'Pxylp1',	'Msl2',	'Anapc13',	'Rab6b',	'Twf2',	'Alas1',	'Rad54l2',	'Rbm15b',	'Rassf1',	'Ifrd2',	'Amigo3',	'1700102P08Rik',	'Plxnb1',	'Cspg5',	'Scap',	'Setd2',	'Prss46',	'Golga4',	'Ctdspl',	'Acaa1a',	'Exog',	'Ctnnb1',	'Trak1',	'Hhatl',	'Higd1a',	'Ano10',	'Abhd5',	'Tmem42',	'Zdhhc3',	'Exosc7',	'Clec3b',	'Cdcp1',	'Limd1',	'Drg1',	'Morc2a',	'Dusp18',	'Slc35e4',	'Ascc2',	'Nf2',	'Gas2l1',	'Emid1',	'Xbp1',	'Camk2b',	'Igfbp3',	'Vps54',	'Bod1',	'Cpeb4',	'Ubtd2',	'Fbxw11',	'Ttc1',	'Adra1b',	'Adam19',	'Nipal4',	'Ifi47',	'Mgat1',	'Tbc1d9b',	'Col23a1',	'Rmnd5b',	'Fstl4',	'44082',	'Gm17334',	'Acsl6',	'Gpx3',	'Trim11',	'Pld6',	'Flcn',	'Rasd1',	'Llgl1',	'Top3a',	'Natd1',	'Map2k3',	'B9d1',	'Epn2',	'Akap10',	'Adora2b',	'Tekt3',	'Shisa6',	'Gas7',	'Dhrs7c',	'Myh10',	'Dnah2',	'Tnfsf13',	'Chrnb1',	'Slc16a11',	'Mis12',	'Pimreg',	'Tekt1',	'Spns2',	'P2rx1',	'Haspin',	'P2rx5',	'Cluh',	'Mnt',	'Tsr1',	'Srr',	'Hic1',	'Rtn4rl1',	'Rpa1',	'Inpp5k',	'Doc2b',	'Rflnb',	'Dbil5',	'Coro6',	'Proca1',	'Unc119',	'Poldip2',	'Tnfaip1',	'Lyrm9',	'Rhbdl3',	'Cdk5r1',	'Rffl',	'1700020L24Rik',	'Wfdc18',	'Synrg',	'Acaca',	'Aatf',	'Rnft1',	'Tubd1',	'Vmp1',	'Mtmr4',	'Hsf5',	'Trim25',	'Nog',	'Tob1',	'Abcc3',	'Abi3',	'Snx11',	'Sp2',	'Osbpl7',	'Tbkbp1',	'Socs7',	'Srcin1',	'Ppp1r1b',	'Stard3',	'Pgap3',	'Nt5c3b',	'Ttc25',	'Cnp',	'Stat3',	'Coa3',	'Tmem101',	'Asb16',	'Dcakd',	'Hexim2',	'Map3k14',	'Arhgap27',	'Gh',	'Pitpnc1',	'Cacng1',	'Apoh',	'Cdc42ep4',	'Grb2',
                    'Tmem94',	'Wbp2',	'Fbf1',	'Sphk1',	'St6galnac2',	'Pgs1',	'Tbc1d16',	'Endov',	'Chmp6',	'Faap100',	'Cbr2',	'Hmga1b',	'Ccdc57',	'Ptchd3',	'Larp4b',	'Heatr1',
                    'Nid1',	'Ly86',	'Rreb1',	'Bmp6',	'Txndc5',	'Stmnd1',	'Fam8a1',	'Phf2',	'Nfil3',	'Faf2',	'Ddx41',	'Smad5',	'Agtpbp1',	'Golm1',	'Aaed1',	'Rps23',	'Ssbp2',	'Hexb',	'Pik3r1',	'Nln',	'Ipo11',	'Depdc1b',	'Rab3c',	'Cdc20b',	'Gzmk',	'Itga1',	'Gm17509',	'Hmgcs1',	'Klhl29',	'Hs1bp3',	'Fam49a',	'Ntsr2',	'Klf11',	'Adi1',	'Tmem18',	'Hdac9',	'Ahr',	'Ifrd1',	'Cfl2',	'Nemf',	'Nin',	'Psma3',	'Slc38a6',	'Gphb5',	'Plekhh1',	'Zfp36l1',	'Ccdc177',	'Dcaf4',	'Abcd4',	'Ltbp2',	'Tmed10',	'Fos',	'Cipc',	'Pomt2',	'Ahsa1',	'Ism2',	'Nrde2',	'Cpsf2',	'Chga',	'Itpk1',	'Syne3',	'Mok',	'Ankrd9',	'Amn',	'Siva1',	'Fbxo4',	'Egflam',	'Wdr70',	'Prlr',	'Npr3',	'Pop1',	'Ubr5',	'Dcstamp',	'Deptor',	'Ndufb9',	'Trib1',	'Fam49b',	'Tg',	'Ndrg1',	'Ly6d',	'2010109I03Rik',	'Ly6e',	'Cyc1',	'Scx',	'Gpt',	'Zfp647',	'Rbfox2',	'Slc16a8',	'Tmem184b',	'Csnk1e',	'Ddx17',	'Tomm22',	'Rpl3',	'Sgsm3',	'Aco2',	'Desi1',	'Mcat',	'Prr5',	'Fbln1',	'Wnt7b',	'Ppara',	'Fam19a5',	'Brd1',	'Pim3',	'Trabd',	'Plxnb2',	'Sbf1',	'Pdzrn4',	'Pced1b',	'Pfkm',	'Lmbr1l',	'Fam186b',	'Slc11a2',	'Gm5475',	'Pou6f1',	'Cela1',	'Nr4a1',	'Krt80',	'Krt88',	'Tns2',	'Amhr2',	'Nfe2',	'Dnase1',	'Mgrn1',	'Nudt16l1',	'Rogdi',	'Gm5767',	'Clec16a',	'Litaf',	'Txndc11',	'Snx29',	'Cpped1',	'Ercc4',	'Mkl2',	'Ntan1',	'Fgd4',	'Slc7a4',	'Dgcr8',	'Comt',	'Ap2m1',	'2510009E07Rik',	'Ahsg',	'Fetub',	'Rfc4',	'Lpp',	'Tprg',	'Il1rap',	'Slc51a',	'Snx4',	'Hcls1',	'Gsk3b',	'Nr1i2',	'Tmem39a',	'Qtrt2',	'Btla',	'Retnlb',	'Tmem45a2',	'Cmss1',	'Crybg3',	'Arl6',	'Btg3',	'Map3k7cl',	'Bach1',	'Tmem50b',	'Cryzl1',	'Chaf1b',	'Dyrk1a',	'Kcnj15',	'Erg',	'Wrb',	'Ezr',	'Slc22a3',	'Slc22a1',	'Mrpl18',	'Acat2',	'Prss22',	'Kctd5',	'Rnps1',	'Pkd1',	'Dusp1',	'Bak1',	'Ip6k3',	'Taf11',	'Slc26a8',	'Stk38',	'BC004004',	'Fgd2',	'Pim1',	'Pknox1',	'Cryaa',	'Sik1',	'Angptl4',	'H2-Ob',	'H2-Aa',	'Gm20481',	'Ddx39b',	'Vegfa',	'Polr1c',	'Ptk7',	'Mea1',	'Cnpy3',	'Mocs1',	'Gm7334',	'Tbc1d5',	'Kat2b',	'Dpp9',	'Safb2',	'Rfx2',	'C3',	'Ptprm',	'Togaram2',	'Xdh',	'Memo1',	'Cdc42ep3',	'Rmdn2',	'Morn2',	'Map4k3',	'Kcng3',	'Thada',	'Dync2li1',	'Gm10309',	'Epas1',	'Mcfd2',	'Ttc7',	'Calm2',	'Crem',	'Svil',	'Cables1',	'Npc1',	'Cabyr',	'Trappc8',	'Rnf125',	'Sap130',	'Gpr17',	'Gypc',	'Nrep',	'Pkd2l2',	'Sil1',	'Pcdhgc5',	'Diaph1',	'Gnpda1',	'Rbm27',	'Trim36',	'Prr16',	'Prdm6',	'Dctn4',	'Synpo',	'Cd74',	'Hmgxb3',	'Pde6a',	'Ppargc1b',	'Impa2',	'Ptpn2',	'Ldlrad4',	'Fam210a',	'Mex3c',	'Acaa2',	'Dym',	'Ctif',	'Zfp236',	'Tshz1',	'Zadh2',	'Zfp407',	'Rttn',	'Lrp5',	'Gstp3',	'Cabp2',	'Pcx',	'Mrpl11',	'Rab1b',	'AI837181',	'Ap5b1',	'Atg2a',	'Rasgrp2',	'Otub1',	'Fads1',	'Gna14',	'Rfk',	'Anxa1',	'Klf9',	'Fam122a',	'Kcnv2',	'Rcl1',	'Il33',	'Acta2',	'Lipa',	'Kif20b',	'Slc35g1',	'Sorbs1',	'Ccnj',	'Ankrd2',	'Pyroxd2',	'Cwf19l1',	'Pkd2l1',	'Wnt8b',	'Fgf8',	'Cuedc2',	'Actr1a',	'Cnnm2',	'Xpnpep1',	'Dusp5',	'Shoc2',	'Shtn1',	'Eif3a'))
#not accounted
integrate <- AddModuleScore(object = integrate, features = AT2RClean, ctrl = 500, name = "NKXAT2RetainedCleanGenes", search = TRUE, random.seed = 1)
FeaturePlot(object = integrate, features = 'NKXAT2RetainedCleanGenes1', split.by = "ID", cols = c('lightgrey','red'))
FeaturePlot(object = integrate, features = 'NKXAT2RetainedCleanGenes1', cols = c('lightgrey','red'))


AT1ACClean <- list(c('Rb1cc1',	'Prex2',	'Tram1',	'Eya1',	'Rdh10',	'Efhc1',	'Gsta3',	'Smap1',	'Uggt1',	'Arid5a',	'Kansl3',	'Lman2l',	'Mgat4a',	'2010300C02Rik',	'Eif5b',	'Npas2',	'Tbc1d8',	'Il1r1',	'AI597479',	'Stat1',	'Stk17b',	'Gtf3c3',	'Rftn2',	'Als2cr12',	'Tmem237',	'Fzd7',	'Nbeal1',	'Fzd5',	'Mreg',	'Gpbar1',	'Stk36',	'Kcne4',	'Ap1s3',	'Fam124b',	'Nyap2',	'Rhbdd1',	'Col4a3',	'Agfg1',	'Kcnj13',	'Hjurp',	'Rbm44',	'Traf3ip1',	'Twist2',	'Mterf4',	'Clasp1',	'Tmem37',	'Fcmr',	'Tmcc2',	'Dstyk',	'Mybph',	'Adora1',	'Ipo9',	'Zfp281',	'Nek7',	'Ivns1abp',	'Axdnd1',	'Tor3a',	'Prdx6',	'Dnm3',	'Fmo6',	'Mael',	'Mgst3',	'F11r',	'Tagln2',	'Grem2',	'Stum',	'Parp1',	'Susd4',	'Spata17',	'G0s2',	'Cdc123',	'Gm13199',	'Il2ra',	'Mindy3',	'Nebl',	'Pip4k2a',	'Gad2',	'Ehmt1',	'Uap1l1',	'Sohlh1',	'Camsap1',	'Setx',	'Med27',	'Sptan1',	'Wdr34',	'Dolpp1',	'Aif1l',	'Rpl12',	'Cntrl',	'Rab14',	'Gsn',	'Gpr21',	'Dennd1a',	'Epc2',	'Ly75',	'Itgb6',	'Fign',	'Fastkd1',	'Gm26558',	'Rapgef4',	'Wipf1',	'Prkra',	'Ccdc141',	'Sestd1',	'Zfp385b',	'Nup160',	'Psmc3',	'F2',	'Ambra1',	'Chst1',	'Prdm11',	'Cd82',	'Alkbh3',	'Ttc17',	'Nat10',	'Bdnf',	'Lin7c',	'Aven',	'Srp14',	'Inafm2',	'Bahd1',	'Haus2',	'Slc24a5',	'Dut',	'Cops2',	'Ciao1',	'Ttl',	'Slc20a1',	'Stk35',	'Tgm3',	'Cds2',	'Kif16b',	'Rin2',	'Fam110a',	'Slc52a3',	'Cox4i2',	'Ccm2l',	'Hck',	'Tm9sf4',	'Cdk5rap1',	'Snta1',	'Chmp4b',	'Gss',	'Trpc4ap',	'Uqcc1',	'Phf20',	'Cnbd2',	'Soga1',	'Samhd1',	'Manbal',	'Ctnnbl1',	'Arhgap40',	'Pkig',	'Pigt',	'Dnttip1',	'Snx21',	'Slc2a10',	'Cse1l',	'Slc9a8',	'Ptpn1',	'Pard6b',	'Zfp64',	'Bmp7',	'Ss18l1',	'Tcfl5',	'Ythdf1',	'Zgpat',	'Lime1',	'Myt1',	'Plp2',	'Ftsj1',	'Il13ra1',	'Pgrmc1',	'Elf4',	'Gm648',	'Fhl1',	'Haus7',	'Slc6a8',	'Pls3',	'Tbl1x',	'Efnb1',	'Itgb1bp2',	'Phka1',	'Morf4l2',	'Prps1',	'Tsc22d3',	'Phka2',	'Cdkl5',	'1700064H15Rik',	'Lrrc31',	'Kcnmb3',	'Dcun1d1',	'Anxa5',	'Nudt6',	'Noct',	'Lhfp',	'Commd2',	'Eif2a',	'Gpr171',	'P2ry14',	'Dhx36',	'Tmem144',	'Tmem131l',	'Trim2',	'Gatb',	'Rusc1',	'Il6ra',	'Rps27',	'Dennd4b',	'Gatad2b',	'S100a14',	'Them4',	'Cgn',	'Anxa9',	'Setdb1',	'Plekho1',	'Adam30',	'Zfp697',	'Nr1h5',	'Capza1',	'Tmem167b',	'Stxbp3',	'Slc44a3',	'Aimp1',	'Dapp1',	'Tspan5',	'Ttll7',	'Gipc2',	'Acadm',	'Tnni3k',	'Tgs1',	'Rab2a',	'Plekhf2',	'Gm12353',	'Slc7a13',	'Casp8ap2',	'Pnrc1',	'Akirin2',	'B4galt1',	'Spink4',	'Dcaf12',	'Rusc2',	'Creb3',	'Frmpd1',	'Slc25a51',	'Tmod1',	'Tstd2',	'Coro2a',	'Ppp3r2',	'Abca1',	'Fam206a',	'Inip',	'Rgs3',	'Tle1',	'Ttc39b',	'Dennd4c',	'Ror1',	'Prkaa2',	'Mrpl37',	'Cc2d1b',	'Osbpl9',	'Ttc39a',	'Cyp4b1',	'Lurap1',	'Plk3',	'Dmap1',	'Artn',	'Kdm4a',	'Szt2',	'Zfp691',	'Edn2',	'Foxo6',	'Ppt1',	'Inpp5b',	'Gnl2',	'Lsm10',	'Eva1b',	'Sh3d21',	'Thrap3',	'Map7d1',	'Trappc3',	'Ago3',	'Yars',	'Pef1',	'Dnajc8',	'Sfn',	'Man1c1',	'Clic4',	'Ifnlr1',	'Cdc42',	'Alpl',	'Eif4g3',	'Hp1bp3',	'Otud3',	'Tmco4',	'Capzb',	'Ubr4',	'Spen',	'Dhrs3',	'Angptl7',
                     'Casz1',	'Kif1b',	'Dnajc11',	'Klhl21',	'Zbtb48',	'Ski',	'Faap20',	'Mib2',	'Tnfrsf4',	'Fam133b',	'Gatad1',	'Cyp51',	'Sri',	'Slc25a40',	'Rsbn1l',	'Gsap',	'Rheb',	'E130116L18Rik',	'Rbm33',	'Shh',	'Supt7l',	'Fosl2',	'Yes1',	'Mxd4',	'Cfap99',	'Sh3bp2',	'Msantd1',	'Psapl1',	'D5Ertd579e',	'Bloc1s4',	'Wfs1',	'Dhx15',	'Chrna9',	'Limch1',	'Slain2',	'Srd5a3',	'Hopx',	'Anxa3',	'Prkg2',	'Gpat3',	'Cds1',	'Ptpn13',	'Slc10a6',	'Gak',	'Tmem175',	'4930522L14Rik',	'Ddx51',	'Kctd10',	'Sppl3',	'Gm10401',	'Taok3',	'Tesc',	'Fbxw8',	'Oas3',	'Ptpn11',	'Trafd1',	'Adam1b',	'Sh2b3',	'Myl2',	'Ccdc63',	'Orai1',	'Rhof',	'Clip1',	'Hcar2',	'Abcb9',	'2810006K23Rik',	'Ccdc92',	'Ncor2',	'Sfswap',	'Tpst1',	'Rfc2',	'Lat2',	'Tmem120a',	'Ywhag',	'Orai2',	'Sh2b2',	'Azgp1',	'Zkscan1',	'Zscan21',	'Gm5294',	'Sun1',	'Gper1',	'Elfn1',	'Mad1l1',	'Gna12',	'Rnf216',	'Rac1',	'Fam220a',	'Lmtk2',	'Usp12',	'Slc7a1',	'Fry',	'N4bp2l1',	'Pds5b',	'Cav2',	'Cav1',	'Tspan12',	'Lmod2',	'Wasl',	'Lrrc4',	'Rbm28',	'Akr1b3',	'Cnot4',	'Chrm2',	'Klrg2',	'Clec2l',	'Parp12',	'Braf',	'Nfe2l3',	'Snx10',	'Mindy4',	'Chmp3',	'Sftpb',	'Usp39',	'Mat2a',	'Cyp26b1',	'Pradc1',	'Tgfa',	'Pcyox1',	'Anxa4',	'Aak1',	'Antxr1',	'Aplf',	'H1fx',	'Mgll',	'Tpra1',	'Chchd6',	'Hdac11',	'Tmem43',	'Foxp1',	'Itpr1',	'Srgap3',	'Cidec',	'Il17re',	'Rassf4',	'Rasgef1a',	'Ret',	'Bms1',	'Cacna1c',	'Lrtm2',	'Ninj2',	'Slc25a18',	'Foxj2',	'Cd9',	'Tspan9',	'Gabarapl1',	'Crebl2',	'Atf7ip',	'Aebp2',	'Recql',	'Ldhb',	'Sbk3',	'Zfp579',	'Zscan22',	'Nop53',	'Ehd2',	'Bbc3',	'Sae1',	'Fosb',	'Ercc1',	'Clasrp',	'Plaur',	'Capn12',	'Eif3k',	'2200002D01Rik',	'Spint2',	'Dpf1',	'Ap2a1',	'Grin2d',	'Hps5',	'Klf13',	'Selenos',	'Akap13',	'Ntrk3',	'Aen',	'Isg20',	'Mfge8',	'Idh2',	'Zscan2',	'Slc28a1',	'Mex3b',	'Abhd17c',	'Ankrd42',	'Ints4',	'Map6',	'Rps3',	'Arrb1',	'Tpbgl',	'Slco2b1',	'Spcs2',	'Lipt2',	'Ucp3',	'Arap1',	'Rhog',	'Cavin3',	'Rpl27a',	'St5',	'Wee1',	'Swap70',	'Ampd3',	'Rnf141',	'Mrvi1',	'Gprc5b',	'Scnn1g',	'Cog7',	'Slc5a11',	'4930451I11Rik',	'Hirip3',	'Tmem219',	'Zfp747',	'Fbrs',	'Slc5a2',	'Bag3',	'Wdr11',	'Stk32c',	'Inpp5a',	'Rnh1',	'Irf7',	'Muc6',	'Mob2',	'Cttn',	'Tpcn2',	'Vip',	'Sash1',	'Stxbp5',	'Stx11',	'Aig1',	'Heca',	'Arfgef3',	'Map3k5',	'Map7',	'1700020N01Rik',	'Sgk1',	'Slc2a12',	'Moxd1',	'Themis',	'Prdm1',	'Popdc3',	'Mcm9',	'Gja1',	'P4ha1',	'Pla2g12b',	'Oit3',	'Mcu',	'Micu1',	'Spock2',	'Pcbd1',	'Lrrc20',	'H2afy2',	'Col13a1',	'Tacr2',	'Ado',	'Arid5b',	'A930033H14Rik',	'Ccdc6',	'Susd2',	'Derl3',	'Smarcb1',	'2610028H24Rik',	'Ybey',	'Lss',	'Ube2g2',	'Trappc10',	'2610008E11Rik',	'Cnn2',	'Gpx4',	'Sbno2',	'Ap3d1',	'Celf5',	'Appl2',	'Timp3',	'Ttc41',	'Apaf1',	'Nr2c1',	'Tmcc3',	'Cradd',	'Poc1b',	'Rap1b',	'Baz2a',	'Cs',	'Arhgef18',	'Pex11g',	'Fam155a',	'Irs2',	'Col4a2',	'Cars2',	'Arhgef10',	'Kat6a',	'Adam32',	'Plpp5',	'Plpbp',	'Mak16',	'Smim18',	'Gm19410',	'Mtus1',	'Triml2',	'Sorbs2',	'Ccdc110',	'Snx25',
                     'Cfap97',	'Aga',	'Spcs3',	'Wdr17',	'Tmem192',	'Slc18a1',	'Gatad2a',	'Mau2',	'Lsm4',	'Haus8',	'Zfp961',	'Gm10282',	'Klf2',	'Large1',	'Arhgap10',	'Smad1',	'Usp38',	'Tbc1d9',	'Rfx1',	'Cacna1a',	'Junb',	'Siah1a',	'Nkd1',	'Nod2',	'Irx6',	'Ccl22',	'Usb1',	'B3gnt9',	'Elmo3',	'Ranbp10',	'Slc12a4',	'Terf2',	'Atxn1l',	'Ap1g1',	'Tat',	'Mtss1l',	'Bco1',	'Gan',	'Cmip',	'Cdh13',	'Osgin1',	'Mbtps1',	'Map1lc3b',	'Zfpm1',	'Piezo1',	'Pabpn1l',	'Vps9d1',	'Spire2',	'Acta1',	'2310022B05Rik',	'Itgb1',	'Abhd6',	'Pxk',	'Ndst2',	'Vcl',	'Vdac2',	'Anxa11',	'Dennd6a',	'Fam208a',	'Prkcd',	'Ccser2',	'Samd4',	'Ktn1',	'Olfr750',	'Mphosph8',	'Zdhhc20',	'Gm6904',	'Kcnrg',	'Blk',	'Fam167a',	'Pinx1',	'Pnoc',	'Dpysl2',	'Dock5',	'9930012K11Rik',	'Sorbs3',	'Gm21750',	'Esd',	'Lrch1',	'Rgcc',	'Dis3',	'Scel',	'Rbm26',	'Rap2a',	'Ipo5',	'Stk24',	'Dync2h1',	'Cep57',	'Panx1',	'Icam1',	'Ilf3',	'Tmed1',	'Kank2',	'Herpud2',	'Ntm',	'Cdon',	'Nectin1',	'Hspb2',	'Rdx',	'Zc3h12c',	'Kdelc2',	'4930550C14Rik',	'Acsbg1',	'Dnaja4',	'Tspan3',	'Mpi',	'Sema7a',	'Hcn4',	'Gramd2',	'Paqr5',	'Itga11',	'Aagab',	'Smad3',	'Hacd3',	'Zfp609',	'Ppib',	'Dapk2',	'Rnf111',	'Myzap',	'Tex9',	'Nedd4',	'Mapk6',	'Lysmd2',	'Elovl5',	'Fbxo9',	'Nt5e',	'Plod2',	'Gk5',	'Slc25a36',	'Gm1123',	'Pik3cb',	'9630041A04Rik',	'Acad11',	'Poc1a',	'Hemk1',	'Hyal2',	'Sema3b',	'Sema3f',	'Uba7',	'Bsn',	'Rhoa',	'Nckipsd',	'Cdc25a',	'Setd2',	'Rtp3',	'Pdcd6ip',	'Cmtm8',	'Itga9',	'Exog',	'Xirp1',	'Cck',	'Snrk',	'Limk2',	'Sec14l3',	'Lif',	'Pgam2',	'Polm',	'Etaa1',	'B3gnt2',	'Zrsr1',	'Pus10',	'Rel',	'Cfap36',	'Bod1',	'Rhbdf1',	'Nprl3',	'Fgf18',	'Rnf145',	'Rnf130',	'Jade2',	'Cdkn2aipnl',	'Skp1a',	'Hspa4',	'44082',	'Slc22a4',	'P4ha2',	'Cdc42se2',	'Tnip1',	'Fat2',	'Sap30l',	'Sh3bp5l',	'Arf1',	'Mprip',	'Pld6',	'Tmem11',	'Prpsap2',	'Adora2b',	'Fbxw10',	'Tekt3',	'F930015N05Rik',	'Zkscan6',	'Ndel1',	'Rnf222',	'Dlg4',	'Asgr1',	'Pimreg',	'Gm43951',	'Cyb5d2',	'Srr',	'Slc43a2',	'Inpp5k',	'Mrm3',	'Timm22',	'Gm10277',	'Dhrs13',	'Crlf3',	'Atad5',	'Myo1d',	'Tada2a',	'Ypel2',	'Mrps23',	'Tob1',	'Wfikkn2',	'Gm11541',	'Tac4',	'Slc35b1',	'Phb',	'Osbpl7',	'Kpnb1',	'Mrpl45',	'Stard3',	'Tcap',	'Krt31',	'Krt15',	'Krt42',	'Stat5b',	'Stat3',	'Cavin1',	'Mapt',	'Itgb3',	'Ddx5',	'Cep95',	'Smurf2',	'Helz',	'Apoh',	'Slc39a11',	'Unc13d',	'Mettl23',	'Mgat5b',	'Tmc8',	'Pgs1',	'Cep295nl',	'Lgals3bp',	'Ndufaf8',	'Fasn',	'Gdi2',	'Zmynd11',	'Lyst',	'Acot13',	'Aldh5a1',	'Serpinb6b',	'Serpinb9',	'Pxdc1',	'Rpp40',	'Gfod1',	'Jarid2',	'Dtnbp1',	'Stmnd1',	'Kif13a',	'Rnf144b',	'Zfp169',	'A830005F24Rik',	'Fam120a',	'Card19',	'Bicd2',	'Tes3-ps',	'Ogn',	'Nxnl2',	'Nsd1',	'Prelid1',	'H2afy',	'Ercc6l2',	'Habp4',	'Slc6a19',	'Ahrr',	'Fam172a',	'Ckmt2',	'Gm20379',	'Ap3b1',	'Pde8b',	'Aggf1',	'F2r',	'Iqgap2',	'Utp15',	'Mrps27',	'Cd180',	'Ndufaf2',	'Fkbp1b',	'Matn3',	'Ttc32',	'2410004P03Rik',
                     'Cpsf3',	'Cys1',	'Id2',	'Sntg2',	'Agr3',	'Prkd1',	'Strn3',	'Nfkbia',	'Mbip',	'Sav1',	'Max',	'Mpp5',	'Plekhd1',	'Sipa1l1',	'Ptgr2',	'Zfp410',	'Fcf1',	'Eif2b2',	'Zc2hc1c',	'Tmed10',	'Fos',	'Jdp2',	'Batf',	'Ift43',	'Gpatch2l',	'Ston2',	'Dglucy',	'Rin3',	'Chga',	'Itpk1',	'Papola',	'Ccnk',	'Ccdc85c',	'Hhipl1',	'Degs2',	'Slc25a47',	'Wdr25',	'Rtl1',	'Ppp2r5c',	'Hsp90aa1',	'Mok',	'Amn',	'Mark3',	'Adssl1',	'Cep170b',	'Ahnak2',	'Crip1',	'Wdr60',	'Ccdc152',	'Rictor',	'1810049J17Rik',	'Zfp622',	'Sema5a',	'Cpq',	'Stk3',	'Snx31',	'Fzd6',	'Ebag9',	'Mal2',	'Col14a1',	'Fam91a1',	'Nsmce2',	'Trappc9',	'Ago2',	'Tigd5',	'Fam83h',	'Dgat1',	'Cpsf1',	'Rbfox2',	'Apol9b',	'Myh9',	'Tmprss6',	'Baiap2l2',	'Pla2g6',	'Maff',	'Cbx6',	'Sgsm3',	'Tef',	'Srebf2',	'Poldip3',	'Brd1',	'Creld2',	'Pim3',	'Cpne8',	'Prickle1',	'Scaf11',	'Endou',	'Ccdc65',	'Fkbp11',	'Aqp2',	'Aqp5',	'Dip2b',	'Atf1',	'6030408B16Rik',	'Krt80',	'Krt7',	'Krt79',	'Sp1',	'Amhr2',	'Calcoco1',	'Adcy9',	'Pam16',	'Cdip1',	'Anks3',	'Ppl',	'Sec14l5',	'Tekt5',	'Socs1',	'Serpind1',	'Aifm3',	'Slc7a4',	'Comt',	'Txnrd2',	'Hira',	'Klhl24',	'Map6d1',	'Igf2bp2',	'Xxylt1',	'Muc13',	'Umps',	'Hacd2',	'Popdc2',	'Zbtb20',	'Boc',	'Ccdc80',	'Abhd10',	'Cblb',	'Tmem30c',	'4930453N24Rik',	'Zfp654',	'Usp25',	'Btg3',	'Usp16',	'Il10rb',	'Ifngr2',	'Kcne2',	'Hlcs',	'Kcnj15',	'Mx1',	'Prdm15',	'Zdhhc14',	'AC122413.2',	'Prr18',	'Tcte2',	'Fam120b',	'Flywch1',	'Kctd5',	'Pdpk1',	'E4f1',	'Mlst8',	'Slc9a3r2',	'Mslnl',	'Gm8225',	'Ergic1',	'Atp6v0e',	'Crebrf',	'Lemd2',	'Nudt3',	'Taf11',	'Fgd2',	'Zfp763',	'Rab11b',	'Ager',	'Rnf5',	'Tnxb',	'Cdsn',	'2310061I04Rik',	'Rnf39',	'Ppp1r11',	'H2-M3',	'Ptchd4',	'Adgrf5',	'Runx2',	'Nfkbie',	'Capn11',	'Tmem63b',	'1600014C23Rik',	'Ptk7',	'Taf8',	'Rftn1',	'Ndufa11',	'Fbxl17',	'Vapa',	'Ralbp1',	'Ddx11',	'Rab12',	'Tgif1',	'Myl12b',	'Myl12a',	'Vit',	'Strn',	'Cdkl4',	'Abcg5',	'Prkce',	'Gm10309',	'Stpg4',	'Epcam',	'Foxn2',	'Cables1',	'Riok3',	'Npc1',	'Kctd1',	'Dsg2',	'Rprd1a',	'Polr2d',	'Lims2',	'Map3k2',	'Kdm3b',	'Reep2',	'Etf1',	'Pfdn1',	'Diaph1',	'Sh3rf2',	'Dcp2',	'Mcc',	'Ticam2',	'Arl14epl',	'Lox',	'Csnk1g3',	'Ndst1',	'Rps14',	'Arhgef37',	'Afap1l1',	'Adrb2',	'Gm9949',	'Zfp532',	'Lman1',	'Impa2',	'Elac1',	'Mro',	'Acaa2',	'Zbtb7c',	'Adnp2',	'Pqlc1',	'Mbp',	'Zadh2',	'Ppp6r3',	'Kmt5b',	'Aldh3b1',	'Lrfn4',	'Tsga10ip',	'Scyl1',	'Frmd8',	'Rps6ka4',	'Cox8a',	'Lgals12',	'Incenp',	'Fth1',	'Cyb561a3',	'Zp1',	'Osbp',	'Psat1',	'Carnmt1',	'Zfand5',	'Abhd17b',	'Klf9',	'Gm9493',	'Fam189a2',	'Pip5k1b',	'Dock8',	'Kcnv2',	'Jak2',	'Ric1',	'Dkk1',	'Acta2',	'Ifit3',	'Ifit1bl1',	'Ifit1',	'Kif20b',	'Rpp30',	'Pdlim1',	'Sorbs1',	'Mms19',	'Zfyve27',	'Golga7b',	'Scd2',	'Hif1an',	'Kcnip2',	'Trim8',	'Wbp1l',	'Cnnm2',	'Slk',	'Cfap43',	'Gsto1',	'Gsto2',	'Trub1'))

integrate <- AddModuleScore(object = integrate, features = AT1ACClean, ctrl = 500, name = "NKXAT1AcquiredCleanGenes", search = TRUE, random.seed = 1)
FeaturePlot(object = integrate, features = 'NKXAT1AcquiredCleanGenes1', split.by = "ID", cols = c('lightgrey','red'))
FeaturePlot(object = integrate, features = 'NKXAT1AcquiredCleanGenes1', cols = c('lightgrey','red'))


AT1RClean <- list(c('Vcpip1',	'Ncoa2',	'Lactb2',	'Phf3',	'Rpl31',	'Nck2',	'1500015O10Rik',	'Wdr75',	'Slc39a10',	'Tmeff2',	'Gls',	'Nab1',	'Dnah7a',	'Hecw2',	'Ankrd44',	'Plcl1',	'Sgo2a',	'Cdk15',	'Klf7',	'Plekhm3',	'Map2',	'Kansl1l',	'Erbb4',	'Ikzf2',	'Igfbp2',	'Tmbim1',	'Epha4',	'Scg2',	'Spp2',	'Glrp1',	'Arl4c',	'Sh3bp4',	'Gbx2',	'Lrrfip1',	'Klhl30',	'Pdcd1',	'Pam',	'Phlpp1',	'Epb41l5',	'Sctr',	'Nckap5',	'Mapkapk2',	'Slc45a3',	'Lemd1',	'Klhdc8a',	'Cntn2',	'Lax1',	'Prelp',	'Syt2',	'Lmod1',	'Csrp1',	'Trove2',	'Rgs2',	'Rgs13',	'Pla2g4a',	'Hmcn1',	'Ncf2',	'Nmnat2',	'Xpr1',	'Tnfsf18',	'4930558K02Rik',	'Pigc',	'Tada1',	'Uck2',	'Fcgr3',	'Tstd1',	'Atp1a4',	'Fmn2',	'Pld5',	'Akt3',	'Zbtb18',	'Lbr',	'Srp9',	'Capn8',	'Lyplal1',	'Esrrg',	'Kctd3',	'Kcnk2',	'Rps6kc1',	'Plxna2',	'Bend7',	'Phyh',	'Usp6nl',	'Trdmt1',	'Hacd1',	'H2al2a',	'A930004D18Rik',	'Otud1',	'Abi1',	'Spopl',	'Lcn4',	'Brd3',	'Ncs1',	'Qrfp',	'Swi5',	'Stxbp1',	'Slc2a8',	'Garnl3',	'Pbx3',	'Phf19',	'Crb2',	'Zeb2',	'Lypd6',	'Arl5a',	'Prpf40a',	'Acvr1',	'Wdsub1',	'Rbms1',	'Tank',	'Dpp4',	'Gcg',	'Grb14',	'Gpr155',	'Mtx2',	'Ttn',	'Zfp804a',	'Zdhhc5',	'Arfgap2',	'1110051M20Rik',	'Pex16',	'Syt13',	'Ext2',	'Traf6',	'Prr5l',	'Abtb2',	'BC052040',	'Stard7',	'Bcl2l11',	'Gm14025',	'Rnf24',	'Lrrn4',	'Btbd3',	'Ism1',	'Flrt3',	'Insm1',	'Foxa2',	'Cd93',	'Snph',	'Id1',	'Bcl2l1',	'Asxl1',	'Eif2s2',	'Myl9',	'Mroh8',	'Src',	'D630003M21Rik',	'Ppp1r16b',	'Chd6',	'Ptprt',	'Wfdc13',	'Wfdc3',	'Eya2',	'Zmynd8',	'Ptgis',	'Snai1',	'Ripor3',	'Bcas1',	'Fam210b',	'Fam209',	'Lsm14b',	'Lama5',	'Rbbp8nl',	'Tpd52l2',	'Bmp15',	'Dgkk',	'Usp27x',	'Suv39h1',	'Bcor',	'Maob',	'Xiap',	'Stag2',	'Gpc4',	'Gpc3',	'Hprt',	'Plac1',	'Plxnb3',	'Msn',	'Zmym3',	'Hdac8',	'Abcb7',	'Fgf16',	'Pcdh19',	'Tceal9',	'Col4a5',	'Map3k15',	'Pir',	'Tmsb4x',	'Gm21887',	'Lrrcc1',	'Car2',	'Gm9733',	'Pde7a',	'Ect2',	'Fndc3b',	'Tmem212',	'Prkci',	'Sox2',	'Atp11b',	'1810062G17Rik',	'Intu',	'Mgst2',	'Stoml3',	'Smad9',	'Tm4sf4',	'P2ry13',	'Mbnl1',	'Lekr1',	'Schip1',	'Sptssb',	'Golim4',	'Ctso',	'Sfrp2',	'Glt28d2',	'Kirrel',	'Etv3',	'Arhgef2',	'Gon4l',	'Efna1',	'Kcnn3',	'Tchhl1',	'Mcl1',	'Car14',	'Hsd3b2',	'Man1a2',	'Igsf3',	'Slc22a15',	'Magi3',	'Cttnbp2nl',	'Cym',	'Cdc14a',	'Cnn3',	'Fabp2',	'Sec24d',	'Camk2d',	'Enpep',	'Npnt',	'Bdh2',	'Gm5105',	'Pdlim5',	'Gtf2b',	'Hs2st1',	'Znhit6',	'Ddah1',	'Adgrl2',	'Erich3',	'Fam110b',	'Sdcbp',	'Nsmaf',	'Triqk',	'Necab1',	'Ccnc',	'Bach2',	'Aqp3',	'Fbxo10',	'Shb',	'Igfbpl1',	'Slc44a1',	'Zfp462',	'Pakap',	'Slc31a1',	'Tnc',	'Nfib',	'Zdhhc21',	'Cer1',	'Adamtsl1',	'Mllt3',	'Gm10306',	'Jun',	'Hook1',	'E130114P18Rik',	'Patj',	'Kank4',	'Cachd1',	'Tctex1d1',	'Oma1',	'Acot11',	'Ssbp3',	'Gpx7',	'Dmrta2',	'Trabd2b',	'Olfr1342',	'Guca2a',	'Scmh1',	'Ctps',	'Nfyc',	'Mycl',	'Trit1',	'Macf1',	'AU040320',	'Zbtb8b',	'Ahdc1',	'Wasf2',	'Nipal3',	'6030445D17Rik',	'Cda',	'Pqlc2',	'Iffo2',	'Tardbp',	'Rbp7',	'Gpr157',	'Camta1',	'Kcnab2',	
                    'B230104I21Rik',	'Morn1',	'Mmp23',	'Cdk14',	'Sema3e',	'Sema3c',	'Wdr86',	'Rnf32',	'Tmem214',	'Tcf23',	'Mrpl33',	'Rnf4',	'Sorcs2',	'Cytl1',	'C1qtnf7',	'Fgfbp1',	'Qdpr',	'Rbpj',	'Tbc1d19',	'Stim2',	'0610040J01Rik',	'Tlr1',	'Wdr19',	'Apbb2',	'Bend4',	'Gm5108',	'Spata18',	'Rasl11b',	'Fip1l1',	'Kit',	'Kdr',	'Nmu',	'Grsf1',	'Gc',	'Npffr2',	'Cxcl1',	'Sowahb',	'Paqr3',	'Lin54',	'Arhgap24',	'Gm17660',	'Lrrc8c',	'Tgfbr3',	'Fgfrl1',	'Zfp932',	'Mn1',	'Mmab',	'Trpv4',	'Pxn',	'Vsig10',	'Wsb2',	'Fbxo21',	'Rasal1',	'Camkk2',	'Kdm2b',	'Hip1r',	'Zbed5',	'Tmem248',	'Galnt17',	'Gtf2i',	'Serpine1',	'Snx8',	'Lfng',	'Tnrc18',	'Zfp853',	'Baiap2l1',	'Smurf1',	'Mtus2',	'Tex26',	'Wdr95',	'Hsph1',	'Stard13',	'Pon3',	'Slc25a13',	'Glcci1',	'Phf14',	'Vwde',	'Tmem168',	'Foxp2',	'St7',	'Asb15',	'Tmem209',	'Bpgm',	'Cald1',	'Slc23a4',	'Ptn',	'Dgki',	'Creb3l2',	'Tmem213',	'Hipk2',	'Tbxas1',	'Ezh2',	'Gimap3',	'Chn2',	'Crhr2',	'Ccser1',	'Tnip3',	'Ndnf',	'Prdm5',	'Gng12',	'Il23r',	'Atoh8',	'Sh2d6',	'Eva1a',	'Pole4',	'Mthfd2',	'Sfxn5',	'Noto',	'Pcbp1',	'Prokr1',	'Eefsec',	'Wnt7a',	'Gm15737',	'Lrig1',	'Fam19a4',	'Lrrn1',	'Lmcd1',	'Rad18',	'Vgll4',	'Tamm41',	'Rho',	'Ankrd26',	'Dcp1b',	'Fbxl14',	'Wnk1',	'Tuba8',	'Tuba3a',	'Ccnd2',	'Tspan11',	'Etv6',	'Bcl2l14',	'Hebp1',	'Gsg1',	'Pbp2',	'Gm8994',	'E330021D16Rik',	'Spx',	'Sox5',	'Smco2',	'Ppfibp1',	'Rpl28',	'Usp29',	'Arhgap35',	'Zfp626',	'Actn4',	'Sipa1l3',	'Fxyd3',	'Zfp658',	'Sergef',	'Dbx1',	'Nipa2',	'Lrrk1',	'Lins1',	'Arrdc4',	'Mctp2',	'St8sia2',	'Sv2b',	'Anpep',	'Iqgap1',	'Alpk3',	'Mesd',	'Tmem135',	'Me3',	'Nars2',	'Emsy',	'Uvrag',	'Serpinh1',	'Neu3',	'P4ha3',	'Plekhb1',	'Scube2',	'Adm',	'Lyve1',	'Ctr9',	'Usp47',	'Psma1',	'Sox6',	'Smg1',	'Iqck',	'Cdr2',	'Rbbp6',	'Gsg1l',	'Zfp629',	'Fgfr2',	'Tacc2',	'Plekha1',	'Dmbt1',	'Acadsb',	'Ctbp2',	'Adam12',	'Mki67',	'Tcerg1l',	'Dpysl4',	'6430531B16Rik',	'Lsp1',	'Ins2',	'Tspan32',	'Cdkn1c',	'Tnfrsf23',	'Shank2',	'Ccnd1',	'1810010D01Rik',	'Cnksr3',	'Tab2',	'Hivep2',	'Reps1',	'Olig3',	'Stx7',	'Mical1',	'Sec63',	'Prep',	'Dcbld1',	'Ranbp2',	'Sh3rf3',	'44084',	'Anapc16',	'Psap',	'Pald1',	'Mypn',	'Ctnna3',	'Lrrtm3',	'Cabcoco1',	'Fam13c',	'Agpat3',	'Gzmm',	'Nuak1',	'Ric8b',	'Syn3',	'Igf1',	'Cfap54',	'Nudt4',	'Btg1',	'Dusp6',	'Rassf9',	'Ccdc59',	'Nap1l1',	'Tbc1d15',	'Tspan8',	'Cnot2',	'Cct2',	'Frs2',	'9530003J23Rik',	'Mdm2',	'Hmga2',	'Msrb3',	'Ppm1h',	'Agap2',	'Ndufa4l2',	'Ankrd10',	'Angpt2',	'Tcim',	'Hgsnat',	'Dctn6',	'Dusp4',	'Dlc1',	'Tusc3',	'Pdgfrl',	'Acsl1',	'Clcn3',	'Cbr4',	'Psd3',	'Sh2d4a',	'Nwd1',	'Ednra',	'Rbmxl1',	'Zfp827',	'1700011L22Rik',	'Clgn',	'Palm3',	'4930432K21Rik',	'G430095P16Rik',	'Zfp423',	'Adcy7',	'Tox3',	'Aktip',	'Nup93',	'Cdh1',	'Sntb2',	'Mon1b',	'Wwox',	'Atmin',	'Nup133',	'Agt',	'Capn9',	'Egln1',	'Irf2bp2',	'Nrp1',	'Flnb',	'Ptprg',	'Gm49355',	'Top2b',	'Rarb',	'Gng2',	'Usp54',	'Rps24',	'Erc2',	'Rft1',	'Arhgap22',	'Ptpn20',	'Zfp488',	
                    'Opn4',	'Fam213a',	'Cgrrf1',	'Ndrg2',	'Olfr221',	'Mettl3',	'Ajuba',	'Tgm1',	'Atp12a',	'Xpo4',	'Msra',	'Kif13b',	'Zfp395',	'Rhobtb2',	'Xpo7',	'Siah3',	'Serp2',	'Ccdc122',	'Akap11',	'Naa16',	'Wbp4',	'Cnmd',	'Diaph3',	'Spry2',	'Dct',	'Abcc4',	'Tmem123',	'Arhgap42',	'Jrkl',	'Maml2',	'Amotl1',	'Smco4',	'Arhgap32',	'Kcnj5',	'Kcnj1',	'Fli1',	'Tbcel',	'Arhgef12',	'Phldb1',	'Dscaml1',	'Sik3',	'Cadm1',	'Nxpe4',	'Dixdc1',	'Fdxacb1',	'Ppp2r1b',	'Exph5',	'Atm',	'Idh3a',	'Peak1',	'Odf3l1',	'1700017B05Rik',	'Neo1',	'Thsd4',	'Rplp1',	'Coro2b',	'Lctl',	'Megf11',	'Slc24a1',	'Snx22',	'Herc1',	'Lactb',	'Tpm1',	'Rora',	'Anxa2',	'Cgnl1',	'Fam83b',	'Slc17a5',	'Hmgn3',	'Ibtk',	'Plscr1',	'1190002N15Rik',	'Tfdp2',	'Spsb4',	'Cldn18',	'Slco2a1',	'Cdv3',	'Col6a6',	'Dusp7',	'Iqcf5',	'Dag1',	'Ccdc71',	'Eomes',	'Scn10a',	'Lyzl4',	'Zfp651',	'Tcaim',	'Adcy1',	'Gm11985',	'Fignl1',	'Ddc',	'Grb10',	'Egfr',	'Ppp3r1',	'Actr2',	'Bcl11a',	'Eml6',	'Sptbn1',	'Fbxw11',	'Rars',	'Wwc1',	'Tenm2',	'Clint1',	'Fstl4',	'Sparc',	'Igtp',	'Wnt9a',	'Prss38',	'Rai1',	'Srebf1',	'Map2k3',	'Cenpv',	'Trpv2',	'Cox10',	'Gas7',	'Usp43',	'Cfap52',	'Stx8',	'Ccdc42',	'Doc2b',	'1700016K19Rik',	'Slc46a1',	'Pex12',	'Rasl10b',	'Ccl6',	'Hnf1b',	'Bcas3',	'Gm11492',	'Mpo',	'Nog',	'Ankfn1',	'Mmd',	'Tom1l1',	'Chad',	'Igf2bp1',	'Hoxb1',	'Skap1',	'Srcin1',	'Lasp1',	'Cdk12',	'Grb7',	'Krt13',	'Tubg2',	'Fzd2',	'Higd1b',	'C1ql1',	'Mrc2',	'43900',	'Pitpnc1',	'Rgs9',	'Abca9',	'Kcnj16',	'Sox9',	'Llgl2',	'Myo15b',	'Sec14l1',	'Engase',	'Rptor',	'Slc38a10',	'Sectm1a',	'Asb13',	'Wdr37',	'Hecw1',	'Aoah',	'Hist1h4k',	'Hist1h2ak',	'Ripor2',	'Sox4',	'Foxf2',	'Mylk4',	'Serpinb1c',	'Tubb2a',	'Cdyl',	'Ly86',	'Rreb1',	'Dsp',	'Bmp6',	'Nedd9',	'Tmem170b',	'Adtrp',	'Tbc1d7',	'Mcur1',	'Gmpr',	'Spin1',	'Rab24',	'Klhl3',	'2010111I01Rik',	'Hsd17b3',	'Irx2',	'Ndufs6',	'Cmya5',	'Jmy',	'S100z',	'Enc1',	'Gm10260',	'Foxd1',	'Gm10320',	'Gtf2h2',	'Pik3r1',	'Srek1',	'Erbin',	'Adamts6',	'Ipo11',	'Rab3c',
                    'Gapt',	'Plk2',	'Il6st',	'Plpp1',	'BC067074',	'Ndufs4',	'Mocs2',	'Mfsd2b',	'Ldah',	'Osr1',	'Rock2',	'Kidins220',	'Sox11',	'Tpo',	'Snx13',	'Tspan13',	'Ispd',	'Snx6',	'Atp5s',	'Cdkl1',	'Atl1',	'Pygl',	'Six4',	'Ppp1r36',	'Plekhg3',	'Rab15',	'Rad51b',	'Dcaf5',	'Susd6',	'Map3k9',	'Numb',	'Riox1',	'Dnal1',	'Ltbp2',	'Ttll5',	'Esrrb',	'Irf2bpl',	'Ahsa1',	'Ptpn21',	'Calm1',	'Gpr68',	'Gm28051',	'Ppp4r4',	'Serpina16',	'Evl',	'Traf3',	'Inf2',	'Dnah11',	'Sp4',	'Ghr',	'Slc1a3',	'Nadk2',	'Lmbrd2',	'Basp1',	'Myo10',	'Fbxl7',	'Ank',	'Trio',	'Dnah5',	'Dap',	'Ankrd33b',	'Sdc2',	'Ywhaz',	'Ncald',	'Azin1',	'Angpt1',	'Rspo2',	'Med30',	'Ext1',	'Mrpl13',	'Mtbp',	'Has2',	'Atad2',	'Fer1l6',	'Fam84b',	'Asap1',	'Gm6569',	'1110038F14Rik',	'Mb',	'Apol6',	'Txn2',	'Ift27',	'Mkl1',	'Serhl',	'Cyb5r3',	'Arfgap3',	'Rtl6',	'Ppara',	'Celsr1',	'Fam19a5',	'Adamts20',	'Tmem117',	'Ano6',	'Slc38a4',	'Slc48a1',	'Adcy6',	'Tuba1b',	'Lima1',	'Fignl2',	'Krt18',	'Rarg',	'Parn',	'Klhl6',	'Trp63',	'Cldn1',	'Fgf12',	'Mb21d2',	'Atp13a5',	'Atp13a4',	'Opa1',	'Fam43a',	'Dlg1',	'Snx4',	'Zfp148',	'Ccdc14',	'Ildr1',	'Ndufb4',	'Fstl1',	'Lsamp',	'Qtrt2',	'Cd96',	'Nectin3',	'Alcam',	'Zpld1',	'Filip1l',	'Cxadr',	'App',	'Bach1',	'Clic6',	'Runx1',	'Pcp4',	'Snx9',	'Qk',	'Igf2r',	'Afdn',	'Lix1',	'Spaca6',	'D17Wsu92e',	'Ppard',	'Abcg1',	'Cryaa',	'Akap8l',	'Gm8909',	'Trim31',	'2410137M14Rik',	'Ttbk1',	'1700001C19Rik',	'Ccnd3',	'Bysl',	'Lrfn2',	'Mocs1',	'Kif6',	'Satb1',	'Themis3',	'Ptprm',	'Epb41l3',	'Rasgrp3',	'Rmdn2',	'Hnrnpll',	'Cox7a2l',	'Zfp36l2',	'Plekhh2',	'Dync2li1',	'Calm2',	'Gm47283',	'Gata6',	'Ankrd29',	'Lama3',	'Cabyr',	'Impact',	'Cdh2',	'Rnf125',	'2700062C07Rik',	'AW554918',	'Celf4',	'Epb41l4a',	'Ctnna1',	'Ecscr',	'Cystm1',	'Pcdhgc5',	'Nr3c1',	'Pabpc2',	'Tnfaip8',	'Hsd17b4',	'Zfp474',	'Snx2',	'Snx24',	'Cep120',	'Megf10',	'Prrc1',	'Ctxn3',	'Slc12a2',	'Ablim3',	'Apcdd1',	'Alpk2',	'Oacyl',	'Chmp1b',	'Spire1',	'Ldlrad4',	'Fam210a',	'Tcf4',	'Gm9925',	'Lipg',	'Ctif',	'Gm6133',	'Hsbp1l1',	'Nfatc1',	'Cyb5a',	'Timm21',	'Chka',	'Atl3',	'Pla2g16',	'Fads3',	'Myrf',	'Tle4',	'Gnaq',	'Pcsk5',	'Rorb',	'Trpm3',	'1700028P14Rik',	'Ptar1',	'Dmrt1',	'Dmrt3',	'Smarca2',	'Cd274',	'Mbl2',	'Cstf2t',	'Sgms1',	'Pten',	'Slc16a12',	'Ppp1r3c',	'Btaf1',	'Fra10ac1',	'Tm9sf3',	'Gm340',	'Slit1',	'Frat2',	'Gbf1',	'Stn1',	'Sorcs3',	'Adra2a',	'Acsl5',	'Vti1a',	'Tcf7l2',	'Adrb1',	'Rab11fip2'))

integrate <- AddModuleScore(object = integrate, features = AT1RClean, ctrl = 500, name = "NKXAT1RetainedCleanGenes", search = TRUE, random.seed = 1)
FeaturePlot(object = integrate, features = 'NKXAT1RetainedCleanGenes1', split.by = "ID", cols = c('lightgrey','red'))
FeaturePlot(object = integrate, features = 'NKXAT1RetainedCleanGenes1', cols = c('lightgrey','red'))


BACClean <- list(c('Rb1cc1',	'Prdm14',	'Rdh10',	'Ube2w',	'Tmem70',	'Paqr8',	'Tram2',	'Kcnq5',	'Smap1',	'Phf3',	'Prim2',	'Fam168b',	'Hs6st1',	'Uggt1',	'Arid5a',	'Kansl3',	'Cnnm4',	'Ankrd23',	'Gm42417',	'Tmem131',	'Eif5b',	'Rpl31',	'Tbc1d8',	'Il1r1',	'Il18r1',	'Myo1b',	'Nab1',	'Nemp2',	'Mfsd6',	'Ankrd44',	'Hspe1',	'Rftn2',	'Aox1',	'Orc2',	'Tmem237',	'Sumo1',	'Ccnyl1',	'Kansl1l',	'Acadl',	'Atic',	'Smarcal1',	'Cxcr2',	'Vil1',	'Plcd4',	'Cnppd1',	'Tuba4a',	'Asic4',	'Sgpp2',	'Mrpl44',	'Mff',	'A030014E15Rik',	'Cab39',	'Dis3l2',	'Ngef',	'Dgkd',	'Ugt1a6b',	'Arl4c',	'Sh3bp4',	'Rab17',	'Lrrfip1',	'Ramp1',	'Ilkap',	'Hes6',	'Ndufa10',	'Kif1a',	'Sned1',	'Mterf4',	'D2hgdh',	'Gin1',	'Pign',	'Tnfrsf11a',	'Dsel',	'Clasp1',	'Tfcp2l1',	'Epb41l5',	'Tmem37',	'Lypd1',	'Tmem163',	'Cd55',	'Gm29427',	'Srgap2',	'Fam72a',	'Elk4',	'Mfsd4a',	'Ppp1r15b',	'Golt1a',	'Kiss1',	'Sox13',	'Mybph',	'Adora1',	'Rabif',	'Ppp1r12b',	'Rnpep',	'Phlda3',	'Camsap2',	'Zbtb41',	'Pla2g4a',	'Rnf2',	'Tsen15',	'Teddm2',	'Mr1',	'Xpr1',	'Cep350',	'Soat1',	'Tnr',	'4930523C07Rik',	'Rabgap1l',	'Rc3h1',	'Prdx6',	'Dnm3',	'Fmo1',	'Sft2d2',	'Dcaf6',	'Creg1',	'Mgst3',	'Nos1ap',	'Zbtb18',	'Psen2',	'Stum',	'Lin9',	'Acbd3',	'Lefty2',	'Tmem63a',	'Fbxo28',	'Mia3',	'43892',	'Iars2',	'Kctd3',	'Fam71a',	'Atf3',	'Ints7',	'Plxna2',	'Dclre1c',	'Prpf18',	'Optn',	'Il15ra',	'Mindy3',	'Rsu1',	'Nebl',	'Skida1',	'Spopl',	'Noxa1',	'Tor4a',	'Ssna1',	'Man1b1',	'Uap1l1',	'Clic3',	'Tmem141',	'Camsap1',	'Nacc2',	'Gpsm1',	'Dnlz',	'Agpat2',	'Col5a1',	'Ralgds',	'Cel',	'Ttf1',	'Setx',	'Wdr34',	'Miga2',	'Dolpp1',	'Asb6',	'Abl1',	'Lamc3',	'Prrc2b',	'Swi5',	'Dnm1',	'Lcn2',	'St6galnac6',	'Cdk9',	'Ttc16',	'Stxbp1',	'Psmd5',	'Phf19',	'Traf1',	'Hc',	'Gsn',	'Lhx6',	'Ptgs1',	'Rbm43',	'Nmi',	'Gpd2',	'Acvr1',	'Ly75',	'Bbs5',	'Ccdc173',	'Gm26558',	'Tlk1',	'Dcaf17',	'Map3k20',	'Cir1',	'Osbpl6',	'Plekha3',	'Ccdc141',	'Nckap1',	'Rapsn',	'Mybpc3',	'Atg13',	'Pex16',	'Prdm11',	'Trim44',	'Slc1a2',	'Cd44',	'Pdhx',	'Cat',	'Hipk3',	'Mettl15',	'Aven',	'Eif2ak4',	'Pak6',	'Bahd1',	'Rmdn3',	'Chac1',	'Exd1',	'Vps39',	'Tgm5',	'Ckmt1',	'Hypk',	'Secisbp2l',	'Cops2',	'Hdc',	'Usp50',	'Stard7',	'Acoxl',	'Slc20a1',	'Snrpb',	'Vps16',	'Spef1',	'Ap5s1',	'Rassf2',	'Slc23a2',	'Cds2',	'Trmt6',	'Jag1',	'Dstn',	'Rrbp1',	'Rin2',	'Ralgapa2',	'Kiz',	'Apmap',	'Pygb',	'Fam110a',	'Srxn1',	'Defb20',	'H13',	'Bcl2l1',	'Asxl1',	'Bpifb1',	'Pxmp4',	'Chmp4b',	'Raly',	'Eif2s2',	'Dynlrb1',	'Trp53inp2',	'Acss2',	'Myh7b',	'Uqcc1',	'Spag4',	'Scand1',	'Cnbd2',	'Src',	'Tti1',	'Rprd1b',	'Ada',	'Wfdc12',	'Sdc4',	'Pigt',	'Snx21',	'Zfp335',	'Slc35c2',	'Elmo2',	'Eya2',	'Arfgef2',	'Cse1l',	'Ube2v1',	'Ripor3',	'Adnp',	'Bmp7',	'Gm20721',	'Lsm14b',	'Osbpl2',	'Adrm1',	'Cables2',	'Ythdf1',	'Ptk6',	'Helz2',	'Zfp512b',	'Myt1',	'Wdr13',	'Gm14862',	'1810030O07Rik',	'Cask',	'Fundc1',	'Kdm6a',	'Araf',	'Lonrf3',	'Lamp2',	'Rap2c',	'Fhl1',	'Fmr1',	'Gabrq',	'Trex2',	'Bgn',	'Taz',	'Pls3',	'Prkx',	'Pdk3',	'Ophn1',	'Efnb1',	'Pja1',	'Phka1',	'Magee1',	'Atrx',	'Morf4l2',	'Tbc1d8b',	'Prps1',	'Il13ra2',	'Shroom2',	'Ap1s2',	'Msl3',	'Tpd52',	'Zbtb10',	'Impa1',	'Chmp4c',	'Mtfr1',	'Gyg',	'Tbl1xr1',	'Ect2',	'Nceh1',	'Tmem212',	'Eif5a2',	'Gpr160',	'Zmat3',	'Gnb4',	'Ttc14',	'Acad9',	'Anxa5',	'Fgf2',	'Spry1',	'Ankrd50',	'Larp1b',	'Noct',	'Spg20',	'Nbea',	'Tm4sf4',	'Igsf10',	'Arhgef26',	'Ssr3',	'Ptx3',	'Mfsd1',	'Il12a',	'Fnip2',	'Ppid',	'Plrg1',	'Gm37240',	'Fam160a1',	'Rps3a1',	'Gm38392',	'Slc25a44',	'Sema4a',	'Mex3a',	'Rxfp4',	'Rit1',	'Gon4l',	'Msto1',	'Dap3',	'Slc50a1',	'Ube2q1',	'Il6ra',	'Rps27',	'Dennd4b',	'Snapin',	'Tdrkh',	'Cgn',	'Selenbp1',	'Zfp687',	'Golph3l',	'Ensa',	'Adamtsl4',	'Ciart',	'Anp32e',	'Rnf115',	'Pdzk1',	'Acp6',	'Chd1l',	'Fmo5',	'Adam30',	'Reg4',	'Cd101',	'Ptgfrn',	'Atp1a1',	'Tspan2',	'Bcas2',	'Syt6',	'Olfml3',	'Hipk1',	'Dclre1b',	'Slc16a1',	'St7l',	'Rap1a',	'I830077J02Rik',	'Dennd2d',	'Lamtor5',	'Gstm6',	'Sort1',	'Mybphl',	'Stxbp3',	'Fndc7',	'Prpf38b',	'Extl2',	'Trmt13',	'Bcar3',	'Sec24d',	'Npnt',	'Gstcd',	'Tet2',	'Cisd2',	'Adh5',	'Eif4e',	'Tspan5',	'Rap1gds1',	'Clca2',	'Ctbs',	'Spata1',	'Usp33',	'Srsf11',	'Lrrc40',	'Tmem68',	'Lyn',	'Tox',	'Plekhf2',	'Dpy19l4',	'Osgin2',	'Atp6v0d2',	'Ggh',	'Tstd3',	'Fbxl4',	'Map3k7',	'Ube2j1',	'Ddx58',	'Smim27',	'Bag1',	'Aqp7',	'Ube2r2',	'Dcaf12',	'Fam219a',	'Fam214b',	'Unc13b',	'Rusc2',	'Car9',	'Tln1',	'Npr2',	'Olfr71',	'Spaar',	'Glipr2',	'Zcchc7',	'Slc25a51',	'Xpa',	'Anp32b',	'Coro2a',	'Sec61b',	'Nr4a3',	'Msantd3',	'Ppp3r2',	'Abca1',	'Slc44a1',	'Fktn',	'Zfp462',	'Klf4',	'Tmem245',	'Zkscan16',	'Ugcg',	'Snx30',	'Atp6v1g1',	'Tmem268',	'Tnfsf15',	'Tnc',	'Trim32',	'Rasef',	'Kdm4c',	'Ttc39b',	'Snapc3',	'Focad',	'Atg4c',	'Jak1',	'Slc35d1',	'Oma1',	'Prkaa2',	'Gpx7',	'Btf3l4',	'Osbpl9',	'Ttc39a',	'Dmrta2',	'Bend5',	'Gm12830',
                   'Cyp4b1',	'Efcab14',	'Mob3c',	'Lurap1',	'Tspan1',	'Plk3',	'Rps8',	'Atp6v0b',	'Dph2',	'Ipo13',	'Artn',	'St3gal3',	'Szt2',	'Elovl1',	'AU022252',	'Rimkla',	'Guca2a',	'Slfnl1',	'Ctps',	'Kcnq4',	'Cap1',	'Macf1',	'Ndufs5',	'Inpp5b',	'Yrdc',	'Zc3h12a',	'Stk40',	'Sh3d21',	'Thrap3',	'Tekt2',	'AU040320',	'Tmem35b',	'Azin2',	'Rnf19b',	'Zbtb8a',	'Dcdc2b',	'Ptp4a2',	'Pef1',	'Hcrtr1',	'Fabp3',	'Sdc3',	'Tmem200b',	'Ptafr',	'Slc9a1',	'Pigv',	'Sh3bgrl3',	'Slc30a2',	'Mtfr1l',	'Selenon',	'Clic4',	'Ifnlr1',	'Pnrc2',	'Gale',	'Cdc42',	'Rap1gap',	'Ece1',	'Hp1bp3',	'Pink1',	'Camk2n1',	'Otud3',	'Epha2',	'Spen',	'Tmem82',	'Plekhm2',	'Agmat',	'Dhrs3',	'Tnfrsf1b',	'Fv1',	'Mfn2',	'Plod1',	'2510039O18Rik',	'Mthfr',	'Fbxo6',	'Fbxo2',	'Ubiad1',	'Pgd',	'Ctnnbip1',	'Clstn1',	'Slc25a33',	'Spsb1',	'H6pd',	'Rere',	'Park7',	'Dnajc11',	'Thap3',	'Phf13',	'Klhl21',	'Plekhg5',	'Kcnab2',	'Dffb',	'Lrrc47',	'Smim1',	'Tprgl',	'Tnfrsf14',	'Gabrd',	'Gnb1',	'Mxra8',	'B3galt6',	'Isg15',	'Ankib1',	'Fzd1',	'Cldn12',	'Cfap69',	'Tmem243',	'Gnat3',	'Rsbn1l',	'Gsap',	'Napepld',	'Tomm7',	'Atg9b',	'Tmub1',	'Asb10',	'Nub1',	'Kmt2c',	'Paxip1',	'Insig1',	'Rbm33',	'Ost4',	'Khk',	'Gm29609',	'Mrpl33',	'Rbks',	'Nelfa',	'Zfyve28',	'Fam193a',	'Mfsd10',	'Msantd1',	'Lrpap1',	'Acox3',	'Sh3tc1',	'Psapl1',	'Tada2b',	'Man2b2',	'Zbtb49',	'Slc2a9',	'Wdr1',	'Zfp518b',	'Cd38',	'Sel1l3',	'Rell1',	'Tlr1',	'Tmem156',	'Rbm47',	'Nipal1',	'Dcun1d4',	'Chic2',	'Clock',	'Hopx',	'Rufy3',	'Ankrd17',	'Btc',	'Rchy1',	'Cdkl2',	'Cxcl10',	'Scarb2',	'Anxa3',	'Paqr3',	'Prkg2',	'Nkx6-1',	'Wdfy3',	'1700016H13Rik',	'Nudt9',	'Gm17660',	'Zfp326',	'Fam69a',	'Cplx1',	'Gak',	'Idua',	'Chfr',	'Pgam5',	'Pxmp2',	'Galnt9',	'Noc4l',	'Ddx51',	'Ep400',	'Tpst2',	'Coro1c',	'Kctd10',	'Ube3b',	'Ankrd13a',	'Oasl1',	'Rnf10',	'Gatc',	'Rab35',	'Cit',	'Prkab1',	'Suds3',	'Taok3',	'Fbxo21',	'Tpcn1',	'Hectd4',	'Aldh2',	'Atxn2',	'Sh2b3',	'Myl2',	'Atp2a2',	'Ift81',	'Rnf34',	'Rhof',	'Diablo',	'Gm49027',	'Clip1',	'Zcchc8',	'Hcar2',	'Hip1r',	'Vps37b',	'Kmt5a',	'Snrnp35',	'Rilpl1',	'Zfp664',	'Scarb1',	'Glt1d1',	'Ran',	'Nupr1l',	'Tpst1',	'Kctd7',	'Rabgef1',	'Tmem248',	'Ncf1',	'Lat2',	'Eif4h',	'Stx1a',	'Tmem120a',	'Srrm3',	'Hspb1',	'Orai2',	'Gm20605',	'Sap25',	'Irs3',	'Cyp3a13',	'Mblac1',	'Sun1',	'Gpr146',	'Gper1',	'Mafk',	'Snx8',	'Ap5z1',	'Papolb',	'Rac1',	'Cyth3',	'Usp42',	'Ccz1',	'Ocm',	'Bhlha15',	'Smurf1',	'Arpc1b',	'Zkscan14',	'Zfp655',	'Wasf3',	'Rasl11a',	'Lnx2',	'Slc7a1',	'Ubl3',	'Gm42791',	'N4bp2l1',	'Casd1',	'Pon1',	'Slc25a13',	'C1galt1',	'Tspan12',	'Cadps2',	'Wasl',	'Rbm28',	'Calu',	'Kcp',	'Ube2h',	'Klf14',	'Mkln1',	'Plxna4',	'Akr1b7',	'Tmem140',	'Stra8',	'Fam180a',	'Zc3hav1',	'Ubn2',	'Clec2l',	'Tra2a',	'Fam221a',	'Osbpl3',	'Hibadh',	'Tax1bp1',	'Cpvl',	'Prr15',	'Plekha8',	'Znrf2',	'Nod1',	'Aqp1',	'Gng12',	'Gadd45a',	'Krcc1',	'Rnf103',	'Kdm3a',	'Reep1',	'Sftpb',	'Rnf181',	'Vamp8',	'Mat2a',	'Sh2d6',	'Suclg1',	'Mrpl19',	'Pole4',	'Hk2',	'Dok1',	'Loxl3',	'Mogs',	'Dctn1',	'Mthfd2',	'Mob1a',	'Bola3',	'Smyd5',	'Add2',	'Pcbp1',	'Asprv1',	'Mxd1',	'Snrnp27',	'Gmcl1',	'Aak1',	'Rab43',	'Cnbp',	'Rab7',	'Rpn1',	'Sec61a1',	'Abtb1',	'Chchd6',	'Txnrd3',	'Zxdc',	'Klf15',	'Aldh1l1',	'Slc41a3',	'Nup210',	'Hdac11',	'Fbln2',	'Tmem43',	'Xpc',	'Nr2c2',	'Prickle2',	'Suclg2',	'Mitf',	'Rybp',	'Gxylt2',	'Edem1',	'Thumpd3',	'Lhfpl4',	'Prrt3',	'Irak2',	'Tatdn2',	'Atp2b2',	'Tamm41',	'Mkrn2',	'Raf1',	'Tmem40',	'Rpl32',	'43898',	'Fxyd4',	'Rasgef1a',	'Adipor2',	'Il17ra',	'Foxj2',	'Chd4',	'Nop2',	'Gapdh',	'Ltbr',	'Scnn1a',	'Plekhg6',	'Cd9',	'Tspan9',	'Fkbp4',	'Cd69',	'Gabarapl1',	'Lrp6',	'Crebl2',	'Strap',	'Recql',	'Ldhb',	'Lrmp',	'Bhlhe41',	'Ccdc91',	'Amn1',	'Cacng7',	'Leng9',	'Tnnt1',	'Ppp6r1',	'Shisa7',	'Zfp580',	'Zfp787',	'Selenow',	'Nop53',	'Ccdc9',	'Bbc3',	'Sae1',	'Zc3h4',	'Slc1a5',	'Ppp5c',	'Opa3',	'Vasp',	'Ercc2',	'Mark4',	'Smg9',	'Arhgef1',	'Grik5',	'Pou2f2',	'Dedd2',	'Ceacam1',	'Tmem91',	'Tgfb1',	'Cyp2b10',	'Egln2',	'Numbl',	'Sptbn4',	'Akt2',	'Zfp36',	'Med29',	'Catsperg1',	'2200002D01Rik',	'Dpf1',	'Sdhaf1',	'Scn1b',	'Lsm14a',	'Faap24',	'Pdcd5',	'Dpy19l3',	'Vsig10l',	'Mybpc2',	'Spib',	'Kcnc3',	'Myh14',	'Ap2a1',	'Fcgrt',	'Aldh16a1',	'Gys1',	'Bcat2',	'Rasip1',	'Kdelr1',	'Ccdc114',	'Sergef',	'Uevld',	'Gas2',	'Nipa2',	'Herc2',	'Klf13',	'E030018B13Rik',	'Fan1',	'Lysmd4',	'Synm',	'Akap13',	'Aen',	'Hapln3',	'Pex11a',	'Zfp710',	'Fes',	'Slc28a1',	'Rps17',	'Cemip',	'Abhd17c',	'Fah',	'Prss23',	'Picalm',	'Ccdc83',	'Kctd21',	'Alg8',	'Ndufc2',	'Ints4',	'Acer3',	'Mogat2',	'Map6',	'Arrb1',	'Tpbgl',	'Ucp3',	'Ucp2',	'Arhgef17',	'Pde2a',	'Clpb',	'Il18bp',	'Nup98',	'Trim6',	'Smpd1',	'Apbb1',	'Hpx',	'Trim3',	'Rpl27a',	'Nrip3',	'Dennd5a',	'Tmem41b',	'Ipo7',	'Swap70',	'Mrvi1',	'Gm4353',	'Rps13',	'Pik3c2a',	'Arl6ip1',	'Smg1',	'Coq7',	
                   'Gde1',	'Gprc5b',	'Lyrm1',	'Abca14',	'Usp31',	'Scnn1b',	'Lcmt1',	'Il4ra',	'D430042O09Rik',	'Gsg1l',	'Atxn2l',	'Sult1a1',	'Tmem219',	'Kctd13',	'Kif22',	'Zfp768',	'Ctf1',	'Kat8',	'Armc5',	'BC017158',	'Rgs10',	'Bag3',	'Inpp5f',	'Fgfr2',	'Nsmce4a',	'Fam53b',	'Nps',	'9430038I01Rik',	'Glrx3',	'Ppp2r2d',	'Zfp941',	'Scgb1c1',	'Irf7',	'Tmem80',	'Pnpla2',	'Mob2',	'Dusp8',	'Kcnq1',	'Slc22a18',	'Osbpl5',	'Shank2',	'Ppfia1',	'Mrgprf',	'Ipcef1',	'Cnksr3',	'Ginm1',	'Utrn',	'Ltv1',	'Reps1',	'Arfgef3',	'Tnfaip3',	'Il22ra2',	'Map3k5',	'Pde7b',	'Ahi1',	'Akap7',	'Rnf146',	'Tspyl1',	'Tspyl4',	'Mfsd4b4',	'Mfsd4b5',	'Armc2',	'Foxo3',	'Sec63',	'Scml4',	'Bend3',	'Gm9803',	'Ascc3',	'Mcm9',	'Fam184a',	'Msl3l2',	'Lims1',	'Edar',	'44084',	'Mcu',	'Eif4ebp2',	'Lrrc20',	'Aifm2',	'Tspan15',	'Tacr2',	'Hk1',	'4930507D05Rik',	'Ccar1',	'Mypn',	'Lrrtm3',	'A930033H14Rik',	'Tfam',	'Bcr',	'Upb1',	'Susd2',	'Gstt1',	'Smarcb1',	'S100b',	'Pcnt',	'2610028H24Rik',	'Lss',	'1810043G02Rik',	'Trappc10',	'Agpat3',	'Cdc34',	'Bsg',	'Misp',	'Tmem259',	'Abca7',	'Arhgap45',	'Polr2e',	'Gpx4',	'Dazap1',	'Rps15',	'2310011J03Rik',	'Adamtsl5',	'Mknk2',	'Ap3d1',	'Dot1l',	'Plekhj1',	'Gng7',	'4930442H23Rik',	'Hmg20b',	'Smim24',	'Nfic',	'Aes',	'Glt8d2',	'Appl2',	'Tcp11l2',	'Polr3b',	'Timp3',	'Nt5dc3',	'Parpbp',	'Scyl2',	'Uhrf1bp1l',	'Hal',	'Nr2c1',	'Tmcc3',	'Nudt4',	'Poc1b',	'Dusp6',	'Zdhhc17',	'Atxn7l3b',	'Tbc1d15',	'Rab21',	'Frs2',	'Wif1',	'Gns',	'Tbk1',	'Slc16a7',	'Agap2',	'B4galnt1',	'Itga7',	'Stxbp2',	'Lig4',	'Col4a1',	'Naxd',	'Cars2',	'Tubgcp3',	'Atp11a',	'Adprhl1',	'Dcun1d2',	'Tmem255b',	'Gas6',	'Rasa3',	'Champ1',	'Cln8',	'Angpt2',	'Atp7b',	'Nkx6-3',	'Adam32',	'Adam9',	'Plpp5',	'Hook3',	'Brf2',	'Rab11fip1',	'Got1l1',	'Tti2',	'Mak16',	'Wrn',	'Smim18',	'Dlc1',	'Pdgfrl',	'Asah1',	'Ccdc110',	'Ankrd37',	'Snx25',	'Cfap97',	'Enpp6',	'Wwc2',	'Lzts1',	'Nr2c2ap',	'Ell',	'Rab3a',	'Ccdc124',	'Use1',	'Pgls',	'Klf2',	'Med26',	'Pkn1',	'Ptger1',	'Nanos3',	'Mri1',	'Dnase2a',	'Junb',	'Hook2',	'Vps35',	'Orc6',	'Dnaja2',	'Itfg1',	'Siah1a',	'N4bp1',	'Brd7',	'Chd9',	'Rpgrip1l',	'Herpud1',	'Nlrc5',	'Cpne2',	'Cx3cl1',	'Mmp15',	'Ccdc113',	'Cmtm4',	'Terb1',	'Cbfb',	'D230025D16Rik',	'Nol3',	'4931428F04Rik',	'Exoc3l',	'Lrrc29',	'Fhod1',	'Tppp3',	'Hsd11b2',	'Ripor1',	'Nfatc3',	'Cyb5b',	'Hp',	'Ap1g1',	'Marveld3',	'Ldhd',	'Zfp1',	'Bcar1',	'Tmem231',	'Terf2ip',	'Syce1l',	'Dynlrb2',	'Cdyl2',	'Sdr42e1',	'Hsbp1',	'Mlycd',	'Osgin1',	'Hsdl1',	'Cotl1',	'Klhl36',	'Gse1',	'Map1lc3b',	'Zcchc14',	'Piezo1',	'Pabpn1l',	'Ankrd11',	'Spata2l',	'Rhou',	'Abcb10',	'Capn9',	'Gnpat',	'Sipa1l2',	'Map10',	'Coa6',	'Itgb1',	'Pxk',	'Gm49355',	'Psmd6',	'Ngly1',	'Rpl15',	'Ube2e1',	'Ecd',	'Usp54',	'Fut11',	'Vcl',	'Lrmda',	'Ppif',	'Anxa11',	'Wnt5a',	'Cacna1d',	'Tkt',	'Prkcd',	'Rft1',	'Sfmbt1',	'Nisch',	'Phf7',	'Gprin2',	'Fam25c',	'Opn4',	'Wapl',	'Ccser2',	'Tspan14',	'Fam213a',	'Sftpd',	'Gmfb',	'Dlgap5',	'Atg14',	'Ktn1',	'Eddm3b',	'Hnrnpc',	'Ppp1r3e',	'Bcl2l2',	'Myh6',	'Nrl',	'Tm9sf1',	'Tgm1',	'Dhrs1',	'Khnyn',	'Cenpj',	'Gm6904',	'Cdadc1',	'Ebpl',	'Ctsb',	'Fdft1',	'Fam167a',	'Ints9',	'Nuggc',	'Dpysl2',	'Dock5',	'Chmp7',	'9930012K11Rik',	'Pdlim2',	'Sorbs3',	'Ppp3cc',	'Lgi3',	'Fam160b2',	'Fndc3a',	'Lrch1',	'Siah3',	'Tsc22d1',	'Dnajc15',	'Dgkh',	'Vwa8',	'Rgcc',	'Diaph3',	'Acod1',	'Cln5',	'Dzip1',	'Stk24',	'Gpr18',	'Tm9sf2',	'Tmem123',	'Maml2',	'Amotl1',	'Piwil4',	'Fut4',	'Zfp426',	'S1pr2',	'Icam1',	'Pde4a',	'Ilf3',	'Dnm2',	'Spc24',	'Kank2',	'Zfp872',	'Zfp810',	'Herpud2',	'Adamts8',	'Zbtb44',	'Prdm10',	'Esam',	'Nectin1',	'Rnf26',	'Bcl9l',	'Ddx6',	'Zbtb16',	'Usp28',	'Ncam1',	'Plet1',	'Dlat',	'Dixdc1',	'Cryab',	'Ppp2r1b',	'Fdx1',	'Rdx',	'Zc3h12c',	'Acat1',	'Dmxl2',	'Idh3a',	'Dnaja4',	'Wdr61',	'Chrnb4',	'Snx33',	'Imp3',	'Ptpn9',	'Cox5a',	'Tmem202',	'Gramd2',	'Fem1b',	'Dennd4a',	'Hacd3',	'Dpp8',	'Rasl12',	'Mtfmt',	'Plekho2',	'Zfp609',	'Gcnt3',	'Adam10',	'Ccpg1os',	'Ccpg1',	'Khdc3',	'Fam214a',	'Tmod3',	'Elovl5',	'Ick',	'Slc17a5',	'Myo6',	'Tbc1d2b',	'Plscr5',	'Xrn1',	'Tfdp2',	'Zbtb38',	'Pccb',	'Ryk',	'Rab6b',	'Cdv3',	'Nudt16',	'Ppm1m',	'Alas1',	'Dusp7',	'Abhd14b',	'Pcbp4',	'Parp3',	'Rrp9',	'Tex264',	'Mapkapk3',	'Cish',	'Hemk1',	'Zmynd10',	'Tusc2',	'Sema3b',	'Rbm6',	'Mst1',	'Apeh',	'Gpx1',	'P4htm',	'Prkar2a',	'Celsr3',	'Shisa5',	'Plxnb1',	'Dhx30',	'Cspg5',	'Scap',	'Ptpn23',	'Nradd',	'Ccdc12',	'Rtp3',	'Lrrfip2',	'Clasp2',	'Dync1li1',	'Itga9',	'Ctdspl',	'Acaa1a',	'Xylb',	'Gorasp1',	'Trak1',	'Cck',	'Higd1a',	'Snrk',	'Ano10',	'Abhd5',	'Zdhhc3',	'Slc6a20a',	'Fyco1',	'Patz1',	'Limk2',	'Rnf185',	'Morc2a',	'Sec14l4',	'Mtfp1',	'Sec14l2',	'Ascc2',	'Uqcr10',	'Ap1b1',	'Gas2l1',	'Dbnl',	'Pgam2',	'Gck',	'Ykt6',	'Ogdh',	'Ppia',	'Purb',	'Grb10',	'Fbxo48',	'Pno1',	'Cep68',	'Sertad2',	'Zrsr1',	'Efemp1',	'Psme4',
                   'Erlec1',	'Bod1',	'Cpeb4',	'Mpg',	'Fbxw11',	'Slit3',	'Rars',	'Mat2b',	'Nudcd2',	'Ccng1',	'Adra1b',	'Rnf145',	'Rack1',	'Gm12184',	'Olfr1393',	'Rnf130',	'Sqstm1',	'Mgat4b',	'Col23a1',	'Sec24a',	'Sar1b',	'Ube2b',	'Cdkl3',	'Ppp2ca',	'Hspa4',	'Aff4',	'Gm9945',	'Kif3a',	'Irf1',	'P4ha2',	'Acsl6',	'Fnip1',	'Cdc42se2',	'Tnip1',	'Fam114a2',	'Irgm2',	'4930438A08Rik',	'Zfp672',	'Trim11',	'Iba57',	'2310033P09Rik',	'Arf1',	'Wnt3a',	'Myo15',	'Mief2',	'Tmem11',	'Natd1',	'Map2k3',	'Aldh3a1',	'Slc47a2',	'B9d1',	'Epn2',	'Ulk2',	'Akap10',	'Adora2b',	'Trim16',	'Tekt3',	'Pmp22',	'Hs3st3b1',	'Hs3st3a1',	'Elac2',	'1700086D15Rik',	'Myocd',	'Zkscan6',	'Gas7',	'Stx8',	'Ndel1',	'Per1',	'Cntrob',	'Tmem102',	'Gps2',	'Slc16a11',	'Med11',	'Inca1',	'Zfp3',	'Pimreg',	'Pitpnm3',	'Med31',	'Mybbp1a',	'Cyb5d2',	'Atp2a3',	'Camkk1',	'Shpk',	'Mettl16',	'Mnt',	'Sgsm2',	'Hic1',	'Rtn4rl1',	'Serpinf1',	'Wdr81',	'Rilp',	'Scarf1',	'Slc43a2',	'Myo1c',	'Doc2b',	'Vps53',	'Dbil5',	'Sdf2',	'Poldip2',	'Nos2',	'Lgals9',	'Ccl8',	'Mmp28',	'Ccl5',	'Heatr6',	'Ppm1d',	'Rnft1',	'Tubd1',	'Vmp1',	'Cltc',	'Gdpd1',	'Trim25',	'Gm45716',	'Stxbp4',	'Cox11',	'Spag9',	'Wfikkn2',	'Cacna1g',	'Spata20',	'Mycbpap',	'Rsad1',	'Slc35b1',	'B130006D01Rik',	'Phospho1',	'Snx11',	'Cdk5rap3',	'Pnpo',	'Tbkbp1',	'Npepps',	'Mrpl45',	'Gpr179',	'Arhgap23',	'Cwc25',	'B230217C12Rik',	'Fbxl20',	'Med1',	'Ppp1r1b',	'Stard3',	'Tcap',	'Erbb2',	'Ormdl3',	'Csf3',	'Med24',	'Thra',	'Nr1d1',	'Krt42',	'Nkiras2',	'Stat5a',	'Stat3',	'Cavin1',	'Naglu',	'Retreg3',	'Dhx8',	'Hdac5',	'Slc25a39',	'Dcakd',	'Hexim1',	'Plekhm1',	'Nsf',	'Cyb561',	'Ace',	'Map3k3',	'Limd2',	'Gm10840',	'Strada',	'Ccdc47',	'Cd79b',	'Polg2',	'Helz',	'Cacng1',	'Apoh',	'Gna13',	'9930022D16Rik',	'Cdc42ep4',	'Nat9',	'Cdr2l',	'Mrpl58',	'Mif4gd',	'Llgl2',	'Myo15b',	'Smim5',	'Unk',	'Unc13d',	'Mrpl38',	'Srp68',	'Ube2o',	'Rhbdf2',	'Cygb',	'Prcd',	'Mettl23',	'Sec14l1',	'6030468B19Rik',	'Tk1',	'Socs3',	'Cant1',	'C1qtnf1',	'Cbx4',	'Sgsh',	'Rnf213',	'Endov',	'Chmp6',	'Ndufaf8',	'Faap100',	'Gcgr',	'Pcyt2',	'Dcxr',	'Ccdc57',	'Csnk1d',	'Metrnl',	'Gdi2',	'Akr1c14',	'Pfkp',	'Gtpbp4',	'Larp4b',	'Zmynd11',	'Lgals8',	'Ggps1',	'Cdk13',	'Yae1d1',	'Stard3nl',	'BC005537',	'Acot13',	'Tubb2a',	'Fam50b',	'Prpf4b',	'Rpp40',	'Slc35b3',	'Tbc1d7',	'Gfod1',	'Nol7',	'Ranbp9',	'Mcur1',	'Gmpr',	'Nup153',	'Phf2',	'Card19',	'Tes3-ps',	'Iars',	'Sema4d',	'Auh',	'Arl10',	'Tspan17',	'Grk6',	'Ddx46',	'H2afy',	'Tifab',	'Klhl3',	'2210016F16Rik',	'Fbp2',	'2010111I01Rik',	'Ell2',	'Ttc37',	'Slf1',	'Cetn3',	'Rasa1',	'Gm20379',	'Zfyve16',	'Ap3b1',	'Pde8b',	'Col4a3bp',	'Tnpo1',	'Ptcd2',	'Serf1',	'Erbin',	'Kif2a',	'Il6st',	'BC067074',	'Parp8',	'Emb',	'Paip1',	'Hmgcs1',	'Nim1k',	'Rab10',	'Kif3c',	'Ncoa1',	'Klhl29',	'Apob',	'Ldah',	'Hs1bp3',	'Pum2',	'Laptm4a',	'Ttc32',	'Smc6',	'Rock2',	'Pqlc3',	'Cpsf3',	'Iah1',	'Adam17',	'Taf1b',	'Cys1',	'Mboat2',	'Kidins220',	'Rnf144a',	'Rsad2',	'Tpo',	'Ccdc71l',	'Nampt',	'Snx13',	'Agmo',	'Arl4a',	'Lsmem1',	'Zfp277',	'Immp2l',	'Dnajb9',	'Stxbp6',	'Sptssa',	'Ppp2r3c',	'Fkbp3',	'Rpl36al',	'Klhdc2',	'Vcpkmt',	'Sos2',	'Abhd12b',	'Actr10',	'Arid4a',	'Mthfd1',	'Zbtb25',	'Max',	'Gphn',	'Plekhh1',	'Arg2',	'Psen1',	'Pnma1',	'Elmsan1',	'Rnf113a2',	'Aldh6a1',	'Zc2hc1c',	'Tmed10',	'Fos',	'Batf',	'Erg28',	'Adck1','Ston2','Fbln5',	'Rin3',	'Lgmn',	'Chga',	'Itpk1',	'Ifi27',	'Ppp4r4',	'Bdkrb2',	'Gskip',	'Ccdc85c',	'Slc25a29',	'Slc25a47',	'Wdr25',	'Ppp2r5c',	'Hsp90aa1',	'Zfp839',	'Cinp',	'Tecpr2',	'Traf3',	'Mark3',	'Trmt61a',	'Bag5',	'Ppp1r13b',	'Adssl1',	'Siva1',	'Zbtb42',	'Cdca4',	'Tex22',	'Mta1',	'Crip2',	'Crip1',	'Ccdc152',	'Fbxo4',	'Oxct1',	'Rictor',	'Egflam',	'Gdnf',	'Dnajc21',	'1810049J17Rik',	'Zfp622',	'Fbxl7',	'Dap',	'Vps13b',	'Spag1',	'Rrm2b',	'Azin1',	'Atp6v1c1',	'Abra',	'Eif3e',	'Eny2',	'Col14a1',	'Sntb1',	'Zhx2',	'Derl1',	'9130401M01Rik',	'Fam91a1',	'Rnf139',	'Washc5',	'Trib1',	'Efr3a',	'Ago2',	'Dennd3',	'Gpr20',	'Zc3h3',	'Tigd5',	'Pycrl',	'Scrib',	'Nrbp2',	'Eppk1',	'BC024139',	'Plec',	'Parp10',	'Smpd5',	'Hgh1',	'Bop1',	'Fbxl6',	'Kifc2',	'Ppp1r16a',	'1110038F14Rik',	'Apol6',	'Apol8',	'Sstr3',	'Triobp',	'Micall1',	'Polr2f',	'Pla2g6',	'Maff',	'Tmem184b',	'Kdelr3',	'Sun2',	'Dnal4',	'Apobec3',	'Pdgfb',	'Chadl',	'Tef',	'Polr3h',	'Desi1',	'Naga',	'Ndufa6',	'Cyp2d22',	'Pacsin2',	'Ttll1',	'Bik',	'Mcat',	'Ttll12',	'Rtl6',	'5031439G07Rik',	'Wnt7b',	'Ppara',	'Alg12',	'Creld2',	'Pim3',	'Plxnb2',	'C730034F03Rik',	'Mapk8ip2',	'Gxylt1',	'Pus7l',	'Twf1',	'Endou',	'Rapgef3',	'Slc48a1',	'Hdac7',	'Asb8',	'Cacnb3',	'Ccdc65',	'Kmt2d',	'Tmbim6',	'Nckap5l',	'Faim2',	'Aqp2',	'Aqp5',	'Racgap1',	'Smarcd1',	'Gpd1',	'Mettl7a1',	'Gm5475',	'Fignl2',	'Nr4a1',	'Krt80',	'Tns2',	'Soat2',	'Zfp740',	'Sp1',	'Itga5',	'Gtsf1',	'Dnaja3',	'Hmox2',	'4930562C15Rik',	'Ubald1'
                   ,	'Mgrn1',	'Nudt16l1',	'AU021092',	'Gm5767',	'Emp2',	'Litaf',	'Zc3h7a',	'Rsl1d1',	'Cpped1',	'Parn',	'Pla2g10',	'Pdxdc1',	'Nde1',	'Cebpd',	'Mapk1',	'Ppil2',	'Scarf2',	'Dgcr6',	'Prodh',	'Hira',	'Camk2n2',	'Ehhadh',	'Tmem41a',	'Liph',	'Igf2bp2',	'Atp13a4',	'Pak2',	'Pcyt1a',	'Fyttd1',	'Heg1',	'Muc13',	'Adcy5',	'Sec22a',	'Iqcb1',	'Lrrc58',	'Gsk3b',	'Atp6v1a',	'Usf3',	'Spice1',	'Slc35a5',	'Abhd10',	'Cd47',	'Cep97',	'Zbtb11',	'Olfr172',	'Riox2',	'4930453N24Rik',	'Usp25',	'Btg3',	'App',	'Map3k7cl',	'Tiam1',	'Sod1',	'Eva1c',	'Synj1',	'Paxbp1',	'Ifnar2',	'Ifnar1',	'Ifngr2',	'Tmem50b',	'Dnajc28',	'Itsn1',	'Kcne2',	'Clic6',	'Brwd1',	'Ripk4',	'Zdhhc14',	'Serac1',	'Sytl3',	'Ezr',	'AC122413.2',	'Sft2d1',	'Prr18',	'Agpat4',	'Igf2r',	'Acat3',	'Wtap',	'Afdn',	'Gm3448',	'Chd1',	'Lnpep',	'Zfp948',	'Zfp943',	'Pdpk1',	'Amdhd2',	'Tbc1d24',	'Caskin1',	'Nthl1',	'Npw',	'Eme2',	'Tekt4',	'Rab40c',	'Capn15',	'Tmem8',	'Dusp1',	'Atp6v0e',	'Crebrf',	'Bnip1',	'Lemd2',	'AI413582',	'Taf11',	'Zfp523',	'Ppard',	'Fkbp5',	'Mapk14',	'Kctd20',	'Stk38',	'Cdkn1a',	'BC004004',	'Gm28043',	'Slc37a1',	'Wdr4',	'Rrp1b',	'Cyp4f13',	'Adamts10',	'Kank3',	'Ndufa7',	'H2-K1',	'Tap1',	'Gm15821',	'H2-Ab1',	'Notch4',	'Pbx2',	'Atf6b',	'Neu1',	'H2-D1',	'H2-Q2',	'Ddr1',	'Ier3',	'Atat1',	'Trim15',	'Rnf39',	'Cyp39a1',	'Enpp4',	'Tmem63b',	'Gtpbp2',	'Polr1c',	'Zfp318',	'Cul9',	'Gnmt',	'1700001C19Rik',	'Tomm6',	'Pgc',	'Kif6',	'Tbc1d5',	'Zfp119b',	'Stap2',	'Sema6b',	'Dpp9',	'Nrtn',	'Prr22',	'Rfx2',	'1700061G19Rik',	'Mllt1',	'Vav1',	'Fbxl17',	'A930002H24Rik',	'Pja2',	'Ppp4r1',	'Twsg1',	'Ankrd12',	'Washc1',	'Mtcl1',	'Arhgap28',	'Myl12a',	'Emilin2',	'Clip4',	'Ypel5',	'Xdh',	'Ttc27',	'Fam98a',	'Strn',	'Prkd3',	'Atl2',	'Hnrnpll',	'Sos1',	'Cox7a2l',	'Dync2li1',	'Lrpprc',	'Prkce',	'Ttc7',	'Ston1',	'Cul2',	'Arhgap12',	'Kif5b',	'Mpp7',	'Wac',	'Usp14',	'Rock1',	'Mib1',	'Osbpl1a',	'Ss18',	'Kctd1',	'Rnf138',	'Sft2d3',	'Reep5',	'Cdc23',	'Egr1',	'Paip2',	'Ecscr',	'Ube2d2a',	'Cxxc5',	'Pcdhga2',	'Pcdhgb5',	'Pcdhga10',	'Rnf14',	'Nr3c1',	'Trim36',	'Snx2',	'Snx24',	'Csnk1g3',	'Tex43',	'Iigp1',	'Synpo',	'Ndst1',	'Rps14',	'Cd74',	'Csf1r',	'Hmgxb3',	'Pde6a',	'Arhgef37',	'Pcyox1l',	'Grpel2',	'Ablim3',	'Adrb2',	'Fbxo38',	'Fech',	'Nars',	'Nedd4l',	'Zfp532',	'Cidea',	'Afg3l2',	'Psmg2',	'Ldlrad4',	'Mbd2',	'Ctif',	'Zbtb7c',	'Hdhd2',	'Adnp2',	'Pqlc1',	'Ctdp1',	'Mbp',	'Cndp2',	'Cyb5a',	'Cpt1a',	'Tesmin',	'Lrp5',	'Chka',	'Unc93b1',	'Cdk2ap2',	'Coro1b',	'Pold4',	'Rhod',	'2010003K11Rik',	'Pcx',	'Lrfn4',	'Rin1',	'Cd248',	'Mus81',	'Snx32',	'Ap5b1',	'Kcnk7',	'Ehbp1l1',	'Sssca1',	'Ltbp3',	'Majin',	'Cdc42bpg',	'Slc22a12',	'Ccdc88b',	'Gpr137',	'Cox8a',	'Slc3a2',	'Lrrn4cl',	'B3gat3',	'Rom1',	'Fth1',	'Best1',	'Myrf',	'Cpsf7',	'Vps37c',	'Ms4a8a',	'Stx3',	'Carnmt1',	'D030056L22Rik',	'Trpm6',	'Rorb',	'Klf9',	'Fam189a2',	'Smarca2',	'Vldlr',	'Pum3',	'Glis3',	'Rcl1',	'Jak2',	'Mlana',	'Uhrf2',	'Pcgf5',	'43895',	'Ccnj',	'Lcor',	'Rrp12',	'Exosc1',	'Ankrd2',	'Pi4k2a',	'Sfrp5',	'R3hcc1l',	'Pyroxd2',	'Got1',	'Erlin1',	'Bloc1s2',	'Scd1',	'Lzts2',	'Gm17018',	'Fbxw4',	'Mgea5',	'Hps6',	'Ldb1',	'Sufu',	'Trim8',	'Wbp1l',	'Cyp17a1',	'Ina',	'Calhm2',	'Sh3pxd2a',	'Sfr1',	'Vti1a',	'Casp7',	'Nhlrc2',	'Ccdc186',	'Vwa2',	'Fam160b1',	'Atrnl1',	'Shtn1',	'Pdzd8',	'Cacul1',	'Zfp950'))

integrate <- AddModuleScore(object = integrate, features = BACClean, ctrl = 500, name = "NKXBAcquiredCleanGenes", search = TRUE, random.seed = 1)
FeaturePlot(object = integrate, features = 'NKXBAcquiredCleanGenes1', split.by = "ID", cols = c('lightgrey','red'))
FeaturePlot(object = integrate, features = 'NKXBAcquiredCleanGenes1', cols = c('lightgrey','red'))


BRClean <- list(c('Cops5',	'Arfgef1',	'Ncoa2',	'Tram1',	'Lactb2',	'Eya1',	'Msc',	'Terf1',	'Ly96',	'Col9a1',	'Ptp4a1',	'Ccdc115',	'Fam178b',	'Inpp4a',	'2010300C02Rik',	'Lipt1',	'Lyg2',	'Rev1',	'Aff3',	'Pdcl3',	'Map4k4',	'Mrps9',	'AI597479',	'Nck2',	'Uxs1',	'Dnah7b',	'Stat4',	'Hibch',	'Pgap1',	'Plcl1',	'Satb2',	'Gm973',	'Fam117b',	'Nrp2',	'Idh1',	'Lancl1',	'Spag16',	'Xrcc5',	'Rpl37a',	'Igfbp5',	'Catip',	'Cnot9',	'Zfp142',	'Wnt10a',	'Ihh',	'Epha4',	'Acsl3',	'Irs1',	'Rhbdd1',	'Daw1',	'A630001G21Rik',	'Armc9',	'Ptma',	'Spp2',	'Glrp1',	'Gbx2',	'Ackr3',	'Col6a3',	'Ube2f',	'Asb1',	'Hdac4',	'Gpc1',	'Crocc2',	'Stk25',	'Bok',	'Pdcd1',	'Slco4c1',	'Pam',	'Phlpp1',	'Gli2',	'Steap3',	'Actr3',	'Mcm6',	'Cd55b',	'Zp3r',	'Pigr',	'Dyrk3',	'Rassf5',	'Ikbke',	'Slc26a9',	'Lax1',	'Optc',	'Fmod',	'Btg2',	'Cyb5r1',	'Ube2t',	'Lmod1',	'Tmem9',	'Cacna1s',	'Kif14',	'Ptprc',	'Dennd1b',	'Trove2',	'Hmcn1',	'Ivns1abp',	'Ncf2',	'Smg7',	'Shcbp1l',	'Dhx9',	'Npl',	'Rnasel',	'Cacna1e',	'Ier5',	'BC034090',	'Abl2',	'Fam20b',	'Ralgps2',	'Pappa2',	'4930558K02Rik',	'Mettl11b',	'Xcl1',	'Uck2',	'Aldh9a1',	'Lrrc52',	'Fcgr4',	'Fcgr3',	'Tomm40l',	'Alyref2',	'Cfap45',	'Cadm3',	'Fmn2',	'Grem2',	'Pld5',	'Rbm8a2',	'Akt3',	'1700016C15Rik',	'Kif26b',	'Smyd3',	'Nvl',	'Lbr',	'Srp9',	'Bpnt1',	'Kcnk2',	'Cenpf',	'Ptpn14',	'Smyd2',	'Prox1',	'Sertad4',	'G0s2',	'Camk1g',	'Cd46',	'Cr1l',	'Nmt2',	'Acbd7',	'Camk1d',	'Proser2',	'Gata3',	'Taf3',	'Itih5',	'Sfmbt2',	'Trdmt1',	'Pdss1',	'Psd4',	'Dph7',	'Nrarp',	'Ccdc187',	'Sec16a',	'Wdr5',	'Fcnb',	'Mrps2',	'Spaca9',	'Cfap77',	'Med27',	'Set',	'Ptpa',	'Ier5l',	'Cstad',	'Ncs1',	'Ass1',	'Exosc2',	'Fam102a',	'Dpm2',	'Ak1',	'Stom',	'Ndufa8',	'Nr6a1',	'Arhgap15',	'Acvr2a',	'Mbd5',	'Epc2',	'Rnd3',	'Arl5a',	'Stam2',	'Fmnl2',	'Prpf40a',	'Rprm',	'Dpp4',	'Ifih1',	'Kcnh7',	'Fign',	'Slc38a11',	'Scn3a',	'Scn2a',	'Stk39',	'Myo3b',	'Gad1',	'Cybrd1',	'Metap1d',	'Dlx2',	'Rapgef4',	'Sp9',	'Wipf1',	'Lnpk',	'Nfe2l2',	'Ttc30a1',	'Rbm45',	'Cwc22',	'Ube2e3',	'Prdx6b',	'Zfp804a',	'Tfpi',	'Zdhhc5',	'Gm19426',	'Prg3',	'Ptprj',	'Arfgap2',	'1110051M20Rik',	'Lrp4',	'Phf21a',	'Accs',	'Api5',	'B230118H07Rik',	'Prr5l',	'Commd9',	'Abtb2',	'Caprin1',	'A930018P22Rik',	'Qser1',	'Them7',	'Rcn1',	'Pax6',	'Mpped2',	'Arl14ep',	'Kif18a',	'Lin7c',	'Arhgap11a',	'Rasgrp1',	'Fsip1',	'Bmf',	'Bub1b',	'Chst14',	'Ccdc32',	'Rpusd2',	'Knl1',	'Rad51',	'Dnajc17',	'Chp1',	'Oip5',	'Itpka',	'Rpap1',	'Capn3',	'Zfp106',	'Ubr1',	'Zscan29',	'Pdia3',	'Mfap1b',	'Mfap1a',	'Wdr76',	'Shf',	'AA467197',	'1810024B03Rik',	'Kcnip3',	'Mall',	'Anapc1',	'Mertk',	'Zc3h8',	'Gm14025',	'Ptpra',	'Cenpb',	'Pank2',	'Pcna',	'Chgb',	'Bmp2',	'Btbd3',	'Macrod2',	'Rbbp9',	'Dtd1',	'Insm1',	'Nkx2-2',	'Foxa2',	'Zfp442',	'Acss1',	'Rspo4',	'Csnk2a1',	'Zcchc3',	'Id1',	'Tpx2',	'Plagl2',	'Dnmt3b',	'Snta1',	'Cbfa2t2',	'E2f1',	'Gm14226',	'Ahcy',	'Itch',	'Edem2',	'Soga1',	'Rpn2',	'Blcap',	'Vstm2l',	'Lbp',	'Snhg11',	'Emilin3',	'Chd6',	'Jph2',	'Tomm34',	'Sys1',	'Ncoa5',	'Zfp663',	'Zmynd8',	'Ncoa3',	'Trp53rkb',	'Znfx1',	'Slc9a8',	'Pard6b',	'Zfp217',	'Bcas1',	'Rae1',	'Ankrd60',	'Zfp831',	'Edn3',	'Zfp931',	'Cdh26',	'Rps21',	'Rbbp8nl',	'Slco4a1',	'Mrgbp',	'Fndc11',	'Sox18',	'Shroom4',	'Usp27x',	'Usp9x',	'Gpr82',	'Maoa',	'Jade3',	'Uba1',	'Ube2a',	'Nkap',	'Ocrl',	'Bcorl1',	'Enox2',	'Mbnl3',	'Gpc4',	'Gpc3',	'Hprt',	'Plac1',	'Zfp449',	'1700020N15Rik',	'Cd99l2',	'Flna',	'Rpl10',	'Arx',	'Pcyt1b',	'Maged1',	'Yipf6',	'Stard8',	'Kif4',	'Gdpd2',	'Slc7a3',	'Snx12',	'Ogt',	'Rtl5',	'Pin4',	'Ercc6l',	'Hdac8',	'Tspan6',	'Cstf2',	'Rpl36a',	'Armcx1',	'Armcx5',	'Bhlhb9',	'Bex4',	'Tceal8',	'Tceal9',	'Tceal3',	'Rnf128',	'Kcne1l',	'Pak3',	'Capn6',	'Rps6ka3',	'Tceanc',	
                  'Egfl6',	'Tmsb4x',	'Arhgap6',	'Gm21887',	'Gm8797',	'Mrps28',	'Pag1',	'Ythdf3',	'Pde7a',	'Tnfsf10',	'Fndc3b',	'Pld1',	'Mynn',	'Actl6a',	'Ccdc39',	'Fxr1',	'Sox2',	'Atp11b',	'D3Ertd254e',	'Slc7a11',	'Maml3',	'Cog6',	'Stoml3',	'Frem2',	'Commd2',	'Pfn2',	'Tsc22d2',	'Mbnl1',	'Rap2b',	'Plch1',	'Shox2',	'Ift80',	'Arl14',	'Nmd3',	'Etfdh',	'Tmem144',	'Fam198b',	'Pdgfc',	'Ctso',	'Gucy1b1',	'Trim2',	'Gm6525',	'Fhdc1',	'Glt28d2',	'Kirrel',	'Etv3',	'Prcc',	'Hdgf',	'Rrnad1',	'Nes',	'Arhgef2',	'Syt11',	'Clk2',	'Thbs3',	'Trim46',	'Krtcap2',	'Efna4',	'Flad1',	'Kcnn3',	'Hax1',	'Tpm3',	'Crtc2',	'Ints3',	'Ilf2',	'Rptn',	'Rfx5',	'Psmd4',	'Vps72',	'Scnm1',	'Cdc42se1',	'Setdb1',	'Hormad1',	'Tars2',	'Rprd2',	'Bola1',	'Hist2h2aa2',	'Hist2h4',	'Fcgr1',	'Hfe2',	'Ttf2',	'Cd2',	'Igsf3',	'Vangl1',	'Sycp1',	'Nras',	'Trim33',	'Rsbn1',	'Magi3',	'Ddx20',	'Adora3',	'Kcnc4',	'Gstm7',	'Atxn7l2',	'Psrc1',	'5330417C22Rik',	'Taf13',	'Cdc14a',	'Rtca',	'Dbt',	'Agl',	'Alg14',	'Cnn3',	'Arhgap29',	'Synpo2',	'Elovl6',	'Egf',	'Ostc',	'Cyp2u1',	'Papss1',	'Aimp1',	'Cenpe',	'Bdh2',	'Gm21962',	'Bmpr1b',	'Pdlim5',	'Gtf2b',	'Lmo4',	'Sh3glb1',	'Znhit6',	'Cyr61',	'Ssx2ip',	'Adgrl2',	'Dnajb4',	'Miga1',	'St6galnac3',	'Erich3',	'Wls',	'Gm11808',	'Penk',	'Fam110b',	'Car8',	'Rbm12b2',	'Manea',	'Akirin2',	'Aco1',	'Ubap2',	'Ubap1',	'Enho',	'Fam166b',	'Trmt10b',	'Ncbp1',	'Anks6',	'Tex10',	'Rnf20',	'Fsd1l',	'Rad23b',	'D630039A03Rik',	'Inip',	'Zfp37',	'Cdc26',	'Wdr31',	'Bspry',	'4933430I17Rik',	'Rgs3',	'Kif12',	'Megf9',	'Tle1',	'Lurap1l',	'Mpdz',	'Nfib',	'Bnc2',	'Adamtsl1',	'Hacd4',	'Klhl9',	'Mtap',	'Fggy',	'E130114P18Rik',	'Nfia',	'Dock7',	'Pgm2',	'Ube2u',	'Cachd1',	'Raver2',	'Ak4',	'Sgip1',	'Tctex1d1',	'Plpp3',	'Fam151a',	'Acot11',	'Ssbp3',	'Lrrc42',	'Elavl4',	'Spata6',	'Cmpk1',	'Tex38',	'Nsun4',	'Uqcrh',	'Rad54l',	'Tmem69',	'Nasp',	'Kif2c',	'Tmem53',	'Cdc20',	'Tmem125',	'Cfap57',	'Slc2a1',	'Zfp691',	'Cldn19',	'Ccdc30',	'Scmh1',	'Mycl',	'Pou3f1',	'9930104L06Rik',	'Dnali1',	'Snip1',	'Ago1',	'Clspn',	'5730409E04Rik',	'Zmym4',	'Sfpq',	'Gja4',	'Gjb3',	'Tmem54',	'Zbtb8os',	'Kpna6',	'Khdrbs1',	'Gm10300',	'Taf12',	'Rab42',	'Med18',	'Smpdl3b',	'Rpa2',	'Ppp1r8',	'Ahdc1',	'Wasf2',	'Cd164l2',	'Tmem222',	'Kdf1',	'Hmgn2',	'Lin28a',	'Pdik1l',	'Pafah2',	'Ldlrap1',	'Rhd',	'Nipal3',	'Fuca1',	'E2f2',	'Hnrnpr',	'C1qa',	'Zbtb40',	'Alpl',	'Pla2g2d',	'Pla2g5',	'Nbl1',	'Pax7',	'Arhgef10l',	'Rcc2',	'Padi6',	'Sdhb',	'Rsg1',	'Arhgef19',	'Clcnka',	'Casp9',	'Tmem51',	'Kazn',	'Cenps',	'Rbp7',	'Eno1',	'Vamp3',	'Ttc34',	'Mmel1',	'Plch2',	'Ski',	'Prkcz',	'C1qtnf12',	'Ttll10',	'9430015G10Rik',	'Cdk6',	'Cyp51',	'Sema3e',	'Pclo',	'Sema3c',	'Gnai1',	'Ptpn12',	'Fgl2',	'Kcnh2',	'Chpf2',	'Actr3b',	'Rnf32',	'Dpysl5',	'Agbl5',	'Cad',	'Ift172',	'Ywhah',	'Spon2',	'Uvssa',	'Slbp',	'Ablim2',	'Sorcs2',	'Grpel1',	'Cpeb2',	'Bst1',	'Fgfbp1',	'Tapt1',	'Slit2',	'Adgra3',	'Slc34a2',	'Tbc1d19',	'0610040J01Rik',	'Klf3',	'Klhl5',	'Ugdh',	'Smim14',	'Chrna9',	'9130230L23Rik',	'Limch1',	'Phox2b',	'Gm5108',	'Nfxl1',	'Lrrc66',	'Spata18',	'Rasl11b',	'Lnx1',	'Tmem165',	'Aasdh',	'Slc4a4',	'Gc',	'Npffr2',	'Afp',	'Rassf6',	'Mthfd2l',	'Epgn',	'Areg',	'Shroom3',	'Sowahb',	'2010109A12Rik',	'Fras1',	'4930524J08Rik',	'Hnrnpdl',	'Gpat3',	'Lrrc8c',	'Zfp644',	'Cdc7',	'Tgfbr3',	'Gfi1',	'Ube2d2b',	'Mtf2',	'Slc26a1',	'Crlf2',	'Gm15446',	'Zfp932',	'Plcxd1',	'Hscb',	'Sez6l',	'Myo18b',	'Sart3',	'Ssh1',	'Mmab',	'Gm10401',	'Dynll1',	'Pla2g1b',	'Sirt4',	'Pxn',	'Rplp0',	'Vsig10',	'Wsb2',	'Rfc5',	'Rnft2',	'2410131K14Rik',	'Med13l',	'Tmem116',	'Acad12',	'Cux2',	'B3gnt4',	'Kntc1',	'Arl6ip4',	'Pitpnm2',	'Cdk2ap1',	'Sbno1',	'Tmed2',	'Tmem132c',	'Galnt17',
                  'Rcc1l',	'Rfc2',	'Eln',	'Cldn3',	'Abhd11',	'Mlxipl',	'Baz1b',	'Rhbdd2',	'Ssc4d',	'Rasa4',	'Polr2j',	'Myl10',	'Col26a1',	'Fis1',	'Plod3',	'Ap1s1',	'Trim56',	'Ufsp1',	'Ephb4',	'Tsc22d4',	'Gm5294',	'Lfng',	'Gna12',	'Foxk1',	'Slc29a4',	'Fscn1',	'Fam220a',	'Nptx2',	'Pomp',	'Mtus2',	'Alox5ap',	'Medag',	'Tex26',	'Wdr95',	'B3glct',	'N4bp2l2',	'Kl',	'Asb4',	'Dync1i1',	'Dlx5',	'Asns',	'Mios',	'Glcci1',	'Ica1',	'Vwde',	'2610001J05Rik',	'St7',	'Wnt2',	'Asz1',	'Fam3c',	'Atp6v1f',	'Tnpo3',	'Tspan33',	'Zc3hc1',	'Klhdc10',	'Cpa4',	'Exoc4',	'Dgki',	'Creb3l2',	'Atp6v0a4',	'Tmem213',	'Ttc26',	'Rab19',	'Arhgef5',	'Ezh2',	'Pdia4',	'Igf2bp3',	'Mpp6',	'Gsdme',	'Nfe2l3',	'Snx10',	'Skap2',	'Creb5',	'Tril',	'Chn2',	'Ghrhr',	'Nap1l5',	'Fam13a',	'Il12rb2',	'Rpia',	'Mrpl35',	'Ptcd3',	'St3gal5',	'Usp39',	'Tmem150a',	'Ggcx',	'Capg',	'Tcf7l1',	'Kcmf1',	'Dnah6',	'Sema4f',	'Wbp1',	'Zfp638',	'Cyp26b1',	'Noto',	'Nat8f4',	'Fam136a',	'Snrpg',	'Tia1',	'Anxa4',	'Nfu1',	'Gfpt1',	'Gkn3',	'Mgll',	'Iqsec1',	'4930590J08Rik',	'Slc25a26',	'Lrig1',	'Fam19a4',	'Lmod3',	'Foxp1',	'Lmcd1',	'Sec13',	'Tsen2',	'Plxnd1',	'Washc2',	'Zfp422',	'Ret',	'Zfp9',	'Dcp1b',	'Fbxl14',	'B4galnt3',	'D330020A13Rik',	'Hdhd5',	'Phc1',	'Rimklb',	'Emg1',	'Phb2',	'Grcc10',	'Usp5',	'Ing4',	'Tnfrsf1a',	'Vwf',	'Ntf3',	'Ccnd2',	'Parp11',	'Klrb1c',	'Klrb1b',	'Ybx3',	'Etv6',	'Mansc1',	'Pbp2',	'Emp1',	'Grin2b',	'Gm8994',	'Atf7ip',	'Plbd1',	'Hist4h4',	'Eps8',	'Mgst1',	'Pik3c2g',	'Slco1c1',	'Cmas',	'Sox5',	'Arntl2',	'Klhl42',	'Pthlh',	'Caprin2',	'Eps8l1',	'Hspbp1',	'Kmt5c',	'Zfp667',	'Zfp418',	'Rnf225',	'Zfp324',	'Zbtb45',	'Chmp2a',	'Slc8a2',	'Prkd2',	'Calm3',	'Mypop',	'Irf2bp1',	'Six5',	'Fosb',	'Exoc3l2',	'Relb',	'Clptm1',	'Bcam',	'Zfp61',	'Lypd3',	'Zfp526',	'Erf',	'Cic',	'Pafah1b3',	'Megf8',	'Ccdc97',	'Rab4b',	'Mia',	'Map3k10',	'Zfp607b',	'Zfp626',	'Hnrnpl',	'Ech1',	'Capn12',	'Zfp790',	'Zfp566',	'Cox7a1',	'E130208F15Rik',	'Lrfn3',	'Nfkbid',	'U2af1l4',	'Zbtb32',	'Upk1a',	'Lgi4',	'Pdcd2l',	'Kctd15',	'Lrp3',	'Zfp536',	'Uri1',	'Ccne1',	'1600014C10Rik',	'Syt3',	'Emc10',	'Nr1h2',	'Cpt1c',	'Prmt1',	'Scaf1',	'Slc17a7',	'Ccdc155',	'Bax',	'Ppp1r15a',	'Mamstr',	'Cyth2',	'Abcc6',	'Kcnj11',	'Kcnc1',	'Spty2d1',	'Ptpn5',	'E2f8',	'Mtmr10',	'Tm2d3',	'Lrrk1',	'Asb7',	'Cers3',	'Adamts17',	'Fam169b',	'Arrdc4',	'Gm10295',	'Mctp2',	'St8sia2',	'Blm',	'3110040N11Rik',	'Mesd',	'Arnt2',	'Rab38',	'Eed',	'Ccdc90b',	'Nars2',	'Kctd14',	'Myo7a',	'Emsy',	'Thap12',	'Wnt11',	'Uvrag',	'Serpinh1',	'Kcne3',	'C2cd3',	'Plekhb1',	'Stard10',	'Phox2a',	'Rnf121',	'Trim34a',	'Tub',	'Ascl3',	'Zfp143',	'Sbf2',	'Usp47',	'Dkk3',	'Spon1',	'Rras2',	'Sox6',	'1110004F10Rik',	'Iqck',	'Acsm4',	'Rexo5',	'Mosmo',	'Scnn1g',	'Rbbp6',	'Tnrc6a',	'Il21r',	'Mapk3',	'Hirip3',	'Cdipt',	'Pagr1a',	'Cd2bp2',	'E430018J23Rik',	'Bcl7c',	'Orai3',	'Zfp646',	'Cox6a2',	'Sec23ip',	'Ate1',	'Tacc2',	'Plekha1',	'Edrf1',	'Adam12',	'Ptpre',	'Mki67',	'Ebf3',	'Tcerg1l',	'Inpp5a',	'Cfap46',	'Lrrc56',	'Pidd1',	'Ctsd',	'Syt8',	'Ins2',	'Tnfrsf22',	'Mrgprd',	'Iyd',	'Armt1',	'Ust',	'Stx11',	'Phactr2',	'Aig1',	'Cited2',	'Perp',	'Map7',	'1700020N01Rik',	'L3mbtl3',	'Arhgap18',	'Cenpw',	'Ncoa7',	'Hddc2',	'Frk',	'Hdac2',	'Marcks',	'Fam229b',	'Traf3ip2',	'Rpf2',	'Gtf3c6',	'Cdk19',	'Mettl24',	'Cdc40',	'Cd164',	'Afg1l',	'Cd24a',	'Prdm1',	'Dcbld1',	'Slc35f1',	'Pkib',	'Anapc16',	'Chst3',	'Psap',	'Gm17455',	'Unc5b',	'Ppa1',	'H2afy2',	'Col13a1',	'Dna2',	'Hnrnph3',	'Pbld2',	'Pbld1',	'Reep3',	'Zfp365',	'Cabcoco1',	'Cdk1',	'Ccdc6',	'Rsph14',	'Rab36',	'Mif',	'Gm5134',	'Zfp280b',	'Prmt2',	'Pofut2',	'Plpp2',	'Cnn2',	'Sbno2',	'Stk11',	'Ndufs7',	'Reep6',	'Tcf3',	'Slc39a3',	'Eef2',	'Nmrk2',
                  'Matk',	'Tjp3',	'Tle2',	'Zfp873',	'Gm4924',	'Nfyb',	'Nuak1',	'Ric8b',	'Mterf2',	'Cry1',	'Ascl4',	'Fbxo7',	'Stab2',	'Gnptab',	'Sycp3',	'Spic',	'Gas2l3',	'Anks1b',	'Tmpo',	'Cfap54',	'Lta4h',	'Snrpf',	'Cradd',	'Dcn',	'4930430F08Rik',	'Nts',	'Ptprq',	'Otogl',	'Pawr',	'E2f7',	'Bbs10',	'Nap1l1',	'Phlda1',	'Ptprb',	'Cnot2',	'Myrfl',	'Grip1',	'Hmga2',	'Usp15',	'Lrig3',	'Avil',	'43899',	'Kif5a',	'Gli1',	'Ndufa4l2',	'Nab2',	'Atp5b',	'Timeless',	'Pan2',	'Cnpy2',	'Cs',	'Zc3h10',	'Rpl41',	'Pa2g4',	'Erbb3',	'Rps26',	'Ikzf4',	'Dgka',	'Pym1',	'Dnajc14',	'Mettl7b',	'Insr',	'Xab2',	'Ccl25',	'Cers4',	'Efnb2',	'Myo16',	'Rab20',	'Ankrd10',	'Pcid2',	'Cdc16',	'Kbtbd11',	'Ckap2',	'Vps36',	'Polb',	'Ap3m2',	'Kat6a',	'Zmat4',	'Plekha2',	'Plpbp',	'Rnf122',	'Fut10',	'Tex15',	'Gtf2e2',	'Ppp1r3b',	'Prag1',	'Micu3',	'Mtus1',	'Pcm1',	'Frg1',	'Mtnr1a',	'F11',	'Tlr3',	'Sorbs2',	'Pdlim3',	'Lrp2bp',	'Primpol',	'Casp3',	'Sap30',	'Mfap3l',	'Clcn3',	'Nek1',	'Cbr4',	'Apela',	'Naf1',	'Psd3',	'Sh2d4a',	'Atp6v1b2',	'Zfp930',	'Ddx49',	'Crlf1',	'Jund',	'Kcnn1',	'Abhd8',	'Fcho1',	'Large1',	'Mcm5',	'Tmem184c',	'Zfp827',	'Smad1',	'Smarca5',	'Il15',	'Tbc1d9',	'Scoc',	'Adgrl1',	'1700067K01Rik',	'Ier2',	'Lyl1',	'Rad23a',	'Tnpo2',	'Neto2',	'Adcy7',	'Rbl2',	'Fto',	'Irx3',	'Amfr',	'Arl2bp',	'Pllp',	'Polr2c',	'Dok4',	'Adgrg1',	'Cngb1',	'Tepp',	'Gins3',	'Cklf',	'Nae1',	'Cdh16',	'B3gnt9',	'Slc9a5',	'Zdhhc1',	'Enkd1',	'Cenpt',	'Psmb10',	'Slc12a4',	'Esrp2',	'Slc7a6',	'Smpd3',	'Cdh3',	'Cdh1',	'Vps4a',	'Psmd7',	'Zfhx3',	'Vac14',	'Mlkl',	'Fa2h',	'Cfdp1',	'Adat1',	'Gm6793',	'Mon1b',	'Wwox',	'Cdh13',	'Fam92b',	'Klhdc4',	'Slc7a5',	'Mvd',	'Galnt2',	'Pcnx2',	'Kcnk1',	'Slc35f3',	'Pard3',	'Fam107a',	'4930452B06Rik',	'Thrb',	'Kcnk5',	'Nudt13',	'Vdac2',	'Zfp503',	'Dlg5',	'Arf4',	'Nt5dc2',	'Sh3bp5',	'Ankrd28',	'Dph3',	'Oxnad1',	'Arhgap22',	'Sftpa1',	'Gm15217',	'Cdkn3',	'Tmem260',	'Klhl33',	'Rnase12',	'Rnase2a',	'Ang4',	'Mettl17',	'Ndrg2',	'Sall2',	'Lrp10',	'Cdh24',	'Slc7a8',	'Cmtm5',	'Ngdn',	'Zfhx2',	'Thtpa',	'Nynrin',	'Mcpt8',	'Parp4',	'Pspc1',	'Zmym5',	'Zmym2',	'Fgf9',	'Spata13',	'Mipep',	'Tnfrsf19',	'Arl11',	'Wdfy2',	'Pinx1',	'Msra',	'Kif13b',	'Pnoc',	'Scara5',	'Pbk',	'Ccdc25',	'Nefm',	'Nefl',	'Adam7',	'Nkx3-1',	'R3hcc1',	'Pebp4',	'Npm2',	'Dok2',	'Lpar6',	'Lcp1',	'Cog3',	'Serp2',	'Naa16',	'Kbtbd6',	'Slain1',	'Ednrb',	'Spry2',	'Slitrk1',	'Gpc5',	'Sox21',	'Cldn10',	'Rap2a',	'Dock9',	'Timm8a2',	'Tmtc4',	'Alkbh8',	'Gucy1a2',	'Dync2h1',	'Trpc6',	'Sesn3',	'Endod1',	'Cep295',	'Ubl5',	'Dnmt1',	'Mrpl4',	'Icam4',	'Fdx1l',	'Ap1m2',	'Qtrt1',	'Ldlr',	'Rab3d',	'Glb1l2',	'Jam3',	'Igsf9b',	'Opcml',	'Ntm',	'Snx19',	'Aplp2',	'Kcnj5',	'Kirrel3',	'Cdon',	'Pus3',	'Stt3a',	'Slc37a2',	'Nrgn',	'Scn3b',	'Hspa8',	'Sc5d',	'Tecta',	'Arhgef12',	'Pou2f3',	'Vps11',	'Atp5l',	'Ube4a',	'Jaml',	'Cep164',	'Pcsk7',	'Tagln',	'Pafah1b2',	'Sik3',	'Apoa1',	'Sik2',	'Colca2',	'1810046K07Rik',	'Ddx10',	'Exph5',	'Kdelc2',	'4930550C14Rik',	'Rcn2',	'Tspan3',	'Lingo1',	'Snupn',	'Sin3a',	'Ppcdc',	'Edc3',	'Arid3b',	'Pml',	'Cd276',	'Nptn',	'Pkm',	'Senp8',	'Thsd4',	'Uaca',	'Tle3',	'Rplp1',	'Kif23',	'Paqr5',	'Coro2b',	'Map2k5',	'Iqch',	'Aagab',	'Snapc5',	'Tipin',	'Dis3l',	'Clpx',	'Oaz2',	'Snx1',	'Dapk2',	'Usp3',	'Rps27l',	'Anxa2',	'Myo1e',	'Aldh1a2',	'Polr2m',	'Myzap',	'Zfp280d',	'Rfx7',	'Prtg',	'Pygo1',	'Wdr72',	'Myo5c',	'Bmp5',	'Hcrtr2',	'Fam83b',	'Col12a1',	'Irak1bp1',	'Sh3bgrl2',	'Elovl4',	'Ube2cbp',	'Cep162',	'Adamts7',	'Plscr1',	'Spsb4',	'Clstn2',	'Nme9',	'Dzip1l',	'Slco2a1',	'Tmem108',	'Acpp',	'Pik3r4',	'Col6a6',	'Poc1a',	'Rpl29',	'Manf',	'Cacna2d2',	'Hyal2',	'Sema3f',	'Mst1r',	'Traip',	'Amigo3',	'Rhoa',	'Lamb2',	
                  'Qrich1',	'Dalrd3',	'Klhl18',	'Stt3b',	'Scn5a',	'Eif1b',	'Entpd3',	'Ulk4',	'Pomgnt2',	'Topaz1',	'Tcaim',	'Limd1',	'Sacm1l',	'Zmat5',	'Nf2',	'Emid1',	'Kremen1',	'Znrf3',	'Polm',	'H2afv',	'Gm11985',	'Igfbp1',	'Spata48',	'Fignl1',	'Ddc',	'Wdr92',	'Ugp2',	'Fam161a',	'4931440F15Rik',	'Sptbn1',	'Ubtd2',	'Kcnmb1',	'Thg1l',	'Sox30',	'Irgm1',	'Olfr1396',	'Cby3',	'Zfp879',	'Zfp2',	'Zfp354b',	'Zfp354a',	'Skp1a',	'Gm10447',	'Pdlim4',	'Gm2a',	'Fat2',	'Sparc',	'Atox1',	'G3bp1',	'Zfp692',	'Guk1',	'Wnt9a',	'Cops3',	'Rai1',	'Llgl1',	'Zswim7',	'Trpv2',	'Lrrc75a',	'F930015N05Rik',	'Dnah9',	'Tmem220',	'Dhrs7c',	'Ntn1',	'Pik3r6',	'Mfsd6l',	'Rpl26',	'Aurkb',	'Borcs6',	'Tmem107',	'Hes7',	'Aloxe3',	'Gucy2e',	'Chd3',	'Kdm6b',	'Dnah2',	'Wrap53',	'Eif5a',	'Cldn7',	'Dlg4',	'Pelp1',	'Tekt1',	'Ggt6',	'Spns2',	'P2rx1',	'Haspin',	'Rflnb',	'Glod4',	'Timm22',	'Ssh2',	'Coro6',	'Ankrd13b',	'Git1',	'Fam222b',	'Traf4',	'Rpl23a',	'Aldoc',	'Wsb1',	'Gm9964',	'Suz12',	'Crlf3',	'Rnf135',	'Psmd11',	'Spaca3',	'Taf15',	'Wfdc21',	'Hnf1b',	'Ddx52',	'Car4',	'Tbx2',	'Gm11444',	'Brip1',	'Med13',	'Ska2',	'Rad51c',	'Tex14',	'Rnf43',	'Supt4a',	'Dynll2',	'Cuedc1',	'Mrps23',	'Msi2',	'Akap1',	'Scpep1',	'Gm525',	'Ankfn1',	'Pctp',	'Tmem100',	'Hlf',	'Tom1l1',	'Utp18',	'Epn3',	'Samd14',	'Dlx3',	'Tac4',	'Spop',	'Nxph3',	'Phb',	'Prr15l',	'4933428G20Rik',	'Srcin1',	'Epop',	'Mllt6',	'Pcgf2',	'Lasp1',	'Fbxo47',	'Plxdc1',	'Pgap3',	'Grb7',	'Cdc6',	'Rara',	'Top2a',	'Tns4',	'Krt20',	'Krt23',	'Krtap1-5',	'Krt19',	'Hap1',	'P3h4',	'Atp6v0a1',	'Coasy',	'Plekhh3',	'Ccr10',	'Aarsd1',	'Arl4d',	'Etv4',	'Dusp3',	'Nags',	'Gm11627',	'Fzd2',	'Adam11',	'Eftud2',	'Gfap',	'Nmt1',	'Fmnl1',	'Map3k14',	'43900',	'Prr29',	'Icam2',	'1810010H24Rik',	'Pitpnc1',	'Axin2',	'Arsg',	'Wipi1',	'Fam20a',	'Abca8b',	'Map2k6',	'Slc39a11',	'Sstr2',	'Atp5h',	'Jpt1',	'Tmem94',	'Recql5',	'Itgb4',	'Wbp2',	'St6galnac2',	'Srsf2',	'Engase',	'Rbfox3',	'Cbx2',	'Nptx1',	'Rptor',	'Slc25a10',	'P4hb',	'Arhgdia',	'Aspscr1',	'Rfng',	'Tbcd',	'Calml3',	'Net1',	'Akr1e1',	'Ero1lb',	'Psma2',	'Mplkip',	'Nme8',	'Elmo1',	'Trim27',	'Zscan12',	'Zkscan4',	'Hist1h2bl',	'Hist1h4j',	'Zfp184',	'Abt1',	'Btn1a1',	'Hist1h4h',	'Hist1h2af',	'Hist1h3g',	'Hist1h2bh',	'Hist1h4f',	'Hist1h1d',	'Hist1h3e',	'Hist1h2ae',	'Hist1h4d',	'Hist1h2be',	'Hist1h2ac',	'Hist1h2bc',	'Hist1h4c',	'Hist1h3c',	'Hist1h4b',	'Hist1h4a',	'Sox4',	'Cdkal1',	'Agtr1a',	'Hus1b',	'Gmds',	'Serpinb6b',	'Serpinb1b',	'Bphl',	'Slc22a23',	'Pxdc1',	'Cdyl',	'Dsp',	'Bmp6',	'Ofcc1',	'Phactr1',	'Jarid2',	'Atxn1',	'Rnf144b',	'Id4',	'Fam120a',	'S1pr3',	'Shc3',	'Secisbp2',	'Diras2',	'Sptlc1',	'Thoc3',	'Gprin1',	'Unc5a',	'Slc34a1',	'Dbn1',	'Pdlim7',	'B4galt7',	'B230219D22Rik',	'Txndc15',	'Spock1',	'Idnk',	'Ntrk2',	'Dapk1',	'Zfp808',	'6720489N17Rik',	'Platr25',	'Gm5141',	'Ptch1',	'Hsd17b3',	'Slc35d2',	'Zfp367',	'Aaed1',	'Uqcrb',	'Zfp429',	'Zfp738',	'Zfp85',	'Mtrr',	'1700001L19Rik',	'Adamts16',	'Irx1',	'Irx2',	'Ndufs6',	'Tert',	'Pou5f2',	'Edil3',	'Hapln1',	'Fam151b',	'Serinc5',	'Cmya5',	'Bhmt',	'Wdr41',	'Aggf1',	'Poc5',	'Ankdd1b',	'Fam169a',	'Gm10260',	'Tmem171',	'Fcho2',	'Cartpt',	'Mccc2',	'Taf9',	'Mrps36',	'Cd180',	'Srek1',	'Adamts6',	'Ndufaf2',	'Ccno',	'Cdc20b',	'Gpx8',	'Snx18',	'Fst',	'Zfp131',	'Asxl2',	'Sdc1',	'Fam49a',	'Mycn',	'Nbas',	'Ntsr2',	'E2f6',	'2410004P03Rik',	'Kcnf1',	'Atp6v1c2',	'Rrm2',	'Id2',	'Cmpk2',	'Adi1',	'Tmem18',	'Slc26a3',	'Pik3cg',	'Agr3',	'Tspan13',	'Dock4',	'Prkd1',	'Coch',	'Nubpl',	'Akap6',	'Eapp',	'Brms1l',	'Nkx2-1',	'Nkx2-9',	'Foxa1',	'Trappc6b',	'Trim9',	'Psma3',	'Tomm20l',	'Dact1',	'Daam1',	'Lrrc9',	'Pcnx4',	'Trmt5',	'Hif1a',	'Wdr89',	'Esr2',	'Plekhg3',	'Rab15',	'Gm6657',	
                  'Tmem229b',	'Zfyve26',	'Dcaf5',	'Slc39a9',	'Ccdc177',	'Susd6',	'Srsf5',	'Map3k9',	'Sipa1l1',	'Rgs6',	'Dpf3',	'Dcaf4',	'Zfyve1',	'Fam161b',	'Lin52',	'Npc2',	'Isca2',	'Ttll5',	'Tgfb3',	'Irf2bpl',	'Pomt2',	'Galc',	'Ttc8',	'Foxn3',	'Efcab11',	'Psmc1',	'Calm1',	'Rps6ka5',	'Slc24a4',	'Serpina6',	'Clmn',	'Setd3',	'Mok',	'Lbhd2',	'Gm266',	'Ckb',	'Aspg',
                  'Akt1',	'Cep170b',	'Tedc1',	'Wdr60',	'Cdca7l',	'Dab2',	'Nadk2',	'Tars',	'Basp1',	'Dnah5',	'Ctnnd2',	'43896',	'Cct5',	'Sema5a',	'Matn2',	'Ankrd46',	'Grhl2',	'Ubr5',	'Nudcd1',	'Eif3h',	'Rad21',	'Med30',	'Ext1',	'Taf2',	'Fam83a',	'Atad2',	'Ndufb9',	'Mtss1',	'Nsmce2',	'Tmem71',	'St3gal1',	'Jrk',	'Slurp1',	'Mafa',	'Exosc4',	'Adck5',	'Slc39a4',	'Rbfox2',	'Ift27',	'Mpst',	'Kctd17',	'Gga1',	'Pdxp',	'Ddx17',	'Gtpbp1',	'Cbx7',	'Syngr1',	'Mkl1',	'Dnajb7',	'Zc3h7b',	'Tob2',	'Xrcc6',	'Srebf2',	'Cenpm',	'Poldip3',	'Cyb5r3',	'Arfgap3',	'Arhgap8',	'Nup50',	'Upk3a',	'Trmu',	'Celsr1',	'Rabl2',	'Pphln1',	'Arid2',	'Scaf11',	'Slc38a1',	'Adcy6',	'Tuba1b',	'Prph',	'Spats2',	'Kcnh3',	'Lima1',	'Dip2b',	'Slc11a2',	'Slc4a8',	'Atg101',	'6030408B16Rik',	'Eif4b',	'Mfsd5',	'Espl1',	'Hoxc4',	'Cbx5',	'Hnrnpa1',	'Nfe2',	'Gpr84',	'Eef2kmt',	'Abat',	'Tekt5',	'Ciita',	'Fgd4',	'Top3b',	'Ccdc116',	'Hic2',	'Lrrc74b',	'Smpd4',	'Klhl22',	'Trmt2a',	'Comt',	'Gp1bb',	'Abcc5',	'Ap2m1',	'Fam131a',	'Vps8',	'Etv5',	'Crygs',	'Adipoq',	'St6gal1',	'Bcl6',	'Lpp',	'Tprg',	'P3h2',	'Mb21d2',	'Atp13a5',	'Fam43a',	'Bdh1',	'Dlg1',	'Zdhhc19',	'Tfrc',	'Iqcg',	'Lmln',	'Umps',	'Ropn1',	'Hacd2',	'Fam162a',	'Stfa3',	'Slc15a2',	'Gpr156',	'Zdhhc23',	'Gramd1c',	'Sidt1',	'Ccdc80',	'Nectin3',	'Morc1',	'Nit2',	'Cmss1',	'Nsun3',	'Cxadr',	'N6amt1',	'Scaf4',	'Mis18a',	'4932438H23Rik',	'Rcan1',	'Runx1',	'Setd4',	'Morc3',	'Chaf1b',	'Igsf5',	'Pcp4',	'Dscam',	'Bace2',	'Mx1',	'A630089N07Rik',	'Cldn20',	'Tagap',	'Rsph3a',	'Pacrg',	'Dact2',	'Spaca6',	'Zfp53',	'Zfp13',	'Bicdl2',	'Tnfrsf12a',	'Dcpp3',	'Kctd5',	'Noxo1',	'Tpsg1',	'Sstr5',	'Mettl26',	'Mrpl28',	'Luc7l',	'Cuta',	'D17Wsu92e',	'Tead3',	'Brpf3',	'Ppil1',	'Pim1',	'Dnah8',	'Tmprss3',	'Pde9a',	'Ndufv3',	'Hsf2bp',	'Ephx3',	'Akap8l',	'Cyp4f17',	'Zbtb22',	'Rgl2',	'H2-DMb2',	'Lsm2',	'Vars',	'Abhd16a',	'Prrc2a',	'H2-Q10',	'Gtf2h4',	'Tubb5',	'H2-T23',	'Mog',	'9130008F23Rik',	'Cd2ap',	'Enpp5',	'Supt3',	'Cdc5l',	'Tmem151b',	'Slc35b2',	'Hsp90ab1',	'Mrpl14',	'Mad2l1bp',	'Ubr2',	'Mrps10',	'Guca1a',	'Taf8',	'1700122O11Rik',	'1700067P10Rik',	'Nfya',	'Lrfn2',	'Satb1',	'Mpnd',	'Fem1a',	'Ticam1',	'Gtf2f1',	'Slc25a23',	'Lama1',	'Epb41l3',	'Zbtb14',	'Tgif1',	'Yipf4',	'Fez2',	'Sult6b1',	'Ndufaf7',	'Rmdn2',	'Galm',	'Gemin6',	'Dhx57',	'Cdkl4',	'Pkdcc',	'Eml4',	'Kcng3',	'Plekhh2',	'Camkmt',	'Mcfd2',	'Epcam',	'Msh2',	'Msh6',	'Gtf2a1l',	'Gm47283',	'Bambi',	'Rbbp8',	'Ankrd29',	'Zfp521',	'Aqp4',	'Dtna',	'Zfp24',	'AW554918',	'Celf4',	'Ammecr1l',	'Bin1',	'Camk4',	'Epb41l4a',	'Apc',	'Brd8',	'Lrrtm2',	'Matr3',	'Spata24',	'Dnajc18',	'Psd2',	'Nrg2',	'Cystm1',	'Pcdhgc3',	'Pcdhgc5',	'Yipf5',	'Prelid2',	'Rbm27',	'Ccdc112',	'Sema6a',	'Dtwd2',	'Srfbp1',	'Lox',	'Ppic',	'Cep120',	'Phax',	'Megf10',	'Ctxn3',	'Fbn2',	'Chsy3',	'Slc6a7',	'Ppargc1b',	'Afap1l1',	'Sh3tc2',	'Pmaip1',	'Seh1l',	'Tcf4',	'Rab27b',	'Smad4',	'Lipg',	'Rpl17',	'Atp5a1',	'Gm6133',	'Pard6g',	'Txnl4a',	'Timm21',	'Acy3',	'Rps6kb2',	'Zdhhc24',	'Mrpl11',	'Npas4',	'Banf1',	'Tigd3',	'Capn1',	'Sac3d1',	'Macrod1',	'Otub1',	'Mark2',	'2700081O15Rik',	'Atl3',	'Pla2g16',	'Bscl2',	'Ganab',	'Eef1g',	'Fads3',	'Syt7',	'Cd6',	'Tmem132a',	'Tmem109',	'Mrpl16',	'Dtx4',	'Tle4',	'Cep78',	'Prune2',	'Gm3443',	'Trpm3',	'Tmem252',	'Kank1',	'4430402I18Rik',	'Pdcd1lg2',	'Ranbp6',	'Dkk1',	'Rnls',	'Slc16a12',	'Pank1',	'Ppp1r3c',	'Btaf1',	'Cpeb3',	'Noc3l',	'Aldh18a1',	'Opalin',	'Slit1',	'Mms19',	'Cyp2c23',	'Cwf19l1',	'Slf2',	'Sfxn3',	'Btrc',	'Poll',	'Npm3',	'Pprc1',	'Gm6970',	'Stn1',	'Cfap58',	'Xpnpep1',	'Add3',	'Mxi1',	'Dusp5',	'Smc3',	'Tectb',	'Acsl5',	'Tcf7l2',	'Habp2',	'Gfra1',	'Hspa12a',	'Eno4',	'Emx2',	'Csf2ra'))

integrate <- AddModuleScore(object = integrate, features = BRClean, ctrl = 500, name = "NKXBothRetainedCleanGenes", search = TRUE, random.seed = 1)
FeaturePlot(object = integrate, features = 'NKXBothRetainedCleanGenes1', split.by = "ID", cols = c('lightgrey','red'))
FeaturePlot(object = integrate, features = 'NKXBothRetainedCleanGenes1', cols = c('lightgrey','red'))


ProgMoreClean <- list(c('Xkr4',	'Jph1',	'Fam178b',	'Gm3646',	'Pou3f3',	'Nck2',	'Rpe',	'Erbb4',	'Igfbp5',	'Acsl3',	'Pid1',	'Glrp1',	'Cops8',	'Twist2',	'Serpinb8',	'Cxcr4',	'Lemd1',	'Ube2t',	'Lgr6',	'Cdc73',	'Hmcn1',	'Teddm2',	'Fam163a',	'Brinp2',	'Cd247',	'Rxrg',	'1700056E22Rik',	'Lyplal1',	'Tgfb2',	'Esrrg',	'Smyd2',	'Prox1',	'Plxna2',	'Camk1d',	'St8sia6',	'Hacd1',	'Nebl',	'Notch1',	'Sardh',	'Rxra',	'Ppp1r26',	'Barhl1',	'Med27',	'Zbtb43',	'Crb2',	'Nr5a1',	'Fmnl2',	'Nr4a2',	'Dpp4',	'Zfp385b',	'Prdx6b',	'Zfp804a',	'Slc43a3',	'Slc1a2',	'Wt1',	'Them7',	'Pax6',	'Arl14ep',	'Ryr3',	'Rasgrp1',	'Frmd5',	'Sord',	'Gatm',	'Sqor',	'Gm9913',	'Ncaph',	'Bcl2l11',	'Mrps26',	'Bmp2',	'Snap25',	'Ovol2',	'Syndig1',	'Gm14226',	'Src',	'Dhx35',	'Mafb',	'Tox2',	'Kcng1',	'Sall4',	'Tfap2c',	'Bmp7',	'Ankrd60',	'Stx16',	'Ccdc160',	'Asb12',	'Snx12',	'Pak3',	'Zfp704',	'Pag1',	'Dnajc5b',	'Fndc3b',	'Tmem212',	'Egfem1',	'Slc7a14',	'Sox2',	'Mab21l1',	'Gpr87',	'Plch1',	'Iqcj',	'Rxfp1',	'Pdgfc',	'Sfrp2',	'Tmem131l',	'Kirrel',	'Fcgr1',	'Vangl1',	'Cym',	'A630076J17Rik',	'Col11a1',	'Cdc14a',	'Plppr5',	'Dpyd',	'Ptbp2',	'Ugt8a',	'Elovl6',	'Lef1',	'Adh6a',	'Unc5c',	'Pkn2',	'Col24a1',	'Cyr61',	'St6galnac5',	'Triqk',	'Fut9',	'Lingo2',	'Pax5',	'Igfbpl1',	'Xpa',	'Rad23b',	'Ambp',	'Astn2',	'Trim32',	'Brinp1',	'Gm11232',	'Slc24a2',	'Mtap',	'Jun',	'Hook1',	'E130114P18Rik',	
                        'Lepr',	'Pde4b',	'Dmrta2',	'3100002H09Rik',	'Nkain1',	'Sdc3',	'Matn1',	'Ptpru',	'Ephb2',	'Wnt4',	'Igsf21',	'Camta1',	'A430005L14Rik',	'Fam133b',	'Steap4',	'Speer4c',	'Mnx1',	'Cad',	'Babam2',	'Cpz',	'Sorcs2',	'Zfp518b',	'Gm16223',	'Fgfbp1',	'Prom1',	'Smim20',	'9130230L23Rik',	'Lnx1',	'Adgrl3',	'Npffr2',	'Fras1',	'Prkg2',	'Mapk10',	'Asphd2',	'Sez6l',	'Myo18b',	'Aym1',	'Gcn1l1',	'Hspb8',	'Ksr2',	'2410131K14Rik',	'Rbm19',	'Rflna',	'Tmem132c',	'Fzd10',	'Galnt17',	'Gtf2ird1',	'Cldn4',	'Ephb4',
                        'Slc25a13',	'Glcci1',	'Aass',	'Mest',	'Plxna4',	'Chchd3',	'Hipk2',	'Tmem178b',	'Tpk1',	'Cntnap2',	'Gimap7',	'Igf2bp3',	'Snx10',	'Creb5',	'Herc6',	'Atoh1',	'Mad2l1',	'Hk2',	'Gfpt1',	'Iqsec1',	'1700123L14Rik',	'Frmd4b',	'Rybp',	'Lrrn1',	'Lmcd1',	'Hrh1',	'Zfp637',	'Ntf3',	'Etv6',	'Grin2b',	'Gucy2c',	'Ptpro',	'Lmo3',	'Pde3a',	'Etnk1',	'Usp29',	'Ccdc8',	'Tshz3',	'Kcnj11',	'Htatip2',	'Chrna7',	'St8sia2',	'Tmem135',	'Rab30',	'Emsy',	'Wnt11',	'Rassf10',	'Spon1',	'Insc',	'Acsm1',	'Acsm4',	'Eef2k',	'Zkscan2',	'Wdr11',	'Nps',	'Mgmt',	'Kndc1',	'Ifitm10',	'Mrpl23',	'Cttn',	'Akap12',	'Myb',	'Echdc1',	'Amd2',	'Crybg1',	'Chst3',	'Ctnna3',	'Lrrtm3',	'Phyhipl',	'Pofut2',	'Chst11',	'Appl2',	'1700113H08Rik',	'Cep83',	'Btg1',	'Lum',	'Kitl',	'Otogl',	'Bbs10',	'Ptprr',	'Hmga2',	'Efnb2',	'Mcf2l',	'Csmd1',	'Unc5d',	'Cldn23',	'Cenpu',	'Enpp6',	'Dctd',	'Tenm3',	'Galnt7',	'Cbr4',	'Cers1',	'Comp',	'Plvap',	'Klf2',	'Hhip',	'Il15',	'Ndufb7',	'Sall1',	'Tox3',	'Rbl2',	'Irx5',	'Drc7',	'Setd6',	'Cdh8',	'Cdh11',	'Cdh3',	'Psmd7',	'Zfhx3',	'Fa2h',	'Adamts18',	'Maf',	'Tldc1',	'Foxl1',	'Map3k21',	'Tomm20',	'Nrp1',	'Top2b',	
                        'Gng2',	'Slc35f4',	'Tnfrsf19',	'Gata4',	'Gfra2',	'Htr2a',	'Olfm4',	'Pcdh20',	'Pcdh9',	'Klf12',	'Ipo5',	'Stk24',	'Nalcn',	'Fgf14',	'Sesn3',	'Jam3',	'Igsf9b',	'Opcml',	'Barx2',	'St3gal4',	'Pknox2',	'Tbcel',	'Grik4',	'Pou2f3',	'Nxpe4',	'Elmod1',	'Cspg4',	'Loxl1',	'Uaca',	'Igdcc3',	'Rora',	'Impg1',	'Sh3bgrl2',	'Slc9a9',	'Rasa2',	'Mrps22',	'Mrpl3',	'Arpp21',	'Trim71',	'Scn5a',	'Mobp',	'Pkd1l1',	'Hus1',	'Vwc2',	'Meis1',	'Bcl11a',	'Ccdc85a',	'Sptbn1',	'Srebf1',	'Dnah9',	'Glp2r',	'Ntn1',	'Smtnl2',	'1700016K19Rik',	'Wsb1',	'Mrm1',	'Bcas3',	'Nog',	'Ankfn1',	'Pctp',	'Hils1',	'Tac4',	'Phb',	'Igf2bp1',	'Ikzf3',	'Krt20',	'Krt13',	'G6pc3',	'Wnt9b',	'1700012B07Rik',	'Map2k6',	'Kcnj16',	'Sox9',	'Sdk2',	'Tnrc6c',	'Cyth1',	'Cbx4',	'Klf6',	'Chrm3',	'Elmo1',	'Nrsn1',	'Cdkal1',	'Mylk4',
                        'Hivep1',	'Phactr1',	'Rnf144b',	'Id4',	'Sema4d',	'Ror2',	'Msx2',	'Pitx1',	'Spock1',	'Agtpbp1',	'Gas1',	'Dapk1',	'Fbp2',	'Ercc6l2',	'Hsd17b3',	'Irx2',	'Poc5',	'Cartpt',	'Naip6',	'Isl1',	'B430218F22Rik',	'Dnmt3a',	'Pomc',	'Dnajc27',	'Fam49a',	'Sox11',	'Sntg2',	'Twist1',	'Sostdc1',	'Dgkb',	'Npas3',	'Egln3',	'Dact1',	'Six1',	'Slc38a6',	'Actn1',	'Dpf3',	'Jdp2',	'Tgfb3',	'Esrrb',	'Sel1l',	'Dlk1',	'Akt1',	'Ghr',	'Slc1a3',	'Adamts12',	'Npr3',	'Basp1',	'Otulin',	'Ankrd33b',	'Trps1',	'Sntb1',	'Tmem65',	'Myc',	'Fam49b',	'St3gal1',	'Slc45a4',	'Gpr20',	'Ly6h',	'Mb',	'Tcf20',	'Rtl6',	'Fam118a',	'Fam19a5',	'Syt10',	'Tmem117',	'Slc38a4',	'Col2a1',	'Smug1',	'Sec14l5',	'Eef2kmt',	'Rtl10',	'Klhl6',	'Il1rap',	'Gmnc',	'Fgf12',	'Opa1',	'Fam43a',	'Tfrc',	'Hacd2',	'Csta1',	'Igsf11',	'Lsamp',	'Bach1',	'Setd4',	'Ets2',	'B3galt5',	'Pcp4',	'Dact2',	'Cldn9',	'1520401A03Rik',	'Lemd2',	'Glp1r',	'Pde9a',	'H2-Q2',	'Spats1',	'1600014C23Rik',	'Ptprm',	'Arhgap28',	'Dlgap1',	'Tgif1',	'Crim1',	'Thada',	'Slc3a1',	'Six2',	'Srbd1',	'Zfp521',	'Cdh2',	'B4galt6',	'Arap3',	'Pabpc2',	'Ppp2r2b',	'Spink1',	'Sema6a',	'Eno1b',	'Cep120',	'Prrc1',	'St8sia3',	'Pmaip1',	'Tcf4',	'Ccdc68',	'Dcc',	'Timm21',	'Nrxn2',	'Gnaq',	'Rfk',	'Trpm3',	'Mamdc2',	'Dmrt3',	'Gm815',	'Dkk1',	'Pten',	'Kif20b',	'Gm340',	'Sorcs3',	'Sorcs1',	'Add3',	'Tectb',	'Tcf7l2',	'Habp2',	'Emx2'))

integrate <- AddModuleScore(object = integrate, features = ProgMoreClean, ctrl = 500, name = "NKXProgMoreCleanGenes", search = TRUE, random.seed = 1)
FeaturePlot(object = integrate, features = 'NKXProgMoreCleanGenes1', split.by = "ID", cols = c('lightgrey','red'))
FeaturePlot(object = integrate, features = 'NKXProgMoreCleanGenes1', cols = c('lightgrey','red'))


ProgLessClean <- list(c('Arfgef1',	'Slco5a1',	'Stau2',	'Pkhd1',	'Col9a1',	'Tmem131',	'Inpp4a',	'2010300C02Rik',	'Rpl31',	'Il1rl2',	'1500015O10Rik',	'Slc39a10',	'1700019D03Rik',	'Stk17b',	'Spats2l',	'Acadl',	'Ikzf2',	'Rpl37a',	'Tmbim1',	'Speg',	'Fam124b',	'Dner',	'Cab39',	'Kcnj13',	'Gpc1',	'Slc35f5',	'Gpr39',	'Nckap5',	'Tmem163',	'Pigr',	'Fam72a',	'Slc26a9',	'Cntn2',	'Nfasc',	'Mdm4',	'Pik3c2b',	'Prelp',	'Arl8a',	'Fam129a',	'Edem3',	'Lamc1',	'Gm10031',	'Fam20b',	'Angptl1',	'4930523C07Rik',	'Kifap3',	'Dpt',	'Mpzl1',	'Gpa33',	'Rgs5',	'B4galt3',	'Fmn2',	'Grem2',	'1700016C15Rik',	'Rrp15',	'Spata17',	'Dtl',	'Hhat',	'A130010J15Rik',	'Cd34',	'Phyh',	'Optn',	'Usp6nl',	'Dnajc1',	'Msrb2',	'Arhgap21',	'Zdhhc12',	'Hmcn2',	'Qrfp',	'Lmx1b',	'Mvb12b',	'Pbx3',	'Dab2ip',	'Strbp',	'Nek6',	'Lypd6b',	'Rbm43',	'Fign',	'Scn7a',	'Stk39',	'Tlk1',	'Cdca7',	'Itgav',	'Tfpi',	'Ctnnd1',	'Ypel4',	'Ube2l6',	'Gm9821',	'Chst1',	'Prdm11',	'Ehf',	'Depdc7',	'Prrg4',	'Aven',	'Meis2',	'Fsip1',	'Bmf',	'Bub1b',	'Ccdc32',	'Ino80',	'Ehd4',	'Eid1',	'Ap4e1',	'Mrps5',	'Gpcpd1',	'Foxa2',	'Pygb',	'Id1',	'Bcl2l1',	'Efcab8',	'Bpifb5',	'Chmp4b',	'Nnat',	'Adig',	'Chd6',	'Ncoa3',	'Arfgef2',	'Tshz2',	'Pck1',	'Rab22a',	'Zfp831',	'Cables2',	'Clcn5',	'Gripap1',	'Maob',	'Slc9a7',	'44080',	'Xpnpep2',	'Gpc4',	'Hprt',	'Abcd1',	'Prkx',	'Arhgef9',	'Amer1',	'Pja1',	'Ogt',	'Nexmif',	'Gm379',	'Gm6377',	'Pof1b',	'Chm',	'Hnrnph2',	'Gprasp1',	'Bex4',	'Tceal8',	'Rnf128',	'Huwe1',	'Spin2c',	'Eif1ax',	'Gemin8',	'Msl3',	'Mrps28',	'Tpd52',	'Zbtb10',	'Car13',	'Tnik',	'Mecom',	'Actl6a',	'Gm11549',	'Bbs12',	'Spry1',	'Pgrmc2',	'P2ry13',	'Mbnl1',	'Lekr1',	'Rapgef2',	'Gucy1b1',	'Rbm46',	'Dchs2',	'Fhdc1',	'Arfip1',	'Tigd4',	'Fam160a1',	'Etv3',	'Mrpl24',	'Rhbg',	'Atp8b2',	'Vps45',	'Mtmr11',	'Acp6',	'Reg4',	'Hmgcs2',	'Ptgfrn',	'Igsf3',	'Casq2',	'Syt6',	'Olfml3',	'Magi3',	'Wnt2b',	'S1pr1',	'Abca4',	'Gclm',	'Bcar3',	'1810037I17Rik',	'Camk2d',	'Rrh',	'Bdh2',	'Slc39a8',	'Bank1',	'Mttp',	'4930579F01Rik',	'Pdlim5',	'Ssx2ip',	'Miga1',	'Zzz3',	'Wls',	'Fam110b',	'Asph',	'Ccne2',	'Gpr63',	'Unc13b',	'Atp8b5',	'Glipr2',	'Gne',	'Fbxo10',	'Tmod1',	'Trmo',	'Nans',	'Tal2',	'Klf4',	'D630039A03Rik',	'Susd1',	'Slc46a2',	'Slc31a2',	'Rgs3',	'Col27a1',	'Pappa',	'Tlr4',	'Lurap1l',	'Mpdz',	'Nfib',	'Plin2',	'Dennd4c',	'Ttc22',	'Lrrc42',	'Foxd2',	'Ptprf',	'Foxo6',	'Kcnq4',	'Mfsd2a',	'Ak2',	'C77080',	'Bsdc1',	'Epb41',	'Trim63',	'Srrm1',	'Rcan3',	'Ifnlr1',	'Rap1gap',	'Szrd1',	'Dnajc16',	'Pdpn',	'Dhrs3',	'Lzic',	'Car6',	'Acot7',	'B230104I21Rik',	'Mmel1',	'Fam213b',	'Morn1',	'Cdk14',	'Dmtf1',	'Sema3e',	'Srpk2',	'Shh',	'Tbc1d14',	'C1qtnf7',	'Cc2d2a',	'Qdpr',	'Stim2',	'N4bp2',	'Rbm47',	'Limch1',	'Slain2',	'Cwh43',	'Rasl11b',	'Thegl',	'Cxcl15',	'Parm1',	'Scarb2',	'Shroom3',	'Ccni',	'Gpat3',	'Ptpn13',	'Cdc7',	'Tgfbr3',	'Fgfrl1',	'Crybb3',	'Ssh1',	'Med13l',	'Atp2a2',	'Tmem120b',	'Tmem132d',	'Gtf2i',	'Sh2b2',	'Cux1',	'Myl10',	'Prkar1b',	'Slc29a4',	'Tnrc18',	'Rbak',	'Lmtk2',	'Bhlha15',	'Alox5ap',	'Fry',	'Pds5b',	'Stard13',	'Ppp1r9a',	'Asb4',	'Cftr',	'Lmod2',	'Tspan33',	'Cep41',	'Mkln1',
                        'Lrguk',	'Bpgm',	'Agbl3',	'Dgki',	'Ephb6',	'Gsdme',	'Nfe2l3',	'Adcyap1r1',	'Thnsl2',	'Sema4f',	'Rtkn',	'Tet3',	'Ankrd53',	'Tgfa',	'Antxr1',	'Plxna1',	'Slc6a6',	'Prickle2',	'Kbtbd8',	'Mitf',	'Foxp1',	'Eif4e3',	'Shq1',	'Rad18',	'Setd5',	'Atp2b2',	'Vgll4',	'Cand2',	'Wnt5b',	'C1s2',	'Acrbp',	'Iffo1',	'A2ml1',	'Styk1',	'E330021D16Rik',	'Atf7ip',	'Pik3c2g',	'Plekha5',	'St8sia1',	'Smco2',	'Klhl42',	'Bicd1',	'Zfp773',	'Dact3',	'Zfp60',	'Zfp568',	'Plekhf1',	'Med25',	'Irf3',	'Emp3',	'Spty2d1',	'Dbx1',	'Mkrn3',	'Igf1r',	'Nr2f2',	'Fam174b',	'Slco3a1',	'Akap13',	'Isg20',	'Wdr73',	'Mex3b',	'Ctsc',	'Fzd4',	'Prss23',	'Nars2',	'Uvrag',	'Serpinh1',	'P4ha3',	'Arhgef17',	'Pde2a',	'Olfr571',	'Scube2',	'Galnt18',	'Rps15a',	'Ndufab1',	'Plk1',	'Tbx6',	'Zfp629',	'Uros',	'Tcerg1l',	'Pwwp2b',	'Tspan32',	'Cdkn1c',	'Shank2',	'Ano1',	'Ccnd1',	'Iyd',	'Vnn3',	'Ctgf',	'Ptprk',	'Marcks',	'Wisp3',	'Traf3ip2',	'Foxo3',	'Afg1l',	'Cd24a',	'Rtn4ip1',	'Prdm1',	'Dcbld1',	'Pln',	'Asf1a',	'Psap',	'Unc5b',	'Pcbd1',	'Tacr2',	'Srgn',	'Nrbf2',	'Ccdc6',	'Specc1l',	'Mmp11',	'Zfp280b',	'Dnmt3l',	'AU041133',	'Nfyb',	'Nuak1',	'Chpt1',	'Nr1h4',	'Tmpo',	'Nudt4',	'Atp2b1',	'Poc1b',	'Lin7a',	'Tph2',	'Zfc3h1',	'Frs2',	'Cpsf6',	'Rap1b',	'Grip1',	'Tbc1d30',	'Slc16a7',	'Shmt2',	'Baz2a',	'Rab20',	'Tfdp1',	'Cdc16',	'Kbtbd11',	'Xkr5',	'Smim19',	'Eri1',	'Mtnr1a',	'Tlr3',	'Ccdc110',	'Irf2',	'Ing2',	'Cldn22',	'Sap30',	'Cpe',	'Pbx4',	'Mmaa',	'Adgrl1',	'Ier2',	'G430095P16Rik',	'Dand5',	'Dnaja2',	'Ces1g',	'Ogfod1',	'Pllp',	'Ccdc102a',	'Cfap20',	'Cmtm4',	'Terf2',	'Tmem170',	'Sdr42e1',	'Gins2',	'Rhou',	'Rab4a',	'Sipa1l2',	'Pcnx2',	'Pard3',	'Flnb',	'Kctd6',	'Dlg5',	'Il17rd',	'Hacl1',	'Gm8113',	'Bmp4',	'Cdkn3',	'Prmt5',	'Rnaseh2b',	'Zfp395',	'Trim35',	'Bnip3l',	'Dock5',	'Chmp7',	'Sorbs3',	'Lpar6',	'Siah3',	'Cog3',	'Serp2',	'Dnajc15',	'Kctd12',	'Pou4f1',	'Spry2',	'Dct',	'Mbnl2',	'Farp1',	'Gm11168',	'Gm10718',	'Ets1',	'Fez1',	'Scn3b',	'Mpzl3',	'Scn2b',	'Sik3',	'Cadm1',	'Zbtb16',	'Ppp2r1b',	'Colca2',	'Tmem266',	'Isl2',	'Peak1',	'Stra6',	'Adpgk',	'Lrrc49',	'Anp32a',	'Coro2b',	'Calml4',	'Pias1',	
                        'Smad6',	'Plekho2',	'Pclaf',	'Gtf2a2',	'Adam10',	'Hmgcll1',	'Klhl31',	'Gcm1',	'Gsta2',	'Eef1a1',	'Irak1bp1',	'Spsb4',	'Dbr1',	'Slc35g2',	'Pccb',	'Cep63',	'Amotl2',	'Srprb',	'Atp2c1',	'Wdr82',	'Cmtm8',	'Stt3b',	'Tgfbr2',	'Ctdspl',	'Scn10a',	'Lyzl4',	'Snrk',	'Zfp105',	'Limd1',	'Ccr5',	'Kremen1',	'Rtn4',	'Bod1',	'Sh3pxd2b',	'Ubtd2',	'Fbxw11',	'Ranbp17',	'Slit3',	'Rars',	'Wwc1',	'Mat2b',	'Trim7',	'Csf2',	'Ccdc69',	'Galnt10',	'Larp1',	'Btnl10',	'Rai1',	'Zfp286',	'Hs3st3b1',	'Cox10',	'Shisa6',	'Dhrs7c',	'Myh10',	'Pfas',	'Scimp',	'Med31',	'Rap1gap2',	'Ccdc92b',	'Srr',	'Timm22',	'Rab34',	'Lyrm9',	'Rasl10b',	'Car4',	'Hsf5',	'Rnf43',	'Kat7',	'B4galnt2',	'Lasp1',	'Csf3',	'Krt25',	'Krt36',	'Plekhh3',	'Ccr10',	'Meox1',	'Cd300lg',	'Mpp2',	'BC030867',	'Ubtf',	'Arhgap27',	'Mettl2',	'Tanc2',	'Cyb561',	'Ern1',	'Kpna2',	'Bptf',	'Pitpnc1',	'Apoh',	'Wipi1',	'Nat9',	'Mgat5b',	'Slc16a3',	'Zmynd11',	'Arid4b',	'Inhba',	'Trim27',	'Gpx6',	'Hist1h2bl',	'Hist1h2an',	'Btn1a1',	'Hist1h2bh',	'Hist1h3e',	'Hist1h2bf',	'Hist1h4d',	'Hist1h3a',	'Sox4',	'Agtr1a',	'Uqcrfs1',	'Foxq1',	'Bphl',	'Rreb1',	'Sycp2l',	'Nedd9',	'Edn1',	'Tbc1d7',	'Rnf182',	'Jarid2',	'Mylip',	'Atxn1',	'Nup153',	'Zfp169',	'Gadd45g',	'Syk',	'Cplx2',	'Ddx46',	'Aaed1',	'Gm28041',	'Irx1',	'Glrx',	'Lhfpl2',	'Nsa2',	'Arhgef28',	'Utp15',	'Fcho2',	'Plk2',	'Snx18',	'Hspb3',	'Arl15',	'Parp8',	'Gm17509',	'Apob',	'Osr1',	'Rock2',	'Cys1',	'Rsad2',	'Tmem18',	'Alkal2',	'Immp2l',	'Lrrn3',	'Stxbp6',	'Mbip',	'Pole2',	'Klhdc1',	'Atp5s',	'Tmx1',	'Frmd6',	'Six4',	'Tmem30b',	'Prkch',	'Syne2',	'Max',	'Dcaf5',	'Map3k9',	'Rgs6',	'Fcf1',	'Ttll5',	'Ston2',	'Gpr68',	'Ccdc88c',	'Golga5',	'Ppp4r4',	'Dicer1',	'Bdkrb1',	'Hsp90aa1',	'Ccdc152',	'Osmr',	'Nadk2',	'Rai14',	'Myo10',	'Nipal2',	'Zfp706',	'Azin1',	'Abra',	'Eif3h',	'Deptor',	'Mrpl13',	'Zhx2',	'Fer1l6',	'Nsmce2',	'Fam84b',	'Ndrg1',	'Adgrb1',	'Eppk1',	'Hsf1',	'Rbfox2',	'Myh9',	'Sstr3',	'Cdc42ep1',	'Syngr1',	'Chadl',	'Atxn10',	'Gramd4',	'Ano6',	'Arid2',	'Slc38a1',	'Nckap5l',	'Lima1',	'Dip2b',	'Sp1',	'Gpr84',	'Zfp385a',	'Crebbp',	'Ercc4',	'Fopnl',	'Map3k13',	'Igf2bp2',	'P3h2',	'Mb21d2',	'Dlg1',	'Mylk',	'Stxbp5l',	'Gsk3b',	'Upk1b',	'Abhd10',	'Phldb2',	'Morc1',	'Nsun3',	'4930453N24Rik',	'Robo1',	'Robo2',	'Btg3',	'Adamts1',	'Grik1',	'Mis18a',	'Mx1',	'Tmem181a',	'Dynlt1a',	'Rsph3b',	'Pacrg',	'Agpat4',	'Afdn',	'Zfp820',	'Zfp947',	'Zfp945',	'Traf7',	'Neurl1b',	'Tulp1',	'Rsph1',	'Cryaa',	'Brd2',	'C2',	'Ddr1',	'Ier3',	'Ppp1r10',	'Adgrf5',	'Dnph1',	'Unc5cl',	'Pp2d1',	'Ptprs',	'Man2a1',	'Themis3',	'Vit',	'Qpct',	'Cdc42ep3',	'Eml4',	'Lrpprc',	'Svil',	'Ccny',	'Dtna',	'Nrep',	'Reep2',	'Sil1',	'Psd2',	'Hbegf',	'Pcdh1',	'Nr3c1',	'Gm5689',	'Stk32a',	'Snx24',	'Tex43',	'Sh3tc2',	'Wdr7',	'Nedd4l',	'Oacyl',	'Grp',	'Lman1',	'Rab27b',	'Smad4',	'Mro',	'Gm9925',	'Myo5b',	'Acaa2',	'Zbtb7c',	'Nfatc1',	'Zfp516',	'Cndp1',	'Ppp6r3',	'Kdm2a',	'Mus81',	'Pcnx3',	'Pla2g16',	'Chrm1',	'Fads2',	'Anxa1',	'1110059E24Rik',	'Fam189a2',	'Pum3',	'Slc16a12',	'Hells',	'Ccnj',	'Tm9sf3',	'Frat1',	'Ina',	'Sfr1',	'Gsto1',	'Afap1l2',	'Pnliprp1',	'Prdx3'))

integrate <- AddModuleScore(object = integrate, features = ProgLessClean, ctrl = 500, name = "NKXProgLessCleanGenes", search = TRUE, random.seed = 1)
FeaturePlot(object = integrate, features = 'NKXProgLessCleanGenes1', split.by = "ID", cols = c('lightgrey','red'))
FeaturePlot(object = integrate, features = 'NKXProgLessCleanGenes1', cols = c('lightgrey','red'))

#save file
saveRDS(integrate, file = "/Users/drlittle/Desktop/Agg12controls_8-6-20/Agg12_Sox9AT1AT2noprolif_mod_score.rds")


####### AT1 to Prog monocle without prolif genes #######
integrate1 <- readRDS( file = "/Users/drlittle/Desktop/Agg12_Epi_4woprolif.rds")
integrate <-readRDS( file = "/Path/to/file/Agg12_Sox9AT1AT2noprolif_mod_score.rds")

data <- as(as.matrix(integrate@assays$RNA@data), 'sparseMatrix')
pd <- new('AnnotatedDataFrame', data=integrate@meta.data)
fData <- data.frame(gene_short_name=row.names(data),row.names = row.names(data))
fd <- new('AnnotatedDataFrame', data=fData)
HSMM <- newCellDataSet(data,
                       phenoData=pd,
                       featureData=fd,
                       expressionFamily = uninormal())

ordering_genes <- integrate1@assays$RNA@var.features[1:1000]

s.genes <- c("Mcm5",     "Pcna" ,    "Tyms",     "Fen1",     "Mcm2",     "Mcm4",     "Rrm1"   ,  "Ung" ,     "Gins2",    "Mcm6" ,    "Cdca7" ,   "Dtl",      "Prim1", "Uhrf1",    "Mlf1ip",   "Hells",  "Rfc2", "Rpa2", "Nasp",     "Rad51ap1", "Gmnn",     "Wdr76" ,   "Slbp"    , "Ccne2",    "Ubr7"    , "Pold3", "Msh2",     "Atad2",    "Rad51",    "Rrm2" ,    "Cdc45" ,   "Cdc6" ,    "Exo1" ,    "Tipin"  ,  "Dscc1"  ,  "Blm"    ,  "Casp8ap2", "Usp1"  ,   "Clspn", "Pola1"   , "Chaf1b" ,  "Brip1" ,   "E2f8")

g2m.genes <- c("Hmgb2",   "Cdk1" ,   "Nusap1" , "Ube2c",   "Birc5"  , "Tpx2" ,   "Top2a",   "Ndc80"  , "Cks2"   , "Nuf2"   , "Cks1b"  , "Mki67",   "Tmpo" ,   "Cenpf",   "Tacc3",   "Fam64a" , "Smc4" ,   "Ccnb2",   "Ckap2l" , "Ckap2",   "Aurkb",   "Bub1" ,   "Kif11"  , "Anp32e" , "Tubb4b" , "Gtse1"  , "Kif20b" , "Hjurp",   "Cdca3",   "Hn1", "Cdc20",   "Ttk"  ,   "Cdc25c" , "Kif2c"  , "Rangap1", "Ncapd2" , "Dlgap5" , "Cdca2"  , "Cdca8"  , "Ect2" ,   "Kif23",   "Hmmr" ,   "Aurka"  , "Psrc1"  , "Anln", "Lbr"  ,   "Ckap5"  , "Cenpe"  , "Ctcf" ,   "Nek2" ,   "G2e3" ,   "Gas2l3" , "Cbx5",    "Cenpa" )
####this makes ordering genes actually 1918 not 2000
###this makes ordering 939 not 1000
integrate1 <- 0
integrate <- 0
ordering_genes <- ordering_genes[!ordering_genes %in% c(g2m.genes, s.genes)]
HSMM <- setOrderingFilter(HSMM, ordering_genes)
HSMM <- reduceDimension(HSMM,norm_method="none", 
                        reduction_method="DDRTree",
                        max_components=2,
                        scaling=TRUE,
                        pseudo_expr=0)
HSMM <- orderCells(HSMM) 

#Figure 4C
plot_cell_trajectory(HSMM, color_by = "ID", show_branch_points = F) + facet_wrap(~ID, nrow = 1)

#Figure 4D left side
plot_cell_trajectory(HSMM, color_by = "State", show_branch_points = F) + facet_wrap(~ID, nrow = 1)

#Figure 4D right side
plot_cell_trajectory(HSMM, markers="Spock2", use_color_gradient = T, show_branch_points = F, show_backbone = F) + scale_color_gradient(low="lightgrey", high="red")
plot_cell_trajectory(HSMM, markers="Lamp3", use_color_gradient = T, show_branch_points = F, show_backbone = F) + scale_color_gradient(low="lightgrey", high="red")
plot_cell_trajectory(HSMM, markers="Sox9", use_color_gradient = T, show_branch_points = F, show_backbone = F)  + scale_color_gradient(low="lightgrey", high="red")

#we cannot infer order from pseudotime so we can switch this to the biological root state of sox9 progenitors
HSMM <- orderCells(HSMM, root_state = 2)

#verify change
plot_cell_trajectory(HSMM, color_by = "Pseudotime")

HSMMDF <- as.data.frame(HSMM$Pseudotime)
HSMMDF$Pseudotime <- HSMM$Pseudotime
HSMMDF$NKXAT2AcquiredGenes1 <- HSMM$NKXAT2AcquiredGenes1
HSMMDF$NKXAT2RetainedGenes1<- HSMM$NKXAT2RetainedGenes1
HSMMDF$NKXAT1AcquiredGenes1 <- HSMM$NKXAT1AcquiredGenes1
HSMMDF$NKXAT1RetainedGenes1<- HSMM$NKXAT1RetainedGenes1
HSMMDF$NKXBAcquiredGenes1 <- HSMM$NKXBAcquiredGenes1
HSMMDF$NKXBothRetainedGenes1<- HSMM$NKXBothRetainedGenes1
HSMMDF$NKXProgMoreGenes1 <- HSMM$NKXProgMoreGenes1
HSMMDF$NKXProgLessGenes1<- HSMM$NKXProgLessGenes1
HSMMDF$ID<- HSMM$ID
HSMMDF$State<- HSMM$State
HSMMDF$cell_type <- HSMM$cell_type
HSMMDF$NFeatures <- HSMM$nFeature_RNA
HSMMDF$seurat_clusters <- HSMM$seurat_clusters
write.csv(HSMMDF, file = "/Path/to/file/Agg12_Mod_score_vs_pseudotime.csv")
HSMMDF <- read.csv(file = "/Users/drlittle/Desktop/Agg12_Mod_score_vs_pseudotime_corrrected.csv")
###need to split to two plots then pseudo color? 

plot_1 <- ggplot(HSMMDF, aes(HSMMDF$Pseudotime, HSMMDF$NKXAT2AcquiredGenes1)) + geom_point(aes(colour=HSMMDF$cell_type))+labs(y="NKXAT2AcquiredGenes1", x="Pseudotime") # geom_text(aes(DBG_logFC, DEG_logFC, label= GNAME,fontface=2, size=6), data= LabelName)
plot_1

plot_2 <- ggplot(HSMMDF, aes(HSMMDF$Pseudotime, HSMMDF$NKXAT2RetainedGenes1)) + geom_point(aes(colour=HSMMDF$cell_type))+labs(y="NKXAT2RetainedGenes1", x="Pseudotime") # geom_text(aes(DBG_logFC, DEG_logFC, label= GNAME,fontface=2, size=6), data= LabelName)
plot_2

plot_3 <- ggplot(HSMMDF, aes(HSMMDF$Pseudotime, HSMMDF$NKXAT1AcquiredGenes1)) + geom_point(aes(colour=HSMMDF$State))+labs(y="NKXAT1AcquiredGenes1", x="Pseudotime") # geom_text(aes(DBG_logFC, DEG_logFC, label= GNAME,fontface=2, size=6), data= LabelName)
plot_3

plot_4 <- ggplot(HSMMDF, aes(HSMMDF$Pseudotime, HSMMDF$NKXAT1RetainedGenes1)) + geom_point(aes(colour=HSMMDF$State))+labs(y="NKXAT1RetainedGenes1", x="Pseudotime") # geom_text(aes(DBG_logFC, DEG_logFC, label= GNAME,fontface=2, size=6), data= LabelName)
plot_4

plot_5 <- ggplot(HSMMDF, aes(HSMMDF$Pseudotime, HSMMDF$NKXBAcquiredGenes1)) + geom_point(aes(colour=HSMMDF$State))+labs(y="NKXBAcquiredGenes1", x="Pseudotime") # geom_text(aes(DBG_logFC, DEG_logFC, label= GNAME,fontface=2, size=6), data= LabelName)
plot_5

plot_6 <- ggplot(HSMMDF, aes(HSMMDF$Pseudotime, HSMMDF$NKXBothRetainedGenes1)) + geom_point(aes(colour=HSMMDF$State))+labs(y="NKXBothRetainedGenes1", x="Pseudotime") # geom_text(aes(DBG_logFC, DEG_logFC, label= GNAME,fontface=2, size=6), data= LabelName)
plot_6

plot_6 <- ggplot(HSMMDF, aes(HSMMDF$NFeatures, HSMMDF$ID)) + geom_point(aes(colour=HSMMDF$ID))+labs(y="NKXBothRetainedGenes1", x="Pseudotime") # geom_text(aes(DBG_logFC, DEG_logFC, label= GNAME,fontface=2, size=6), data= LabelName)
plot_6

plot_7 <- ggplot(HSMMDF, aes(HSMMDF$Pseudotime, HSMMDF$NKXProgMoreGenes1)) + geom_point(aes(colour=HSMMDF$State))+labs(y="NKXProgMoreGenes1", x="Pseudotime") # geom_text(aes(DBG_logFC, DEG_logFC, label= GNAME,fontface=2, size=6), data= LabelName)
plot_7

plot_8 <- ggplot(HSMMDF, aes(HSMMDF$Pseudotime, HSMMDF$NKXProgLessGenes1)) + geom_point(aes(colour=HSMMDF$State))+labs(y="NKXProgLessGenes1", x="Pseudotime") # geom_text(aes(DBG_logFC, DEG_logFC, label= GNAME,fontface=2, size=6), data= LabelName)
plot_8


###### This section is for Figure 5 and Supplement Figure 5 #####
#E18.5 control and Yap/Taz CKO progenitor-specific mutant samples.

library(Seurat)
library(cowplot)
library(dplyr)
library(ggplot2)
library(sctransform)
library(monocle)
library(VGAM)
library(grid)
library(EnhancedVolcano)
library(stringr)
library(MAST)

data <- Read10X(data.dir = "/Path/to/file/feature_bc_matrix/")

integrate <- CreateSeuratObject(data, min.cells = 3, min.features = 200) %>% PercentageFeatureSet(pattern = "^mt-", col.name = "percent.mt") %>% NormalizeData() %>% FindVariableFeatures() %>% ScaleData()
VlnPlot(integrate, features = c("nFeature_RNA","nCount_RNA","percent.mt"))
integrate <- subset(integrate, subset = nFeature_RNA > 200 & nFeature_RNA < 5000 & percent.mt < 15)

integrate <- ScaleData(integrate) %>% RunPCA(npcs = 30) %>% RunUMAP(dims=1:30) %>% FindNeighbors() %>% FindClusters()

#label control versus mutant in metadata
integrate$geno <- substr(colnames(integrate@assays$RNA@counts), 18, 18)
integrate$geno <- plyr::mapvalues(x = integrate$geno,
                                  from = 1:2,
                                  to = c("ctrl","SYT"))
rm(data)
#confirm new labels
DimPlot(integrate, group.by = "seurat_clusters", split.by = "geno", label = T)

#identify where epithelial cell types are
FeaturePlot(integrate, features = c("Spock2", "Lamp3", "Foxj1", "Scgb1a1"))
#identify lineage clusters and proliferating clusters
FeaturePlot(integrate, features = c("Col3a1", "Cdh5", "Ptprc", "Cdh1", "Mki67"))

#add meta data for clusters into lineages
DimPlot(integrate, label =T)
integrate@meta.data$lineage <- integrate@meta.data$seurat_clusters
integrate@meta.data$lineage <- plyr::mapvalues(x= integrate@meta.data$lineage,
                                               from=c("0","1","2","3","4","5","6","7","8","9","10","11","12","13","14" , "15", "16", "17", "18", "19", "20", "21", "22", "23", "24", "25", "26","27"),
                                               to= c("Cdh1","Cdh1", "Cdh5", "Col3a1", "Cdh1", "Ptprc","Cdh1", "Col3a1", "Cdh1", "Ptprc", "Col3a1", "Cdh5", "Ptprc", "Ptprc", "Col3a1", "Cdh5", "Cdh1", "Ptprc", "Cdh5", "Ptprc", "Cdh1", "Col3a1", "Col3a1", "Cdh5", "Col3a1", "Col3a1", "Cdh5", "Cdh5"))
#confirm new names
Idents(object = integrate) <- "lineage"
DimPlot(integrate, label = T, group.by = "lineage")
#save file
saveRDS(integrate, file ="/Path/to/file/SYT_all_lineages.rds")

#subset epithelial cell types, confirm subset, and repeat clustering
integrate_epi <- subset(integrate, idents=c("Cdh1"))
DimPlot(integrate_epi, label =T)
integrate_epi <- ScaleData(integrate_epi) %>% RunPCA(npcs = 30) %>% RunUMAP(dims=1:30) %>% FindNeighbors() %>% FindClusters()
DimPlot(integrate_epi, label =T)

#check control vs mutant
DimPlot(integrate_epi, group.by = "seurat_clusters", split.by = "geno")
DimPlot(integrate_epi, group.by = "seurat_clusters", split.by = "geno", label = T)

#recheck for doublets lineage genes that should not be present
FeaturePlot(integrate_epi, features = c("Col3a1", "Cdh5", "Ptprc", "Cdh1", "Mki67"))
#check epithelial markers
FeaturePlot(integrate_epi, features = c("Spock2", "Lamp3", "Scgb1a1", "Foxj1"), cols = c('lightgrey','red'))


#add metadata renaming clusters as cell types
integrate_epi@meta.data$cell_type <- integrate_epi@meta.data$seurat_clusters
integrate_epi@meta.data$cell_type <- plyr::mapvalues(x= integrate_epi@meta.data$cell_type,
                                                     from=c("0","1","2","3","4","5","6","7","8","9","10","11","12", "13"),
                                                     to= c("AT1","AT1", "AT2", "AT2", "AT2", "AT2","AT1", "AT2", "AT2", "Secretory", "AT1", "Ciliated", "AT2", "AT2" ))
#verify 
Idents (integrate_epi) <- "cell_type"
DimPlot(integrate_epi, label =T, split.by = "geno")

#Figure 5F UMAP
DimPlot(integrate_epi, label = F, split.by = "geno", group.by = "cell_type", cols = c('green4','darkmagenta','goldenrod1', 'salmon'))

Idents(integrate_epi) <- "geno"
Ctrl <- subset(integrate_epi, idents=c("ctrl"))
table(Ctrl$cell_type)
#AT1       AT2 Secretory  Ciliated 
#2040       881        72        32 


Mutant <- subset(integrate_epi, idents=c("SYT"))
table(Mutant$cell_type)
#AT1       AT2 Secretory  Ciliated 
#521      2672       165        81 

#Figure 5F Dotplot
Idents(integrate_epi) <- "cell_type"
DotPlot(integrate_epi, features = c("Spock2","Lamp3","Scgb1a1","Foxj1"), group.by = "cell_type") + coord_flip()


### saved epithelial cell object 
saveRDS(integrate_epi, file ="/Path/to/file/SYT_subsetEpi.rds")
integrate_epi <-readRDS( file ="/Path/to/file/SYT_subsetEpi.rds")

#Sup. Figure 5
FeaturePlot(integrate_epi, features = c("Spock2", "Lamp3", "Foxj1", "Scgb1a1") ,cols = c('lightgrey','red'))
FeaturePlot(integrate_epi, features = c("Ager", "Il33", "Sox9", "Mki67") ,cols = c('lightgrey','red'))


#Figure 5G
#Differential gene expression analysis AT1vAT1
Idents(integrate_epi) <-"cell_type"
integrate_AT1 <- subset(integrate_epi, idents=c("AT1"))
integrate_AT1 <- FindVariableFeatures(integrate_AT1)
Idents(integrate_AT1) <- "geno"
DEG_cAT1vmAT1   <- FindMarkers(integrate_AT1, ident.1 = c("SYT"), ident.2 = c("ctrl"), min.pct = 0.05, logfc.threshold = 0.0, test.use = "MAST", pseudocount.use = 0.001)
write.csv(DEG_cAT1vmAT1, file = "/Path/to/file/SYT_cAT1vmAT1.csv")

#Figure 5G left panel Volcanoe plot cAT1vmAT1
GNAME <- c("list", "of", "genes", "you", "want", "labeled")
EnhancedVolcano(DEG_cAT1vmAT1, lab = rownames(DEG_cAT1vmAT1), x='avg_logFC', y='p_val_adj', FCcutoff = 0.5, pCutoff = 10e-5, 
                xlim=c(-max(abs(DEG_cAT1vmAT1$avg_logFC))-.1,max(abs(DEG_cAT1vmAT1$avg_logFC))+.1), 
                gridlines.major = F, gridlines.minor = F, xlab = "ln(fold change)", colAlpha = 1,
                legendVisible = F, title=NULL, subtitle=NULL, col=c('black','black','black','red'), selectLab = GNAME, transcriptLabSize = 2.0)
#without annotation
GNAME <- 0


#Figure 5G right panel Volcanoe plot cAT2vmAT2
#Differential gene expression analysis AT2vAT2
integrate_AT2 <- subset(integrate_epi, idents=c("AT2"))
integrate_AT2 <- FindVariableFeatures(integrate_AT2)
Idents(integrate_AT2) <- "geno"
DEG_cAT2vmAT2   <- FindMarkers(integrate_AT2, ident.1 = c("SYT"), ident.2 = c("ctrl"), min.pct = 0.05, logfc.threshold = 0.0, test.use = "MAST", pseudocount.use = 0.001)
write.csv(DEG_cAT2vmAT2, file = "/Path/to/file/SYT_cAT2vmAT2.csv")

GNAME <- c("list", "of", "genes", "you", "want", "Il33")
EnhancedVolcano(DEG_cAT2vmAT2, lab = rownames(DEG_cAT2vmAT2), x='avg_logFC', y='p_val_adj', FCcutoff = 0.5, pCutoff = 10e-5, 
                xlim=c(-max(abs(DEG_cAT2vmAT2$avg_logFC))-.1,max(abs(DEG_cAT2vmAT2$avg_logFC))+.1), 
                gridlines.major = F, gridlines.minor = F, xlab = "ln(fold change)", colAlpha = 1,
                legendVisible = F, title=NULL, subtitle=NULL, col=c('black','black','black','red'), selectLab= GNAME, transcriptLabSize = 2.0)
#without annotation
GNAME <- 0


#Sup. Figure 5 
Idents(object = integrate_epi) <- "geno"
DimPlot(integrate_epi, label = T)
Mutant <- subset(integrate_epi, idents=c("SYT"))
Idents(object = Mutant) <- "cell_type"
MutantAT2 <- subset(Mutant, idents=c("AT2"))
Ctrl <- subset(integrate_epi, idents=c("ctrl"))
Idents(object = Ctrl) <- "cell_type"
DimPlot(Ctrl, label = T)
CtrlAT1 <- subset(Ctrl, idents=c("AT1"))

mAT2andcAT1 <- merge(CtrlAT1, y = MutantAT2, add.cell.ids = c("cAT1", "mAT2"), project = "SYTEnVol")
mAT2andcAT1 <- FindVariableFeatures(mAT2andcAT1)
mAT2andcAT1 <- ScaleData(mAT2andcAT1) %>% RunPCA(npcs = 30) %>% RunUMAP(dims=1:30) %>% FindNeighbors() %>% FindClusters()
DimPlot(mAT2andcAT1, label = T, group.by = "cell_type")
Idents(mAT2andcAT1) <- "cell_type"
saveRDS(mAT2andcAT1, file = "/Path/to/file/SYT_cAT1mAT2.rds")
DEG_cAT1vmAT2   <- FindMarkers(mAT2andcAT1, ident.1 = c("AT2"), ident.2 = c("AT1"), min.pct = 0.05, logfc.threshold = 0.0, test.use = "MAST", pseudocount.use = 0.001)
write.csv(DEG_cAT1vmAT2, file = "/Path/to/file/SYT_mAT2vcAT1.csv")

#Volvanoe plot cAT1vmAT2
GNAME <- c("list", "of", "genes", "you", "want", "labeled")
EnhancedVolcano(DEG_cAT1vmAT2, lab = rownames(DEG_cAT1vmAT2), x='avg_logFC', y='p_val_adj', FCcutoff = 0.5, pCutoff = 10e-5, 
                xlim=c(-max(abs(DEG_cAT1vmAT2$avg_logFC))-.1,max(abs(DEG_cAT1vmAT2$avg_logFC))+.1), 
                gridlines.major = F, gridlines.minor = F, xlab = "ln(fold change)", colAlpha = 1,
                legendVisible = F, title=NULL, subtitle=NULL, col=c('black','black','black','red'), selectLab = GNAME, transcriptLabSize = 2)
#without annotation
GNAME <- 0


#subset AT1 and AT2 for monocle 
Idents(integrate_epi) <-"cell_type"
integrate_AT1AT2 <- subset(integrate_epi, idents=c("AT1", "AT2"))
integrate_AT1AT2 <- FindVariableFeatures(integrate_AT1AT2)
#save object for entry into monocle
saveRDS(integrate_AT1AT2, file ="/Path/to/file/SYT_AT1AT2.rds")


####add module score to seurat object for genes associated with increased NKX2-1 binding and genes associated with decreased NKX2-1 binding
#####compare to binding add module score to just AT1 and AT2 cells
integrate_AT1AT2 <- readRDS( file ="/Path/to/file/SYT_AT1AT2.rds")

#As we can only add a score for genes detected in scRNA-seq we want to export a list of all genes to cross refernce with ChIP-seq annotated genes
allgenesaccountedfor <- as.data.frame(integrate_AT1AT2@assays[["RNA"]]@counts@Dimnames[[1]])
write.csv(allgenesaccountedfor, file = "/Path/to/file/SYT_allgenesdetected.csv")
genestocheck  <- as.data.frame(integrate_AT1AT2@assays$RNA@var.features)
write.csv(genestocheck, file = "/Path/to/file/SYT_variablegenes.csv")

#After cross referencing the list of genes in the ChIP-seq annotation top 20% of increased NKX2-1 binding peaks associated genes and top 20% of decreased NKX2-1 binding peaks associated genes we want to add them to the seurat object's variable features to reference
allgenesforanalysis <- c('S100a9',	'Scgb3a2',	'BC100530',	'Retnlg',	'Scgb1a1',	'Stfa1',	'Spp1',	'Stfa2',	'Gm5483',	'Igkc',	'Stfa2l1',	'Ngp',	'Stfa3',	'Cxcl2',	'Ccl21a',	'Camp',	'Ltf',	'C1qb',	'Wfdc17',	'C1qa',	'Fabp4',	'Dcn',	'Ccl4',	'Upk3b',	'S100a8',	'Pf4',	'Krt15',	'Cd74',	'Acta2',	'Actc1',	'Dynlrb2',	'Olfm4',	'C1qc',	'Il1b',	'H2-Aa',	'2010005H15Rik',	'Ptn',	'H2-Ab1',	'Saa3',	'Wfdc21',	'Meig1',	'Myl4',	'Myl7',	'Tnnt2',	'Tagln',	'Upk3a',	'Ccl3',	'Tnni3',	'H2-Eb1',	'Tppp3',	'Cxcl12',	'Ccdc153',	'Ccl12',	'Ccl5',	'Hba-a1',	'Myh6',	'Ckm',	'Marco',	'Hp',	'Ttn',	'Ccl2',	'Tnnc1',	'Hbb-bs',	'Car4',	'Chil3',	'Igfbp5',	'Ccl6',	'Sln',	'Plac8',	'Actg2',	'Rsph1',	'Tnni1',	'Gkn2',	'Myl9',	'Hbb-bt',	
                         'Cbr2',	'Cox6a2',	'Lypd2',	'H19',	'Tpm2',	'Mlf1',	'AU040972',	'Sostdc1',	'Csrp3',	'1110017D15Rik',	'Cyp2f2',	'Mmrn1',	'Cpa3',	'3300002A11Rik',	'Fabp3',	'Fam183b',	'Lum',	'Cd79a',	'Higd1b',	'Ccl9',	'Lrrc23',	'Ighm',	'Sfrp2',	'Ankrd1',	'Cxcl10',	'Rgs1',	'Gzma',	'Mcpt8',	'Postn',	'Lyz1',	'Col1a1',	'1700012B09Rik',	'Riiad1',	'Ube2c',	'Ly6d',	'Igfbp2',	'Iglc1',	'Deup1',	'Ifitm6',	'1500015O10Rik',	'Nppa',	'Itm2a',	'Ccl17',	'Col2a1',	'Atp6v0d2',	'Osm',	'Smpx',	'Ifitm1',	'Ptx3',	'Pifo',	'Cytl1',	'Gm12840',	'Mfap5',	'Aldh1a2',	'Des',	'Prss34',	'Actn2',	'Pgam2',	'Myh11',	'Fbln2',	'Nkg7',	'Myl1',	'Gdf15',	'Cx3cr1',	'Trbc2',	'Gucy1a1',	'Cxcl13',	'Hspb7',	'Stmn2',	'Vwf',	'Cpe',	'Clec4n',	'Sh3bgr',	'Tm4sf1',	'Fcrls',	'Col11a1',	'Timp4',	'Gm5416',	'Nov',	'Ccdc113',	'Vpreb3',	'Reg3g',	'Myoz2',	'1700024G13Rik',	'Ifi27l2a',	'Mt2',	'Trdn',	'Cd7',	'Col1a2',	
                         'Col3a1',	'Ccna1',	'Plp1',	'Tubb4b',	'Sntn',	'Col9a3',	'Cnn1',	'Mpz',	'Gpnmb',	'Tbx20',	'Asprv1',	'Pnoc',	'Lrg1',	'Ccdc17',	'Scgb3a1',	'Igfbp3',	'1700007K13Rik',	'Gal',	'Nme5',	'Mmp12',	'Foxj1',	'4933434E20Rik',	'Cd3g',	'Slpi',	'Erich2',	'Cenpf',	'Cxcl1',	'Aldh1a1',	'Apoe',	'Lcn2',	'Inhba',	'Dnah12',	'Tnf',	'Tpm1',	'Pln',	'4833427G06Rik',	'Fbxl22',	'Gm19935',	'Stab1',	'Pdzd2',	'Mybpc3',	'Ccdc39',	'Cma1',	'Il1a',	'Alas2',	'Col9a1',	'AA467197',	'Col9a2',	'Egfl8',	'Htra4',	'Rgs5',	'Mmp9',	'Calca',	'Tekt1',	'Lpl',	'Tpsb2',	'Agr2',	'Ccl7',	'Casq2',	'8430408G22Rik',	'Tctex1d4',	'Efcab10',	'Ccrl2',	'Cfap126',	'Mrc1',	'Enpp2',	'Cd14',	'Cited1',	'Birc5',	'Rpl39l',	'5430431A17Rik',	'Mmp8',	'Iglc3',	'Pclaf',	'Aif1',	'Klrd1',	'Ckb',	'Atf3',	'Aspn',	'Acan',	'Obscn',	'Klra5',	'Fth1',	'Gpm6a',	'BC051019',	'S100a4',	'Ly6a',	'Lgals7',	'Gm9946',	'Ascl1',	'Sulf1',
                         'Ebf1',	'Reln',	'Top2a',	'Cyp11a1',	'Retnla',	'1700001C02Rik',	'Ctla2a',	'Eln',	'Cd200r3',	'Nme9',	'Hist1h1b',	'1700016K19Rik',	'Eef1a2',	'Lyve1',	'Ms4a7',	'Odf3b',	'Gkn3',	'Ppbp',	'Sbspon',	'Il1rn',	'Mybphl',	'Ms4a1',	'Ms4a4c',	'Arhgdig',	'Clec4e',	'Wfdc2',	'Ptgs2',	'Spa17',	'Areg',	'Hdc',	'Capsl',	'Sost',	'Cd69',	'Eno3',	'Cxcl3',	'Pde5a',	'Hhip',	'Fbxo36',	'Mki67',	'Inmt',	'Ctsk',	'Ptprb',	'Ndufa4l2',	'Hmox1',	'Cd3d',	'Csrp2',	'Dmkn',	'Tmem212',	'Trbc1',	'Slurp1',	'Pcp4',	'Tmem107',	'Mfap4',	'Cldn5',	'Gucy1b1',	'Tph1',	'Gja4',	'Lrrc26',	'Lrrc51',	'Dnali1',	'Pon1',	'Pcdh17',	'Fgl2',	'Myb',	'Hist1h2ae',	'Col8a1',	'Col11a2',	'Zmynd10',	'Fcna',	'Fam92b',	'Sfrp1',	'Ms4a4b',	'Ms4a6b',	'Tcrg-C1',	'Pthlh',	'Lbhd2',	'Ly6g',	'Serpine1',	'Tekt2',	'Cenpa',	'A2ml1',	'Plet1',	'Cfap206',	'Fosb',	'Hspa2',	'Pdgfrb',	'Fhl2',	'Ppp1r36',	'Nupr1',	'Hilpda',	'Morn5',
                         'Cdc20',	'Slc22a1',	'Selenbp1',	'Scn7a',	'Ms4a2',	'Mt1',	'Plek',	'Ch25h',	'Nusap1',	'Agtr2',	'Fabp7',	'Dnah5',	'Cdk1',	'Gp9',	'Ly6c2',	'C1qtnf3',	'Nppc',	'Emcn',	'Hspb1',	'Tpsab1',	'Igf1',	'F13a1',	'Fam161a',	'Mzb1',	'Tnnt1',	'Htra1',	'Isg15',	'Sftpc',	'Lyz2',	'Ucma',	'Aplnr',	'Ltb',	'Wif1',	'Lgals3',	'Nrgn',	'C2cd4b',	'Drc3',	'Pcsk1',	'Synpo2l',	'Mest',	'Nppb',	'Ednrb',	'Cd163l1',	'Ear2',	'Spag6l',	'Lrrn4',	'Calcrl',	'AW112010',	'Cox4i2',	'Psd',	'Ccnb1',	'Lgals2',	'Clu',	'Tmem158',	'3110079O15Rik',	'Sftpa1',	'Sftpd',	'Mustn1',	'Mycl',	'Cdkn1c',	'Calb2',	'Ldb3',	'Cdh5',	'Il7r',	'Cfap45',	'1700007G11Rik',	'Tm4sf5',	'Ramp2',	'Ccl11',	'F7',	'Ptprcap',	'Angptl7',	'Pglyrp1',	'Hist1h2ap',	'Prc1',	'Cthrc1',	'Dkk2',	'Tuba1a',	'Clec1a',	'Gm29538',	'H2afx',	'Sox17',	'Cldn10',	'Muc5b',	'Agr3',	'Srgn',	'Hspa1a',	'1700001L19Rik',	'Slc7a11',	'Gpc3',	'H2al1b',	
                         'Ackr3',	'Pdgfra',	'Nts',	'Ift22',	'Spef2',	'Dnah6',	'Cym',	'Il1rl1',	'Gm867',	'Scd1',	'Pou3f1',	'Resp18',	'Cd93',	'Cybb',	'Adprhl1',	'Egfl7',	'Klk10',	'Tcf21',	'Trdc',	'Lrriq1',	'Hspa4l',	'Sprr1a',	'Cyp2s1',	'Cd160',	'Dapl1',	'2410004P03Rik',	'Map3k7cl',	'Agt',	'Ccna2',	'Lilr4b',	'Chia1',	'Serpinb1a',	'Tgfbi',	'Mt3',	'Rrm2',	'Fgf12',	'Il2rb',	'Thbs4',	'Dusp2',	'Fcgr2b',	'Clec14a',	'Calml4',	'Ltbp2',	'Rsph9',	'Cfap53',	'Krt14',	'Dab2',	'Penk',	'Epha5',	'Ctgf',	'Cdca8',	'Kcnrg',	'F10',	'Arg1',	'Il18',	'Ly6h',	'Itga1',	'Ms4a6c',	'Ccnb2',	'Ccno',	'Hmgb2',	'Hsp90aa1',	'Bcl2a1d',	'Treml1',	'Tcap',	'Cxcr6',	'Ak7',	'Cxcl15',	'Mgp',	'Hpgd',	'Angptl1',	'Slc40a1',	'Gpihbp1',	'Dnah10',	'Cks2',	'Klf2',	'Cfap44',	'Cd83',	'Krt17',	'Spag17',	'Cd68',	'Muc16',	'Dnah3',	'Ctss',	'Dpt',	'Mcpt2',	'6430531B16Rik',	'1110004E09Rik',	'2610028H24Rik',	'S100g',	'Gsta3',	
                         'Hba-a2',	'Myl3',	'Cfap54',	'Hspa1b',	'G0s2',	'Gjc3',	'Iqcg',	'Fst',	'Kdr',	'Shisa8',	'Pitx2',	'Tekt4',	'Myom2',	'Bcl2a1b',	'Cpa1',	'Il6',	'Tnfaip3',	'Upk1b',	'Gm13889',	'Gzmb',	'Pkhd1l1',	'Apobec2',	'Iglc2',	'Csf1r',	'Popdc2',	'4833423E24Rik',	'Mcemp1',	'Plvap',	'Tnxb',	'Hopx',	'Pecam1',	'Dnajb13',	'Cnmd',	'Cav3',	'Cd163',	'Cfap52',	'Cenpe',	'Cdkn1a',	'Dnah9',	'Hydin',	'Col5a3',	'Pla2g7',	'Cmtm5',	'Tmem100',	'Cdca3',	'Xcl1',	'Rspo1',	'Fmo2',	'Crybb1',	'Mafb',	'Lgals1',	'Ctsg',	'Fcer1g',	'Thy1',	'Nkain4',	'Sec14l3',	'Cd177',	'Nudc',	'Phgdh',	'Vpreb1',	'Cd34',	'Ctsw',	'Icam2',	'Spink2',	'Pdcd1',	'Rbp4',	'Chrm2',	'Skap1',	'Gsta4',	'Rptoros',	'Wt1',	'Dlk1',	'Pdlim3',	'Trim63',	'Krt83',	'Clec3b',	'Clec1b',	'Entpd2',	'Calcb',	'Tpx2',	'Ccr2',	'Mlip',	'Aldh1b1',	'Ly6c1',	'Cyr61',	'Sfn',	'Mns1',	'Ngb',	'C1ql3',	'Cby1',	'Chil1',	'Spaca9',	'Gas1',	'Npy',	'Ncr1',
                         'Akr1c18',	'Socs3',	'Nrk',	'Snca',	'Ppil6',	'Ecscr',	'Thbs1',	'Fam162b',	'Aqp5',	'Ccl24',	'Cd52',	'Olr1',	'Mcidas',	'Hmmr',	'Sgk1',	'Rasd1',	'Art3',	'Enkur',	'Tlr2',	'Il33',	'Matn3',	'Wdr78',	'Fam13a',	'Ogn',	'Cldn15',	'Cfap77',	'Fhad1',	'Klhdc4',	'Mapk10',	'Ager',	'Cebpe',	'Ccdc78',	'Cd79b',	'Pnliprp1',	'Cd24a',	'Plbd1',	'Egr1',	'Basp1',	'Cetn4',	'Ckmt2',	'Fbln5',	'Cyp2e1',	'Naaa',	'Acta1',	'Il4i1',	'Slc7a2',	'Dnaic2',	'Krtap17-1',	'Ccdc30',	'Ear1',	'Sfrp5',	'Trac',	'Fcer1a',	'Lst1',	'Cldn19',	'Mia',	'Ier3',	'Lox',	'Akap5',	'Cd209a',	'Edn1',	'Pcdh18',	'Coro1a',	'Lrrc10',	'Kcng2',	'Selenom',	'Col14a1',	'Ascc1',	'Ctsd',	'Pycard',	'Rarres2',	'Smc2',	'6820408C15Rik',	'Nnat',	'Tnfrsf13c',	'Cotl1',	'Moxd1',	'Sox18',	'Gm30146',	'Aqp4',	'Tyrobp',	'Gm43409',	'Ccpg1os',	'Cep126',	'Aurkb',	'Ccdc146',	'Fbln1',	'Elof1',	'Spic',	'Fndc1',	'Bex2',	'Cd3e',	'Pttg1',
                         'Dnajb1',	'Tceanc',	'Klrk1',	'Trem2',	'C6',	'Hsd11b1',	'Comp',	'Hcar2',	'Cyyr1',	'Mpeg1',	'Adh1',	'Itih5',	'Fibin',	'Csf2rb',	'C130080G10Rik',	'Pimreg',	'Spata24',	'Adamts5',	'Cd300lf',	'Plaur',	'Msr1',	'Aard',	'Stmnd1',	'Fam47e',	'Arhgdib',	'Hapln1',	'Ms4a6d',	'Chad',	'Ccdc34',	'Vcan',	'Mylk3',	'Cox7a1',	'Gp5',	'Rhox5',	'Scg5',	'Spink5',	'Efcab1',	'Tnc',	'Msln',	'Spi1',	'4932443I19Rik',	'Hes1',	'C3ar1',	'Ptgis',	'Anxa8',	'Alox5ap',	'Rrad',	'Mmp19',	'Cdkn3',	'Fam166b',	'Gda',	'Mgl2',	'Igll1',	'Prr29',	'Foxd3',	'Col5a1',	'Cytip',	'Cox8b',	'Bmp10',	'Fcgr3',	'Krtdap',	'Kif11',	'Thbs2',	'Trim55',	'Laptm5',	'Itgb2',	'Prox1',	'Tcrg-C4',	'Ifit3',	'Drc1',	'Pmvk',	'Ace',	'Wdr66',	'Wdr63',	'Olfr1369-ps1',	'Tbx18',	'Itgb8',	'Gm30211',	'Ryr2',	'Lrrc10b',	'Barx2',	'Cfap70',	'Kcnj8',	'H2-DMb1',	'Gatm',	'Cdhr3',	'Gem',	'Sirpa',	'Kcnj3',	'Lepr',	'Tmem151a',	
                         'Fbn2',	'Rsph4a',	'Hist1h2bc',	'S100a14',	'Ccdc40',	'Cfap36',	'Klrb1c',	'Ndnf',	'Mcpt4',	'Ftl1',	'Ccr9',	'Ccdc151',	'Lrrc71',	'Asb2',	'Mdk',	'Slfn1',	'Hand2',	'Axl',	'Ccr5',	'Notch3',	'Shox2',	'Iigp1',	'Tomm20l',	'Sox10',	'Folr2',	'Cfap65',	'Mcam',	'Fam216a',	'F3',	'Fabp5',	'Vim',	'Krt5',	'Rxfp1',	'Vegfd',	'Tnmd',	'Bcl2a1a',	'Emilin2',	'Racgap1',	'Psap',	'Dnah8',	'Rbp1',	'Prss35',	'Tuba1b',	'Adm',	'Ccnd1',	'1810010H24Rik',	'Gm26522',	'Clca3a2',	'Maf',	'Morn3',	'Bgn',	'Rcn3',	'Igfbp6',	'Itgam',	'Gyg',	'Cd302',	'Slfn2',	'Cst8',	'Pam',	'Macf1',	'Sptssb',	'Flt4',	'Xlr',	'Ptprr',	'Cxcl14',	'Gng11',	'Rgs2',	'Lrrc46',	'A930007I19Rik',	'Srl',	'Wdr95',	'Cd300c2',	'Kif23',	'Nabp1',	'Lrrc6',	'H2-DMa',	'Hyls1',	'Hist1h1c',	'Sprr2b',	'Cryab',	'Klrc1',	'Podn',	'Apoc1',	'Dcdc2a',	'Gdf10',	'Sftpb',	'Rsad2',	'Cst3',	'Pilra',	'Ereg',	'Gm26870',	'F630028O10Rik',
                         'Plagl1',	'Spata18',	'Aurka',	'Gabra1',	'Il17b',	'Unc79',	'Rtkn2',	'Cdh11',	'Cfap161',	'Gsto1',	'Cyp1a1',	'H1fx',	'Itpka',	'Uckl1os',	'Rasgrp3',	'Trf',	'Wnt2',	'Dpep1',	'Myzap',	'Fyb',	'Tcerg1l',	'Spc24',	'Lbh',	'Tie1',	'Gm21541',	'Cdh15',	'Klre1',	'Adam28',	'Cyp4f18',	'Gm26532',	'Ccr10',	'Ceacam10',	'Krt7',	'Abcg1',	'Saxo2',	'Ahsg',	'Adgrg6',	'Smim5',	'Phox2b',	'Mndal',	'Ppp1r14a',	'Sh2d1a',	'Col13a1',	'Mcm5',	'Slc25a4',	'Fhl1',	'Ly86',	'Gmnn',	'Kit',	'Ropn1l',	'Efhc1',	'Col5a2',	'Itgb2l',	'Sell',	'Crip1',	'Sh2d1b1',	'Mpo',	'Pi16',	'Cabcoco1',	'Tmem89',	'Car8',	'Lrrc74b',	'Gclc',	'Akap12',	'6030408B16Rik',	'Clec4a3',	'Gimap1',	'Dppa3',	'Col4a4',	'Ear6',	'Rasl11a',	'Lmo2',	'Clec4a2',	'Rac2',	'Angptl8',	'Hmcn1',	'Clec9a',	'Ltc4s',	'Ccl20',	'Gm42047',	'Arc',	'Fos',	'Btbd3',	'Ccdc162',	'Il13',	'Dok2',	'Cd209b',	'Retn',	'Gap43',	'C2',	'Plk1',	'Tacc3',
                         'Gpx3',	'Meis2',	'Krt19',	'Cks1b',	'Kcnh7',	'Tmem2',	'Barx1',	'Klk11',	'Tcim',	'Ifi204',	'Ccr1',	'Prss23',	'Fpr2',	'Gm16160',	'Lsp1',	'Ccdc187',	'Mcm3',	'4930444F02Rik',	'Gimap4',	'Cyba',	'Lmo7',	'Capg',	'Irf8',	'Ldlrad1',	'Tubb5',	'Hsph1',	'Bbof1',	'Pparg',	'1700088E04Rik',	'Clic3',	'Spata33',	'Mfap2',	'Fhit',	'Fn1',	'Pbk',	'Aebp1',	'Dusp15',	'Stc1',	'C5ar1',	'Dusp5',	'Stk33',	'Crh',	'Hc',	'Ahr',	'Cdkn2b',	'Prx',	'Ubxn10',	'Sdf2l1',	'Fcgr1',	'Aspm',	'Acp5',	'Vcam1',	'Clic5',	'Tspan1',	'Col6a6',	'Foxf1',	'Bex4',	'Cebpb',	'Fxyd1',	'Tuba4a',	'Pir',	'Tyms',	'Klhl23',	'Cfap69',	'Cald1',	'Slfn5',	'Enpp6',	'Stmn1',	'Odc1',	'Hspb3',	'Tuba1c',	'Egr2',	'Edn2',	'S100b',	'Cxcl5',	'Catip',	'Ighd',	'Lcp1',	'Mapk15',	'Ifi203',	'Map1b',	'Ets1',	'Crocc2',	'Kcnmb2',	'Trim54',	'Rrm1',	'Rab32',	'Grm6',	'Lrrc34',	'Gm34030',	'Grp',	'Pid1',	'Pih1d2',	'Spon2',	'Ccdc33',	'C3',	
                         'Clec12a',	'Clec4d',	'1700028P14Rik',	'Abcd2',	'Bphl',	'Cd44',	'Slit2',	'Atcayos',	'Vegfc',	'Fbn1',	'Nr4a1',	'Krt4',	'Pex5l',	'Gm13571',	'Calml3',	'Abca8a',	'Sla',	'Ldhb',	'Mmp2',	'Islr',	'Crabp2',	'Dync2h1',	'Ank1',	'Pld4',	'Il2ra',	'S1pr1',	'Spc25',	'Lmcd1',	'Kitl',	'Sst',	'Cox6b2',	'Gm973',	'Lmnb1',	'S100a6',	'Msc',	'Gngt2',	'Ifitm3',	'Sod2',	'Sparcl1',	'H2afz',	'Dcstamp',	'Pcolce',	'Nags',	'Bst1',	'Gpx2',	'Il1r2',	'Ckap2l',	'Fank1',	'Mdm1',	'Slc34a2',	'Ccdc74a',	'Maged2',	'Megf6',	'Ifrd1',	'Prox1os',	'Ccr7',	'Acyp1',	'Ptprc',	'Pdpn',	'Ncf4',	'Pcsk1n',	'Uts2b',	'Cnpy1',	'Ly6g6f',	'Phgr1',	'Nkx2-5',	'Ndufaf3',	'Mylk',	'B430010I23Rik',	'Ift74',	'Cfp',	'Grem1',	'Mrgpra2b',	'Myct1',	'Scn3b',	'Traf3ip1',	'Ndn',	'Rasip1',	'Tnfaip2',	'Sbsn',	'Hebp1',	'Igsf6',	'Samd5',	'Insm1',	'Snx10',	'Fam96a',	'Kctd14',	'Cd84',	'Slc16a3',	'Hey1',	'Fam81b',	'Ghrl',	'Tfec',	
                         'Klrb1b',	'Thbd',	'Apela',	'Gadd45g',	'Plin2',	'Fcgr4',	'Aprt',	'BC018473',	'Incenp',	'Lypd1',	'Fmo1',	'Efemp1',	'Cp',	'Enpep',	'Nexn',	'Tbx3',	'Cdc20b',	'Esam',	'Hist3h2ba',	'Acvrl1',	'Peg3',	'Anln',	'Smpd3',	'Gimap3',	'Ran',	'AC160336.1',	'Pi15',	'Siglech',	'Cyp4b1',	'AU020206',	'Rgcc',	'AU015791',	'Junb',	'Cetn2',	'Aldh1a7',	'Mdh1b',	'Uhrf1',	'Arl11',	'Npr3',	'Meox2',	'Heyl',	'Tmem252',	'Eng',	'Krt8',	'Pltp',	'Gm16006',	'Tmem37',	'Esco2',	'Dqx1',	'Knl1',	'Gm14226',	'Cd53',	'4930444P10Rik',	'Zfp36',	'Fxyd5',	'Napsa',	'Krt18',	'Gm21762',	'Wdr27',	'Rsph10b',	'Rapsn',	'Tnni2',	'Ffar2',	'Fbxo5',	'Susd5',	'Ms4a8a',	'Atad2',	'Nostrin',	'Col6a3',	'Masp1',	'Kif20b',	'Kif15',	'Bub1b',	'Rtl8a',	'Lif',	'Ccnd2',	'Eno1',	'Dusp1',	'Ccdc18',	'Clec5a',	'1700013F07Rik',	'Pmaip1',	'Palld',	'Lhx6',	'Rassf4',	'Lhfp',	'Egr4',	'Adh7',	'Gm47969',	'Wnt4',	'Zmynd12',	'Gimap6',	
                         'Itgb1bp2',	'Meg3',	'Gipc2',	'Thrsp',	'Flt1',	'Fpr1',	'Homer2',	'Kcna1',	'Ccdc155',	'Tctex1d2',	'Cndp1',	'BC028528',	'Wfdc1',	'Hcls1',	'Mcm6',	'P2ry14',	'Pou2f2',	'Armc4',	'Fbxl5',	'Serping1',	'Slitrk4',	'Klra2',	'Pcolce2',	'Htra3',	'Podxl',	'Foxn4',	'Col25a1',	'Jaml',	'Nme1',	'Mis18bp1',	'Cav1',	'Selplg',	'Ccdc60',	'Tcrg-C2',	'Gata2',	'Ttc21a',	'Synpo2',	'Nfkbia',	'Acod1',	'Cyp8b1',	'2310007B03Rik',	'Rnase2a',	'Oscp1',	'Ptpn6',	'Gprc5a',	'Col6a5',	'Wnt5a',	'Malt1',	'1700018F24Rik',	'Pbx4',	'Tspan7',	'Wisp2',	'Bex1',	'Lgmn',	'Rsph14',	'Mef2c',	'Rag1',	'Gypa',	'3110039M20Rik',	'Gp6',	'Pdlim4',	'Mid1ip1',	'Frat2',	'Me2',	'Mmd2',	'Csf3r',	'Cybrd1',	'Madcam1',	'Serpinf1',	'Plk3',	'Cdh2',	'Igfbp4',	'Spock3',	'Sirpb1c',	'Crim1',	'Dnaaf1',	'Mrap',	'Nebl',	'C230072F16Rik',	'Scg2',	'Smc4',	'Fam216b',	'1700029J07Rik',	'Dcxr',	'Pamr1',	'Slc36a2',	'Gjc1',	'Pdgfb',	'Mfge8',	
                         'Serpinb6b',	'Gm11697',	'2200002D01Rik',	'44078',	'Fgfbp1',	'Necab2',	'Apob',	'Lrrtm1',	'Cldn4',	'Bves',	'Dapk2',	'Cldn11',	'Ncf2',	'Capn6',	'Gpr182',	'Cd33',	'Nr4a2',	'Cdo1',	'Gm33489',	'Fcrla',	'Col4a3',	'D930007P13Rik',	'Upk1a',	'Ttc25',	'Ttyh1',	'A930003A15Rik',	'Mgst1',	'Bst2',	'Serpina3g',	'Rp1',	'Nhp2',	'Fxyd3',	'Stmn3',	'Col12a1',	'Tmem47',	'Trp63',	'Pde4b',	'Cd4',	'Pvalb',	'Gsr',	'Lat2',	'Hoxd10',	'Ak9',	'Myo1f',	'Abi3bp',	'3830403N18Rik',	'Nfkbid',	'Car7',	'44083',	'Crispld1',	'Arhgap30',	'Bin2',	'Epsti1',	'Ccdc189',	'Lgi1',	'Hexb',	'Cd5l',	'Pla2g1b',	'Itgb7',	'Cavin2',	'Srm',	'Jpt1',	'Pcdh7',	'Tex26',	'Zfp474',	'Aox3',	'Tubb6',	'Tspan18',	'Jcad',	'Cst9',	'Scnn1g',	'Rspo3',	'Tk1',	'Sorbs2',	'Sncg',	'Arg2',	'Spock2',	'Slfn14',	'S100a10',	'Serpina3n',	'Gm15543',	'Gm19990',	'Id4',	'Smyd1',	'Csf3',	'Ces1d',	'Sphkap',	'Cd72',	'Gm17018',	'Kctd12',	'Cxcl9',	
                         'Cavin3',	'Mxd3',	'Ruvbl2',	'Lpar4',	'Gja5',	'Baz1a',	'Cap2',	'Ranbp1',	'Trp53i11',	'Piezo2',	'Sulf2',	'Cd37',	'Cdca2',	'Cfap100',	'Tsga10ip',	'Dut',	'Itgal',	'Ckmt1',	'Cfap57',	'Dbx1',	'Prtn3',	'Tnfrsf1b',	'Slc4a1',	'Iqca',	'Serpinb5',	'Ccdc170',	'Zfp36l2',	'Lig1',	'B4galnt1',	'Fcmr',	'Clec10a',	'Nrxn1',	'Olfr655',	'Apba2',	'Ctsb',	'Cck',	'Onecut2',	'Slc26a3',	'Traf3ip2',	'Emp2',	'Cmtm7',	'Tsnaxip1',	'2210011C24Rik',	'Itgae',	'Ptprz1',	'Vwa1',	'Knstrn',	'Mcm7',	'Hells',	'Ifi209',	'Ccdc80',	'Osr2',	'Gm2a',	'Kif22',	'Ltb4r1',	'Procr',	'Aqp7',	'Ccdc184',	'Olfr78',	'Tgm3',	'Tnfsf9',	'Msrb1',	'Loxl1',	'Adipor2',	'Myo1g',	'Iqub',	'Lamp3',	'Elmod1',	'Dock10',	'Tubb2b',	'Ppp1r16b',	'AC154509.1',	'Egfl6',	'Lrp2',	'Prrx1',	'Ttll6',	'Meltf',	'Cstb',	'Lmna',	'Dach1',	'Ccdc96',	'Klf4',	'Adgre1',	'Tfcp2l1',	'Aqp1',	'Sbk2',	'Adm2',	'Thsd7a',	'Cryba4',	'Sgcg',	'Lama2',	'Insc',	'Prex2',	
                         'Prph',	'Nrp2',	'Fgf21',	'1700003E16Rik',	'Sema6a',	'Gimap5',	'Ctsh',	'Mir155hg',	'Zbp1',	'Slc16a10',	'Adam8',	'Chodl',	'Anxa1',	'Klhl4',	'Gdpd2',	'Map9',	'Myc',	'Pcna',	'Adam33',	'Adamts18',	'Ehf',	'Nlrp3',	'Nrarp',	'Cd248',	'Frem2',	'Plk4',	'Ripply3',	'Nfam1',	'Plk2',	'Clec4a1',	'9330158H04Rik',	'Cxcr4',	'Lpcat1',	'Cyth4',	'Olfml3',	'Wnt6',	'Tceal6',	'Cpz',	'Acot7',	'Kcnk3',	'Sox2',	'Gmfg',	'Ncam1',	'Cd300a',	'Adamts2',	'Fermt3',	'A730046J19Rik',	'Slfn4',	'Npc2',	'Rgs18',	'Cspg4',	'Blvra',	'Tek',	'Gas6',	'Gxylt2',	'Lta',	'Ptcra',	'Arhgef15',	'Ttc16',	'Mgll',	'Mirg',	'Atf5',	'Slc38a5',	'Plpp3',	'Cth',	'Emid1',	'Fam111a',	'Gpc6',	'Il4',	'Kcnj5',	'Crispld2',	'Acox2',	'Cfap157',	'Svep1',	'Cxcr2',	'Tmem163',	'B9d1',	'Spag16',	'Kpna2',	'Sh2d2a',	'Mgst2',	'Lilrb4a',	'Fam46c',	'A330069E16Rik',	'Tm6sf1',	'Epha3',	'Ckap2',	'Lockd',	'Wfdc13',	'Asf1b',	'Gm14964',	
                         'S100a16',	'Rbp2',	'Plgrkt',	'Col18a1',	'Fxyd6',	'44077',	'Efnb2',	'Dpysl3',	'Sall1',	'Serpinb11',	'Ifit1',	'Cd200',	'Bicc1',	'Vwa3a',	'AI593442',	'Cd48',	'Wisp1',	'Sema3d',	'Samsn1',	'Pappa2',	'Oas1e',	'Coro2a',	'Tacstd2',	'Pirb',	'Errfi1',	'Cela1',	'Ntrk2',	'Katnal2',	'Fsip1',	'Oit1',	'Fmod',	'Slc9a3r2',	'Slc39a2',	'Ptprd',	'Slc15a3',	'Ctxn1',	'Alpk2',	'Ap1s2',	'Adgrl4',	'Rnase6',	'Atp2a2',	'Tnn',	'Map4k1',	'Cd164l2',	'Ajap1',	'Trdv4',	'Vegfa',	'Chgb',	'Ntm',	'AI646519',	'Gm29773',	'Syce2',	'Krtap28-13',	'Tspan13',	'Adamts1',	'Serpinb2',	'Ptp4a3',	'Gapdh',	'Gfra2',	'Alx1',	'Nefl',	'Jun',	'Aldh3b1',	'Bok',	'Ffar4',	'Lcp2',	'Klk1',	'Ncapg',	'Crabp1',	'5330417C22Rik',	'Ccdc65',	'Phlda3',	'Arhgap42',	'Dnm3os',	'Flrt2',	'Adgrl2',	'Rbm24',	'Il17ra',	'1700097N02Rik',	'Tnfaip6',	'Ppp1r14c',	'Sox3',	'AU022793',	'Gpr37l1',	'Tmem213',	'Fscn1',	'Klhdc9',	'Gm10260',	'Rnase4',	
                         'Spink4',	'AC140186.1',	'Sptlc3',	'Chrdl1',	'Sema3a',	'Ier2',	'Milr1',	'Frem1',	'Tshb',	'Lmod2',	'Tbx5',	'Gpd1',	'St8sia6',	'Jag1',	'Lbp',	'Ifi207',	'Timp3',	'Spon1',	'Kif2c',	'Tbx21',	'Fam105a',	'Tcf7l2',	'Medag',	'Ankrd66',	'Unc93b1',	'Cenpw',	'Parm1',	'Vsnl1',	'Ccne2',	'Defb1',	'Igfbp7',	'Tpsg1',	'Ldb2',	'Ifi202b',	'Apbb1ip',	'Tgoln1',	'Mymx',	'Apln',	'Olfr558',	'Cryba2',	'Cilp',	'Npl',	'Gm13580',	'Cnn3',	'Exosc2',	'Arhgap18',	'Myh14',	'Adamtsl2',	'Myo18b',	'A330084C13Rik',	'Rgs22',	'Rag2',	'Maats1',	'Ncapd2',	'Trem3',	'Grap',	'Foxd1',	'Fgf5',	'Dlx4',	'Emp3',	'Ednra',	'Cd19',	'Tchh',	'Bub1',	'Nek5',	'Gm29480',	'Il15',	'Mb',	'Hspb2',	'Alkal2',	'Gpr183',	'Piwil4',	'Rassf9',	'Pgk1',	'Ang',	'Fgr',	'Lgi3',	'Gm6377',	'Drd2',	'Itga8',	'Gpr141',	'Fmnl1',	'Fut9',	'Spag8',	'Prr18',	'Tbx2',	'Fabp12',	'Ptpre',	'Kif20a',	'Uchl1',	'Ddx39',	'Lrrc36',	'Pim1',	'Pcdh1',	'Gm16755',	'Slc6a6',
                         'Klhl6',	'Dhh',	'Cycs',	'Sim2',	'Gramd2',	'Nrcam',	'Sox7',	'Nek2',	'Ulk4',	'Dydc2',	'Hspd1',	'AC149090.1',	'Cacna1s',	'Fgd5',	'Rbp7',	'Vtn',	'Nbl1',	'Ttll3',	'Taldo1',	'Slc11a1',	'Tmc5',	'Chchd6',	'Gm31659',	'Chst2',	'Bpifb1',	'Cnp',	'Pkib',	'Loxl2',	'Ltbp4',	'Gpr171',	'4631405K08Rik',	'Nid1',	'Smtnl1',	'Dpy30',	'Fam198b',	'Qpct',	'Mcm2',	'Trpc6',	'Rhoj',	'Ccdc180',	'Cd300e',	'Xirp1',	'Tnfaip8l2',	'Ndc80',	'Ldhc',	'Fkbp11',	'Arhgap9',	'Hbegf',	'Slc43a3',	'Gm16104',	'Clec2d',	'Ube2t',	'Cxcr5',	'Ddo',	'Phlda1',	'Fcor',	'Rab27b',	'Vnn1',	'Dram1',	'Dusp14',	'Rnh1',	'Creb5',	'Elf5',	'Prrx2',	'Actb',	'Tnfrsf23',	'Upb1',	'Cd38',	'Irf5',	'Dnah11',	'Gp1ba',	'Stmn4',	'Myl10',	'Cyp2a5',	'Tdrd9',	'Pdgfrl',	'Angpt1',	'Chac1',	'Runx1',	'Ooep',	'Ccdc181',	'Acoxl',	'Sgo2a',	'P2ry6',	'Fam81a',	'Wfdc3',	'Hoxa5',	'Ncoa4',	'Alox15',	'Matn1',	'Marcks',	'Olfml2b',	'Pou2af1',	'Tbx4',	'Fam49b',	
                         'Drc7',	'Limch1',	'Klri2',	'Myf5',	'Gcnt3',	'Mov10l1',	'Timp1',	'Epas1',	'N4bp3',	'Ddit3',	'Plat',	'4933406B17Rik',	'Itgb4',	'Slc35f1',	'Ttc29',	'Eno4',	'Nfkbiz',	'H2-Q1',	'Ttc36',	'Jhy',	'Kcnn4',	'Osr1',	'Rtp4',	'Slamf9',	'Ifi205',	'Ska1',	'Nkx6-1',	'Gm10851',	'Cdt1',	'1810011H11Rik',	'Stkld1',	'Tmem243',	'Plxnd1',	'Itgb6',	'Adamts17',	'Fxyd4',	'Sdc2',	'Ramp1',	'Aoc3',	'Trem1',	'Gm8369',	'F2r',	'Mxd1',	'Chst4',	'Adgre4',	'Arl4a',	'Fam20c',	'Sct',	'Bambi',	'Matn2',	'Gm26825',	'Ankrd65',	'Parl',	'Cd55',	'Abca8b',	'Asz1',	'Ccdc81',	'Akr1c14',	'Luzp2',	'Fasn',	'Smim6',	'Col15a1',	'Scn10a',	'Snap25',	'Rnf207',	'Dlec1',	'Scnn1a',	'Gusb',	'Snx7',	'Neil3',	'Colec12',	'Gm10457',	'B3gnt4',	'Gp2',	'Gm41300',	'Cyp26a1',	'Arsj',	'Tgfb1',	'Ly6f',	'Aldoa',	'Wdfy4',	'Grrp1',	'Isyna1',	'Glrx',	'Stau2',	'Fam135a',	'Lmbrd1',	'Mgat4a',	'Rpl31',	'Stat4',	'Sf3b1',	'Bzw1',	'Klf7',	'Acadl',	
                         'Tmem169',	'Mogat1',	'Serpine2',	'Cul3',	'Slc16a14',	'Cops7b',	'Sag',	'Mroh2a',	'Cops8',	'Ube2f',	'Hdac4',	'Ankmy1',	'Serpinb12',	'Slc35f5',	'Mgat5',	'Pigr',	'Plekha6',	'Snrpe',	'Csrp1',	'Atp6v1g3',	'1700025G04Rik',	'Rgs8',	'Ier5',	'Mr1',	'Gm2000',	'Soat1',	'Tor3a',	'Astn1',	'Blzf1',	'Vangl2',	'Sdccag8',	'1700016C15Rik',	'Cnst',	'Cdc42bpa',	'Coq8a',	'Ephx1',	'Trp53bp2',	'Tlr5',	'1700056E22Rik',	'Rab3gap2',	'Esrrg',	'Kcnh1',	'Traf3ip3',	'Prpf18',	'Proser2',	'Plxdc2',	'Dnajc1',	'Hnmt',	'Cdk9',	'Garnl3',	'Zbtb34',	'Pbx3',	'Crb2',	'Kif5c',	'Fmnl2',	'Acvr1',	'Grb14',	'Stk39',	'Cers6',	'Dhrs9',	'Gpr155',	'Osbpl6',	'Cwc22',	'Itgav',	'Slc43a1',	'Ptprj',	'Kbtbd4',	'Creb3l1',	'Prdm11',	'Slc1a2',	'Bmf',	'Plcb2',	'Bahd1',	'Dll4',	'Ubr1',	'Zscan29',	'Cep152',	'Slc27a2',	'Mall',	'Bcl2l11',	'Tmem230',	'Gpcpd1',	'Bfsp1',	'Nxt1',	'Cst11',	'Zfp442',	'Acss1',	'Fkbp1a',	'Fam110a',	'Slc52a3',	
                         'Srxn1',	'Sox12',	'H13',	'Bcl2l1',	'a',	'Map1lc3a',	'Pigu',	'Ncoa6',	'Dhx35',	'Gm826',	'L3mbtl1',	'Oser1',	'Pabpc1l',	'Kcns1',	'Elmo2',	'Ocstamp',	'Prex1',	'Nfatc2',	'Zfp217',	'Pmepa1',	'Cdh26',	'Adrm1',	'Ints6l',	'Gm7173',	'Pja1',	'Igbp1',	'Gjb1',	'Kcne1l',	'Zfp704',	'Gm9833',	'Fndc3b',	'Slc2a2',	'Egfem1',	'Lrrc31',	'Acad9',	'Spry1',	'Maml3',	'Mab21l1',	'Rapgef2',	'Pdgfc',	'Dchs2',	'Trim2',	'Fhdc1',	'Fbxw7',	'Arhgef2',	'Rit1',	'Ciart',	'Zfp697',	'Man1a2',	'Ptgfrn',	'Mab21l3',	'Slc22a15',	'Ampd1',	'Dclre1b',	'Dennd2d',	'Prok1',	'Cyb561d1',	'Celsr2',	'Cdc14a',	'Rtca',	'Plppr5',	'Alpk1',	'Rpl34',	'Lef1',	'Arhgef38',	'Slc39a8',	'Sh3glb1',	'Ctbs',	'St6galnac3',	'Lrrc40',	'Tox',	'Gm10604',	'Epha7',	'Bach2',	'Aco1',	'Aptx',	'Phf24',	'Fbxo10',	'Tgfbr1',	'Nr4a3',	'Abca1',	'Slc44a1',	'Ptpn3',	'D630039A03Rik',	'Zkscan16',	'Slc46a2',	'Fkbp15',	'Hdhd3',	'Zfp618',	'Col27a1',	'Pappa',
                         'Trim32',	'Mllt3',	'Oma1',	'Ssbp3',	'Tmem59',	'Ndc1',	'Lrp8',	'Nrd1',	'Pdzk1ip1',	'Ipp',	'Akr1a1',	'Med8',	'Cited4',	'Mfsd2a',	'Zc3h12a',	'Stk40',	'Gjb3',	'Trim62',	'C77080',	'Ptp4a2',	'Wasf2',	'Rpl11',	'Rap1gap',	'Ece1',	'Vwa5b1',	'Rnf186',	'Pqlc2',	'Ubr4',	'Pax7',	'Rcc2',	'Pgd',	'Slc25a33',	'Gpr153',	'Nadk',	'Armc10',	'Lhfpl3',	'Prkag2',	'Insig1',	'Cad',	'Gm9970',	'Mrpl33',	'Fosl2',	'Pisd',	'Ppargc1a',	'Stim2',	'0610040J01Rik',	'Rell1',	'Klb',	'Rbm47',	'Corin',	'Nipal1',	'Tec',	'Fryl',	'Lrrc66',	'Aasdh',	'Dck',	'Rassf6',	'Mthfd2l',	'Stbd1',	'Mrpl1',	'Bmp2k',	'Prkg2',	'Cds1',	'Aff1',	'Lrrc8b',	'Zfp326',	'Fbrsl1',	'Mn1',	'Tchp',	'Git2',	'Bicdl1',	'Taok3',	'Med13l',	'Cux2',	'Hvcn1',	'Gpn3',	'Anapc5',	'Ccdc62',	'Hip1r',	'Arl6ip4',	'Zfp664',	'Tmem132c',	'Auts2',	'Sh2b2',	'Mad1l1',	'Chst12',	'Foxk1',	'Usp42',	'Baiap2l1',	'Trrap',	'Ppp1r9a',	'Asns',	'C1galt1',	'Ica1',	'Met',	'Capza2',	
                         'Wnt16',	'Fam3c',	'Cadps2',	'Flnc',	'Plxna4',	'Slc35b4',	'Akr1b8',	'Mtpn',	'E330009J07Rik',	'Clcn1',	'Zfp282',	'Mindy4',	'Adcyap1r1',	'Pde1c',	'Mogs',	'Cyp26b1',	'D6Ertd527e',	'Rab43',	'Cnbp',	'Abtb1',	'Zxdc',	'Slc41a3',	'Iqsec1',	'Mitf',	'Rybp',	'Edem1',	'Rpusd3',	'Vgll4',	'Tsen2',	'Cand2',	'Rpl32',	'Tmem121b',	'Ntf3',	'Cracr2a',	'Prmt8',	'Tspan11',	'Magohb',	'Bcl2l14',	'Borcs5',	'Grin2b',	'H2afj',	'Rerg',	'Abcc9',	'Sox5',	'Bhlhe41',	'Lilra5',	'Tmem150b',	'Ccdc61',	'Ppp1r37',	'Lipe',	'Eid2b',	'2200002J24Rik',	'Cebpg',	'Kcnc1',	'Cyfip1',	'Selenos',	'Fam169b',	'Slco3a1',	'Rlbp1',	'Crtc3',	'Gm2115',	'Ccdc83',	'Ccdc90b',	'Tsku',	'Clpb',	'Rassf10',	'Psma1',	'Sox6',	'Gde1',	'Tmem159',	'Abca14',	'Otoa',	'Lcmt1',	'Setd1a',	'Prss8',	'Rgs10',	'Bag3',	'Plekha1',	'Fam53b',	'Glrx3',	'Mtg1',	'Eps8l2',	'Kcnq1',	'Tnfrsf26',	'Ppfia1',	'Mthfd1l',	'Sash1',	'Hivep2',	'Cited2',	'Pex7',	'Map3k5',	
                         'Map7',	'Pde7b',	'Vnn3',	'L3mbtl3',	'Themis',	'Fyn',	'Man1a',	'Mcu',	'Tbata',	'Hk1',	'Arid5b',	'A930033H14Rik',	'Ube2d1',	'Col6a1',	'Dazap1',	'Ap3d1',	'Txnrd1',	'Appl2',	'Nuak1',	'Fhl4',	'Btbd11',	'Utp20',	'Gas2l3',	'Nudt4',	'E2f7',	'Kcnc2',	'Tph2',	'Yeats4',	'Il22',	'Grip1',	'BC048403',	'Ppm1h',	'Slc16a7',	'Rdh16f2',	'Sdr9c7',	'Trappc5',	'Arglu1',	'Fam155a',	'Tnfsf13b',	'Tex29',	'Proz',	'Tfdp1',	'Atp7b',	'Thap1',	'Ppp1r3b',	'Mtus1',	'Irf2',	'Stox2',	'Galnt7',	'Psd3',	'Lzts1',	'Ell',	'Mpv17l2',	'Tom1',	'Rasd2',	'Mmaa',	'Chd9',	'Irx5',	'Gnao1',	'Amfr',	'Slc38a7',	'Zfp90',	'Tango6',	'Cyb5b',	'Zfhx3',	'Mtss1l',	'Wwox',	'Rab4a',	'Galnt2',	'Cog2',	'2810004N23Rik',	'Sprtn',	'Sipa1l2',	'Tomm20',	'Pard3',	'Dlg5',	'Rps24',	'Fam208a',	'Rft1',	'Timm23',	'Sh2d4b',	'Mbl1',	'Psmc6',	'Naa30',	'Cdh24',	'Fitm1',	'Tinf2',	'Spata13',	'Dpysl2',	'Bnip3l',	'Egr3',	'Rcbtb2',	'Nudt15',	'Lrch1',	
                         'Abcc4',	'Stk24',	'Timm8a2',	'Jrkl',	'Fat3',	'Eepd1',	'Adamts15',	'Kirrel3',	'St3gal4',	'Zfp202',	'Sorl1',	'Tbcel',	'Ccdc84',	'Scn2b',	'Tmprss4',	'Apoa1',	'Rpl10-ps3',	'Dixdc1',	'Ppp2r1b',	'Btg4',	'Ppcdc',	'Hexa',	'Uaca',	'Tle3',	'Itga11',	'Aagab',	'Smad6',	'Snapc5',	'Uchl4',	'Rab11a',	'Snx1',	'Usp3',	'Tln2',	'Vps13c',	'Rora',	'Gtf2a2',	'Lipc',	'Tcf12',	'Arpp19',	'Myo5c',	'Sh3bgrl2',	'Me1',	'Copb2',	'Ky',	'Anapc13',	'1300017J02Rik',	'Rbm15b',	'Prkar2a',	'Pfkfb4',	'Arpp21',	'Trim71',	'Itga9',	'Oxsr1',	'Ano10',	'Tmem42',	'Sacm1l',	'Ccr3',	'Smtn',	'Sec14l4',	'Ascc2',	'Kremen1',	'Tns3',	'Cobl',	'Etaa1',	'Otx1',	'Bod1',	'Kcnmb1',	'Slit3',	'Pank3',	'Adam19',	'Col23a1',	'Sar1b',	'Fstl4',	'Gm10447',	'Gm12216',	'Csf2',	'Slc36a1',	'Larp1',	'Flcn',	'Myo15',	'Tmem11',	'Map2k3',	'Epn2',	'Adora2b',	'Cenpv',	'Myocd',	'Myh13',	'Asgr2',	'Mink1',	'Spns2',	'Mnt',	'Tsr1',	'Hic1',	'Nxn',	'Proca1',
                         'Lgals9',	'Ksr1',	'Rhbdl3',	'Wfdc18',	'Ddx52',	'Aatf',	'Bcas3',	'Ints2',	'Tubd1',	'Hsf5',	'Msi2',	'Nog',	'Hlf',	'Tac4',	'Gip',	'Snx11',	'Pip4k2b',	'Arl5c',	'Cdk12',	'Klhl11',	'Acly',	'Meox1',	'Asb16',	'Dcakd',	'Hexim2',	'Map3k14',	'Arf2',	'Cyb561',	'Taco1',	'Rgs9',	'Amz2',	'Sox9',	'Sstr2',	'Slc9a3r1',	'Slc25a19',	'Grb2',	'Wbp2',	'Rnf157',	'Mxra7',	'Cep295nl',	'Cbx2',	'Chmp6',	'Bahcc1',	'Klf6',	'Larp4b',	'Chrm3',	'Heatr1',	'Stard3nl',	'Gpx6',	'D130043K22Rik',	'Cdkal1',	'A530084C06Rik',	'Ssr1',	'Txndc5',	'Slc35b3',	'Mylip',	'Atxn1',	'Phf2',	'Wnk2',	'Shc3',	'Sema4d',	'Syk',	'Auh',	'Sptlc1',	'Agtpbp1',	'Golm1',	'Adamts16',	'Slc6a3',	'Ap3b1',	'Mrps27',	'Kif2a',	'Gpbp1',	'Arl15',	'Parp8',	'Gm17509',	'Hmgcs1',	'Ncoa1',	'Sdc1',	'Laptm4a',	'Rdh14',	'Lpin1',	'Greb1',	'Pdia6',	'Itgb1bp1',	'Rnf144a',	'Cog5',	'Hbp1',	'Pik3cg',	'Ccdc71l',	'Strn3',	'Hectd1',	'Akap6',	'Vcpkmt',	'Nin',	'Abhd12b',
                         'Pygl',	'Dact1',	'Syne2',	'Max',	'Plekhh1',	'Rad51b',	'Ttc9',	'Pcnx',	'Sipa1l1',	'Riox1',	'Acot1',	'Abcd4',	'Jdp2',	'Nrde2',	'Calm1',	'Rin3',	'Chga',	'Vrk1',	'Degs2',	'Wars',	'Rtl1',	'Dio3',	'Adssl1',	'Zbtb42',	'Rapgef5',	'Ccdc152',	'Prlr',	'Ctnnd2',	'Cox6c',	'Snx31',	'Azin1',	'Oxr1',	'Slc30a8',	'Ext1',	'Fer1l6',	'Ndufb9',	'Nsmce2',	'Phf20l1',	'St3gal1',	'Ly6i',	'Top1mt',	'Ccdc166',	'Plec',	'Cyc1',	'Hsf1',	'Zfp647',	'Sstr3',	'Ankrd54',	'Rpl3',	'Mkl1',	'Desi1',	'Cyb5r3',	'Creld2',	'Lrrk2',	'Pdzrn4',	'Pus7l',	'Ano6',	'Spats2',	'Prpf40b',	'Dip2b',	'Galnt6',	'Slc4a8',	'Fignl2',	'Krt80',	'Krt1',	'Zfp740',	'Hoxc4',	'Socs1',	'Serpind1',	'Dgcr8',	'Hira',	'Tmem41a',	'Bcl6',	'Il1rap',	'Hrasls',	'Lsg1',	'Smbd1',	'Snx4',	'Kalrn',	'Sec22a',	'Gpr156',	'Gsk3b',	'Btla',	'Trat1',	'Retnlb',	'Cmss1',	'St3gal6',	'Riox2',	'Nrip1',	'Usp25',	'Mrpl39',	'Grik1',	'Scaf4',	'Clic6',	'Setd4',	'Cbr3',	'Morc3',	
                         'Chaf1b',	'Dyrk1a',	'Kcnj15',	'Erg',	'Fam3b',	'Serac1',	'Ezr',	'Pde10a',	'Pacrg',	'Igf2r',	'Afdn',	'Tbc1d24',	'Ccnf',	'Hs3st6',	'Lmf1',	'Neurl1b',	'AI413582',	'Slc26a8',	'Cpne5',	'Gm28043',	'Glp1r',	'Msh5',	'Mrps18a',	'Mrps10',	'Foxp4',	'B430306N03Rik',	'Mocs1',	'Gm7334',	'Shd',	'2410015M20Rik',	'Pja2',	'Ptprm',	'Tgif1',	'Clip4',	'Xdh',	'Ttc27',	'Pkdcc',	'Haao',	'Slc3a1',	'Gm10309',	'Calm2',	'Foxn2',	'Svil',	'Cables1',	'Npc1',	'Kctd1',	'Epb41l4a',	'Sil1',	'Spry4',	'Arhgap26',	'Nr3c1',	'Yipf5',	'Stk32a',	'Ccdc112',	'Sncaip',	'43893',	'Slc12a2',	'Smim3',	'Rps14',	'Grpel2',	'Fam210a',	'Ccdc68',	'Myo5b',	'Acaa2',	'Smad2',	'Ctdp1',	'Nfatc1',	'Tshz1',	'Zadh2',	'Fam69c',	'Timm21',	'Sptbn2',	'Ap5b1',	'Ccdc88b',	'Mark2',	'2700081O15Rik',	'Fads1',	'Sdhaf2',	'Vps37c',	'Tle4',	'Gna14',	'Rfx3',	'Asah2',	'Sgms1',	'Ankrd22',	'Stambpl1',	'Btaf1',	'Slc35g1',	'Cc2d2b',	'Ccnj',	'Tll2',	'Lcor',	'AI606181',	
                         'Pyroxd2',	'Hps1',	'Dnmbp',	'Mfsd13a',	'Actr1a',	'Trim8',	'Cyp17a1',	'Cnnm2',	'Usmg5',	'Sh3pxd2a',	'Rpl13a-ps1',	'Xpnpep1',	'Mxi1',	'Adra2a',	'Gucy2g',	'Nhlrc2',	'Atrnl1',	'Gfra1',	'Shtn1',	'Prdm14',	'Kcnb2',	'Smap1',	'Phf3',	'Arid5a',	'Tgfbrap1',	'AI597479',	'Hecw2',	'Spats2l',	'Stradb',	'Sumo1',	'Erbb4',	'Ikzf2',	'Abca12',	'Rpl37a',	'Tns1',	'Stk36',	'Epha4',	'Fam124b',	'Sh3bp4',	'Twist2',	'D2hgdh',	'Tmem185b',	'Nckap5',	'Lax1',	'Nr5a2',	'Xpr1',	'Brinp2',	'Dnm3',	'F11r',	'Catspere2',	'Lbr',	'Lyplal1',	'Tgfb2',	'Kcnk2',	'Usp6nl',	'Cubn',	'Trdmt1',	'Hacd1',	'Arhgap21',	'Ptgds',	'Setx',	'Sptan1',	'Hmcn2',	'Plpp7',	'Phf19',	'Gsn',	'Stom',	'Dennd1a',	'Ctnnd1',	'Chst1',	'Traf6',	'Ryr3',	'Rasgrp1',	'Sema6d',	'Slc24a5',	'Gm9913',	'Stard7',	'Rassf2',	'Crls1',	'Plcb4',	'Ism1',	'Flrt3',	'Rbbp9',	'Tm9sf4',	'Samhd1',	'Tgm2',	'Zmynd8',	'Ptpn1',	'Ripor3',	'Bcas1',	'Gnas',	'Zfp831',	'Ythdf1',	'Fndc11',	'Dgkk',	'Maob',	
                         'Fundc1',	'Mcts1',	'Stag2',	'Elf4',	'Phf6',	'Hprt',	'Mpp1',	'Srpx2',	'Drp2',	'Dnajc5b',	'Naaladl2',	'Mecom',	'Dcun1d1',	'Nudt6',	'Commd2',	'Arhgef26',	'Dhx36',	'Lekr1',	'Arhgef11',	'Rusc1',	'Reg4',	'Cd101',	'Igsf3',	'Mov10',	'Kcnd3',	'Agl',	'Nfkb1',	'Tspan5',	'Impad1',	'Fam110b',	'Plekhf2',	'Slc7a13',	'Tbc1d2',	'Epb41l4b',	'Akap2',	'Cdk5rap2',	'Megf9',	'Snapc3',	'Psip1',	'Adamtsl1',	'Hacd4',	'Prkaa2',	'Osbpl9',	'Dmrta2',	'Elavl4',	'Gm12830',	'Rnf220',	'Eri3',	'Artn',	'Olfr1342',	'Scmh1',	'Lin28a',	'Hspg2',	'Zfp984',	'Gpr157',	'Pusl1',	'Gatad1',	'Cyp51',	'Gtpbp10',	'Rundc3b',	'Sema3e',	'Fbxl13',	'Kmt2e',	'Smarcd3',	'Rnf32',	'Lmbr1',	'Tmem214',	'Tcf23',	'Adra2c',	'Clnk',	'Adgra3',	'Chic2',	'Gc',	'Gm9958',	'Shroom3',	'Anxa3',	'Hpse',	'Klhl8',	'Hfm1',	'Fam69a',	'Cit',	'Ccdc63',	'Rhof',	'Por',	'Tfr2',	'Pdgfa',	'Tnrc18',	'N4bp2l2',	'Slc25a13',	'Tmem168',	'Mdfic',	'Tes',	'Cped1',	'Pot1a',	'Cep41',	'Klf14',
                         'Dgki',	'Creb3l2',	'Hipk2',	'Braf',	'Tra2a',	'Osbpl3',	'Crhr2',	'Ccser1',	'Pole4',	'Aup1',	'Magi1',	'Cpne9',	'Cacna1c',	'Ninj2',	'Ddx47',	'E330021D16Rik',	'Spx',	'Etnk1',	'Amn1',	'Ube2s',	'Zfp579',	'Ehd2',	'Tmem160',	'Arhgap35',	'Samd4b',	'Capn12',	'Actn4',	'Catsperg2',	'Wtip',	'Pepd',	'Rhpn2',	'Tshz3',	'Atp10a',	'Trpm1',	'Tjp1',	'Mctp2',	'Rgma',	'Fam174b',	'St8sia2',	'Mrps11',	'Polg',	'Cemip',	'Abhd17c',	'Ilk',	'Galnt18',	'Tead1',	'Xylt1',	'Scnn1b',	'Cog7',	'Ino80e',	'Dhx32',	'Adam12',	'Dock1',	'Stk32c',	'Mob2',	'Mrpl23',	'Cnksr3',	'Ust',	'Frk',	'Popdc3',	'Fam184a',	'Msl3l2',	'Lims1',	'Sh3rf3',	'Slc29a3',	'Adamts14',	'Tet1',	'Herc4',	'Ado',	'Sbno2',	'Ttc41',	'Apaf1',	'Atp2b1',	'Otogl',	'Bbs10',	'Tspan8',	'Cnot2',	'Rap1b',	'Os9',	'Col4a1',	'Polb',	'Adam9',	'Brf2',	'Rbpms',	'Mfhas1',	'Pcm1',	'Triml2',	'Ccdc110',	'Tenm3',	'Aga',	'Crtc1',	'Haus8',	'Zfp827',	'Smad1',	'Vps35',	'Zfp423',	'Brd7',	'Snx20',
                         'Nod2',	'Cdh8',	'Atxn1l',	'Atmin',	'Sult5a1',	'Vps9d1',	'Nrp1',	'Gm49355',	'Gng2',	'Adk',	'Vdac2',	'Selenok',	'Nisch',	'Hacl1',	'Ankrd28',	'Arhgap22',	'A630023A22Rik',	'Wapl',	'Ktn1',	'Olfr750',	'Gm6904',	'Msra',	'Zfp395',	'Serp2',	'Dgkh',	'Tdrd3',	'Scel',	'Gpc5',	'Mbnl2',	'Cep57',	'Zfp426',	'Ubl5',	'Dnm2',	'Opcml',	'Arhgef12',	'Dscaml1',	'4930550C14Rik',	'Atm',	'Etfa',	'Pstpip1',	'Ptpn9',	'Sema7a',	'Ice2',	'Adam10',	'Tmod3',	'Hmgn3',	'Tfdp2',	'Pik3cb',	'Cldn18',	'Cep63',	'Tmem108',	'Pik3r4',	'Dag1',	'Ctdspl',	'Csrnp1',	'Ctnnb1',	'Trak1',	'Cdcp1',	'Sf3a1',	'Hormad2',	'Polm',	'Adcy1',	'Gm11985',	'Abca13',	'Ddc',	'Egfr',	'Ehbp1',	'Rtn4',	'Lyrm7',	'Fat2',	'Arf1',	'Wnt3a',	'Srebf1',	'Usp22',	'Rnf112',	'Myh10',	'Ndel1',	'Aipl1',	'Zzef1',	'Rab11fip4',	'Crlf3',	'Rasl10b',	'Ankfn1',	'Pdk2',	'Igf2bp1',	'Med24',	'Wipf2',	'Stat5b',	'Plekhm1',	'Gosr2',	'Helz',	'Rptor',	'Gdi2',	'Dip2c',	'Hecw1',	'Aoah',	'Ripor2',	'Gcnt2',
                         'Tbc1d7',	'Rnf182',	'Gmpr',	'Fam8a1',	'Rnf144b',	'Diras2',	'Hk3',	'Nsd1',	'Spock1',	'Kif27',	'Dapk1',	'Fancc',	'Cdc14b',	'Zfp366',	'Cd180',	'Erbin',	'BC067074',	'Fam49a',	'Cpsf3',	'Adam17',	'Sypl',	'Twistnb',	'Snx13',	'Lrrc72',	'Stxbp6',	'Prkd1',	'Nubpl',	'Arhgap5',	'Tmem229b',	'Ccdc177',	'Dpf3',	'Zfp410',	'Batf',	'Ttll5',	'Ift43',	'Gstz1',	'Sptlc2',	'Gtf2a1',	'Ttc7b',	'Ppp4r4',	'Clmn',	'Wdr25',	'Mok',	'Amn',	'Tex22',	'Ghr',	'Myo10',	'Fbxl7',	'Trio',	'Dap',	'Rims2',	'Rspo2',	'Mrpl13',	'Zhx2',	'Fam84b',	'Rbfox2',	'Grap2',	'Adsl',	'Zc3h7b',	'Pacsin2',	'Wnt7b',	'Tmem117',	'Arid2',	'Scaf11',	'Slc48a1',	'Abcc5',	'Ephb3',	'Mb21d2',	'Bdh1',	'Osbpl11',	'Adcy5',
                         'Pdia5',	'Ndufb4',	'Boc',	'Phldb2',	'Cd47',	'Tbc1d23',	'Filip1l',	'App',	'Ifngr2',	'Itsn1',	'Arid1b',	'Qk',	'Agpat4',	'Zfp13',	'Btbd9',	'Pou5f1',	'Ddr1',	'Tnfrsf21',	'Supt3',	'Mad2l1bp',	'Usp49',	'Prickle4',	'Unc5cl',	'Kcnh8',	'Lpin2',	'Memo1',	'Map4k3',	'Gtf2a1l',	'Mtpap',	'Ccny',	'Mib1',	'Garem1',	'Polr2d',	'Stard4',	'Nrep',	'Kdm3b',	'Lrrtm2',	'Sh3rf2',	'Mcc',	'Arl14epl',	'Eno1b',	'Gm4950',	'Snx2',	'Snx24',	'Csnk1g3',	'Gm4951',	'Ablim3',	'St8sia3',	'Zfp532',	'Mbd2',	'Zbtb7c',	'Ppp6r3',	'Ahnak',	'Zp1',	'Psat1',	'Trpm6',	'Fam189a2',	'Tjp2',	'Kank1',	'Cstf2t',	'Pten',	'Slc16a12',	'Pcgf5',	'Ppp1r3c',	'Ide',	'Sorbs1',	'Gm340',	'Cfap43',	'Gsto2',	'Trub1',	'Rab11fip2')

VariableFeatures(integrate_AT1AT2, assay = NULL) <- allgenesforanalysis
ScaleData(integrate_AT1AT2)
#A few gene names get mislabed in tranfer between files:  44078, 44083, 44077, 43893


#Genes annotated from sites with decreased NKX2-1 binding in the SYT mutant
SYT_NKX21_binding_down <- list( c('Prex2',	'Prdm14',	'Kcnb2',	'Gsta3',	'Smap1',	'Phf3',	'Arid5a',	'Tgfbrap1',	'AI597479',	'Cavin2',	'Hecw2',	'Spats2l',	'Stradb',	'Sumo1',	'Erbb4',	'Ikzf2',	'Abca12',	'Rpl37a',	'Igfbp2',	'Tns1',	'Stk36',	'Epha4',	'Fam124b',	'Col4a4',	'Col4a3',	'Ccl20',	'Pid1',	'Sh3bp4',	'Twist2',	'D2hgdh',	'Tmem185b',	'Marco',	'Nckap5',	'Fcmr',	'Lax1',	'Nr5a2',	'Ptprc',	'Xpr1',	'Brinp2',	'Dnm3',	'F11r',	'Catspere2',	'Lbr',	'Trp53bp2',	'Rab3gap2',	'Lyplal1',	'Tgfb2',	'Esrrg',	'Kcnk2',	'Ucma',	'Usp6nl',	'Cubn',	'Trdmt1',	'Hacd1',	'Arhgap21',	'Clic3',	'Ptgds',	'Setx',	'Sptan1',	'Hmcn2',	'Plpp7',	'Phf19',	'Gsn',	'Stom',	'Dennd1a',	'Lrp2',	'Osbpl6',	'Ctnnd1',	'Chst1',	'Prdm11',	'Traf6',	'Ryr3',	'Rasgrp1',	'Thbs1',	'Sema6d',	'Slc24a5',	'Gm9913',	
                                  'Stard7',	'Rassf2',	'Crls1',	'Plcb4',	'Ism1',	'Flrt3',	'Rbbp9',	'Cd93',	'Nxt1',	'Slc52a3',	'Tm9sf4',	'Samhd1',	'Tgm2',	'Ppp1r16b',	'Zmynd8',	'Sulf2',	'Ptpn1',	'Ripor3',	'Bcas1',	'Gnas',	'Zfp831',	'Ythdf1',	'Fndc11',	'Dgkk',	'Maob',	'Fundc1',	'Mcts1',	'Stag2',	'Elf4',	'Phf6',	'Hprt',	'Fhl1',	'Mpp1',	'Srpx2',	'Drp2',	'Dnajc5b',	'Naaladl2',	'Egfem1',	'Mecom',	'Dcun1d1',	'Nudt6',	'Mgst2',	'Commd2',	'Arhgef26',	'Dhx36',	'Lekr1',	'Rapgef2',	'Fam198b',	'Map9',	'Fbxw7',	'Arhgef11',	'Rusc1',	'Reg4',	'Phgdh',	'Cd101',	'Igsf3',	'Tshb',	'Mov10',	'Kcnd3',	'Agl',	'Nfkb1',	'Slc39a8',	'Tspan5',	'Wdr63',	'Impad1',	'Fam110b',	'Plekhf2',	'Gm10604',	'Slc7a13',	'Atp6v0d2',	'Coro2a',	'Tbc1d2',	'Abca1',	'Epb41l4b',	'Akap2',	'D630039A03Rik',	
                                  'Cdk5rap2',	'Megf9',	'Snapc3',	'Psip1',	'Adamtsl1',	'Hacd4',	'Prkaa2',	'Osbpl9',	'Dmrta2',	'Elavl4',	'Gm12830',	'Rnf220',	'Eri3',	'Artn',	'Olfr1342',	'Scmh1',	'Wasf2',	'Lin28a',	'Hspg2',	'Zfp984',	'Gpr157',	'Pusl1',	'Gatad1',	'Cyp51',	'Gtpbp10',	'Rundc3b',	'Sema3a',	'Sema3e',	'Fbxl13',	'Kmt2e',	'Smarcd3',	'Rnf32',	'Lmbr1',	'Tmem214',	'Tcf23',	'Adra2c',	'Clnk',	'Adgra3',	'Ppargc1a',	'Stim2',	'Phox2b',	'Chic2',	'Gc',	'Gm9958',	'Shroom3',	'Anxa3',	'Hpse',	'Klhl8',	'Hfm1',	'Fam69a',	'Cit',	'Med13l',	'Cux2',	'Ccdc63',	'Rhof',	'Por',	'Sh2b2',	'Tfr2',	'Pdgfa',	'Tnrc18',	'Medag',	'N4bp2l2',	'Slc25a13',	'Tmem168',	'Mdfic',	'Tes',	'Cped1',	'Pot1a',	'Cep41',	'Klf14',	'Ptn',	'Dgki',	'Creb3l2',	'Hipk2',	'Braf',	'Tra2a',	'Osbpl3',
                                  'Snx10',	'Crhr2',	'Aqp1',	'Ccser1',	'Pole4',	'Aup1',	'Mgll',	'Magi1',	'Cpne9',	'Cacna1c',	'Ninj2',	'Il17ra',	'Ddx47',	'Grin2b',	'E330021D16Rik',	'Spx',	'Etnk1',	'Sox5',	'Amn1',	'Ube2s',	'Zfp579',	'Ehd2',	'Tmem160',	'Arhgap35',	'Samd4b',	'Gmfg',	'Capn12',	'Actn4',	'Catsperg2',	'Wtip',	'Pepd',	'Rhpn2',	'Tshz3',	'Atp10a',	'Trpm1',	'Tjp1',	'Mctp2',	'Rgma',	'Fam174b',	'St8sia2',	'Slco3a1',	'Mrps11',	'Mfge8',	'Polg',	'Tm6sf1',	'Cemip',	'Abhd17c',	'Ilk',	'Lyve1',	'Galnt18',	'Tead1',	'Rassf10',	'Psma1',	'Sox6',	'Xylt1',	'Scnn1g',	'Scnn1b',	'Cog7',	'Ino80e',	'Dhx32',	'Adam12',	'Dock1',	'Stk32c',	'Eps8l2',	'Mob2',	'Mrpl23',	'Cnksr3',	'Ust',	'Frk',	'Marcks',	'Traf3ip2',	'Popdc3',	'Fam184a',	'Msl3l2',	'Lims1',	'Sh3rf3',	'Spock2',
```
