## Extended Data Figures for "Lung lineage transcription factor NKX2-1 epigenetically resolves opposing cell fates in vivo"

##### Extended Data Fig. 1: Additional validation of the *Wnt3a^Cre^* and *Sftpc^CreER^* drivers.

**a**, Genetic labeling and FACS purification of AT1 and AT2 nuclei at P7 using $Wnt3a^{Cre}$ and $Sftpc^{CreER}$, respectively. The non-AT1 cells targeted by $Wnt3a^{Cre}$ were mostly immune cells (3.3% of GFP cells), which did not express NKX2-1, and occasional AT2 cells (0.7% of GFP cells). Confocal images of immunostained lungs showing that all alveolar cells express NKX2-1 while AT2 cells are marked by LAMP3 and cuboidal E-Cadherin (ECAD). $Rosa^{Sun1GFP}$ marks the nuclear envelop of recombined cells (arrowheads; green in the diagram). All nuclei are Sytox Blue positive. For $Sftpc^{CreER}$, 250 ug tamoxifen was administrated 3 days before tissue harvest. Scale: 10 um.

**b**, Confocal images of immunostained lungs showing labeling of immune cells (CD45) by $Wnt3a^{Cre}$, which do not express NKX2-1 and thus do not interfere with NKX2-1 ChIP-seq. Scale: 10 um.

**Extended Data Figure 1:**

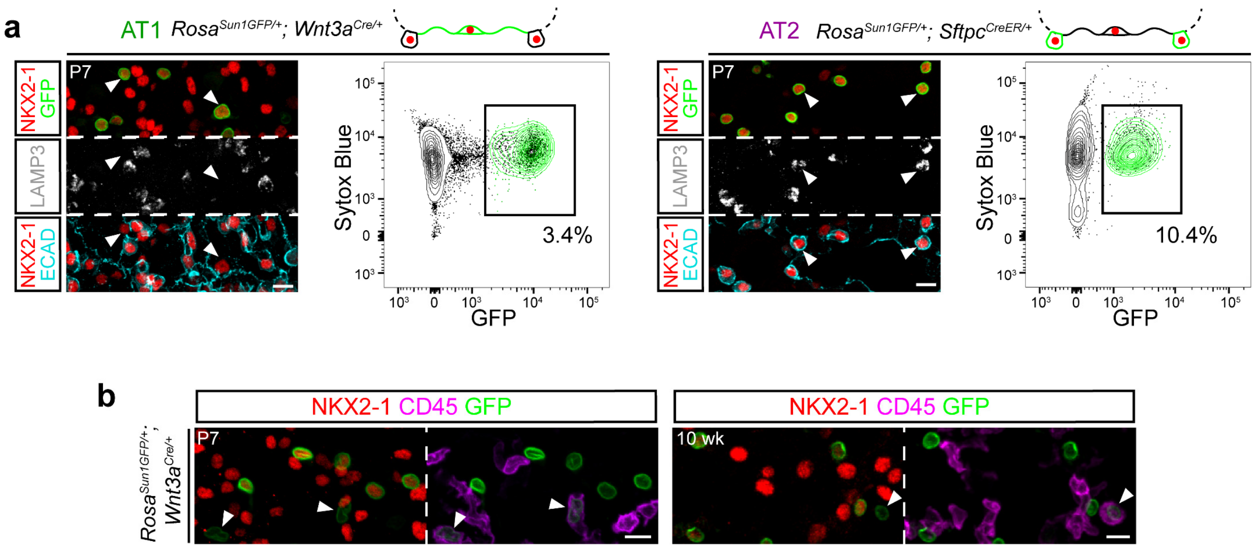

**Extended Data Fig. 2: ScATAC-seq deconvolves NKX2-1 ChIP-seq binding into lineage and housekeeping peaksets.**

**a**, ScATAC-seq UMAPs of indicated cell types (abbreviation in parenthesis for the dot plot) and lineages (abbreviation in parenthesis for **b**, **c**), as supported by the corresponding markers in the dot plot.

**b**, NKX2-1 ChIP-seq heatmaps grouped by differential NKX2-1 binding as in Fig. 1 and cross-referenced to scATAC-seq heatmaps from **a**. Cell-type-specific NKX2-1 binding sites have higher scATAC-seq signals in the corresponding cell type; NKX2-1 lineage sites have higher scATAC-seq signals in the epithelial cell types; whereas NKX2-1 housekeeping sites have comparable scATAC-seq signals across lineages. NKX2-1 lineage sites are sorted by scATAC-seq signals in Sox2 airway cells to show a subset (bottom) with less accessibility in airway cells than alveolar cells.

**c**, Example NKX2-1 binding sites and scATAC-seq signals of the AT1-specific (*Spock2*), AT2-specific (*Lamp3*), lineage (*Cdh1*), and housekeeping (*Gapdh*) sets.

#### Extended Data Figure 2:

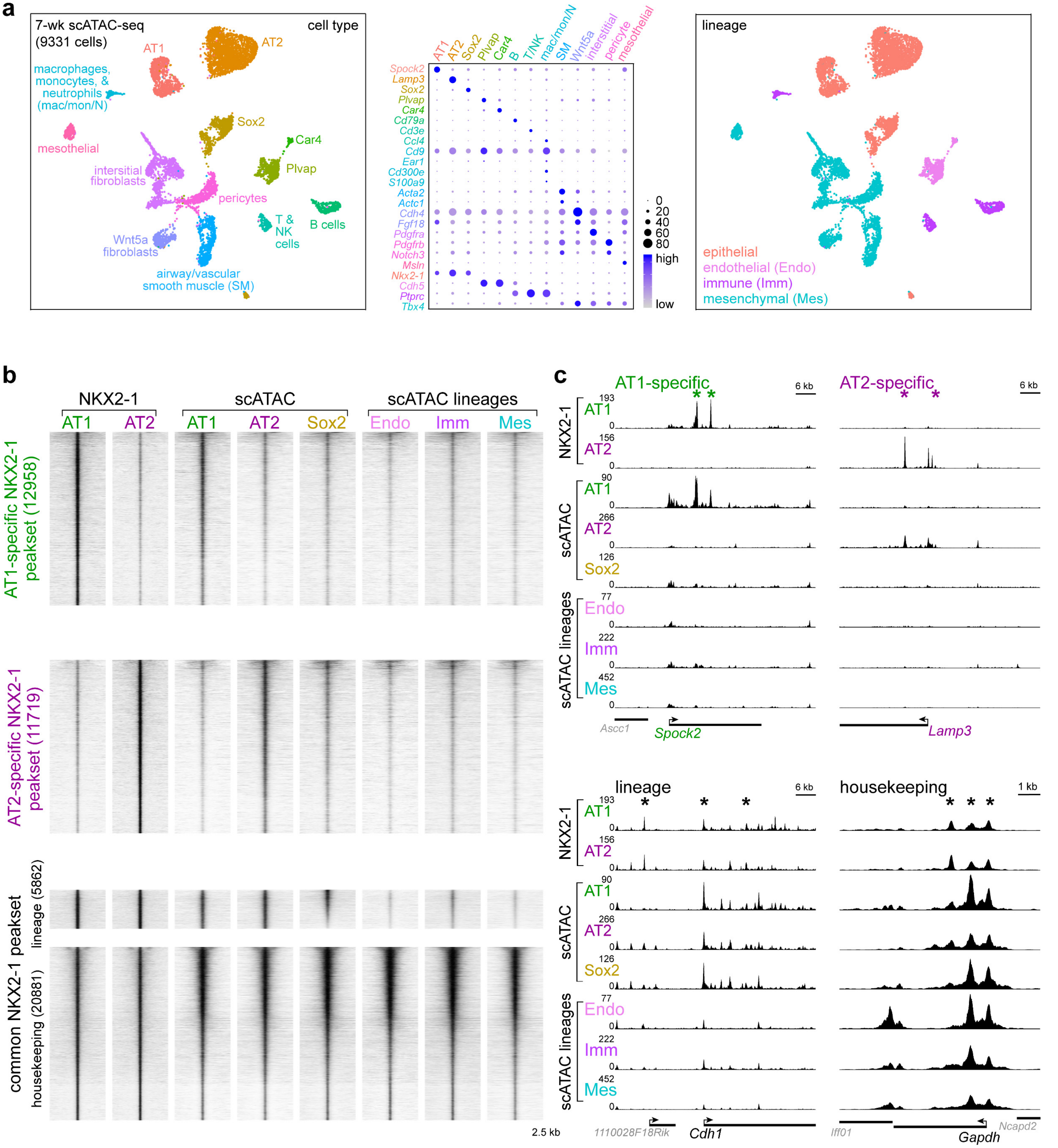

**Extended Data Fig. 3: Generation of *Rtkn2$^{CreER}$* mice and proliferation in *Nkx2-1* mutants.**

**a**, CRISPR (clustered regularly interspaced short palindromic repeats) targeting of the *Rtkn2* locus. The gRNA sequence is shown with the protospacer adjacent motif (PAM) underlined; the translation start of *Rtkn2* (ATG in red) is replaced by that of CreER. Locus-specific PCR genotyping identifies a positive founder mouse.

**b**, Confocal images of immunostained lungs showing *Rtkn2$^{CreER}$* does not target immune (CD45) or endothelial (ERG) cells. Two doses (1.5 mg) of tamoxifen (Tam) were administered for the first panel and one dose (2 mg) for the second panel. Scale: 10 um.

**c**, **d**, Confocal images of immunostained lungs of NKX2-1$^{Rtkn2}$ (c) and NKX2-1$^{Sftpc}$ (d) mutants and their littermate controls, as in Fig. 2, showing ectopic proliferative (KI67) cells 7 and 5 days after tamoxifen (Tam) administration, respectively. Proliferation is rare 5 days after tamoxifen in the NKX2-1$^{Rtkn2}$ mutant. Scale: 10 um.

**Extended Data Figure 3:**

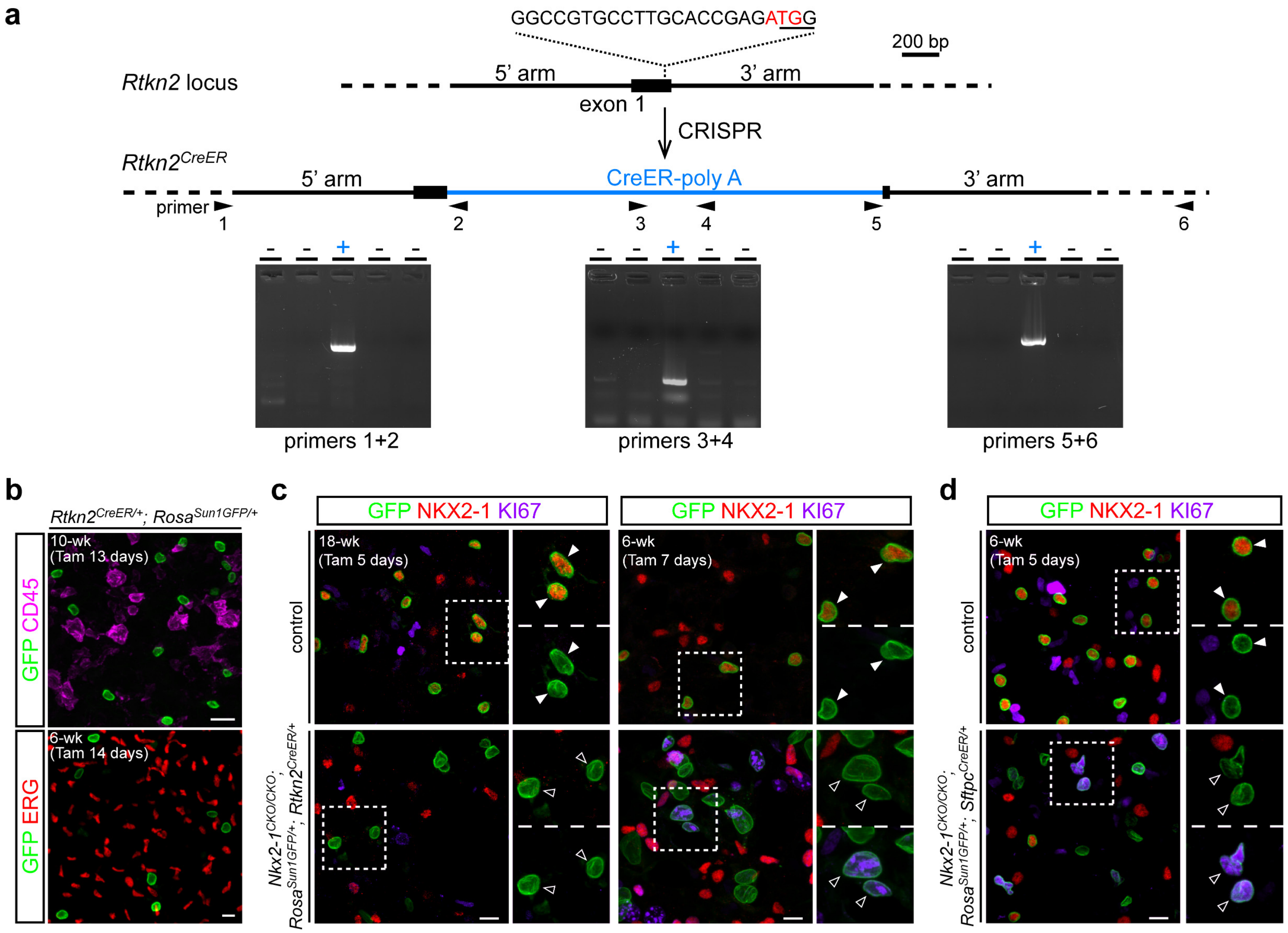

**Extended Data Fig. 4: Additional temporal analysis of NKX2-1 binding.**

**a**, NKX2-1 ChIP-seq heatmaps from E14.5 whole lungs, purified AT1 or AT2 nuclei from P7 or 10-week-old lungs, grouped as common (top) or progenitor-specific (bottom) peaksets and sorted by differential NKX2-1 binding between E14.5 lungs and the average of 10-week-old AT1 and AT2 cells. As in Fig. 3 and 4, the top (acquired or lost) and bottom (retained or reduced) 20% sites (boxed area) are used for subsequent analysis. The acquired common sites have little binding in progenitors and increase over time in both AT1 and AT2 cells, while the retained common sites have comparable binding in progenitor, AT1, and AT2 cells. The lost or reduced progenitor-specific sites have high binding in progenitors but decrease drastically or to a lesser extent, respectively, over time in both AT1 and AT2 cells.

**b**, Example NKX2-1 binding sites (asterisk) of acquired or retained common sites (*Irf2*; both types near the same gene) and lost or reduced sites (*Tinag*).

**c**, NKX2-1 binding signal of the indicated four categories in **a**, showing the kinetics of common and progenitor-specific sites as AT1 (left) versus AT2 (right) cells mature.

**d**, ScRNA-seq feature plots validating progenitor (*Sox9*), AT1 (*Spock2*, *Pdpn*, and *Hopx*), and AT2 (*Sftpb* and *Lamp3*) cells in Fig. 4a.

**e**, Seurat module scores of gene sets associated with acquired or retained common NKX2-1 sites and lost or reduced progenitor-specific sites as defined in **a**, plotted along the Monocle trajectories as in Fig. 4.

**f**, NKX2-1 binding signal of the four categories as in **e** in AT1 or AT2 cells over time, using data from **c**. Acquired common sites are associated with an increase in the module score over time, albeit more gradually in AT1 cells, while retained common sites have relatively constant module scores. Neither lost nor reduced progenitor-specific sites are associated with a decrease in the module score, possibly due to removal of proliferative cells in the Monocle trajectory analysis.

### Extended Data Figure 4:

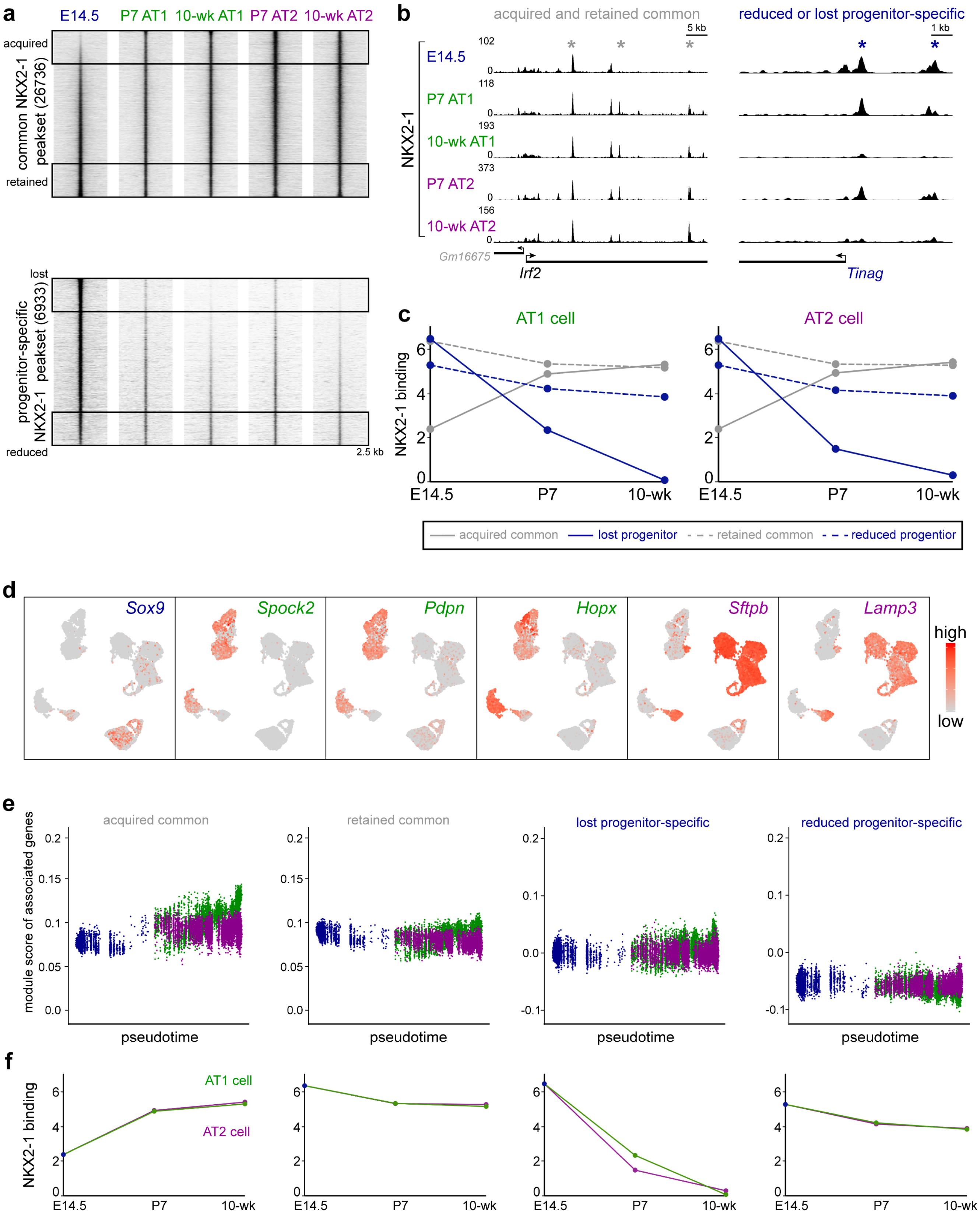

**Extended Data Fig. 5: Additional analysis of YAP/TAZ as partner factors for NKX2-1 during development.**

**a**, Violin plots of scRNA-seq data from Fig. 4 in color-coded cell types, showing enrichment of *Tead1/4* and Hippo signaling target genes *Ctgf* and *Cyr61* in AT1 cells and *Cebpa/d* in AT2 cells.

**b**, Feature plots of scRNA-seq comparison of $Y/T^{Sox9}$ mutant and littermate control lungs, as in Fig. 5. The left half (bracket; exaggerated AT2 cells) of the AT2 cell cluster in the $Y/T^{Sox9}$ mutant loses the low expression of an AT1 gene (*Ager*) but ectopically expresses *Il33* and *Scgb1a1*, the latter of which has a low level of expression in control AT2 cells.

**c**, Volcano plot of scRNA-seq data from Fig. 5 comparing AT1 cells in the control lung to AT2 cells in the $Y/T^{Sox9}$ mutant, which arise in part from AT1 cells upon *Yap/Taz* deletion, consistent with *Yap/Taz* mutant AT1 cells becoming AT2 cells.

**d**, **e**, Confocal images of immunostained lungs showing loss of AT1 markers PDPN and HOPX and wide-spread AT2 markers LAMP3 and SFTPC in the $Y/T^{Sox9}$ mutant. Scale: 10 um.

**Extended Data Figure 5:**

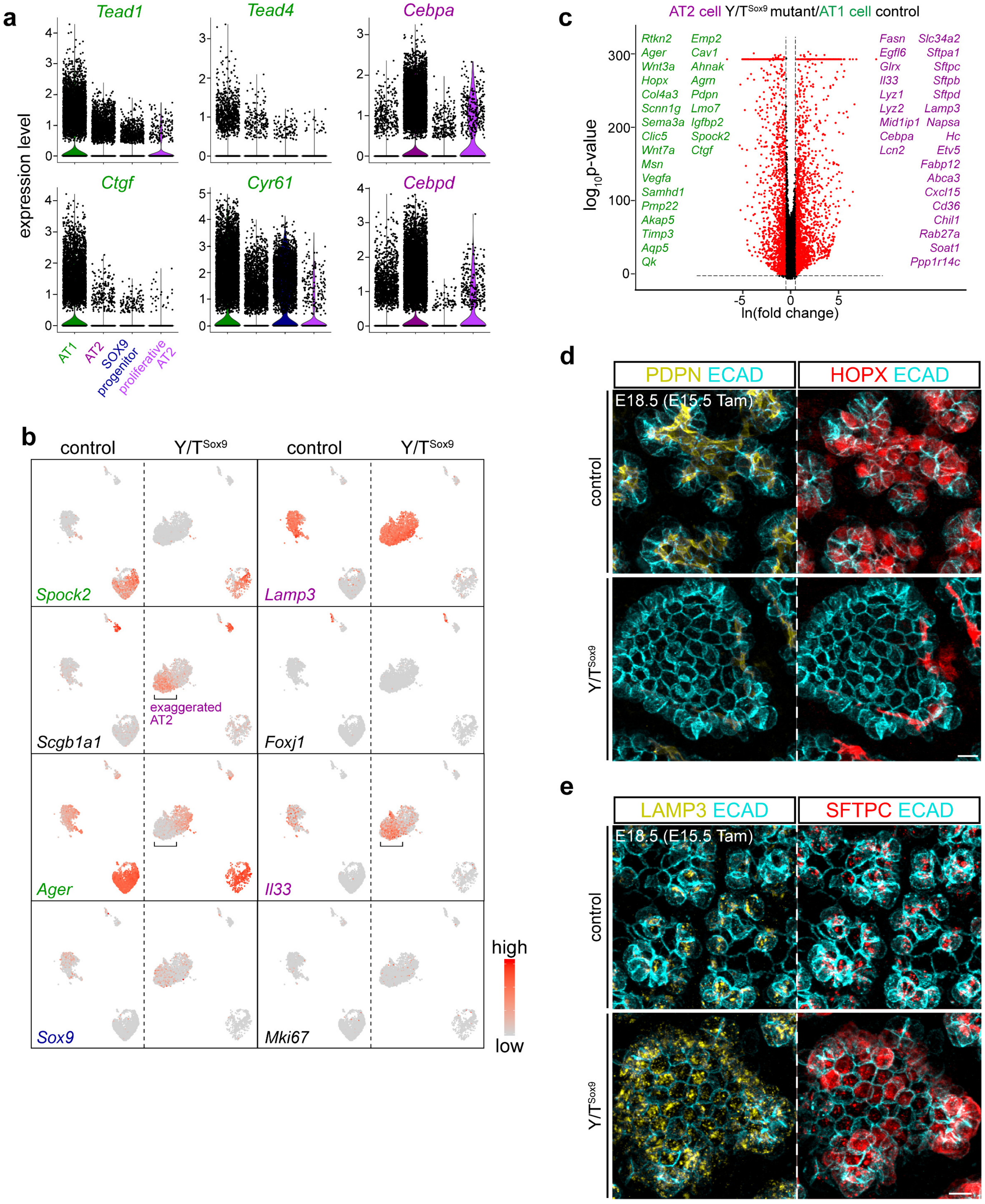

**Extended Data Fig. 6: Additional analysis of the $Y/T^{Wnt3a}$ mutant.**

**a**, Feature plots of scRNA-seq comparison of $Y/T^{Wnt3a}$ mutant and littermate control lungs, as in Fig. 6. The right half (bracket; intermediate) of the AT1 cell cluster in the $Y/T^{Wnt3a}$ mutant ectopically expresses AT2 genes *Sftpc* and *Lamp3*, as well as genes implicated in AT2-to-AT1 conversion during injury-repair such as *Sfn*.

**b**, Volcano plot of scRNA-seq data from Fig. 6 comparing AT2 cells in the control lung to AT1 cells in the $Y/T^{Wnt3a}$ mutant to confirm their expected cell type identity.

**Extended Data Figure 6:**

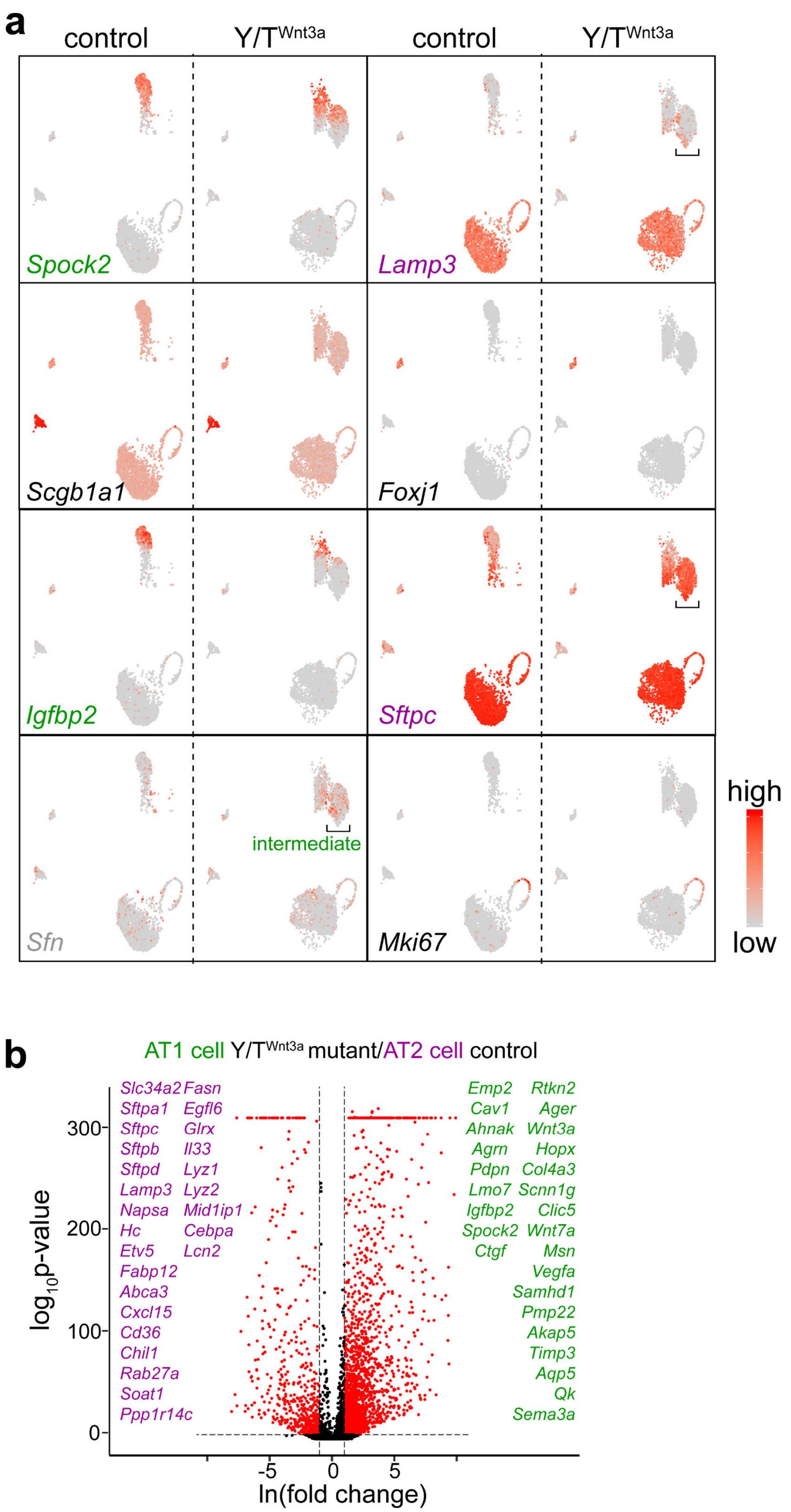

**Extended Data Fig. 7: Additional analysis of increased accessibility sites in the *Nkx2-1* mutants.**

**a**, Example gastrointestinal genes (*Tff1/2/3*) with little increase in accessibility except for a small change near *Tff2* in the NKX2-1$^{Sftpc}$ mutant (asterisk), possibly due to faster adoption of the gastrointestinal fate by AT2 cells than AT1 cells.

**b**, ScRNA-seq UMAPs of our published NKX2-1$^{Aqp5}$ mutant showing the indicated cell types and numbers.

**c**, Violin plots of the scRNA-seq data in **b**, showing increased expression of gastrointestinal genes including *Hnf4a* and *Tff1/2/3* and a transcription factor with the ELF motif (*Elf3*) in *Nkx2-1* mutant AT1 cells.

**Extended Data Figure 6:**

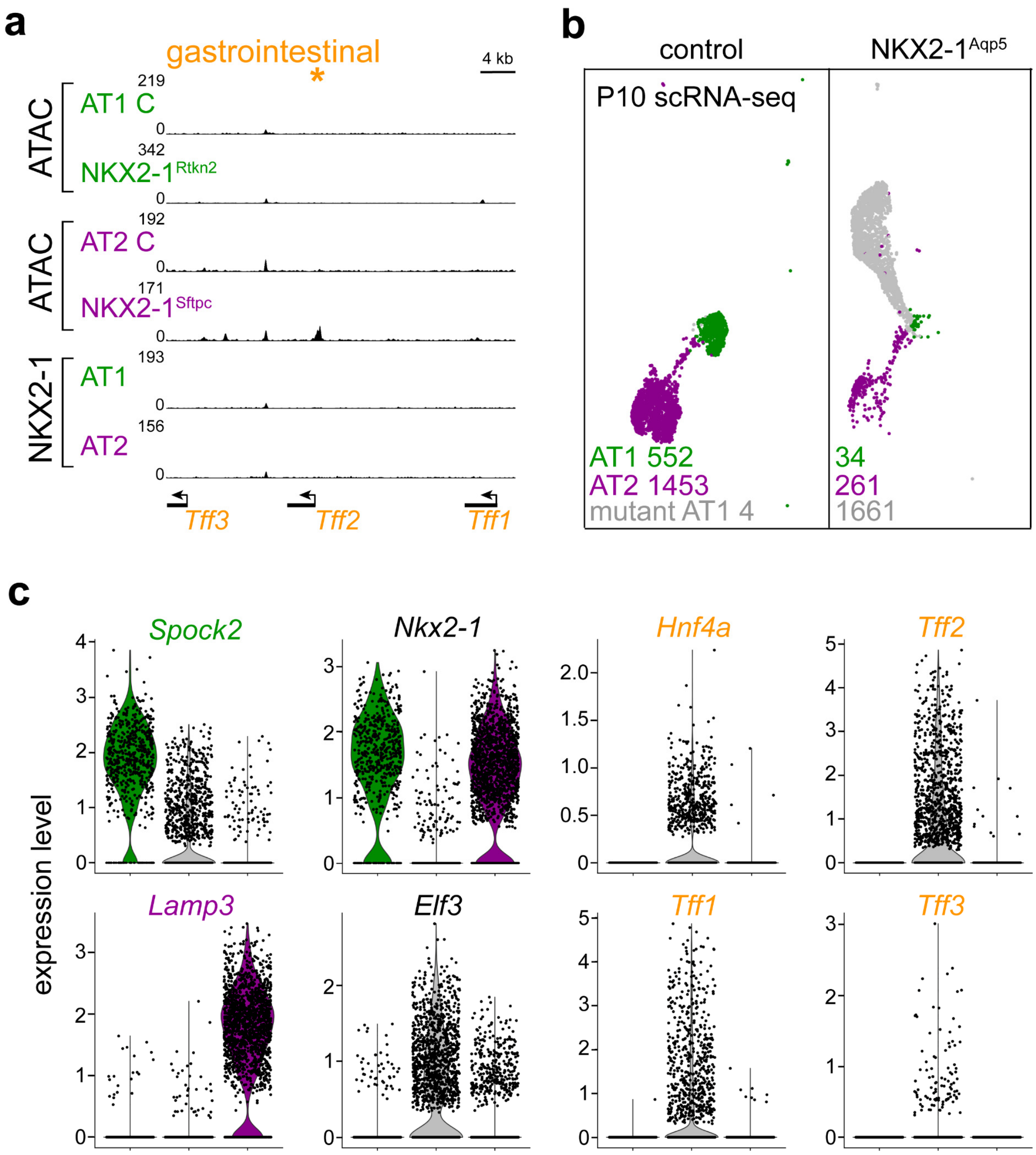

**Supplementary Table 1: Peaksets for NKX2-1 ChIP-seq heatmaps in Fig. 1, 2.**

**Supplementary Table 2: Peaksets for NKX2-1 ChIP-seq heatmaps in Fig. 3.**

**Supplementary Table 3: NKX2-1 ChIP-seq peaksets for acquired or retained AT1/AT2-specific and common categories and lost or reduced progenitor-specific categories.**

**Supplementary Table 4: Peaksets for NKX2-1 ChIP-seq heatmaps in Fig. 5.**

**Supplementary Table 5: Curated AT1 and AT2 genes and scRNA-seq analysis in Fig. 5.**

**Supplementary Table 6: Peaksets for NKX2-1 ChIP-seq heatmaps in Fig. 6.**

**Supplementary Table 7: Curated AT1 and AT2 genes and scRNA-seq analysis in Fig. 6.**

**Supplementary Table 8: Peaksets for NKX2-1$^{Rtkn2}$ ATAC-seq heatmaps in Fig. 7.**

**Supplementary Table 9: Peaksets for NKX2-1$^{Sftpc}$ ATAC-seq heatmaps in Fig. 7.**

**Supplementary Table 10: MA plot values for Fig. 7.**

**Supplemental File 1: Custom script used to generate the figures.**
